# Diversification of Nucleoside Analogues as Potent Antiviral Drug Leads

**DOI:** 10.1101/2023.06.17.545395

**Authors:** Stephan Scheeff, Yan Wang, Mao-Yun Lyu, Behzad Nasiri Ahmadabadi, Sam Chun Kit Hau, Tony K. C. Hui, Yufeng Zhang, Zhong Zuo, Renee Wan Yi Chan, Billy Wai-Lung Ng

**Affiliations:** School of Pharmacy, Faculty of Medicine, The Chinese University of Hong Kong, Hong Kong; Department of Paediatrics, Faculty of Medicine; CUHK – Hub of Paediatric Excellence; The Chinese University of Hong Kong, Hong Kong; Department of Chemistry, The Chinese University of Hong Kong, Hong Kong; Primemax Biotech Ltd., Hong Kong; Li Ka Shing Institute of Health Sciences, Faculty of Medicine, The Chinese University of Hong Kong

## Abstract

Nucleoside analogues are potent antiviral agents, but the continuous emergence of pathogenic viruses demands novel and diverse structures. Herein, we have created a diversified library of highly bioactive and non-cytotoxic nucleoside analogues featuring an unprecedented carbobicyclic core that mimics natural ribonucleoside conformation. These regio- and stereo-divergent analogues exhibit up to 16-fold greater antiviral efficacy than the FDA-approved antiviral, ribavirin. Importantly, the carbobicyclic core structure is critical for the potent antiviral efficacy, thus opening up ample opportunities for further lead optimization and mechanistic investigations.

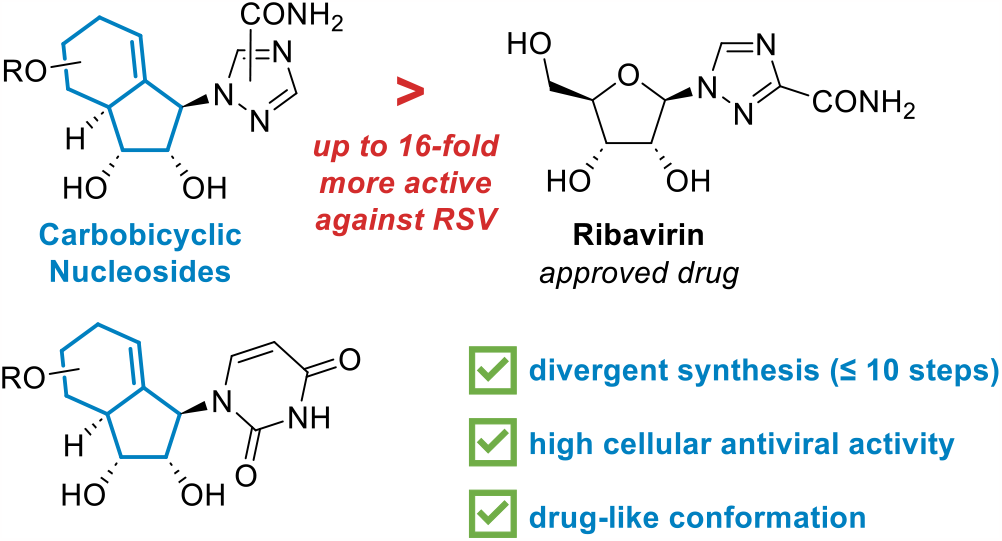

## Introduction

Nucleoside analogues are effective antiviral drugs and are used to treat many pathogenic viral infections such as HIV,^1, 2^ hepatitis^3-5^ and SARS-CoV-2^6-11^ among others.^12-16^ The continuous emergence of new viruses necessitates the urgent development of novel antivirals. Herein, we report the design and synthesis of nucleoside analogues with an unprecedented carbobicyclic core, thereby unlocking their tremendous potential as a conceptually new class of antiviral drug candidates.

Structural diversity of nucleoside analogues is key to overcoming drug resistance and enhancing combination therapies for various viruses. There are four traditional approaches for diversifying nucleoside analogues (Figure 1A): 1) the introduction of artificial nucleobases,^17^ as in ribavirin^18^ or molnupiravir^7, 19, 20^; 2) the simplification of the ribose sugar, as in adefovir^4, 21, 22^; 3) the substitution/addition of azide, fluorine, methyl or cyano groups at the sugar core, as in remdesivir^6, 23^ or zidovudine^2, 24^; 4) the modification of the ribose sugar, as in thionucleosides (i.e. lamivudine^5, 25, 26^). Substitution of the ring oxygen in the ribonucleoside with a methylene group creates an important antiviral class, the carbocyclic nucleoside analogues (Figure 1B). However, the absence of anomeric stabilization can lead to unnatural conformations, resulting in reduced biological activity.^27, 28^ Thus, carbobicyclic nucleoside analogues have been developed to lock the sugar ring in the bioactive conformation.

**Figure 1.**
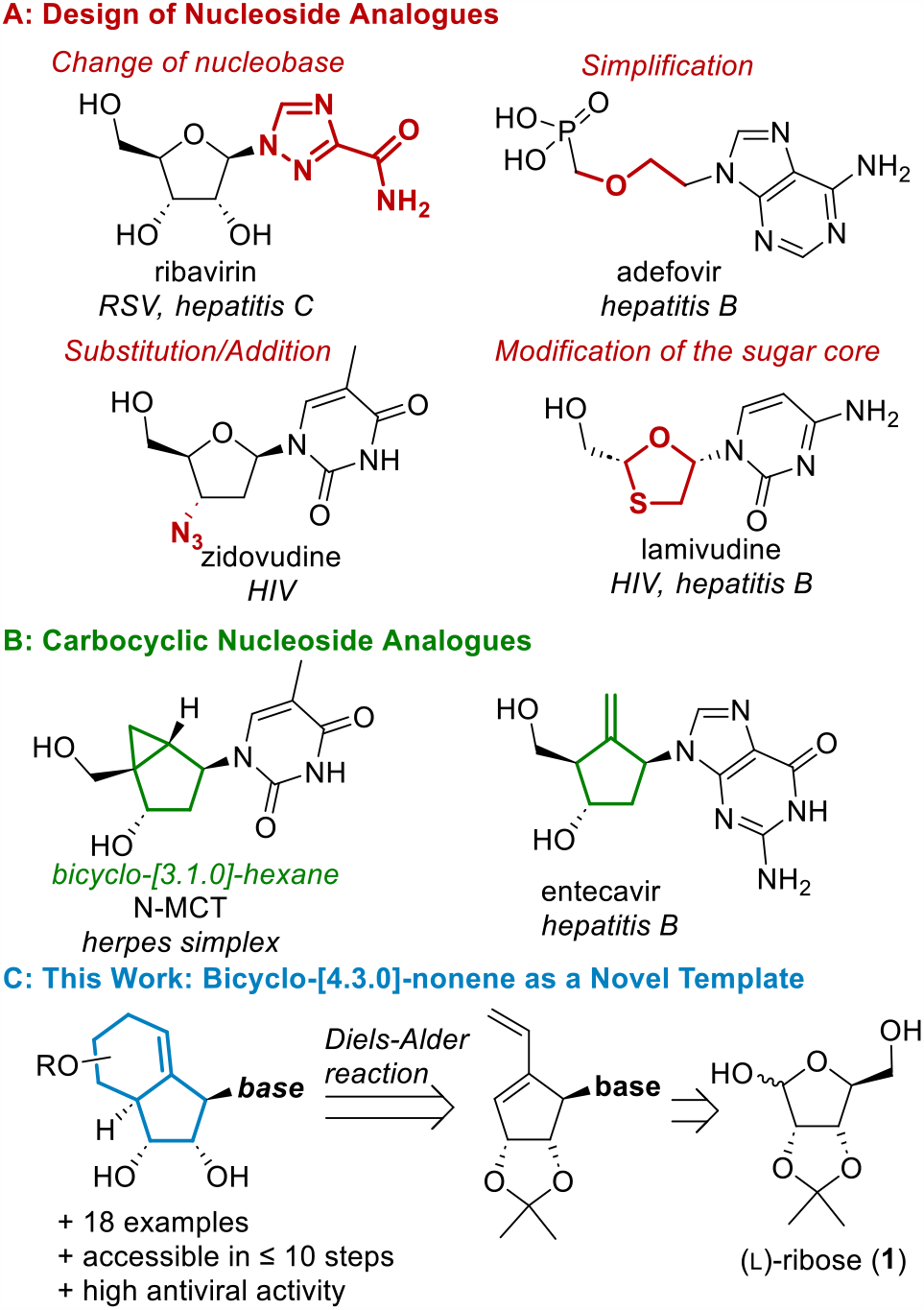
Common strategies in the design of nucleoside analogues and introduction of the bicyclo-[4.3.0]-nonene structure.

In particular, the bicyclo-[3.1.0]-hexane analogues, such as north methanocarbathymidine (N-MCT, Fig. 1B),^29, 30^ have shown superior drug-like properties^31, 32^ and promising antiviral activities. Another strategy to generate a favourable conformation employs an exocyclic double bond to replace the ribose ring-oxygen^33^ as in entecavir (Fig. 1B)^3, 34^. The limited scope for further functionalization of these analogues impedes the in-depth study of structure-activity relationships (SAR). Saturated bicyclo-[4.3.0]-nonane nucleoside analogues provide an excellent entry for functionalization and SAR study; unfortunately, these analogues do not show antiviral activity and their synthesis is lengthy and inefficient (involving 16 to 19 synthetic steps from (d)-ribose, see Fig S1).^35^

We hypothesize that an endocyclic double bond in the unsaturated bicyclo-[4.3.0]-nonene scaffold (Fig. 1C) would render them conformationally similar to the bioactive analogues entecavir and ribavirin. Thus, building upon our expertise in synthetic carbohydrate chemistry and drug design,^36-38^ we have designed a concise and divergent synthetic route towards novel bicyclo-[4.3.0]-nonene nucleoside analogues. Our step-economic synthesis features a Mitsunobu reaction and an intermolecular Diels-Alder reaction, allowing the rapid creation of a regio- and stereo-divergent library of analogues. Importantly, our pilot synthesis has enabled the discovery of an unprecedented class of antiviral drug leads, thus unlocking ample opportunities for further lead optimizations and mechanism of action studies.

## Results and Discussion

Respiratory syncytial virus (RSV) causes upper respiratory tract infections, including severe and life-threatening infections in high-risk patients.^39, 40^ In 2019, 33 million acute respiratory infections in children (0-5 years) were caused by RSV, with about 10% of cases requiring hospitalization and 100 thousand RSV-related deaths in this age group, accounting for 2% of the total mortality rate.^40^ Ribavirin is licensed for the treatment of RSV, however, the inhalation of ribavirin is costly and often inefficient.^41, 42^ Previous attempts to modify ribavirin often resulted in inactive compounds^43, 44^, so it is warranted to develop ribavirin analogues with a new core scaffold. To this end, we will couple the carbobicyclic core with the triazole nucleobase of ribavirin, with the aim to develop a more effective, non-cytotoxic, and orally bioavailable alternative to ribavirin.

**Scheme 1.**
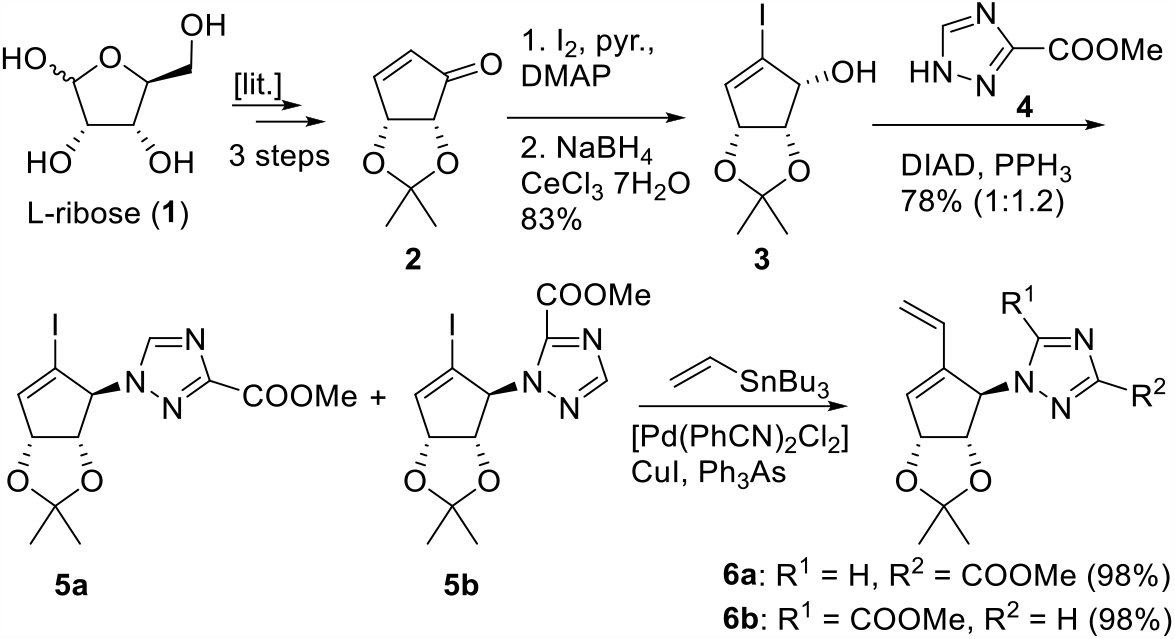
Synthesis of Diels-Alder precursor 6 from readily available sugar 1.

The Diels-Alder precursor **6** was synthesized from the commercially available (l)-ribose-derived enone **2**^45, 46^ in four steps (see Scheme 1). First, a-iodination of the enone **2** under Baylis-Hillman type conditions followed by Luche reduction gave alcohol **3** stereospecifically.^47^ Mitsunobu coupling of **3** with triazole **4** yielded 78% of regioisomers **5a** and **5b**.^18, 48-51^ With THF as the solvent, the *iso*-ribavirin type isomer **5b** was afforded as the major product (ratio **5a**:**5b** = 1:4). With DCM as the solvent, both regioisomers were obtained in a nearly equimolar ratio. They were converted to the corresponding dienes **6a** and **6b** in excellent yield using a Stille cross-coupling.^52^

In the key intermolecular Diels-Alder reaction, an activated dienophile, vinyl boronate, could allow the efficient installation of pseudo-C5′-OH functionality after oxidation of the resulting cycloaddition product (see Scheme 2).^53-55^ A facile oxidative cleavage of the boronate of the Diels-Alder adduct with NaBO3 would then liberate the desired secondary alcohol. During the Diels-Alder reaction, the dienophile approach the diene **6** from the upper side, avoiding the steric hindrance from the isopropylidene acetal group. To our delight, we successfully isolated the desired alcohol **7** directly as a single isomer, together with the other Diels-Alder adducts **8** and **9** as separable isobutyric esters after esterification (for details, see SI Fig. S2).

**Scheme 2.**
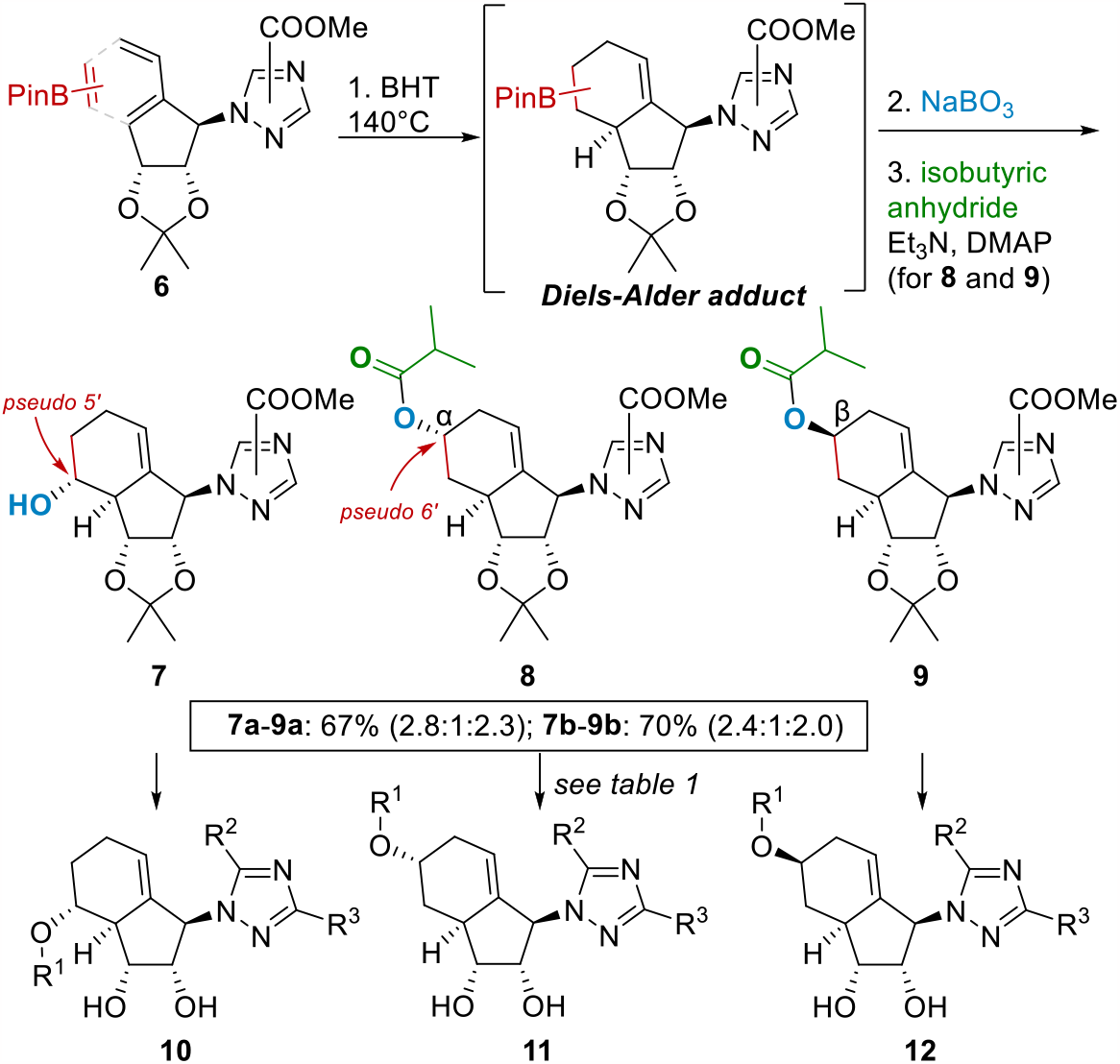
Intermolecular Diels-Alder reaction for the synthesis of analogues **10**-**12**.

The synthesis of the designed ribavirin analogues was then completed with ease (see Table 1). Firstly, the methyl ester of the triazole nucleobase was transformed into the amide^20, 48-50, 56^, and then the nucleoside analogues **10**-**12** were achieved by deprotection. In detail, from Diels-Alder adduct **7**, the ribavirin analogue **10a** and the isomer **10b** were obtained by treatment with methanolic ammonia and subsequent acidic treatment for deprotection; while an additional esterification step^57, 58^ yielded the C5′-OH esters **10c**,**d**. In the case of **8** and **9**, the amidation conditions did not lead to cleavage of the *iso*-butyric acid esters (at pseudo-C6′), and analogues **11c**,**d** and **12c**,**d** were isolated after acidic deprotection. A three-step sequence, including amidation, K2CO3-mediated ester cleavage, and acidic diol-deprotection, gave **11a**,**b** and **12a**,**b** in good yields.

**Table 1.**
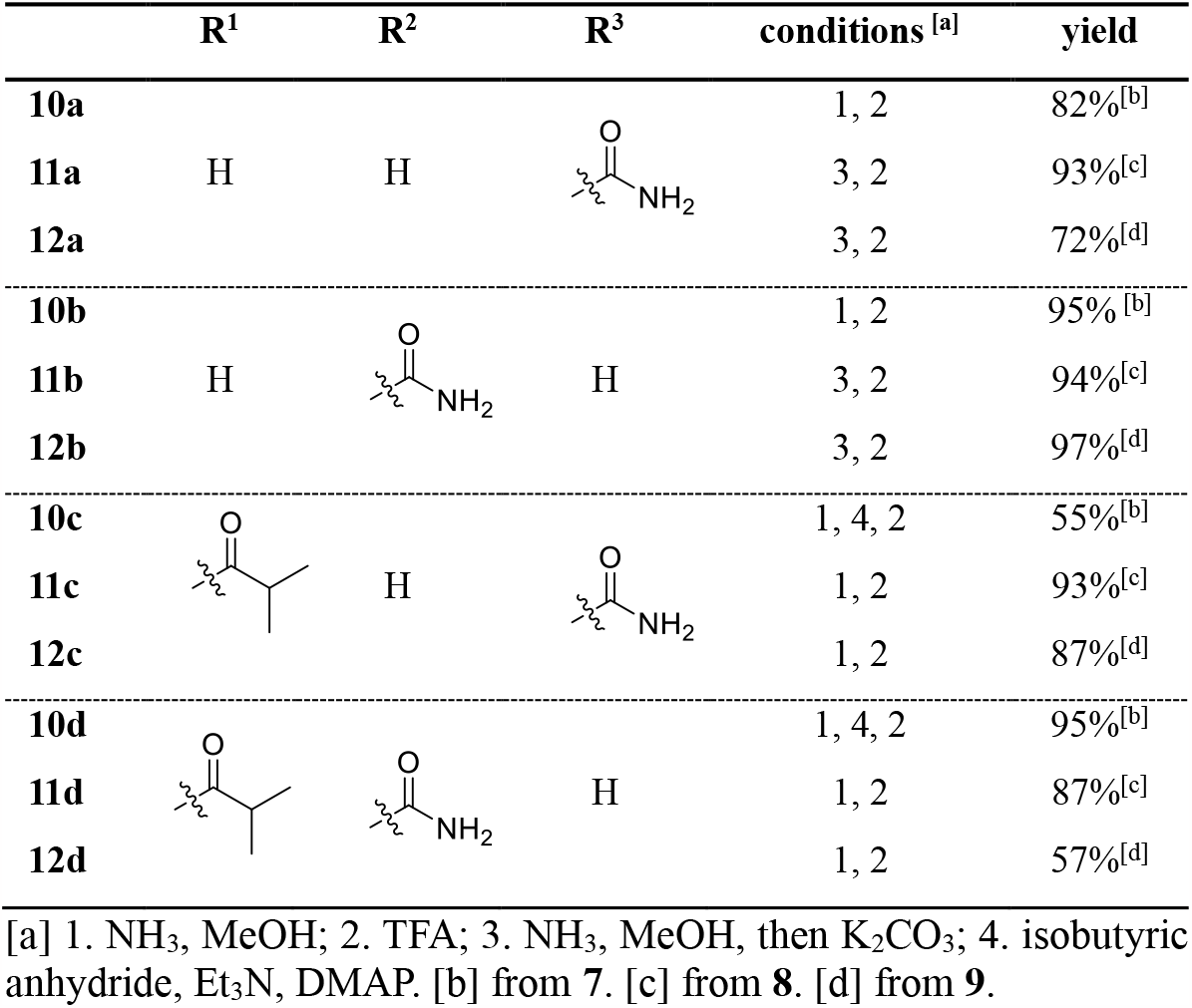
Synthesis of compounds **10-12** from **7-9**.

In short, a total of 12 ribavirin analogues were synthesized in 8-10 steps from the commercially available enone **2**. Of note, the structure of **12b** were unambiguously determined by X-ray single crystal analysis, which completely agreed with our 2D NOE-NMR assignments (see SI). The analysis validated our initial hypothesis that the bicyclic core might have a ribose-like conformation. First, the pseudorotation angle was determined as +24.0**°** which correlates with a C3’-endo (North) conformation (see Fig S3). This conformation is also observed in natural nucleosides and therefore is promising for biological activity.^28^ Second, superposition of the three-dimensional X-ray structure of **12b** with carbasugar entecavir^59, 60^ revealed an identical conformation of the carbasugar core of both structures (Fig. 2). More importantly, the comparison with authentical ribose-based ribavirin^61^ revealed a nearly identical core conformation albeit slightly twisted. Thus, by introducing the double bond into the carbobicyclic system, the nucleoside analogues adopted a favourable drug-like conformation. This is in stark contrast to the previously reported saturated bicyclo-[4.3.0]-nonane nucleoside analogues, which have an biologically inactive C1’-exo conformation.^35^

**Figure 2.**
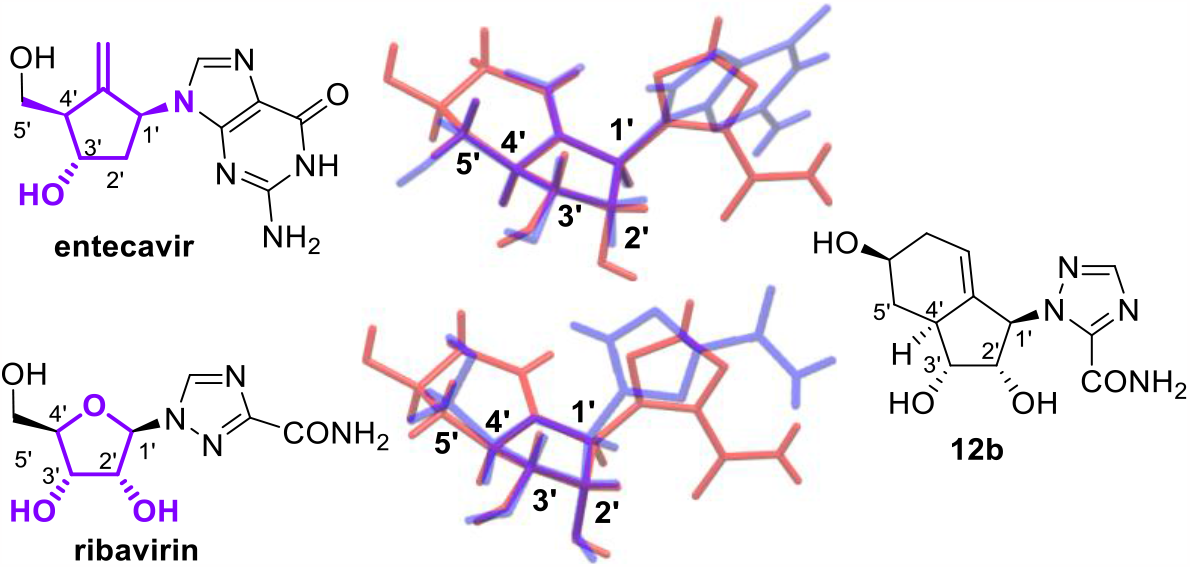
Superposition of X-ray structures of entecavir and ribavirin (both blue) with nucleoside analogue 12b (red).

With the ribavirin analogues in hands, we next evaluated their cellular activities in inhibiting RSV replication using RSV-A-infected HEp-2 cells. In a screening (c = 2 and 40 μM) of compounds **10**-**12a**,**b** all of these compounds showed promising antiviral activities (see Fig. 3A and S4, S5). Furthermore, immunostaining of RSV-infected cells after drug treatment validated the screening results (Fig. 3B left). Whereas ribavirin (2 μM) only slightly reduces RSV replication, the treatment with **12a** (2 μM) leads to an inhibition higher than 80% (Fig. 3B right). Again, at 20 μM also ribavirin showed a high antiviral activity (Fig S5). Of note, our analogues showed negligible cytotoxicity in a parallel toxicity screen using uninfected cells (see Fig. S6). The ribavirin analogues (**10-12a**) and *iso*-ribavirin (**10-12b**) analogues have comparable activity, and the presence of an ester group (**10-12c**, see Fig S7) showed no significant effect on activity. Although there is no significant difference in antiviral activity, these subtle structural differences might impact the pharmacokinetic properties of the molecules. For example, the predicted intestinal permeability of the *iso*-butyrate analogues **11c**,**d** and **12c**,**d** is higher than their counterparts **11a**,**b** and **12a**,**b**. Generally, all analogues **10**-**12** have a higher predicted intestinal permeability than ribavirin (see table S1) suggesting higher oral bioavailability.^62^

**Figure 3.**
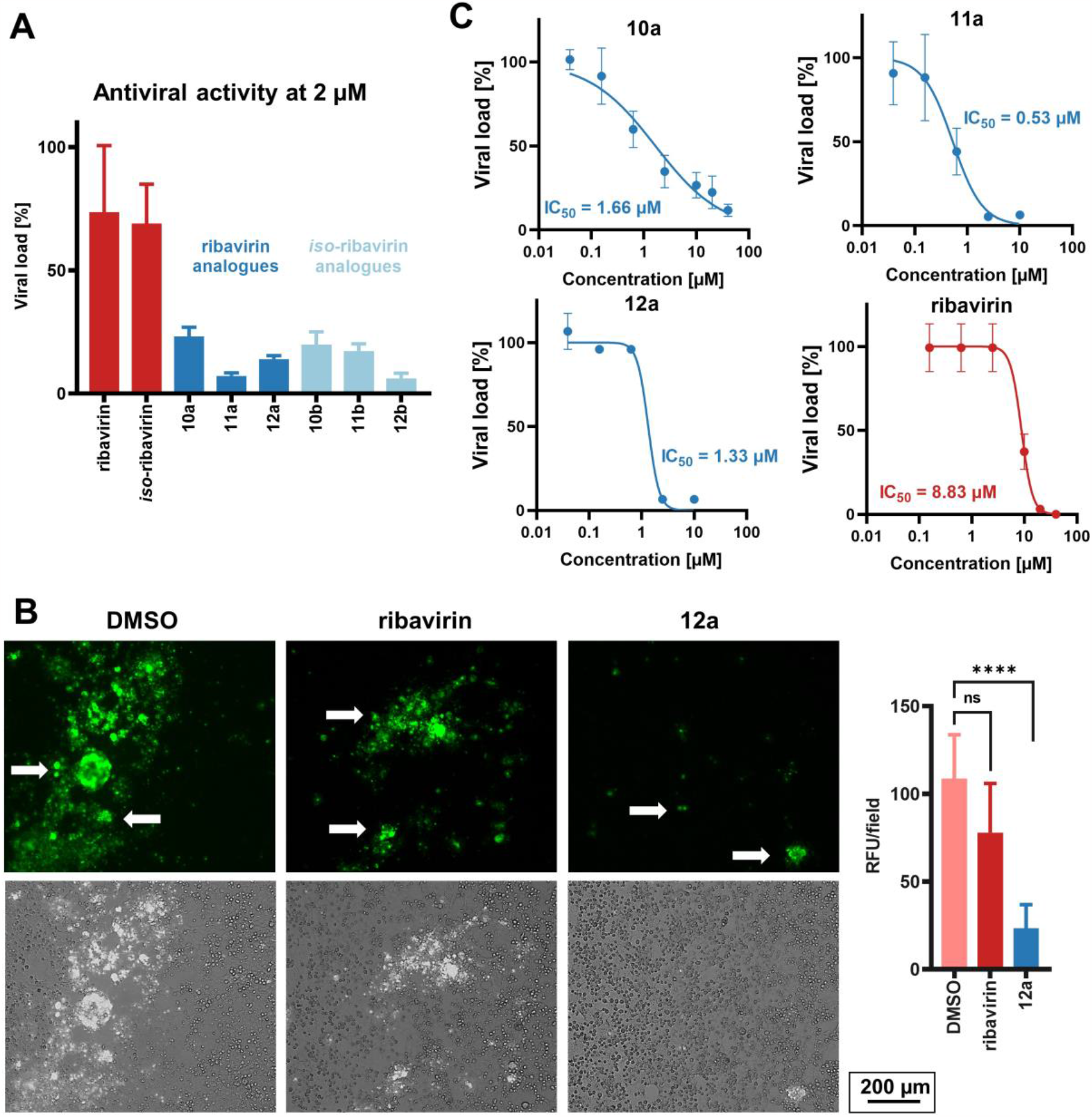
Antiviral activity of ribavirin type analogues against RSV. **A**: TCID_50_ assay (HEp-2 cells, 2 μM, n = 3) revealed high activity of compounds **10**-**12a**,**b**; **B**: Representative fluorescence images and quantification of immunostained RSV (see white arrows) presented in the infected HEp-2 cells (grey) with DMSO, ribavirin (2 μM), and **12a** (2 μM) treatment, respectively (*\*\*\*\** P *<* 0.0001, n ≥ 3); **C**: Antiviral activity (HEp-2 cells, RSV infection) of **10**-**12a** and ribavirin. Data are expressed as the percentage ± SEM of the DMSO control (n = 3).

Encouraged by the extraordinary antiviral activity of analogues **12a**-**d**, we followed up to determine the half-maximal inhibitory concentration (IC50) against RSV and the cytotoxicity (as reflected by CC50). All three derivatives of pseudo-C5′-isomer **10-12a** showed excellent anti-RSV activity with IC50 values at a low micromolar range (0.53-1.66 μM), even lower than that of ribavirin (IC50 = 8.83 μM, Fig. 3C). Analogue **11a** was the most active compound, with 16 times higher activity than ribavirin (IC50 = 0.53 μM). Of note, all three derivatives have a CC50 value >40 μM, indicating a high selectivity and thus a wide therapeutic window (see Fig. S8).

Next, we synthesized carbobicyclic uridine analogues **19**-**21** as “mechanistic probes” to rule out the possibility that the antiviral effect stemmed from the triazole base alone (Scheme 3). As such, uridine is a natural RNA building block and should not display significant antiviral activity.^63^ If these analogues are still active, this would suggest that the carbobicyclic core structure also contributes to its antiviral activity motif and more importantly can be used as a template for novel antivirals. To synthesize the uridine analogues **19**-**21**, alcohol **3** was first coupled with *N*3-benzoyluracil (**13**)^64, 65^ under Mitsunobu conditions. Stille cross-coupling of the Mitsunobu product **14** followed by debenzoylation again gave an excellent yield of the desired Diels-Alder precursor **15**. In our pilot Diels-Alder reaction (scheme not shown), the *N*3-benzoyl group was partially cleaved under the thermal condition, leading to a low isolated yield of the Diels-Alder adduct. Thus, debenzoylation at *N*3-benzoyl prior to the Diels-Alder reaction was essential. The conversion of the uracil diene **15** was relatively retarded compared to its ribavirin analogue; optimized molar ratio of the dienophile furnished the diverse cycloadducts **16**-**18** in moderate yield, with regio- and diastereoselectivity comparable to that of the ribavirin analogues (see also Fig S2). The final esterification and/or deprotection led to the carbobicyclic uracil analogues **19**-**21**.

**Scheme 3.**
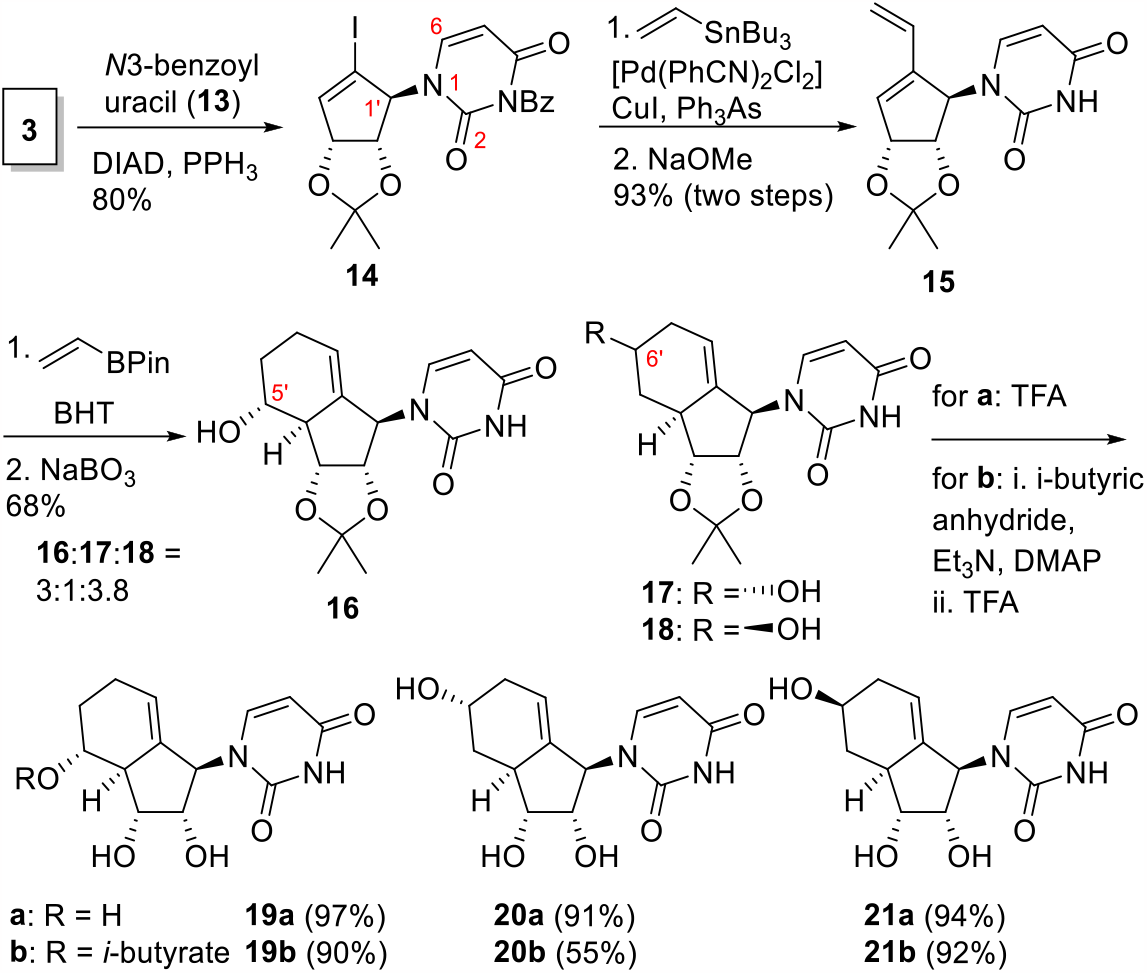
Synthesis of uridine analogues **19**-**21**.

In the subsequent cellular assays, uridine analogues **19**-**21a** showed good antiviral activity (see SI, Fig. S4) and almost no cytotoxicity (see Fig. S6). For example, the pseudo-C5′-OH analogue **19a** showed good antiviral activity (IC50 = 6.94 μM) and low cytotoxicity (CC50 >40 μM; Fig. S8). The uridine analogue is therefore equipotent to ribavirin. Collectively, these data confirmed that the carbobicyclic core contributes to the antiviral activity and can be used as a template for novel antivirals in future studies and, therefore, are promising antiviral drug leads.

During NMR analysis of the uracil analogues, we observed significant peak broadening, suggesting that the analogues exist as two structural conformers at room temperature. Variable temperature NMR studies showed that both conformers merge at elevated temperatures (315 K, see Fig. S9). The desired connectivity (*N*1 vs. *O*2) was confirmed by NMR (characteristic ^1^H and ^13^C shift at C1′)^66^ and NOESY studies (NOE coupling of H6 with the carbasugar core). In addition to NOE studies, the relative configuration was determined in analogy with ribavirin-type analogues, as the identical core structure (e.g. **10** and **19**) shares characteristic ^1^H-NMR chemical shifts and splitting patterns (see Fig S10-S12).

## Conclusion

In summary, we reported herein the design, synthesis, and biological activity of an unprecedented class of nucleoside analogues with a bicyclo-[4.3.0]-nonene core. Taking advantage of our concise and divergent synthetic route, rapid synthesis of a focused library was possible in a minimum of synthetic steps.

These analogues showed extraordinary antiviral activity against RSV, negligible cytotoxicity and potentially enhanced oral bioavailability. Both groups of nucleoside analogues (ribavirin- and uridine-type) showed higher antiviral activity than their parental compounds, suggesting that the carbobicyclic core is critical to their antiviral activities. Thus, this new antiviral structural motif opens tremendous opportunities to combat emerging/future outbreaks of viral diseases and antiviral drug resistance. Our discovery will catalyze future in-depth mechanistic investigation of the mode of action and *in vivo* animal efficacy, pharmacokinetics, and pharmacodynamics studies.

## Supporting information

Supporting Information with experimental details and analytical data

## Supporting Information

Experimental details and analytical data are available in the Supporting Information.

## Author Contributions

All authors have given approval to the final version of the manuscript.

## Notes

S.S., M.-Y.L. and B.W.-L.N. are inventors on a U.S. non-provisional patent application submitted by The Chinese University of Hong Kong which covers the described carbobicyclic core structure.

## Acknowledgment

Dedicated to Professor Tony K. M. Shing: We thank Prof. Tony K. M. Shing for his inspiration and helpful discussions. We thank Prof. Paul PK Chan (Department of Microbiology, CUHK) for providing the RSV virus stock and Dr KP Tao for the technical support in conducting the RSV virus genotyping and assistance in preparing the viral titration assays. We also thank Sarah Ng and Alice Ting In Wong (Department of Chemistry, CUHK) for their technical support in conducting NMR and HRMS analysis.

