## Supporting Information with experimental details and analytical data for "Diversification of Nucleoside Analogues as Potent Antiviral Drug Leads"

### Additional Figures

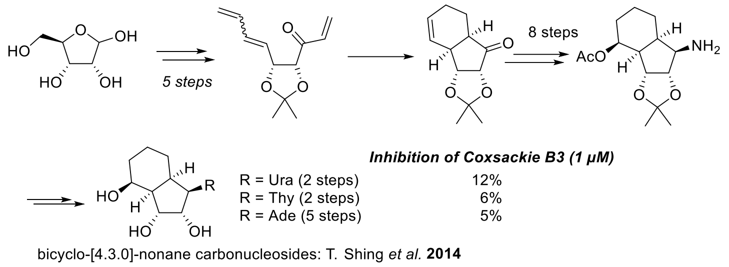

**Figure S1:** Previously reported bicyclic nucleoside analogues. Bicyclo-[4.3.0]-nonane structured nucleoside analogues were synthesized with a long synthetic route (16-19 steps) and low diversity (3 analogues).[1] At the same time the antiviral activity remained low, so this approach is unattractive.

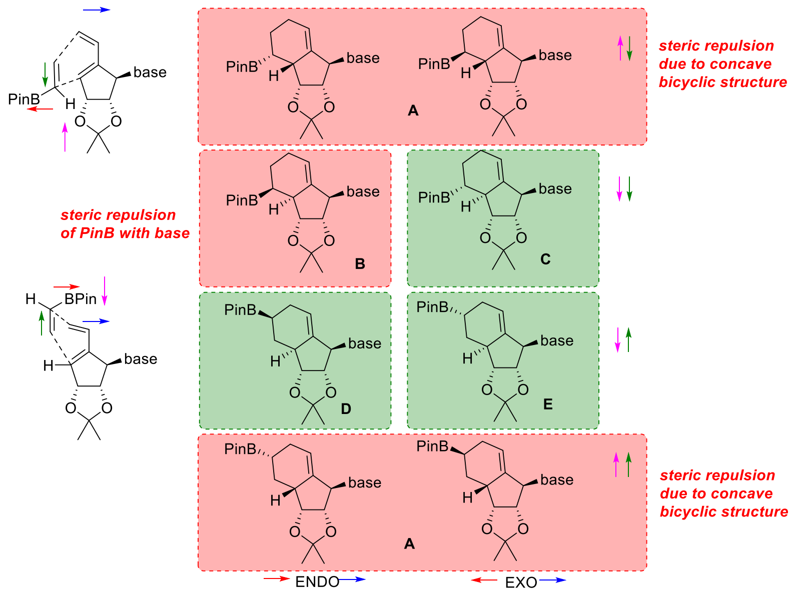

**Figure S2:** Possible isomers during intermolecular Diels-Alder reaction, only green isomers were obtained. In theory, eight regioisomers in the intermolecular Diels-Alder reaction are possible. Experimentally, we found that in all cases dienophile approaches from the top (pink arrow going down). Isomers **A** were not observed when ribavirin was used, however, traces (<5%) were found during the synthesis of uridine type analogues. Of the four remaining isomers, isomer **B** was not observed, probably due to steric interactions between the bulky pinacol residue and the nucleobase. Isomer **C** should be the electronically favoured isomer. Isomer **D** is isolated in significantly higher yields than isomer **E**, it is believed that this isomer is electronically favoured in comparison to isomer **E** as both isomers share similar steric repulsions.

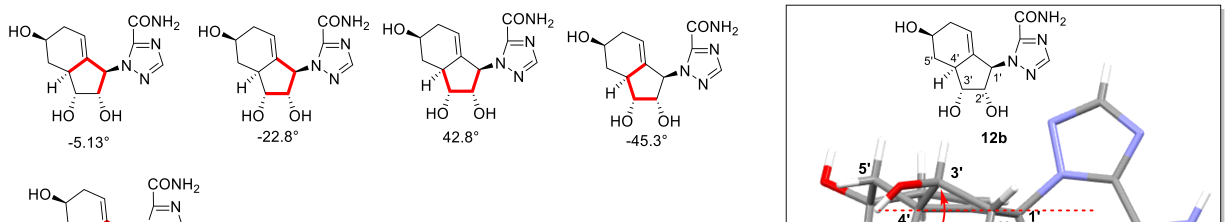

$$\tan(P) = \frac{(31.0°-22.8°)-(-45.3°-5.13°)}{2x\ 42.8°\ (\sin 36° + \sin 72°)} = 0.45$$

P = 24.0°

**Figure S3:** Calculation of angle of pseudorotation P using torsion angles derived from X-ray structure of **12b**.[2] For **12b** the P was determined as +24.0° which correlates with a C3' endo (North) conformation. This conformation is also observed in natural nucleosides and therefore is promising for biological activity.[2]

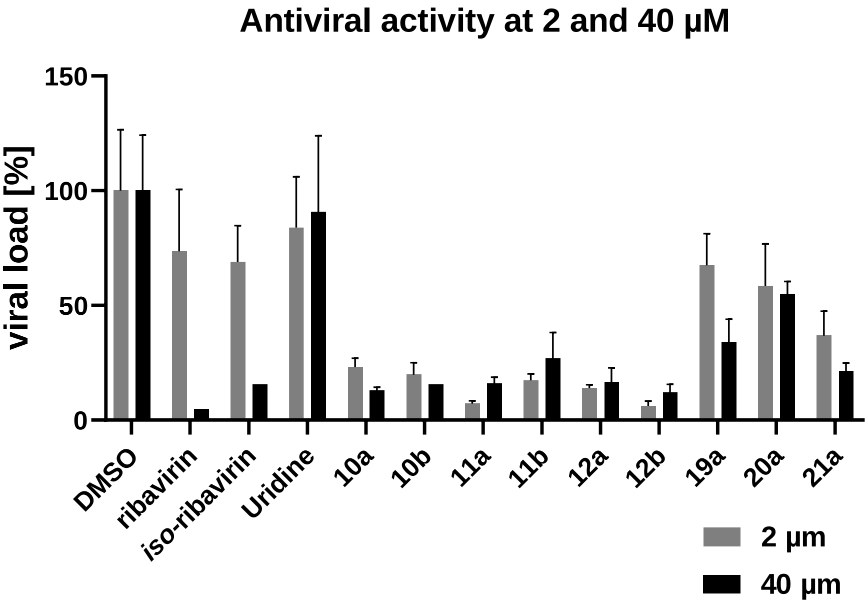

**Figure S4:** Antiviral screening at 2 and 40 µM compound concentration against RSV in $TCID_{50}$ assay using HEp-2 cells (MOI 0.02). Viral load in % of DMSO control (n = 3).

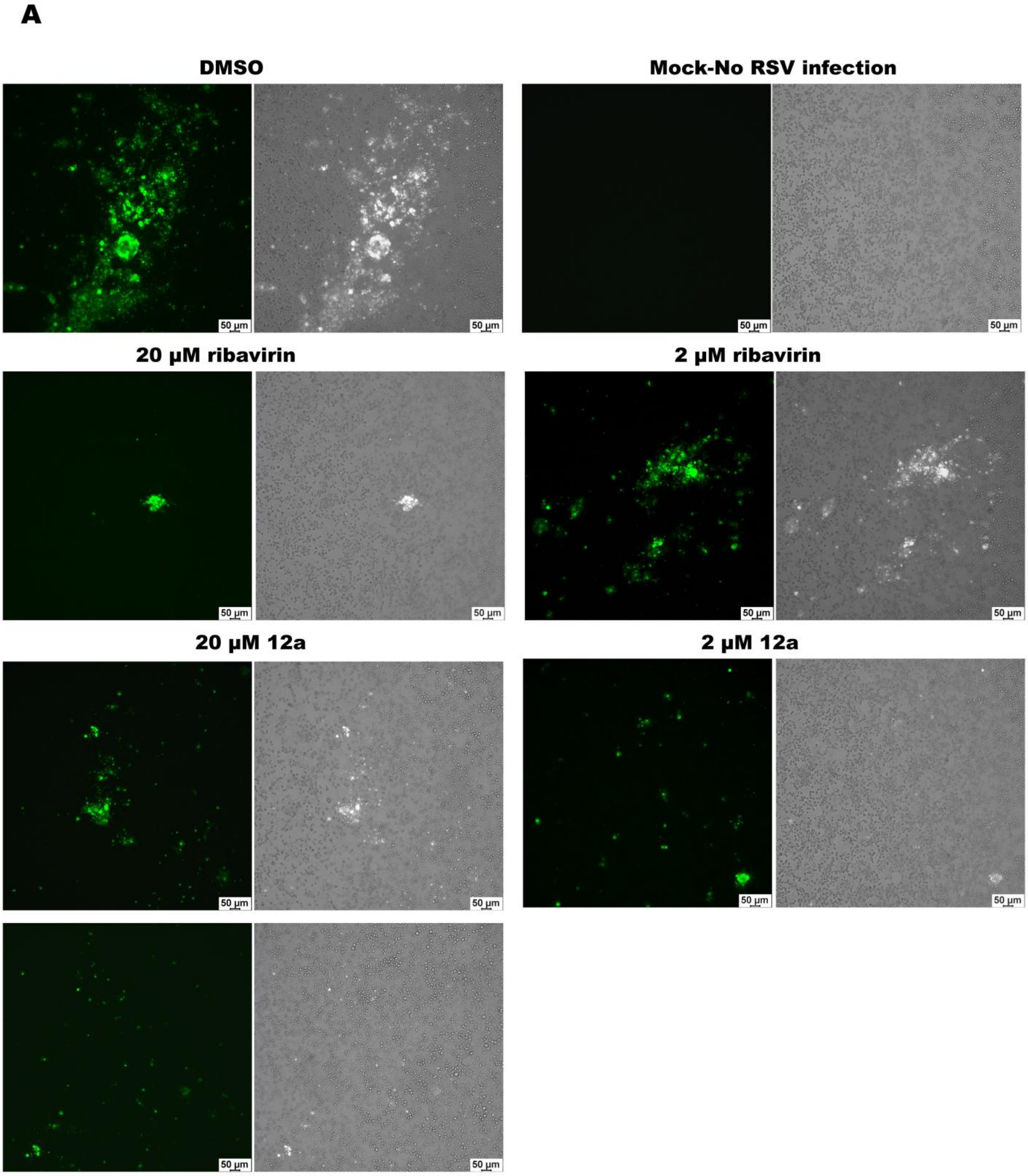
A
DMSO
Mock-No RSV infection
20 μM ribavirin
2 μM ribavirin
20 μM 12a
2 μM 12a
50 μm

B

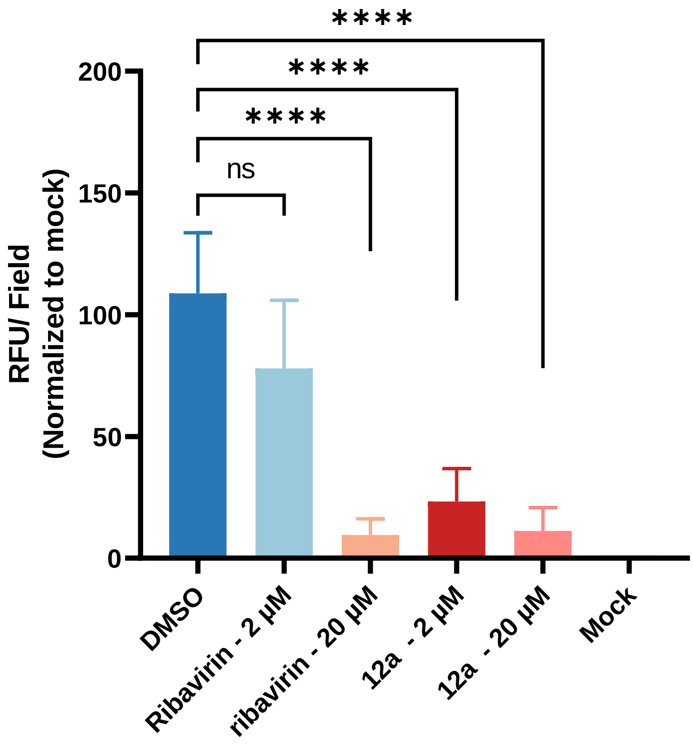

**Figure S5:** Ribavirin and **12a** inhibit RSV replication and rescue RSV infected HEp-2 cells with decreased cytopathic effect (CPE). (A) Representative fluorescence microscopy (10x) of mock-infected or RSV-infected HEp-2 cells with DMSO, ribavirin (2 μM, 20 μM) and **12a** (2 μM, 20 μM) treatments, stained with anti-RSV fusion protein antibody, followed by secondary antibody conjugated with alexa fluor 488. (B) The relative fluorescence unit (RFU)/ field was quantified by image J (Version 1.52, NIH, USA) and normalized to the mock group. Results were means + SD from 3 sets of cultures, 5-6 of random fields were pooled and analyzed for each culture. Statistical significance was analyzed by one-way analysis of variance (ANOVA) using GraphPad Prism 9 (GraphPad Software, San Diego, CA) (**P* < 0.05; ***P* < 0.01; *****P* < 0.0001; ns stands for not statistically significant).

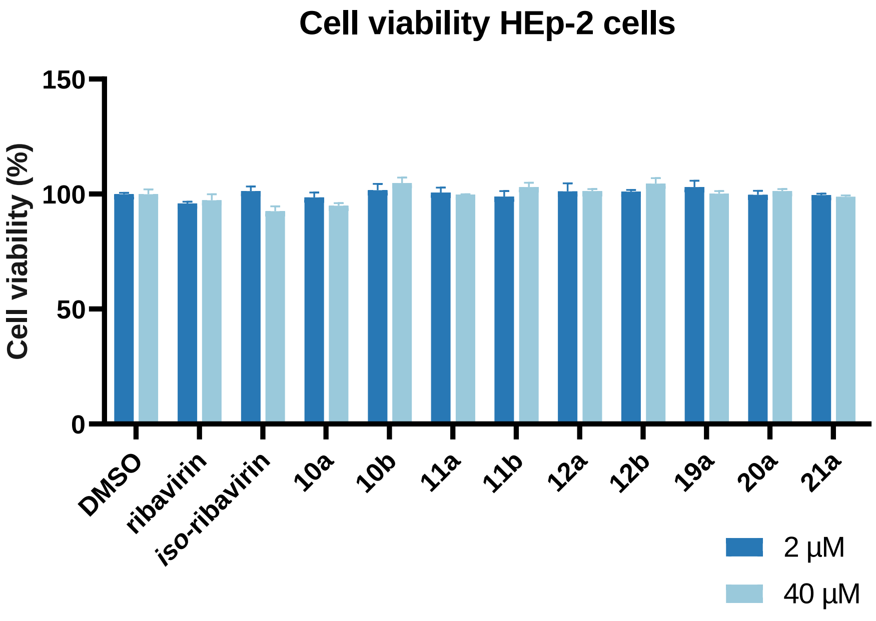

**Figure S6:** Cell viability (HEp-2 cells, MTT assay) of compounds at 2 and 40 µM. Data are expressed as the percentage ± SEM of the DMSO control, n=3.

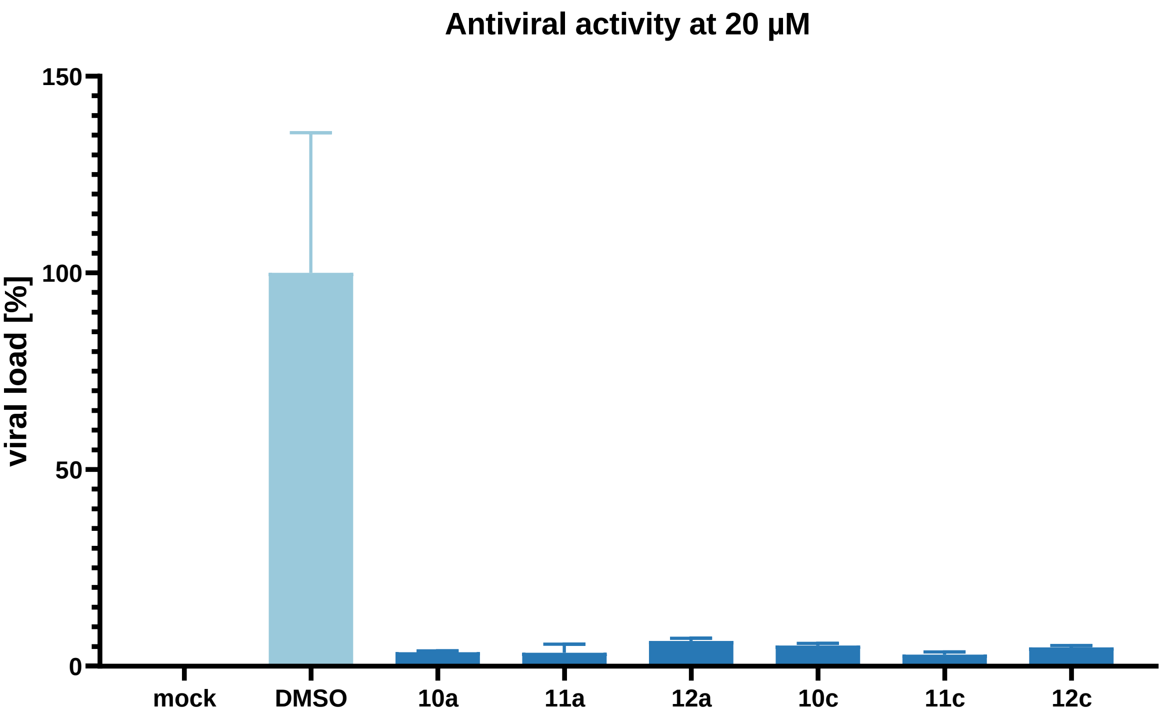

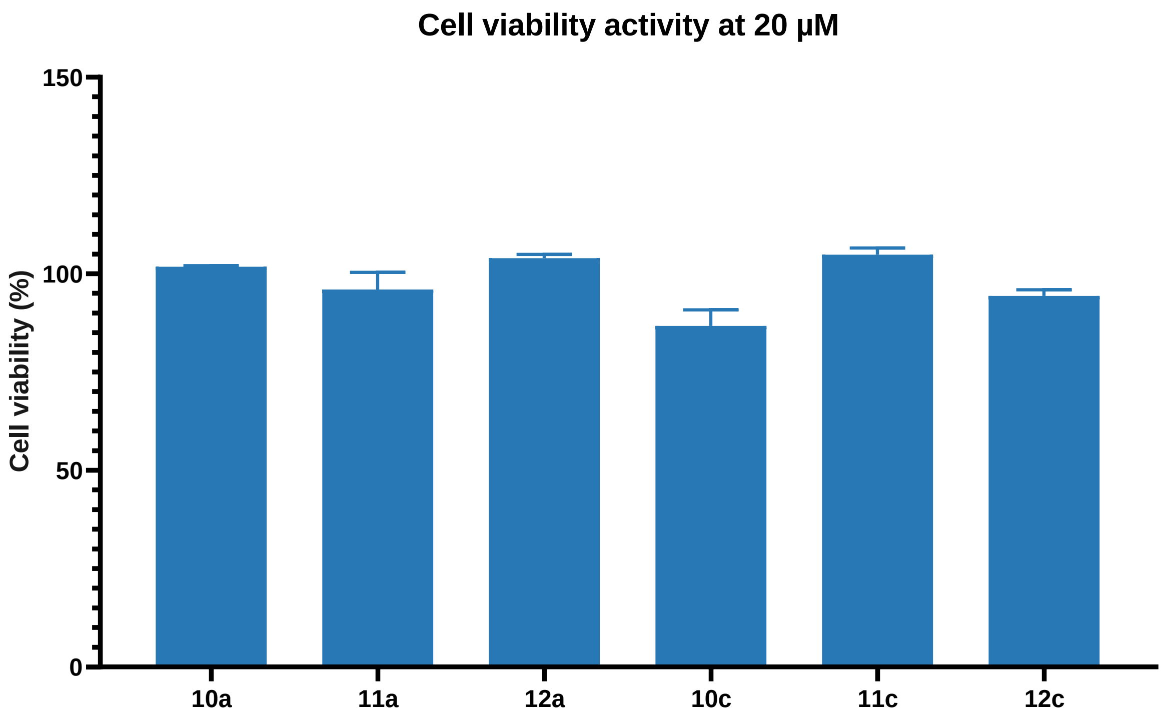

**Figure S7:** Antiviral screening and cell viability at 20 µM compound concentration against RSV in $TCID_{50}$ assay using HEp-2 cells (MOI 0.02). Inhibition and cell viability in % of DMSO control (n = 3).

**Table S1.** Predicted Caco-2 permeability of the tested compounds. The intestinal permeabilities in Caco-2 monolayer model (Caco-2 permeability, $P_{app}$) of these nucleoside analogues were predicted using ADMETlab 2.0.[3] Chemical structures of all tested compounds were converted to SMILES in ChemDraw Professional 16.0 as input for ADMETlab 2.0. The predicted $P_{app}$ of all tested compounds were shown in **Table S1** with propranolol ($P_{app}$ of 2.38781E-05 cm/s) and atenolol ($P_{app}$ of 3.22107E-06 cm/s) as transcellular and paracellular markers, respectively.

| Compound | Predicted $P_{app}$ ($10^{-6}$ cm/s) |
|---|---|
| Ribavirin | 2.21 |
| Uridine | 0.84 |
| Entecavir | 0.52 |
| 10a | 5.30 |
| 10c | 4.36 |
| 11a | 3.18 |
| 11c | 4.14 |
| 12a | 3.95 |
| 12c | 4.33 |
| 10b | 6.46 |
| 10d | 5.92 |
| 11b | 3.77 |
| 11d | 5.33 |
| 12b | 4.57 |
| 12d | 6.41 |
| 19a | 2.65 |
| 19b | 2.45 |
| 20a | 1.18 |
| 20b | 2.19 |
| 21a | 1.66 |
| 21b | 2.52 |

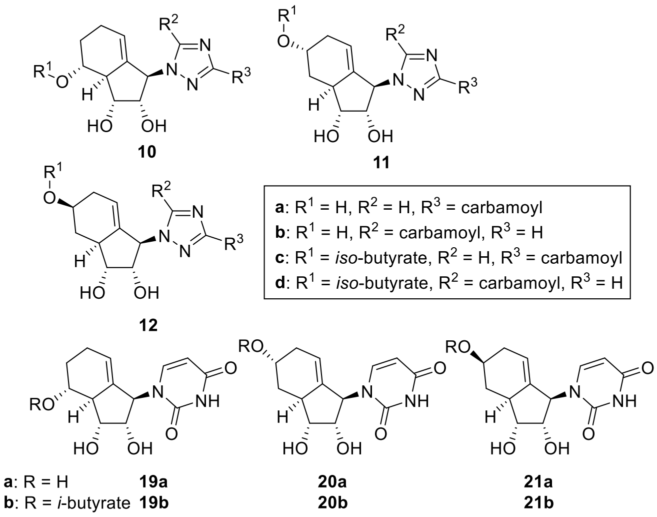

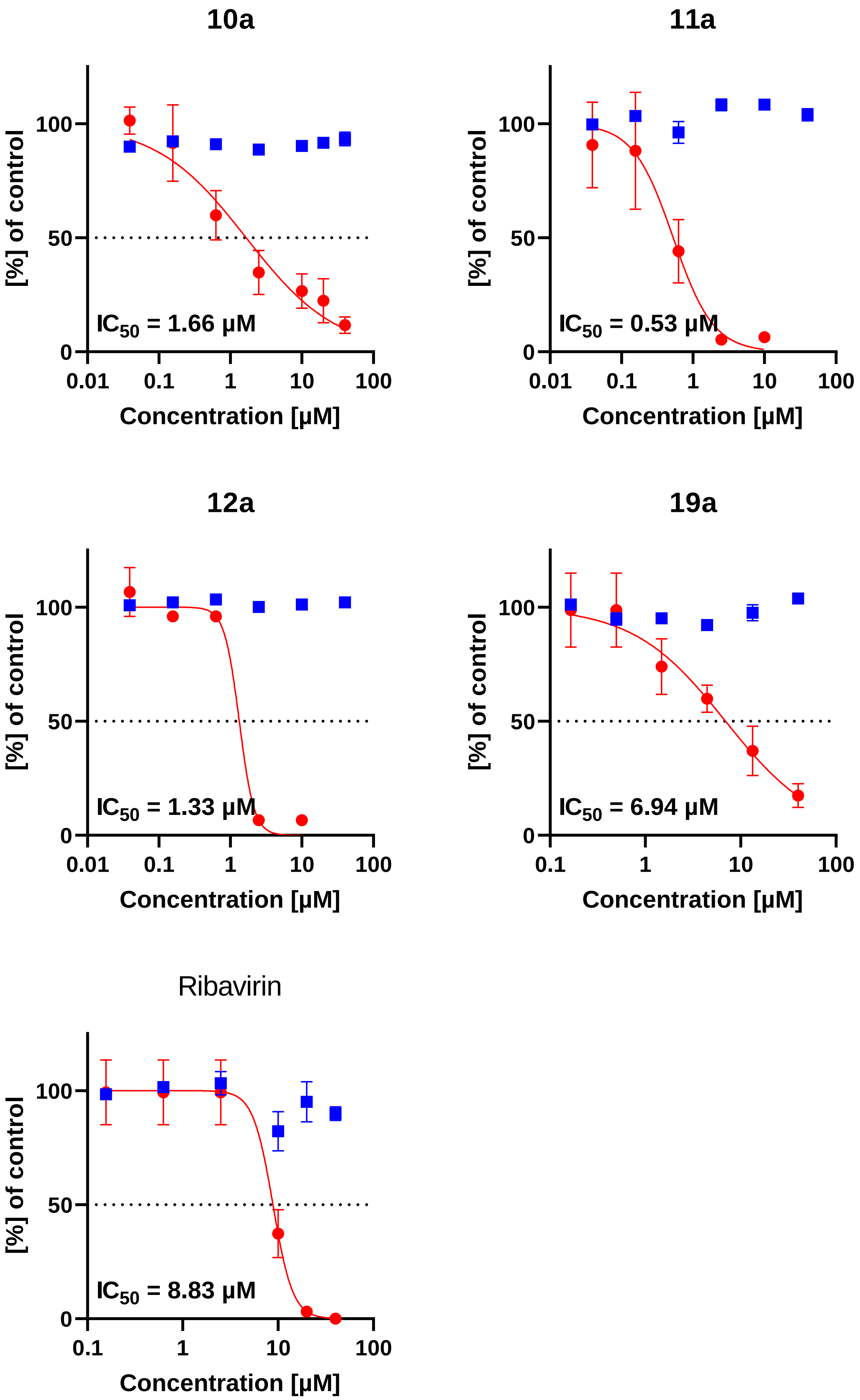

**Figure S8:** Antiviral activity (HEp-2 cells, RSV infection, red) and cytotoxicity (HEp-2 cells, MTT assay, blue) of **10**-**12a**, **19a** and ribavirin. Data are expressed as the percentage ± SEM of the DMSO control. Each data point represents the mean of triplicate values, and the SEM is represented by the error bars.

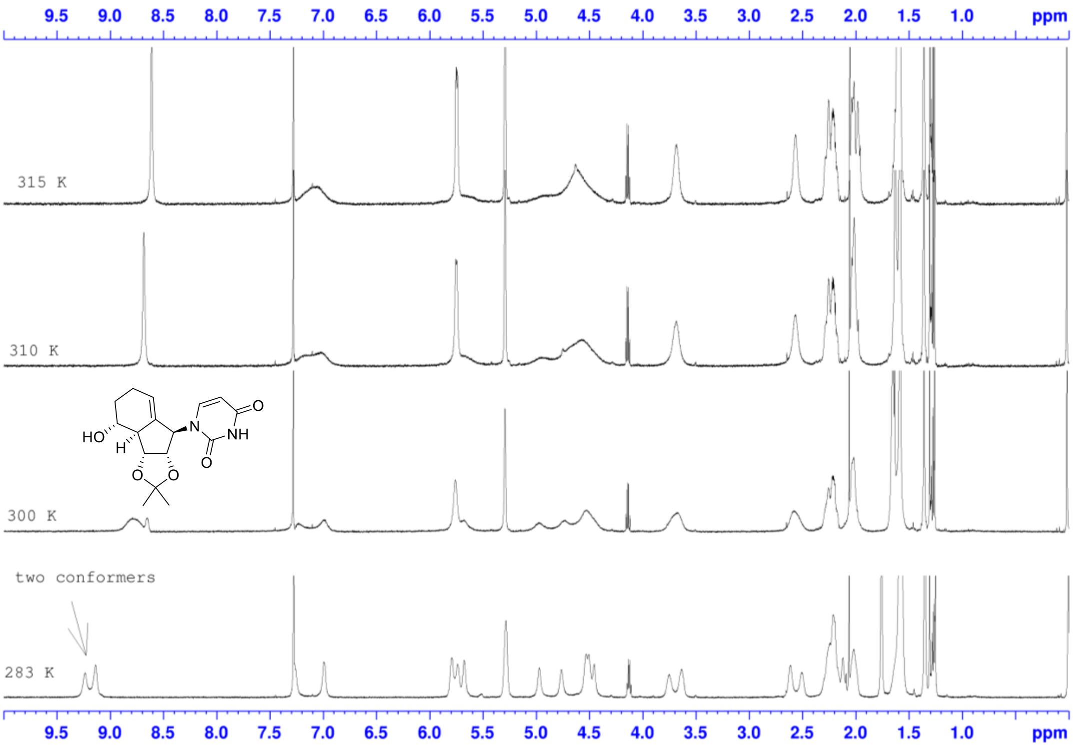

**Figure S9:** Temperature dependent NMR study of compound **16** shows two conformers. For all uridine-type analogues after Mitsunobu coupling, broad signals were observed in proton and carbon NMR. A temperature-dependent study of compound **16** revealed that two conformers are present at temperatures below room temperature, which are slowly interconvert into each other at the temperature range which is available on the NMR machine used for this study. Due to this signal broadening, both proton and carbon NMR are of limited use for identification of the structural identity. In detail, even with prolonged measuring time, 2D NMR studies, different solvents, or different acquisition temperatures, proton NMR was often of limited use and not all carbon signals could be found.

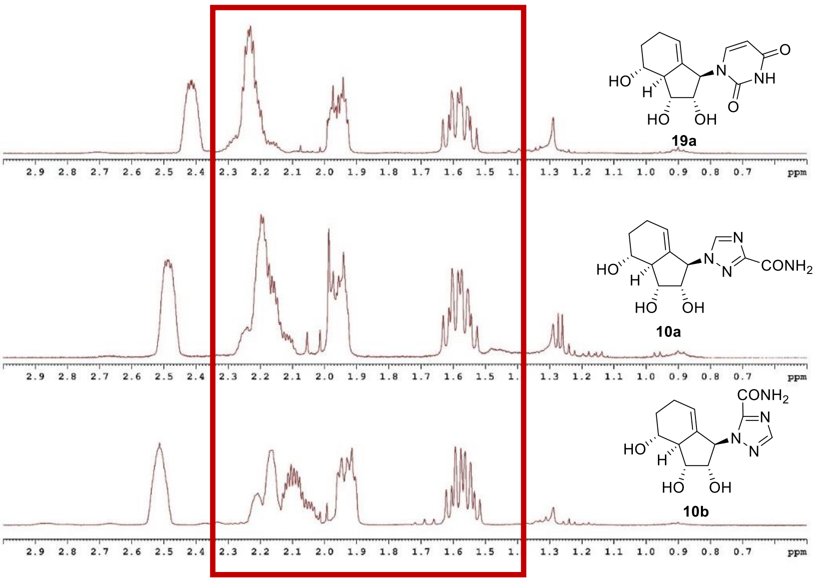

**Figure S10:** NMR for compounds **10a**,**b** and **19a**.

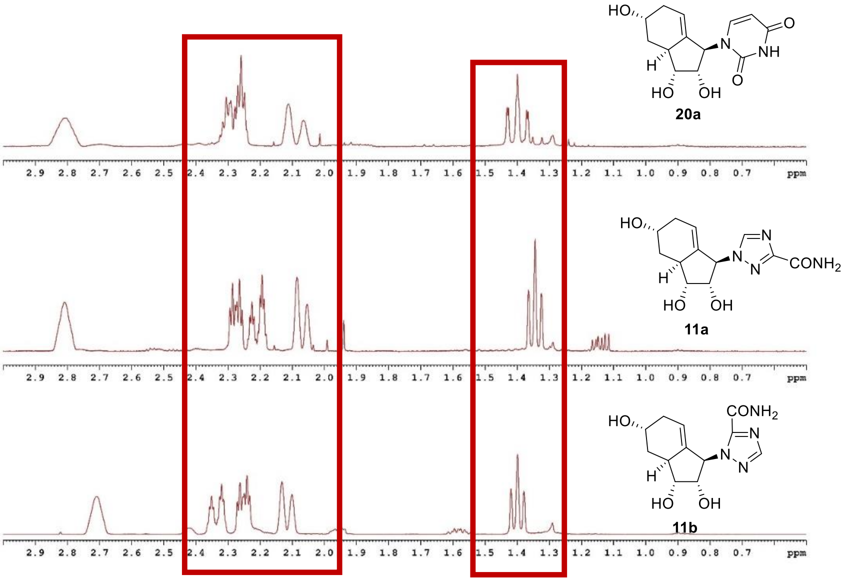

**Figure S11:** NMR for compounds **11a**,**b** and **20a**.

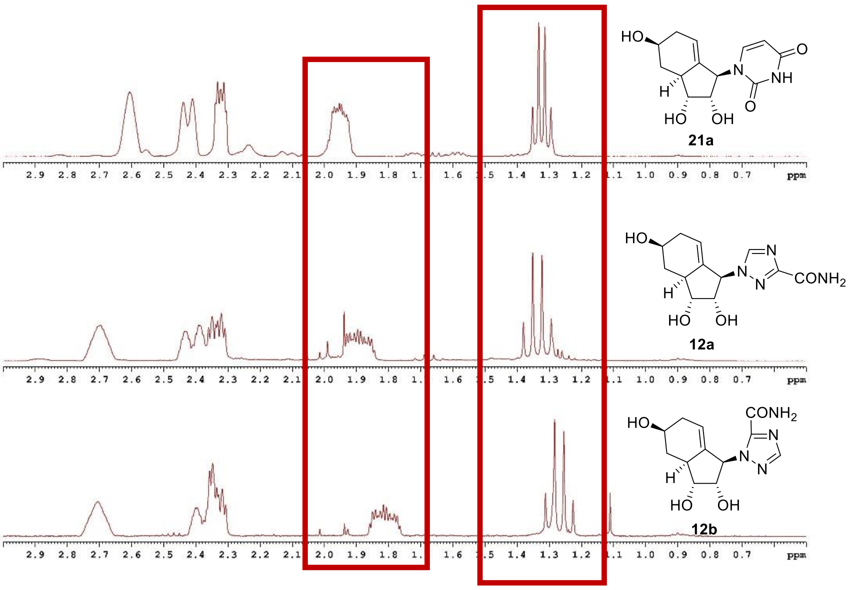

**Figure S12:** NMR for compounds **12a**,**b** and **21a**.

### Cellular Assays

#### Cells

HEp-2 cells (ATCC, CCL-23) were maintained in Dulbecco's modified Eagle's medium (DMEM, Life Technologies, Gaithersburg) with 10% fetal bovine serum (FBS, Life Technologies) and 1% penicillin-streptomycin (PS, Life Technologies).

#### Virus strains

The working RSV-A stock was gifted by Prof. Paul PK Chan (Department of Microbiology, CUHK) and identity was confirmed using the RSV nucleoprotein (N protein) and attachment glycoprotein (G protein) genotyping. The strain used had a high similarity to human RSV Long strain [HRSVA/Maryland.USA/Long/1956, sequence ID: OK649668.1], 99% in positions 1716-2119 (404 bp) and 100% in positions 5186-5590 (405 bp), respectively.

#### Virus propagation

HEp-2 cells were plated in a T175 flask. At 80-90% confluency, cells were washed with phosphate-buffered saline (PBS) and inoculated with RSV-A inoculum for 2 h at 37 °C. Virus growth medium with reduced FBS (2%) was added to end the virus absorption and cells were incubated for days until >70% of cells developed cytopathic effect (CPE). The culture supernatant was collected and centrifuged at 2,000 g at 4 °C for 10 mins, and aliquots of the supernatant were frozen at -80 °C for viral quantification and later use.

#### Virus titration

HEp-2 cells were seeded on 96-well tissue culture plates one day before the viral titration assay. Cells were washed once with PBS. Virus samples or culture supernatants were titrated in serial half-$log_{10}$ dilutions with the corresponding culture medium before adding the diluted virus to the cell plates in quadruplicate. The highest viral dilution leading to CPE was recorded and the 50% tissue culture infectious dose ($TCID_{50}$) was calculated using the Spearman-Karber method.

#### RSV inhibition assays

HEp-2 cells were seeded at 5 × $10^4$ cells per well in 96-well plates the day before and infection at a multiplicity of infection (MOI) of 0.02 at 37 °C for 1 h. The inoculum was discarded and the cells were washed once with PBS. Cell culture medium (DMEM, 2% FBS) with the corresponding compounds at their designated concentration was added. All compounds were dissolved in dimethyl sulfoxide (DMSO, Santa Cruz Biotechnology Inc., Dallas, TX, USA) at 8 mM and further dilute serially in DMEM containing 0.5% DMSO to final concentrations ranging from 39 nM to 40 μM except stated otherwise, with 0.5% DMSO in DMEM as the vehicle control. The supernatant of the cultures was collected and frozen at -80 °C at 72 hours post-infection for viral titration.[4] The potency of the compounds in inhibiting the RSV replication was expressed in half maximal inhibitory concentration ($IC_{50}$).

##### Assessment of cell cytotoxicity by the MTT assay

Cytotoxicity assays were performed with thiazolyl blue tetrazolium bromide (MTT, Thermo Fisher, Carlsbad, CA, M6494) after the cells were incubated with the compounds for 72 hours. The 50% cytotoxic concentration ($CC_{50}$) was defined as a 50% reduction in absorbance measured at 490 nm with a reference wavelength of 690nm by a spectrophotometer (Synergy HTX Multi-mode Microplate Reader; Biotek Instruments, Inc., Winooski, VT, USA) compared to control wells.

$IC_{50}$ and $CC_{50}$ values were calculated by fitting the data to a sigmoidal curve equation in GraphPad Prism 9 (GraphPad Software, San Diego, CA).

##### Immunostaining

HEp-2 cells were seeded at 1.5 × $10^5$ cells per well in 6-well plates the day before infection (MOI = 0.02, at 37 °C for 1 h). The inoculum was discarded and the cells were washed once with PBS. Cell culture medium (DMEM, 2% FBS) with the presence or absence of compounds were added to the cells, with 0.5% DMSO in DMEM as the vehicle control. At 72 hours post-infection, cells were fixed in 4% paraformaldehyde (PFA), permeabilised with 0.1% Triton X-100, blocked with 2% bovine serum albumin (BSA) with 2% normal goat serum (NGS), and stained for RSV F protein with primary antibody (anti-RSV antibody, fusion protein, all type A, B strains, clone 131-2A, MAB8599, 1:500, Merck) followed by secondary antibody (goat anti-mouse IgG (H + L), alexa fluor 488, A-11001, 1:500, Invitrogen). RSV infected or non-infected cells were imaged using a fluorescence inverted microscope (Leica DMI8, Germany) in 10× magnification. Mean fluorescence intensity within each field was measured and quantified by image J (Version 1.52, NIH, USA), the relative fluorescence unit (RFU)/ field was normalized to the mock (non-infection) group.

### Chemical Synthesis

#### *General Methods*

Reagents and solvents were purchased from commercial suppliers (Sigma Aldrich, TCI Chemicals, Meryer Chemicals (Shanghai), Bidepharm, Energy Chemical and Dieckmann Chemical) in the highest purity available. Common solvents used for reactions (MeOH, DCM, THF, NMP, MeCN, pyridine) were purchased in anhydrous conditions and stored over molecular sieve. NMP and toluene were stored over $CaH_2$ and distilled before usage. The reactions were performed under an $N_2$ atmosphere in oven-dried glassware which had been flushed with nitrogen. All reagents and solvents were handled using standard Schlenk techniques. Temperatures above room temperature refer to the oil bath temperatures. For cooling to 0 °C a water/ice bath was used.

Reaction progress was monitored using TLC monitoring and was performed with silica gel pre-coated aluminium sheets (Machery-Nagel "DC-Fertigfolien ALUGRAM® SIL G/$UV_{254}$; 0.20 mm Schichtdicke Kieselgel 60 mit Fluoreszenz-Indikator $UV_{254}$"). Compounds could be visualized by UV light and staining with a solution of CAM (1.0 g $Ce(SO_4)_2$), 2.5 g $(NH_4)_6Mo_7O_{24}$, 8 mL $H_2SO_4$ in 100 mL $H_2O$) with subsequent heating. Column chromatography was performed using silica gel 60 (230-400 mesh) and solvents were used without prior purification.

All NMR spectra were recorded on Bruker spectrometer at the Chinese University of Hong Kong or at Master Dynamic Limited (Hong Kong) with operating frequencies of 100 ($^{13}C$), 125($^{13}C$), 400 ($^{1}H$) and 600 ($^{1}H$) MHz. Deuterated solvents ($CDCl_3$ and MeOD) were manufactured from Eurisotop and purchased from Meryer Chemicals and calibrated to the residual signal of undeuterated solvents.[5]

High-resolution mass spectra were recorded under the supervision of Alice Ting In Wong at the Chemistry Department at the Chinese University of Hong Kong using ESI or APCI ionization on a *Q Exactive Focus Orbitrap* spectrometer from Thermo Scientific.Optical rotations were recorded on a *Bellingham+Stanley ADP440+* polarimeter.

#### *Synthesis of Ribavirin Analogues*

##### Synthesis of (3a*R*,6a*R*)-5-iodo-2,2-dimethyl-3a,6a-dihydro-4*H*-cyclopenta[*d*][1,3] dioxol-4-one (S1)

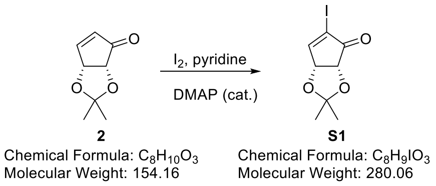

To a stirred solution of unsaturated ketone **2** (0.77 g, 5.00 mmol, 1.00 eq.) in dry DCM (25 mL) was added iodine (3.17 g, 12.5 mmol, 2.50 eq.). Then pyridine (0.50 mL, 6.25 mmol, 1.25 eq.) and DMAP (30.5 mg, 0.25 mmol, 5 mol%) were added subsequently at 0 °C. The reaction was then stirred at room temperature. After 1.5 h the reaction was quenched by aq. $Na_2S_2O_3$ solution (15 g/50 mL $H_2O$) and extracted with DCM (20 mL x 2). The combined organic phase was washed with brine (30 mL), dried over $MgSO_4$ and concentrated to give the crude which was purified by column chromatography on silica gel (Hex/EA 4:1). Iodide **S1** was isolated as white crystalline solid (1.27 g, 0.45 mmol, 90%). **$R_f$** = 0.38 (Hex/EA 4:1); **$^1$H-NMR** (400 MHz, $CDCl_3$) δ [ppm] = 7.96 (d, *J* = 2.6 Hz, 1H), 5.21 (dd, *J* = 5.6, 2.6 Hz, 1H), 4.51 (d, *J* = 5.6 Hz, 1H), 1.40 (s, 3H), 1.36 (s, 3H); **$^{13}$C{$^1$H}-NMR** (101 MHz, $CDCl_3$) δ [ppm] = 197.7, 165.0, 115.9, 105.8, 79.8, 73.8, 27.4, 26.5.

The analytical data were in accordance with previously published data.[6]

##### Synthesis of (3a*R*,6aR)-5-iodo-2,2-dimethyl-3a,6a-dihydro-4*H*-cyclopenta[*d*][1,3] dioxol-4-ol (3)

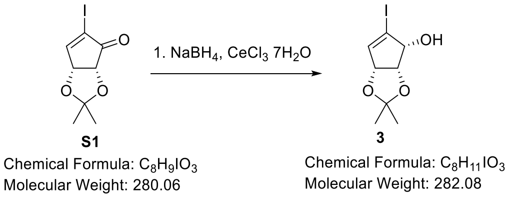

Unsaturated ketone **S1** (1.28 g, 4.53 mmol, 1.00 eq.) was dissolved in MeOH (40 mL) and $CeCl_3$*7$H_2O$ (1.94 g, 5.22 mmol, 1.15 eq.) was added. Then at 0 °C $NaBH_4$ (197 mg, 5.22 mmol, 1.15 eq.) was added. The reaction was stirred for 1h, then quenched with sat. $NH_4Cl$ solution (3 mL). The mixture was concentrated *in vacuo* and the crude was redissolved in an EtOAc/$H_2O$ mixture (1:1, 40 mL). After the separation of the organic phase, the aqueous phase was extracted with EtOAc (3x 15 mL). The combined organic phases were washed with brine (25 mL), dried over $MgSO_4$ and concentrated *in vacuo*. Alcohol **3** (1.18 g, 4.18 mmol, 92%) was isolated as a white solid in sufficient purity.

**R$_f$** = 0.32 (Hex:EtOAc 4:1); **$^1$H-NMR** (400 MHz, $CDCl_3$) δ [ppm] = 6.31 (t, *J* = 1.5 Hz, 1H), 4.92 (dd, *J* = 1.8, 5.6 Hz, 1H), 4.69 (t, *J* = 4.7 Hz, 1H), 4.42 (ddd, *J* = 1.3, 5.6, 10.4 Hz, 1H), 2.81 (d, *J* = 10.5 Hz, 1H), 1.43 (s, 3H), 1.40 (s, 3H).
The analytical data were in accordance with previously published data.[6]

#### Synthesis of Methyl 1-((3a*S*,4*S*,6a*R*)-5-iodo-2,2-dimethyl-3a,6a-dihydro-4*H*-cyclopenta[*d*][1,3]dioxol-4-yl)-1H-1,2,4-triazole-3-carboxylate (5a) and methyl 1-((3a*S*,4*S*,6a*R*)-5-iodo-2,2-dimethyl-3a,6a-dihydro-4*H*-cyclopenta[*d*][1,3]dioxol-4-yl)-1H-1,2,4-triazole-5-carboxylate (5b)

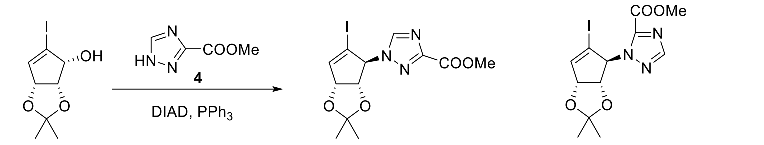

**3** — Chemical Formula: $C_8H_{11}IO_3$, Molecular Weight: 282.08
**5a** — Chemical Formula: $C_{12}H_{14}IN_3O_4$, Molecular Weight: 391.17
**5b** — Chemical Formula: $C_{12}H_{14}IN_3O_4$, Molecular Weight: 391.17

The alcohol **3** (1.00 g, 3.55 mmol, 1.00 eq.) was dissolved in DCM (50 mL). After the addition of the triazole **4** (0.52 g, 4.08 mmol, 1.15 eq.) and 4Å MS the mixture was stirred for 2h. Then at 0 °C first $PPh_3$ (1.12 g, 4.25 mmol, 1.20 eq.) and DIAD (0.84 mL, 4.25 mmol, 1.20 eq.) was added. The reaction was stirred for 18h. Then the reaction was filtered, and the solvent was removed *in vacuo*. Purification by column chromatography ($SiO_2$, Hex:EtOAc 3:1 to 1:1) gave first **5b** as a yellow oil (0.59 g, 1.51 mmol, 42%), later **5a** as a white foam (0.49 g, 1.25 mmol, 36%).

**5a: R$_f$** = 0.13 (Hex:EtOAc 3:1); **$^1$H-NMR** (400 MHz, $CDCl_3$) δ [ppm] = 8.26 (s, 1H), 6.60 (s, 1H), 5.36 (m, 2H), 4.89 (d, *J* = 6.8 Hz, 1H), 3.99 (s, 3H), 1.47 (s, 3H), 1.34 (s, 3H); **$^{13}$C{$^1$H}-NMR** (101 MHz, $CDCl_3$) δ [ppm] =160.0, 156.1, 146.3, 145.3, 113.1, 96.1, 85.0, 82.5, 77.7, 53.0, 27.2, 25.9; **HRMS** (ESI, 3.5kV) calc for $[M+Na]^+$ $C_{12}H_{14}IN_3O_4Na^+$ 413.99212, found 413.99171.

**5b: R$_f$** = 0.24 (Hex:EtOAc 3:1); **$^1$H-NMR** (400 MHz, $CDCl_3$) δ [ppm] = 8.03 (s, 1H), 6.58 (s, 1H), 6.54 (s, 1H), 5.35 (dt, *J* = 1.8, 5.7 Hz, 1H), 4.84 (d, *J* = 5.8 Hz, 1H), 4.06 (s, 3H), 1.50 (s, 3H), 1.37 (s, 3H); **$^{13}$C{$^1$H}-NMR** (101 MHz, $CDCl_3$) δ [ppm] = 158.3, 151.9, 145.4, 145.2, 112.9, 96.9, 85.3, 83.0, 76.8, 53.5, 27.4, 26.1; **HRMS** (ESI, 3.5kV) calc for $[M+Na]^+$ $C_{12}H_{14}IN_3O_4Na^+$ 413.99212, found 413.99183.

#### Synthesis of Methyl 1-((3a*S*,4*S*,6a*R*)-2,2-dimethyl-5-vinyl-3a,6a-dihydro-4*H*-cyclopenta[*d*][1,3]dioxol-4-yl)-1H-1,2,4-triazole-3-carboxylate (6a)

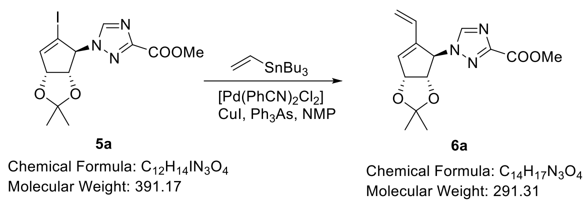

Iodide **5a** (478 mg, 1.22 mmol, 1.00 eq.) was dissolved in NMP (4 mL, freshly distilled) and [Pd(PhCN)$_2$Cl$_2$] (14.1 mg, 36.7 µmol, 3 mol%), CuI (14.0 mg, 73.3 µmol, 6 mol%) and Ph$_3$As (22.5 mg, 73.3 µmol, 6 mol%) was added. Then at 0 °C vinyl tributyltin (430 µL, 1.47 mmol, 1.20 eq.) was added dropwise. The reaction was stirred at this temperature for 3h. The reaction was quenched with sat. NaHCO$_3$ solution (10 mL) and EtOAc (10 mL). After the separation of the organic phase, the aqueous phase was extracted with EtOAc (3x 10 mL). The combined organic phases were washed with brine (15 mL), dried over MgSO$_4$ and concentrated *in vacuo*. Purification by column chromatography (SiO$_2$, Hex:EtOAc 2:1 to 1:1) gave **6a** as a white solid (349 mg, 1.20 mmol, 98%). **R$_f$** = 0.29 (Hex:EtOAc 1:1); **$^1$H-NMR** (400 MHz, CDCl$_3$) δ [ppm] = 8.07 (s, 1H), 6.47 (dd, *J* = 11.2, 17.6 Hz, 1H), 6.21 (s, 1H), 5.68 (s, 1H), 5.42 (d, *J* = 5.4 Hz, 1H), 5.23 (d, *J* = 17.7 Hz, 1H), 4.70 (d, *J* = 5.6 Hz, 1H), 3.99 (s, 3H), 1.42 (s, 3H), 1.33 (s, 3H); **$^{13}$C{$^1$H}-NMR** (101 MHz, CDCl$_3$) δ [ppm] = 160.1, 155.6, 143.9, 138.8, 136.5, 130.0, 119.7, 112.4, 84.4, 83.4, 69.4, 53.0, 27.5, 26.0; **HRMS** (ESI, 3.5kV) calc for [M+Na]$^+$ C$_{14}$H$_{17}$N$_3$O$_4$Na$^+$ 314.11113, found 314.11088.

#### Synthesis of Methyl 1-((3a*R*,3b*R*,4*R*,8*R*,8a*S*)-4-hydroxy-2,2-dimethyl-3a,3b,5,6,8,8a-hexahydro-4*H*-indeno[1,2-*d*][1,3]dioxol-8-yl)-1*H*-1,2,4-triazole-3-carboxylate (7a)

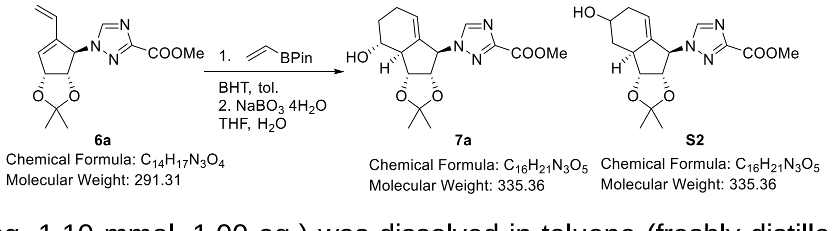

Diene **6a** (320 mg, 1.10 mmol, 1.00 eq.) was dissolved in toluene (freshly distilled over CaH$_2$, 15 mL) and BHT (29.0 mg, 0.11 mmol, 10 mol%) and vinyl boronic acid pinacol ester (477 µL, 2.82 mmol, 2.50 eq.) were added. The reaction was heated to 135 °C (oil bath temperature) in a sealed tube for 96 h. Then the volatiles was removed and the crude mixture was dissolved in THF/H$_2$O (20 mL, 1:1). Then, NaBO$_3$ x 4 H$_2$O (866 mg, 3.56 mmol, 3.00 eq.) was added. After 90 min, the reaction was completed and diluted with EtOAc/brine (1:1, 20 mL). After the separation of the organic phase, the aqueous phase was extracted with EtOAc (3x 20 mL). The combined organic phases were dried over MgSO$_4$ and concentrated *in vacuo*. Purification by column chromatography (SiO$_2$, DCM:EtOAc

1:1 to EtOAc/DCM 1:3 to pure EtOAc ) gave **7a** as a white solid (114 mg, 0.34 mmol, 31%) and **S2** (179 mg, 0.53 mmol, 49%) as a mixture of isomers.

**7a**: **R$_f$** = 0.30 (EtOAc:DCM 1:1); **$^{1}$H-NMR** (400 MHz, $CDCl_3$) δ [ppm] = 8.25 (s, 1H), 5.20 (brs, 1H), 5.06 (dd, *J* = 5.6, 7.7 Hz, 1H), 4.92 (m, 1H), 4.64 (dd, *J* = 5.0, 7.7 Hz, 1H), 4.00 (s, 3H), 3.69 (m, 1H), 2.63 (m, 1H), 2.11-2.20 (m, 2H), 1.95-2.05 (m, 2H), 1.52-1.65 (m, 1H), 1.56 (s, 3H), 1.33 (s, 3H); **$^{13}$C{$^{1}$H}-NMR** (101 MHz, $CDCl_3$) δ [ppm] = 160.2, 155.6, 145.7, 136.4, 122.5, 114.5, 83.3, 82.3, 71.5, 68.6, 53.0, 52.5, 31.3, 27.3, 25.04, 24.97; **HRMS** (ESI, 3.5kV) calc for $[M+Na]^+$ $C_{16}H_{21}N_3O_5Na^+$ 358.13734, found 358.13714.

**S2**: **R$_f$** = 0.16 (EtOAc:DCM 1:1).

#### Synthesis of 1-((1*R*,2*S*,3*R*,3a*R*,4*R*)-2,3,4-trihydroxy-2,3,3a,4,5,6-hexahydro-1*H*-inden-1-yl)-1*H*-1,2,4-triazole-3-carboxamide (10a)

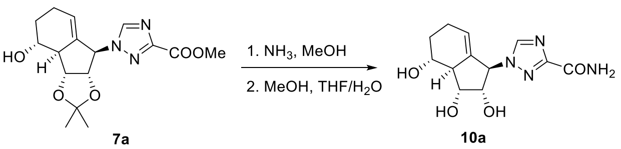

A solution of **7a** (26.0 mg, 77.5 μmol) in MeOH (0.5mL) was prepared and $NH_3$ (7M in MeOH, 0.5 mL) was added. The solution was stirred for 16 h and the volatiles was removed. Then after the crude was dissolved in MeOH (0.5 mL), TFA (10% in $H_2O$, 0.5 mL) was added and the reaction was stirred for another 23 h. Removal of the volatiles was followed by purification by column chromatography ($SiO_2$, EtOAc 5% to 10% MeOH) yielded nucleoside analogue **10a** (17.8 mg, 63.5 μmol, 82% over 2 steps) as a white solid.

**R$_f$** = 0.08 (EtOAc, 5% MeOH); $[\alpha]_D^{25}$ = -37.1 (c = 0.10, MeOH); **$^{1}$H-NMR** (400 MHz, MeOD-d4) δ [ppm] = 8.60 (s, 1H), 5.25 (brs, 1H), 5.03 (m, 1H), 4.25 (dd, *J* = 6.2, 6.8 Hz, 1H), 4.16 (dd, *J* = 5.0, 5.6 Hz, 1H), 3.57 (ddd, *J* = 3.8, 9.7, 11.3 Hz, 1H), 2.48 (m, 1H), 2.08-2.28 (m, 2H), 1.91-2.00 (m, 1H), 1.57 (ddt, *J* = 7.3, 11.3, 11.8 Hz, 1H); **$^{13}$C{$^{1}$H}-NMR** (101 MHz, $CDCl_3$) δ [ppm] = 163.5, 158.1, 147.1, 135.9, 122.7, 76.6, 73.8, 71.8, 68.6, 53.7, 32.3, 26.1 ; **HRMS** (ESI, 3.5kV) calc for $[M+Na]^+$ $C_{12}H_{16}N_4O_4Na^+$ 303.10638, found 303.10631.

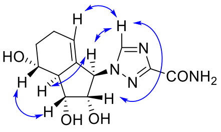

NOESY interactions
**10a**

#### Synthesis of (1*R*,2*S*,3*R*,3a*S*,4*R*)-1-(3-carbamoyl-1*H*-1,2,4-triazol-1-yl)-2,3-dihydroxy-2,3,3a,4,5,6-hexahydro-1*H*-inden-4-yl isobutyrate (10c)

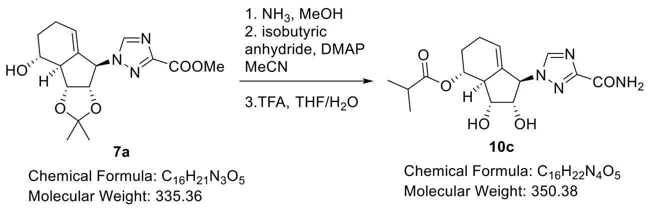

Alcohol **7a** (32 mg, 95.5 µmol, 1.00 eq.) was dissolved in MeOH (0.35 mL) and $NH_3$ (7M in MeOH, 0.35 mL) was added. After 16 h, the solvent was removed *in vacuo*. The residue was dissolved in dry ACN (1.5 mL) and at 0 °C DMAP (14.6 mg, 0.12 mmol, 1.25 eq.) and isobutyric anhydride (22.7 mg, 24 µL, 1.50 eq.) were added. After 20 min, the reaction was quenched with sat. $NaHCO_3$ (5 mL) and ethyl acetate (5 mL). The organic phase was separated, and the aqueous phase was extracted with ethyl acetate (4x 5 mL). The combined organic phases were dried over $MgSO_4$, and the solvent was removed *in vacuo*. Purification by column chromatography ($SiO_2$, DCM:EtOAc 3:1) gave the butyric ester which was directly deprotected by redissolving in MeOH (0.5 mL) and addition of TFA (10% in $H_2O$, 0.5 mL). After 24 h the solvent was removed *in vacuo*. Purification by column chromatography ($SiO_2$, EtOAc 5% MeOH) gave nucleoside analogue **10c** (18.6 mg, 53.1 µmol, 55% over 3 steps) as a white solid.

**$R_f$** = 0.33 (EtOAc 5% MeOH); $[\alpha]_D^{25}$ = -55.4 (c = 0.10, MeOH); **$^1$H-NMR** (400 MHz, $CDCl_3$) δ [ppm] = 8.63 (s, 1H), 5.25 (s, 1H), 5.17 (m, 1H), 4.91 (m, 1H), 4.22 (dd, *J* = 5.6, 5.8 Hz, 1H), 4.08 (dd, *J* = 5.9, 6.0 Hz, 1H), 2.74 (m, 1H), 2.61 (sept, *J* = 6.9 Hz, 1H), 2.14-2.33 (m, 2H), 1.93-2.03 (m, 1H), 1.68 (ddt, *J* = 7.3, 11.4, 11.4 Hz, 1H), 1.21 (d, *J* = 7.2 Hz, 3H), 1.91 (d, *J* = 7.1 Hz, 3H); **$^{13}$C{$^1$H}-NMR** (101 MHz, $CDCl_3$) δ [ppm] = 178.7, 157.6,# 146.9*, 135.6, 123.8, 76.7, 74.4, 74.3, 68.5, 50.5, 35.4, 28.9, 25.8, 19.3 (2x) ; **HRMS** (ESI, 3.5kV) calc for $[M+Na]^+$ $C_{15}H_{22}N_4O_5Na^+$ 373.14824, found 373.14822.

*# found by HMBC, * found by HSQC, note: carbamoyl carbon was not observed.*

**Synthesis of Methyl 1-((3a*R*,3b*S*,5*R*,8*R*,8a*S*)-5-(isobutyryloxy)-2,2-dimethyl-3a,3b,5,6,8,8a-hexahydro-4*H*-indeno[1,2-*d*][1,3]dioxol-8-yl)-1*H*-1,2,4-triazole-3-carboxylate (8a) and methyl 1-((3a*R*,3b*S*,5*S*,8*R*,8a*S*)-5-(isobutyryloxy)-2,2-dimethyl-3a,3b,5,6,8,8a-hexahydro-4*H*-indeno[1,2-*d*][1,3]dioxol-8-yl)-1*H*-1,2,4-triazole-3-carboxylate (9a)**

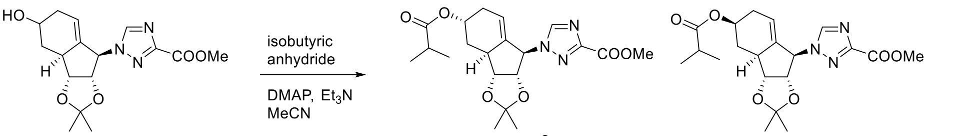

**S2**
Chemical Formula: $C_{16}H_{21}N_3O_5$
Molecular Weight: 335.36

**8a**
Chemical Formula: $C_{20}H_{27}N_3O_6$
Molecular Weight: 405.45

**9a**
Chemical Formula: $C_{20}H_{27}N_3O_6$
Molecular Weight: 405.45

Diastereomeric mixture **S2** (175.5 mg, 0.52 mmol, 1.00 eq.) was dissolved in MeCN (6 mL) and isobutyric anhydride (173 µL, 1.04 mmol, 2.00 eq.) $Et_3N$ (145 µL, 1.04 mmol, 2.00 eq.) and DMAP (6.4 mg, 52.2 µmol, 10 mol%) were added subsequently. After 18 h, the reaction was quenched by the addition of sat. $NaHCO_3$ solution and diluted with EtOAc. The organic phase was separated, and the aqueous phase was extracted with ethyl acetate (3x10 mL). The combined organic phases were washed with brine (15 mL) dried over $MgSO_4$, and the solvent was removed *in vacuo*. Purification by column chromatography ($SiO_2$, Hex:EtOAc 1:1) gave **8a** (48.1 mg, 11.8 µmol, 23%) and **9a** (108.8 mg, 26.8 µmol, 52%) both as white solids.

**8a**: $\mathbf{R_f}$ = 0.49 (Hex:EtOAc 1:2); **$^1$H-NMR** (600 MHz, $CDCl_3$) δ [ppm] = 8.26 (s, 1H), 5.25 (brs, 1H), 5.21 (brs, 1H), 5.08 (dd, *J* = 5.4, 7.3 Hz, 1H), 4.88 (m, 1H), 4.43 (dd, *J* = 7.4, 9.3 Hz, 1H), 3.99 (s, 3H), 2.27 (brs, 1H), 2.50 (hep, *J* = 7.0 Hz, 1H), 2.44 (ddd, *J* = 4.0, 5.3, 13.6 Hz, 1H), 2.32-2.35 (m, 1H), 2.09 (d, *J* = 19.1 Hz, 1H), 1.55 (s, 3H), 1.46 (ddd, *J* = 2.2, 11.9, 13.6 Hz, 1H), 1.31 (s, 3H), 1.14 (d, *J* = 7.0 Hz, 3H), 1.12 (d, *J* = 7.0 Hz, 3H); **$^{13}$C{$^1$H}-NMR** (151 MHz, $CDCl_3$) δ [ppm] = 176.6, 160.2, 155.7, 145.7, 138.4, 119.1, 114.2, 83.5, 83.3, 68.5, 66.7, 53.0, 39.3, 34.2, 30.5, 29.8, 27.4, 25.1, 19.12, 19.05; **HRMS** (ESI, 3.5kV) calc for $[M+Na]^+$ $C_{20}H_{27}N_3O_6Na^+$ 428.17921, found 428.17857.

**9a**: $\mathbf{R_f}$ = 0.56 (Hex:EtOAc 1:2); **$^1$H-NMR** (600 MHz, $CDCl_3$) δ [ppm] = 8.25 (s, 1H), 5.19 (brs, 1H), 5.05 (ddd, *J* = 1.0, 5.1, 7.2 Hz, 1H), 4.95 (m, 1H), 4.91 (m, 1H), 4.41 (ddd, *J* = 0.9, 5.6, 7.1 Hz, 1H), 3.99 (s, 3H), 2.80 (brs, 1H), 2.50 (hep, *J* = 7.0 Hz, 1H), 2.35-2.43 (m, 2H), 1.94-2.03 (m, 1H), 1.51 (m, 3H), 1.47 (q, *J* = 12.0 Hz, 1H), 1.29 (s, 3H), 1.13 (m, 6H); **$^{13}$C{$^1$H}-NMR** (151 MHz, $CDCl_3$) δ [ppm] = 176.7, 160.2, 155.7, 145.7, 138.5, 120.5, 114.1, 83.5, 82.8, 68.9, 67.9, 53.0, 44.2, 34.1, 32.4, 30.2, 27.4, 25.1, 19.04, 19.02; **HRMS** (ESI, 3.5kV) calc for $[M+Na]^+$ $C_{20}H_{27}N_3O_6Na^+$ 428.17921, found 428.17857.

#### Synthesis of 1-((1*R*,2*S*,3*R*,3a*S*,5*R*)-2,3,5-trihydroxy-2,3,3a,4,5,6-hexahydro-1*H*-inden-1-yl)-1*H*-1,2,4-triazole-3-carboxamide (11a)

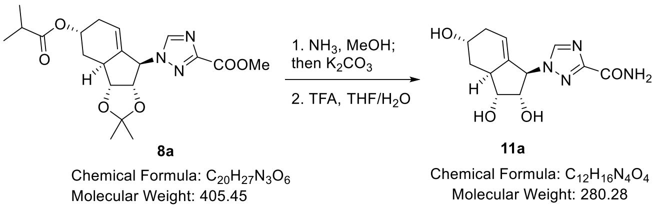

Ester **8a** (24.8 mg, 61.2 µmol, 1.00 eq.) was dissolved in MeOH (0.5 mL) and $NH_3$ (7M in MeOH, 0.5 mL) was added. The reaction was stirred for 16 h and then $K_2CO_3$ (16.9 mg, 0.12 mmol, 2.00 eq.) was added. After further 16 h stirring, a brine/EtOAc (10 mL, 1:1) mixture was added. The organic phase was separated, and the aqueous phase was extracted with EtOAc (4x 4 mL). The combined organic phases were dried over $MgSO_4$ and the solvent was removed *in vacuo*. The crude was redissolved in MeOH (0.5 mL) and TFA (10% in $H_2O$, 0.5 mL) was added. The reaction was stirred for 24 h, then the solvent was removed *in vacuo*. Purification by column chromatography ($SiO_2$, EtOAc 7% MeOH) gave nucleoside analogue **11a** (16.0 mg, 57.1 µmol, 93% over 3 steps) as a white solid.

$\mathbf{R_f}$ = 0.21 (EtOAc, 5% MeOH); $[\alpha]_D^{25}$ = +6.2 (c = 0.12, MeOH); **$^1$H-NMR** (400 MHz, MeOD-d4) δ [ppm] = 8.60 (s, 1H), 5.24 (brs, 1H), 5.11 (m, 1H), 4.25 (dd, *J* = 4.1, 5.7 Hz, 1H), 4.18 (m, 1H), 3.96 (dd, *J* = 5.7, 8.0 Hz, 1H), 2.80 (brs, 1H), 2.23-2.34 (m, 1H), 2.09 (d, *J* = 19.1 Hz, 1H), 1.40 (ddd, *J* = 1.8, 11.8, 13.3 Hz, 1H); **$^{13}$C{$^1$H}-NMR** (101 MHz, MeOD-d4) δ [ppm] = 163.5, 158.2, 148.9, 138.1, 121.2, 77.1, 76.9, 69.1, 65.0, 39.3, 34.2, 34.0; **HRMS** (ESI, 3.5kV) calc for $[M+H]^+$ $C_{12}H_{16}N_4O_4H^+$ 281.12443, found 281.12393.

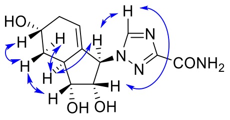

NOESY interactions
**11a**

#### Synthesis of (1*R*,2*S*,3*R*,3a*S*,5*R*)-1-(3-carbamoyl-1*H*-1,2,4-triazol-1-yl)-2,3-dihydroxy-2,3,3a,4,5,6-hexahydro-1*H*-inden-5-yl isobutyrate (11c)

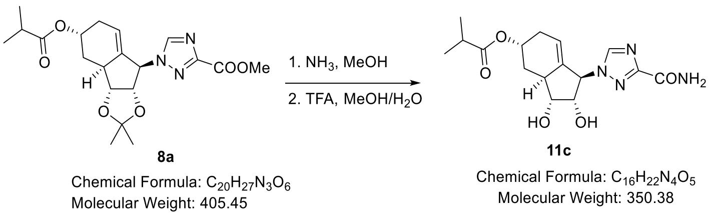

Ester **8a** (20.1 mg, 49.6 μmol) was dissolved in MeOH (0.5 mL) and $NH_3$ (7M in MeOH, 0.5 mL) was added. The reaction was stirred for 16h and then the volatiles was removed *in vacuo*. The crude was redissolved in MeOH (0.75 mL) and TFA (10% in $H_2O$, 0.75 mL) was added. After 24h the solvent was removed *in vacuo*. Purification by column chromatography ($SiO_2$, EtOAc 5% MeOH) gave nucleoside analogue **11c** (16.2 mg, 46.2 μmol, 93% over 2 steps) as a white solid.

**$R_f$** = 0.17 (EtOAc, 5% MeOH); $[\alpha]_D^{25}$ = +3.2 (c = 0.12, MeOH); **$^1$H-NMR** (400 MHz, $CDCl_3$) δ [ppm] = 8.70 (s, 1H), 5.28 (brs, 1H), 5.23 (brs, 1H), 5.17 (brs, 1H), 4.26 (dd, *J* = 4.3, 5.1 Hz, 1H), 3.99 (dd, *J* = 5.8, 8.0 Hz, 1H), 2.77 (brs, 1H), 2.52 (sept, *J* = 7.0 Hz, 1H), 2.31-2.45 (m, 2H), 2.16 (d, *J* = 19.3 Hz, 1H), 1.48 (t, *J* = 12.5 Hz, 1H), 1.16 (d, *J* = 7.0 Hz, 3H, 1.13 (d, *J* = 7.0 Hz, 3H) ; **$^{13}$C{$^1$H}-NMR** (101 MHz, $CDCl_3$) δ [ppm] = # 178.3, 138.3, 121.1, 77.1, 77.0, 69.0, 68.8, 39.7, 35.4, 31.4, 19.34, 19.26; **HRMS** (ESI, 3.5kV) calc for $C_{16}H_{22}N_4O_5Na^+$ [M+Na]$^+$ 373.14824, found 373.14763.
### not all carbon signals could be found due to broad signals

#### Synthesis of 1-((1*R*,2*S*,3*R*,3a*S*,5*S*)-2,3,5-trihydroxy-2,3,3a,4,5,6-hexahydro-1*H*-inden-1-yl)-1*H*-1,2,4-triazole-3-carboxamide (12a)

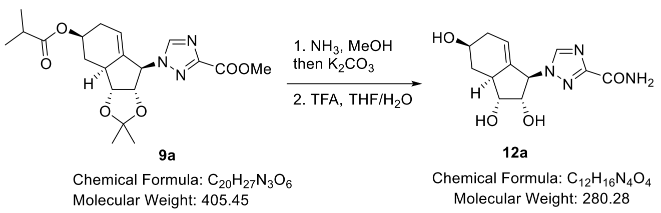

Ester **9a** (21.6 mg, 53.3 μmol, 1.00 eq.) was dissolved in MeOH (0.5 mL) and $NH_3$ (7M in MeOH, 0.5 mL) was added. The reaction was stirred for 17.5h and then $K_2CO_3$ (14.7 mg, 0.11 mmol, 2.00 eq.) was added. After a further 24h, $H_2O$/EtOAc (10 mL, 1:1) was added. The organic phase was separated and the aqueous phase was extracted with EtOAc (4x 4 mL). The combined organic phases were dried over $MgSO_4$ and the solvent was removed *in vacuo*. The crude was redissolved in MeOH (0.5 mL) and TFA (10% in $H_2O$, 0.5 mL) was added. The reaction was stirred for 27h, then the solvent was removed *in vacuo*. Purification by column chromatography ($SiO_2$, EtOAc 7.5% MeOH) gave nucleoside analogue **12a** (10.8 mg, 38.5 μmol, 72% over 3 steps) as a white solid.

**$R_f$** = 0.08 (EtOAc, 7.5% MeOH); $[\alpha]_D^{25}$ = +4.9 (c = 0.13, MeOH); **$^1$H-NMR** (400 MHz, MeOD-d4) δ [ppm] = 8.59 (s, 1H), 5.19 (s, 1H), 5.16 (m, 1H), 4.24 (dd, *J* = 3.8, 5.5 Hz, 1H), 3.95 (dd, *J* = 5.9, 8.3 Hz, 1H), 3.90 (m, 1H), 2.70 (m, 1H), 2.37-2.46 (m, 1H), 2.34 (ddd, *J* = 3.6, 5.2, 11.6 Hz, 1H), 1.83-1.94 (m, 1H), 1.34 (q, *J* = 11.6 Hz, 1H); **$^{13}$C{$^1$H}-NMR** (101 MHz, $CDCl_3$) δ [ppm] = 163.5, 158.3, 146.9, 138.3, 122.6, 77.5, 76.6, 68.6, 68.2, 45.3, 37.2, 35.4 ; **HRMS** (ESI, 3.5kV) calc for [M+H]$^+$ $C_{12}H_{16}N_4O_4H^+$ 281.12375, found 281.12443.

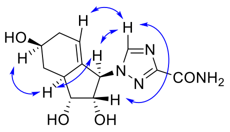

NOESY interactions
**12a**

#### Synthesis of (1*R*,2*S*,3*R*,3a*S*,5*S*)-1-(3-carbamoyl-1*H*-1,2,4-triazol-1-yl)-2,3-dihydroxy-2,3,3a,4,5,6-hexahydro-1*H*-inden-5-yl isobutyrate (12c)

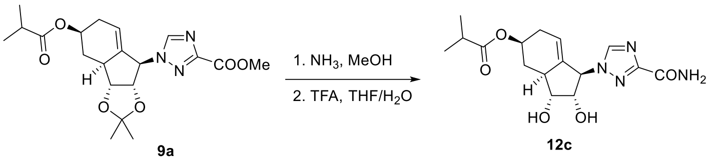

**9a** — Chemical Formula: $C_{20}H_{27}N_3O_6$, Molecular Weight: 405.45

**12c** — Chemical Formula: $C_{16}H_{22}N_4O_5$, Molecular Weight: 350.38

Ester **9a** (20.8 mg, 51.3 µmol) was dissolved in MeOH (0.5 mL) and $NH_3$ (7M in MeOH, 0.5 mL) was added. The reaction was stirred for 17.5h and then the volatiles were removed *in vacuo*. The crude was redissolved in MeOH (0.5 mL) and TFA (10% in $H_2O$, 0.5 mL) was added. After 24h the solvent was removed *in vacuo*. Purification by column chromatography ($SiO_2$, EtOAc 3% MeOH) gave nucleoside analogue **12c** (15.7 mg, 44.8 µmol, 87% over 2 steps) as a white solid.

**R$_f$** = 0.37 (EtOAc, 5% MeOH); $[\alpha]_D^{25}$ = +5.9 (c = 0.09, MeOH); **$^1$H-NMR** (600 MHz, $CDCl_3$) δ [ppm] = 8.61 (s, 1H), 5.22 (brs, 1H), 5.19 (m, 1H), 5.00 (dddd, *J* = 3.8, 6.4, 9.7, 12.3, 1H), 4.24 (dd, *J* = 3.5, 5.6 Hz, 1H), 3.97 (dd, *J* = 5.7, 8.4 Hz, 1H), 2.77 (brs, 1H), 2.54 (hep, *J* = 7.0 Hz, 1H), 2.46-2.51 (m, 1H), 2.37 (ddd, *J* = 3.5, 5.4, 11.4 Hz, 1H), 1.99-2.06 (m, 1H), 1.47 (q, *J* = 11.5 Hz, 1H), 1.15 (d, *J* = 7.0 Hz, 3H), 1.14 (d, *J* = 7.0 Hz, 3H) ; **$^{13}$C{$^1$H}-NMR** (151 MHz, $CDCl_3$) δ [ppm] = 178.4, 163.4, 158.3, 147.0, 138.5, 122.1, 77.5, 76.6, 71.1, 68.9, 44.6, 35.3, 33.4, 31.9, 19.3 (2x); **HRMS** (ESI, 3.5kV) calc for $[M+Na]^+$ $C_{16}H_{22}N_4O_5Na^+$ 373.14824, found 373.14768.

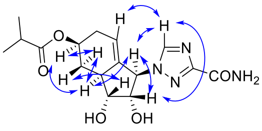

NOESY interactions
**12c**

#### Synthesis of Methyl 1-((3a*S*,4*S*,6a*R*)-2,2-dimethyl-5-vinyl-3a,6a-dihydro-4*H*-cyclopenta[*d*][1,3]dioxol-4-yl)-1H-1,2,4-triazole-5-carboxylate (6b)

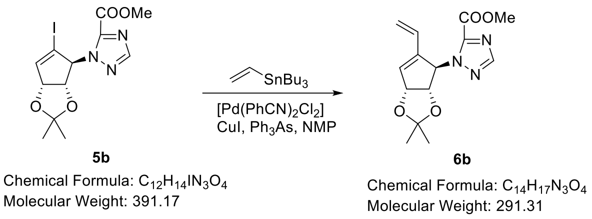

Iodide **5b** (400 mg, 1.02 mmol, 1.00 eq.) was dissolved in NMP (2 mL, freshly distilled) and [Pd(PhCN)$_2$Cl$_2$] (11.8 mg, 30.7 µmol, 3 mol%), CuI (11.7 mg, 61.4 µmol, 6 mol%) and Ph$_3$As (18.8 mg, 61.4 µmol, 6 mol%) was added. Then at 0 °C vinyl tributyltin (330 µL, 1.12 mmol, 1.10 eq.) was added dropwise. The reaction was stirred at this temperature for 1h. The reaction was quenched with sat. NaHCO$_3$ solution (10 mL) and EtOAc (10 mL). After the separation of the organic phase, the aqueous phase was extracted with EtOAc (3x 10 mL). The combined organic phases were washed with brine (15 mL), dried over MgSO$_4$ and concentrated *in vacuo*. Purification by column chromatography (SiO$_2$, Hex:EtOAc 2:1 to 1:1) gave **6b** as a yellow gum (290 mg, 1.00 mmol, 98%). **R$_f$** = 0.18 (Hex:EtOAc 3:1); **$^1$H-NMR** (400 MHz, CDCl$_3$) δ [ppm] = 7.97 (s, 1H), 6.71 (s, 1H), 6.42 (dd, *J* = 11.0, 17.9 Hz, 1H), 6.16 (s, 1H), 5.50 (d, *J* = 5.7 Hz, 1H), 5.13 (d, *J* = 11.9 Hz, 1H), 5.12 (d, *J* = 17.0 Hz, 1H), 4.61 (d, *J* = 5.6 Hz, 1H), 4.07 (s, 3H), 1.47 (s, 3H), 1.35 (s, 3H); **$^{13}$C{$^1$H}-NMR** (101 MHz, CDCl$_3$) δ [ppm] = 158.7, 151.5, 144.5, 140.7, 131.1, 130.3, 118.9, 112.3, 84.9, 84.1, 68.8, 53.6, 27.6, 26.2; **HRMS** (ESI, 3.5kV) calc for [M+H]$^+$ $C_{14}H_{17}N_3O_4H^+$ 292.12918, found 292.12868.

#### Synthesis of Methyl 1-((3a*R*,3b*R*,4*R*,8*R*,8a*S*)-4-hydroxy-2,2-dimethyl-3a,3b,5,6,8,8a-hexahydro-4*H*-indeno[1,2-*d*][1,3]dioxol-8-yl)-1*H*-1,2,4-triazole-5-carboxylate (7b)

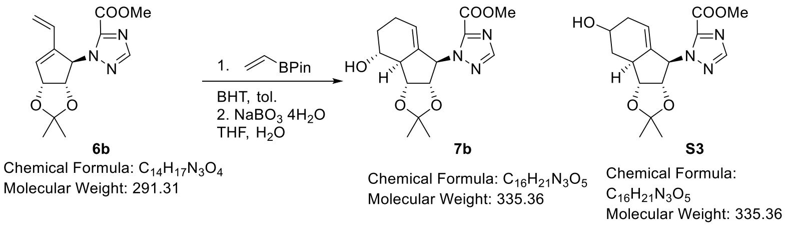

Diene 6b (291 mg, 1.00 mmol, 1.00 eq.) and BHT (25.8 mg, 0.10 mmol, 10 mol%) were dissolved in toluene (10 mL) and then vinyl pinacol borate (0.42 mL, 2.50 mmol, 2.50 eq.) was added. The reaction was stirred in a sealed tube for 3d at 140°C (oil bath temperature). Then, the solvent was removed and the residue was dissolved in THF (10 mL) and ph7 buffer (10 mL). Then NaBO$_3$ 4 H$_2$O (614 mg, 4.00 mmol, 4.00 eq.) was added and the reaction was stirred for 1 h. Then saturated Na$_2$S$_2$O$_3$ solution (10 mL) and EtOAc (10 mL) were added. After phase separation, the aqueous phase was extracted with EtOAc (3x15 mL). The combined organic phases were dried over $MgSO_4$ and concentrated *in vacuo*. Purification by column chromatography ($SiO_2$, Hex:EtOAc 1:1 to 1:3 to EtOAc) gave **7b** as a pure compound (104 mg, 0.31 mmol, 31%) and **S3** as a diastereomeric mixture (140 mg, 0.42 mmol, 42%).

**7b**: **R$_f$** = 0.21 (Hex:EtOAc 1:1); **$^1$H-NMR** (600 MHz, $CDCl_3$) δ [ppm] = 8.03 (s, 1H), 6.25 (s, 1H), 5.11 (dd, *J* = 5.1, 7.3 Hz, 1H), 4.77 (s, 1H), 4.67 (dd, *J* = 5.0, 7.1 Hz, 1H), 4.00 (s, 3H), 3.70 (dd, *J* = 10.5, 10.7 Hz, 1H), 2.66 (brs, 1H), 2.05-2.21 (m, 2H), 1.90-2.02 (m, 2H), 1.56-1.71 (m, 1H), 1.58 (s, 3H), 1.35 (s, 3H); **$^{13}$C{$^1$H}-NMR** (151 MHz, $CDCl_3$) δ [ppm] = 158.4, 151.4, 145.3, 137.3, 121.3, 114.1, 84.0, 82.6, 71.7, 67.8, 53.4, 52.9, 31.3, 27.6, 25.4, 25.0 ; **HRMS** (ESI, 3.5kV) calc for $[M+Na]^+$ $C_{16}H_{21}N_3O_5Na^+$ 358.13734, found 358.13681.

**S3**: **R$_f$** = 0.18 (Hex:EtOAc 1:1).

#### Synthesis of 1-((1*R*,2*S*,3*R*,3a*R*,4*R*)-2,3,4-trihydroxy-2,3,3a,4,5,6-hexahydro-1*H*-inden-1-yl)-1*H*-1,2,4-triazole-5-carboxamide (10b)

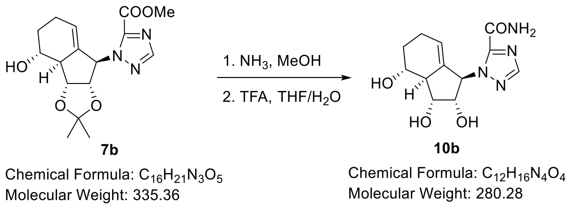

Ester **7b** (78.7 µmol, 26.4 mg) was dissolved in MeOH (0.3 mL) and $NH_3$ (7M in MeOH, 0.3 mL) was added. The reaction was stirred for 16h and the solvent was then removed *in vacuo.* The crude was redissolved in MeOH (1 mL) and TFA (50 µL) was added. After 30h the solvent was removed and the remaining TFA was removed by co-evaporation with MeOH (3x 2 mL). Purification by column chromatography ($SiO_2$, EtOAc with 3% MeOH) gave **10b** as a white solid (21.1 mg, 75.2 µmol, 95%).

**R$_f$** = 0.23 (EtOAc/MeOH 3%); $[\alpha]_D^{25}$ = -64.5 (c = 0.15, MeOH) **$^1$H-NMR** (400 MHz, $CDCl_3$) δ [ppm] = 7.98 (s, 1H), 6.28 (s, 1H), 4.91 (m, 1H), 4.33 (dd, *J* = 5.7, 5.7 Hz, 1H), 4.19 (dd, *J* = 5.9, 6.0 Hz, 1H), 3.56 (ddd, *J* = 3.7, 9.2, 11.4 Hz, 1H), 2.51 (brs, 1H), 2.14-2.24 (m, 1H), 2.00-2.13 (m, 1H), 1.89-1.97 (m, 1H), 1.57 (ddt, *J* = 6.9, 11.4, 11.5, 1H); **$^{13}$C{$^1$H}-NMR** (101 MHz, $CDCl_3$) δ [ppm] = 161.0, 151.2, 148.8, 136.6, 122.3, 76.8, 75.2, 72.6, 68.1, 53.1, 32.3, 26.1; **HRMS** (ESI, 3.5kV) calc for $[M+Na]^+$ $C_{12}H_{16}N_4O_4Na^+$ 303.10638, found 303.10612.

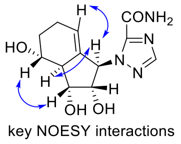

key NOESY interactions
**10b**

#### Synthesis of (1*R*,2*S*,3*R*,3a*S*,4*R*)-1-(5-carbamoyl-1*H*-1,2,4-triazol-1-yl)-2,3-dihydroxy-2,3,3a,4,5,6-hexahydro-1*H*-inden-4-yl isobutyrate (10d)

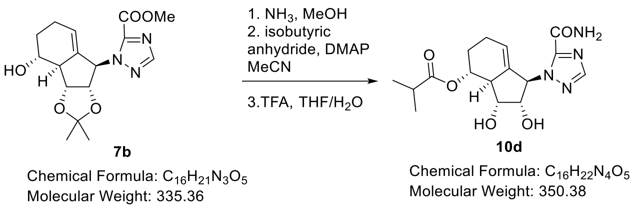

Alcohol **7b** (20.0 mg, 59.6 µmol, 1.00 eq.) was dissolved in MeOH (0.5 mL) and $NH_3$ (7M in MeOH, 0.5 mL) was added. After 17 h the solvent was removed *in vacuo*. The residue was dissolved in dry ACN (1.5 mL) and at 0 °C DMAP (9.1 mg, 74.5 µmol, 1.25 eq.) and *iso*-butyric anhydride (14.2 mg, 15 µL, 1.50 eq.) were added. After 2 h, the reaction was quenched with sat. $NaHCO_3$ (5 mL) and ethyl acetate (5 mL). The organic phase was separated, and the aqueous phase was extracted with ethyl acetate (4x 5 mL). The combined organic phases were dried over $MgSO_4$, and the solvent was removed *in vacuo*. Purification by column chromatography ($SiO_2$, DCM:EtOAc 3:1) gave the butyric ester which was directly deprotected by redissolving in MeOH (0.75 mL) and addition of TFA (10% in $H_2O$, 0.25 mL). After 40 h the solvent was removed *in vacuo*. Purification by column chromatography ($SiO_2$, EtOAc 3% MeOH) gave nucleoside analogue **10d** (19.8 mg, 56.5 µmol, 95% over 3 steps) as a white solid.

**$R_f$** = 0.54 (EtOAc, 5% MeOH); $[\alpha]_D^{25}$ = -66.4 (c = 0.10, MeOH); **$^1$H-NMR** (400 MHz, $CDCl_3$) δ [ppm] = 7.98 (s, 1H), 6.29 (s, 1H), 5.04 (m, 1H), 4.87 (m, 1H), 4.27 (dd, *J* = 4.4, 5.5 Hz, 1H), 4.14 (dd, *J* = 5.9, 7.4 Hz, 1H), 2.75 (m, 1H), 2.60 (sept, *J* = 7.0 Hz, 1H), 2.19-2.29 (m, 1H), 2.07-2.19 (m, 1H), 1.89-1.98 (m, 1H), 1.67 (ddt, *J* = 6.8, 11.6, 11.9 Hz, 1H), 1.20 (d, *J* = 7.0 Hz, 3H), 1.18 (d, *J* = 7.0 Hz, 3H); **$^{13}$C{$^1$H}-NMR** (101 MHz, $CDCl_3$) δ [ppm] = 178.7, 161.0, 151.3, 148.5, 136.5, 123.2, 77.0, 75.7, 74.8, 67.9, 50.0, 35.4, 28.9, 25.8, 19.4, 19.3; **HRMS** (ESI, 3.5kV) calc for $[M+Na]^+$ $C_{16}H_{22}N_4O_5Na^+$ 373.14824, found 373.14786.

**Synthesis of Methyl 1-((3a*R*,3b*S*,5*R*,8*R*,8a*S*)-5-(isobutyryloxy)-2,2-dimethyl-3a,3b,5,6,8,8a-hexahydro-4*H*-indeno[1,2-*d*][1,3]dioxol-8-yl)-1*H*-1,2,4-triazole-5-carboxylate (8b) and methyl 1-((3a*R*,3b*S*,5*S*,8*R*,8a*S*)-5-(isobutyryloxy)-2,2-dimethyl-3a,3b,5,6,8,8a-hexahydro-4*H*-indeno[1,2-*d*][1,3]dioxol-8-yl)-1*H*-1,2,4-triazole-5-carboxylate (9b)**

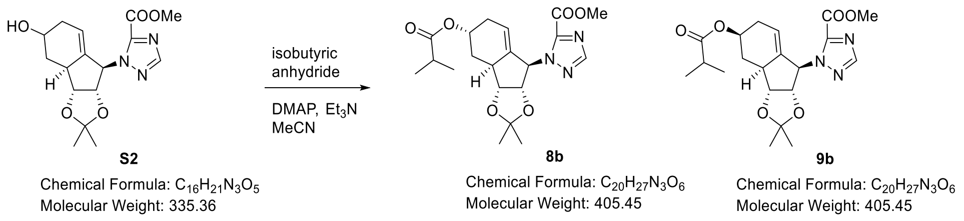

Alcohol **S2** (47.3 mg, 0.14 mmol, 1.00 eq.) was dissolved in acetonitrile (5 mL) and isobutyric anhydride (35 µL, 0.21 mmol, 1.50 eq.), $Et_3N$ (30 µL, 0.21 mmol, 1.50 eq.) and DMAP (1.7 mg, 14.1 µmol, 10 mol%) were added. The reaction was stirred for 19 h. Then acetonitrile was removed and the crude was redissolved in EtOAc/sat. $NaHCO_3$ (1:1, 16 mL). After phase separation, the aqueous phase was extracted with EtOAc (3x10 mL). The combined organic phases were washed with brine (20 mL), dried over $MgSO_4$ and concentrated *in vacuo*. Purification by column chromatography ($SiO_2$, Hex:EtOAc 3:2) gave first **9b** (36.2 mg, 89.3 µmol, 63%) and then **8b** (17.9 mg, 44.1 µmol, 31%).

**8b**: $\mathbf{R_f}$ = 0.40 (Hex:EtOAc 1:1); **$^1$H-NMR** (400 MHz, $CDCl_3$) δ [ppm] = 8.04 (s, 1H), 6.33 (brs, 1H), 5.22 (brs, 1H), 5.09 (dd, *J* = 4.9, 7.1 Hz, 1H), 4.76 (m, 1H), 4.46 (dd, *J* = 5.5, 7.0 Hz, 1H), 4.01 (s, 3H), 2.81 (brs, 1H), 2.53 (hep, *J* = 6.9 Hz, 1H), 2.40-2.49 (m, 1H), 2.24-2.35 (m, 1H), 2.09 (d, *J* = 18.9 Hz, 1H), 1.58 (s, 3H), 1.49 (ddd, *J* = 1.7, 11.6, 15.1 Hz, 1H), 1.33 (s, 3H), 1.17 (d, *J* = 7.0 Hz, 3H), 1.14 (d, *J* = 6.9 Hz, 3H); **$^{13}$C{$^1$H}-NMR** (100 MHz, $CDCl_3$) δ [ppm] = 176.8, 158.4, 151.5, 145.3, 139.4, 118.1, 113.8, 84.1, 83.8, 67.6, 66.7, 53.4, 39.6, 34.2, 30.7, 29.9, 27.6, 25.4, 19.2, 19.1 ; **HRMS** (ESI, 3.5kV) calc for $[M+Na]^+$ $C_{20}H_{27}N_3O_6Na^+$ 406.19726, found 406.19643.

**9b**: $\mathbf{R_f}$ = 0.60 (Hex:EtOAc 1:1); **$^1$H-NMR** (600 MHz, $CDCl_3$) δ [ppm] = 8.03 (s, 1H), 6.23 (brs, 1H), 5.10 (dd, *J* = 4.7, 7.3 Hz, 1H), 4.97 (dddd, *J* = 3.5, 6.0, 9.6, 12.3 Hz, 1H), 4.76 (m, 1H), 4.43 (dd, *J* = 5.6, 7.0 Hz, 1H), 4.00 (s, 3H), 2.85 (brs, 1H), 2.51 (hep, *J* = 7.0 Hz, 1H), 2.35-2.45 (m, 2H), 1.92-1.99 (m, 1H), 1.55 (s, 3H), 1.50 (q, *J* = 11.6 Hz, 1H), 1.31 (s, 3H), 1.13 (d, *J* = 7.0 Hz, 3H), 1.12 (d, *J* = 7.0 Hz, 3H); **$^{13}$C{$^1$H}-NMR** (151 MHz, $CDCl_3$) δ [ppm] = 176.7, 158.3, 151.5, 145.3, 139.4, 119.3, 113.8, 84.1, 83.1, 69.1, 67.1, 53.4, 44.5, 34.2, 32.6, 30.3, 27.6, 25.5, 19.1, 19.0 ; **HRMS** (ESI, 3.5kV) calc for $[M+H]^+$ $C_{20}H_{27}N_3O_6H^+$ 406.19726, found 406.19646.

#### Synthesis of 1-((1*R*,2*S*,3*R*,3a*S*,5*R*)-2,3,5-trihydroxy-2,3,3a,4,5,6-hexahydro-1*H*-inden-1-yl)-1*H*-1,2,4-triazole-5-carboxamide (11b)

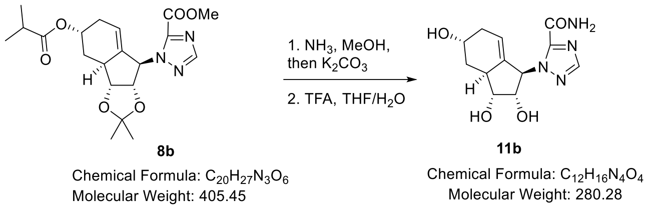

Precursor **8b** (16.1 mg, 39.7 μmol, 1.00 eq.) was dissolved in MeOH (0.5 mL) and $NH_3$ (7M in MeOH, 0.5 mL) was added. The reaction was stirred for 17h, and then $K_2CO_3$ (11.0 mg, 79.4 μmol, 2.00 eq.) was added. After a further 30h, $H_2O$/EtOAc (10 mL, 1:1) was added. The organic phase was separated and the aqueous phase was extracted with EtOAc (5x 4 mL). The combined organic phases were dried over $MgSO_4$ and the solvent was removed *in vacuo*. The crude was redissolved in MeOH (0.5 mL) and TFA (10% in $H_2O$, 0.5 mL) was added. The reaction was stirred for 19h, then the solvent was removed *in vacuo*. Purification by column chromatography ($SiO_2$, EtOAc 5% MeOH) gave nucleoside analogue **11b** (10.4 mg, 37.1 μmol, 94% over 3 steps) as a white solid.

**$R_f$** = 0.21 (EtOAc, 5% MeOH); $[\alpha]_D^{25}$ = -28.1 (c = 0.14, MeOH); **$^1$H-NMR** (600 MHz, $CDCl_3$) δ [ppm] = 7.95 (s, 1H), 6.29 (s, 1H), 4.97 (s, 1H), 4.32 (dd, *J* = 3.3, 5.6 Hz, 1H), 4.15 (brs, 1H), 4.02 (dd, *J* = 5.5, 8.9 Hz, 1H), 2.81 (brs, 1H), 2.28 (ddd, *J* = 3.7, 5.3, 12.9 Hz, 1H), 2.17-2.24 (m, 1H), 2.07 (d, *J* = 19.2 Hz, 1H), 1.34 (t, *J* = 11.9 Hz, 1H); **$^{13}$C{$^1$H}-NMR** (151 MHz, $CDCl_3$) δ [ppm] = 161.1, 151.3, 148.5, 138.8, 120.3, 77.8, 77.3, 68.5, 65.1, 39.1, 34.4, 30.0; **HRMS** (ESI, 3.5kV) calc for $[M+Na]^+$ $C_{12}H_{16}N_4O_4Na^+$ 303.10638, found 303.10603.

#### Synthesis of (1*R*,2*S*,3*R*,3a*S*,5*R*)-1-(5-carbamoyl-1*H*-1,2,4-triazol-1-yl)-2,3-dihydroxy-2,3,3a,4,5,6-hexahydro-1*H*-inden-5-yl isobutyrate (11d)

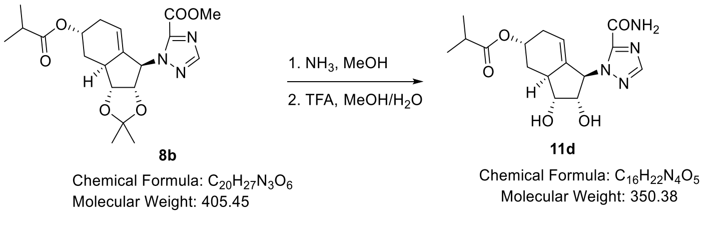

Ester **8b** (20.5 mg, 50.6 μmol) was dissolved in MeOH (0.5 mL) and $NH_3$ (7M in MeOH, 0.5 mL) was added. The reaction was stirred for 22h and then the volatiles were removed *in vacuo*. The crude was redissolved in MeOH (0.5 mL) and TFA (10% in $H_2O$, 0.5 mL) was added. After 22 h the solvent was removed *in vacuo*. Purification by column chromatography ($SiO_2$, EtOAc 3% MeOH) gave nucleoside analogue **11d** (15.2 mg, 43.4 μmol, 87% over 2 steps) as a white solid.

**R$_f$** = 0.53 (EtOAc, 7.5% MeOH); $[\alpha]_D^{25}$ = -30.3 (c = 0.11, MeOH); **$^1$H-NMR** (400 MHz, MeOD-d4) δ [ppm] = 7.96 (s, 1H), 6.31 (brs, 1H), 5.21 (brs, 1H), 5.03 (brs, 1H), 4.30 (dd, *J* = 2.8, 5.5 Hz, 1H), 4.04 (dd, *J* = 5.6, 9.2 Hz, 1H), 2.77 (brs, 1H), 2.54 (sept, *J* = 7.0 Hz, 1H), 2.40 (ddd, *J* = 4.5, 4.6, 13.3 Hz, 1H), 2.23-2.34 (m, 1H), 2.13 (d, *J* = 19.7 Hz, 1H), 1.40 (ddd, *J* = 1.5, 11.6, 13.2 Hz, 1H), 1.16 (d, *J* = 7.2 Hz, 3H), 1.14 (d, *J* = 7.2 Hz, 3H); **$^{13}$C{$^1$H}-NMR** (101 MHz, MeOD-d4) δ [ppm] = 177.1, 159.8, 150.1, 147.1, 137.9, 119.0, 76.6, 76.1, 67.6, 67.1, 38.1, 34.1, 30.3, 30.2, 18.1, 18.0; **HRMS** (ESI, 3.5kV) calc for [M+Na]$^+$ $C_{16}H_{22}N_4O_5Na^+$ 373.14824, found 373.14778.

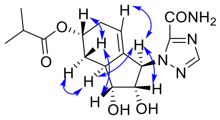

key NOESY interactions
**11d**

#### Synthesis of 1-((1*R*,2*S*,3*R*,3a*S*,5*S*)-2,3,5-trihydroxy-2,3,3a,4,5,6-hexahydro-1*H*-inden-1-yl)-1*H*-1,2,4-triazole-5-carboxamide (12b)

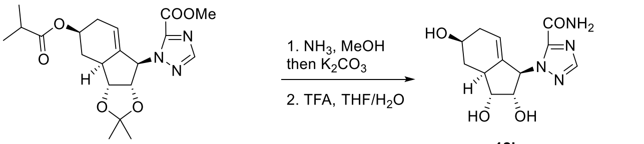

Ester **9b** (25.2 mg, 62.2 μmol, 1.00 eq.) was dissolved in MeOH (0.5 mL) and $NH_3$ (7M in MeOH, 0.5 mL) was added. The reaction was stirred for 16 h and then $K_2CO_3$ (17.2 mg, 0.12 mmol, 2.00 eq.). After further 16 h brine/EtOAc (10 mL, 1:1) was added. The organic phase was separated and the aqueous phase was extracted with EtOAc (4x 4 mL). The combined organic phases were dried over $MgSO_4$ and the solvent was removed *in vacuo*. The crude was redissolved in MeOH (0.5 mL) and TFA (10% in $H_2O$, 0.5 mL) was added. The reaction was stirred for 24 h, then the solvent was removed *in vacuo*. Purification by column chromatography ($SiO_2$, EtOAc 5% MeOH) gave nucleoside analogue **12b** (16.9 mg, 60.3 μmol, 97% over 3 steps) as a white solid.

**R$_f$** = 0.35 (EtOAc, 5% MeOH); $[\alpha]_D^{25}$ = -84.6 (c = 0.09, MeOH); **$^1$H-NMR** (400 MHz, $CDCl_3$) δ [ppm] = 7.96 (s, 1H), 6.23 (brs, 1H), 4.99 (m, 1H), 4.30 (dd, *J* = 2.6, 5.6 Hz, 1H), 3.99 (dd, *J* = 5.5, 9.2 Hz, 1H), 3.90 (m, 1H), 2.71 (brs, 1H), 2.28-2.44 (m, 2H), 1.81 (m, 1H), 1.27 (q, *J* = 11.5 Hz, 1H); **$^{13}$C{$^1$H}-NMR** (101 MHz, $CDCl_3$) δ [ppm] = 161.1, 151.3, 148.3, 139.1, 121.7, 77.8, 77.5, 68.2, 68.0, 45.1, 37.4, 35.5; **HRMS** (ESI, 3.5kV) calc for [M+Na]$^+$ $C_{12}H_{16}N_4O_4Na^+$ 303.10638, found 303.10603.

#### Synthesis of (1*R*,2*S*,3*R*,3a*S*,5*S*)-1-(5-carbamoyl-1*H*-1,2,4-triazol-1-yl)-2,3-dihydroxy-2,3,3a,4,5,6-hexahydro-1*H*-inden-5-yl isobutyrate (12d)

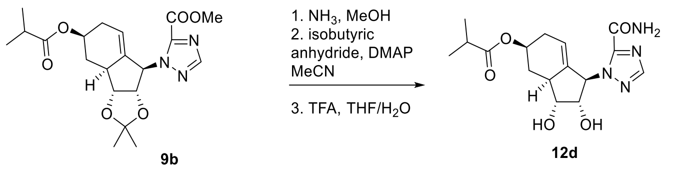

**9b**

Chemical Formula: $C_{20}H_{27}N_3O_6$
Molecular Weight: 405.45

**12d**

Chemical Formula: $C_{16}H_{22}N_4O_5$
Molecular Weight: 350.38

Precursor **9b** (28.0 mg, 69.1 μmol) was dissolved in MeOH (0.5 mL) and $NH_3$ (7M in MeOH, 0.5 mL) was added. The reaction was stirred for 17h, then the solvent was removed *in vacuo*. The crude was redissolved in MeOH (0.5 mL) and TFA (10% in $H_2O$, 0.5 mL) was added. The reaction was stirred for 25h, then the solvent was removed *in vacuo*. Purification by column chromatography ($SiO_2$, EtOAc 2% MeOH) gave nucleoside analogue **12d** (13.8 mg, 39.4 μmol, 57% over 2 steps) as a white solid.

**$R_f$** = 0.56 (EtOAc, 7.5% MeOH); $[\alpha]_D^{25}$ = -42.9 (c = 0.10, MeOH); **$^1$H-NMR** (400 MHz, $CDCl_3$) δ [ppm] = 7.97 (s, 1H), 6.27 (s, 1H), 5.04 (m, 1H), 4.94-5.02 (m, 1H), 4.33 (dd, *J* = 2.4, 5.5 Hz, 1H), 4.03 (dd, *J* = 5.4, 9.2 Hz, 1H), 2.78 (brs, 1H), 2.53 (sept, *J* = 7.0 Hz, 1H), 2.41-2.50 (m, 1H), 2.37 (ddd, *J* = 4.5, 5.0, 11.5 Hz, 1H), 1.89-2.00 (m, 1H), 1.40 (q, *J* = 11.5 Hz, 1H), 1.14 (d, *J* = 7.0 Hz, 3H), 1.13 (d, *J* = 7.0 Hz, 3H); **$^{13}$C{$^1$H}-NMR** (101 MHz, $CDCl_3$) δ [ppm] = 178.4, 161.0, 151.4, 148.4, 139.3, 121.2, 77.8, 77.4, 71.2, 68.0, 44.5, 35.3, 33.6, 31.9, 19.3; **HRMS** (ESI, 3.5kV) calc for $[M+Na]^+$ $C_{16}H_{22}N_4O_5Na^+$ 373.14824, found 373.14765.

#### *Synthesis of Uridine Analogues*

##### Synthesis of 3-benzoyl-1-((3a*S*,4*S*,6a*R*)-5-iodo-2,2-dimethyl-3a,6a-dihydro-4*H*-cyclopenta[*d*][1,3]dioxol-4-yl)pyrimidine-2,4(1*H*,3*H*)-dione (14)

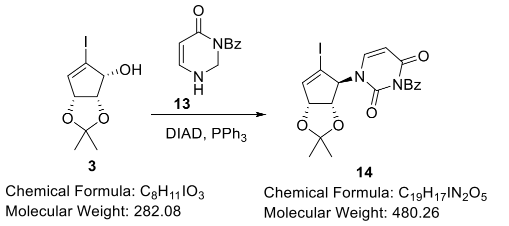

A solution of alcohol **3** (705 mg, 2.50 mmol, 1.00 eq.) and protected uracil **13** (649 mg, 3.00 mmol, 1.2 eq.) in THF (50 mL) was prepared and activated 4Å MS MS was added and the mixture was stirred for 2h. Then at 0 °C first $PPh_3$ (0.98 g, 3.75 mmol, 1.50 eq.) then DIAD (0.74 mL, 3.75 mmol, 1.50 eq.) were added. The reaction was stirred for 16h. Then the reaction was filtered, and the solvent was removed *in vacuo*. Purification by column chromatography ($SiO_2$, Hex:EtOAc 4:1 to 1:1) gave **14** (0.96 g, 2.00 mmol, 80%) as a white foam.

$\mathbf{R_f}$ = 0.45 (Hex/EA 1:1); **$^1$H-NMR** (400 MHz, $CDCl_3$) δ [ppm] = 7.93 (d, *J* = 7.5 Hz, 2H), 7.66 (tt, *J* = 1.2, 7.5 Hz, 1H), 7.50 (t, *J* = 7.9 Hz, 2H), 7.20 (brs, 1H), 6.53 (brs, 1H), 5.87 (d, *J* = 8.0 Hz, 1H), 5.22 (d, *J* = 5.7 Hz, 1H), 4.97 (brs, 1H), 4.55 (brs, 1H), 1.45 (s, 3H), 1.33 (s, 3H); **$^{13}$C{$^1$H}-NMR** (101 MHz, $CDCl_3$) δ [ppm] = 168.3, 162.0, 149.1, 145.8, 135.5, 131.1, 130.7, 129.4, 112.7, 103.1, 85.6, 80.6, 27.1, 25.7; **HRMS** (ESI, 3.5kV) calc for $[M+Na]^+$ $C_{19}H_{17}IN_2O_5Na^+$ 503.00743, found 503.00744.
*Note: Broad signals were observed in the proton spectra as well as in the carbon spectra. Therefore, not all carbon signals could be found despite intensive 2D NMR experiments.*

##### Synthesis of 3-benzoyl-1-((3a*S*,4*S*,6a*R*)-2,2-dimethyl-5-vinyl-3a,6a-dihydro-4*H*-cyclopenta[*d*][1,3]dioxol-4-yl)pyrimidine-2,4(1*H*,3*H*)-dione (S4)

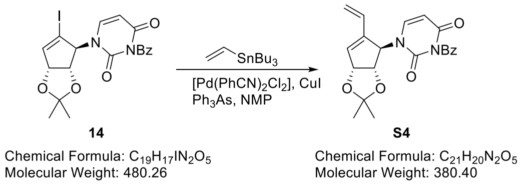

Vinyliodide **14** (1.70 g, 3.54 mmol, 1.00 eq.), $[Pd(PhCN)_2Cl_2]$ (67.9 mg, 0.18 mmol, 5 mol%), CuI (67.4 mg, 0.35 mmol, 10 mol%) and $Ph_3As$ (108.4 mg, 0.35 mmol, 10 mol%) were dissolved in freshly distilled NMP (over $CaH_2$, 11 mL). Then at 0 °C tributyl(vinyl)stannane (1.35 g, 1.24 mL, 4.25 mmol, 1.20 eq.) was added and the reaction was stirred for 4h at this temperature. Then the reaction was quenched with half sat. $NaHCO_3$ solution (20 mL) and the aqueous phase was extracted with EtOAc (3x 30 mL). The combined organic phase was washed with brine (30 mL) and brine was reextracted with EtOAc (1x 15 mL). The combined organic phases were dried over $MgSO_4$ and concentrated to give the crude. The crude product was purified by column chromatography on silica gel (first column: Hex/EA 2:1 to 1:2, second column to remove residual NMP and impurities: DCM/EA 3:1). The diene **S4** was isolated as a white crystalline solid (1.28 g, 3.36 mmol, 95%).

**$R_f$** = 0.53 (Hex/EA 1:2); **$^1$H-NMR** (400 MHz, $CDCl_3$) δ [ppm] = 7.91 (d, *J* = 7.3 Hz, 2H), 7.66 (tt, *J* = 1.2, 7.5 Hz, 1H), 7.50 (t, *J* = 8.0 Hz, 2H), 6.96 (brs, 1H), 6.47 (dd, *J* = 11.0, 17.8 Hz, 1H), 6.19 (brs, 1H), 5.79 (d, *J* = 8.1 Hz, 1H), 5.69 (brs, 1H), 5.39 (d, *J* = 11.0 Hz, 1H), 5.34 (d, *J* = 4.6 Hz, 1H), 5.25 (d, *J* = 18.0 Hz, 1H), 4.63 (brs, 1H), 1.41 (s, 3H), 1.34 (s, 3H); **$^{13}$C{$^1$H}-NMR** (101 MHz, $CDCl_3$) δ [ppm] = 168.5, 162.0, 149.8, 139.8, 136.7, 135.3, 131.5, 130.6, 130.0, 129.3, 120.3, 112.4, 102.8, 84.4, 83.3, 65.8, 27.4, 25.8; **HRMS** (ESI, 3.5kV) calc for $[M+Na]^+$ $C_{21}H_{20}N_2O_5Na^+$ 403.12644, found 403.12626.

*Note: Broad signals were observed in the proton spectra as well as in the carbon spectra. Therefore, not all carbon signals could be found despite long measurement time.*

#### Synthesis of 1-((3a*S*,4*R*,6a*R*)-2,2-dimethyl-5-vinyl-3a,6a-dihydro-4H-cyclopenta[*d*][1,3]dioxol-4-yl)pyrimidine-2,4(1H,3H)-dione (15)

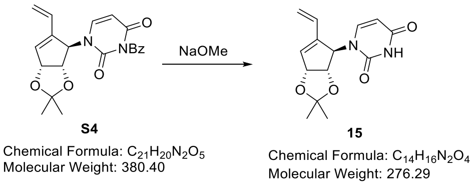

To a solution of ester **S4** (1.23 g, 3.24 mmol, 1.00 equiv.) in dry MeOH (32 mL) was added NaOMe (0.52 g, 9.71 mmol, 3.00 equiv.) at 0°C. The reaction was stirred at room temperature for 21h, then sat. $NH_4Cl$ solution was added (50 mL). The mixture was extracted with EtOAc (4x 50 mL), and the combined organic phases dried over $MgSO_4$ and concentrated *in vacuo*. Purification by column chromatography ($SiO_2$, 2:1 to 1:2 DCM/EtOAc) delivered deprotected **15** as a white solid (0.88 g, 3.18 mmol, 98%).

**$R_f$** = 0.19 (DCM/EA 3:1); **$^1$H-NMR** (400 MHz, $CDCl_3$) δ [ppm] = 9.17 (s, 1H), 6.84 (brs, 1H), 6.43 (dd, *J* = 11.0, 17.6 Hz, 1H), 6.17 (s, 1H), 5.74 (brs, 1H), 5.68 (dd, *J* = 2.1, 8.0 Hz, 1H), 5.31 (m, 2H), 5.20 (d, *J* = 17.8 Hz, 1H), 4.56 (d, *J* = 3.9 Hz, 1H), 1.43 (s, 3H), 1.34 (s, 3H); **$^{13}$C{$^1$H}-NMR** (101 MHz, $CDCl_3$) δ [ppm] = 163.3, 150.8, 139.9 (2x), 136.4, 129.8, 120.6, 112.3, 102.8, 84.5, 83.2, 65.1, 27.4, 25.8; **HRMS** (ESI, 3.5kV) calc for $[M+Na]^+$ $C_{14}H_{16}N_2O_4Na^+$ 299.10023, found 299.09993.

| # | C [ppm] | H [ppm] | 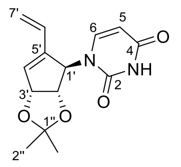 |
|---|---|---|---|
| 2 | 150.8 | -- | |
| 4 | 163.3 | -- | |
| 5 | 102.8 | 5.68 | |
| 6 | 139.9 | 6.84 | |
| 1’ | 65.1 | 5.74 | |
| 2’ | 83.2 | 5.31 | |
| 3’ | 84.5 | 4.56 | |
| 4’ | 136.4 | 6.17 | |
| 5’ | 139.9 | -- | |
| 6’ | 129.8 | 6.43 | |
| 7’ | 120.6 | 5.31, 5.20 | |
| 1’’ | 112.3 | -- | |
| 2’’ | 27.4/25.8 | 1.43, 1.34 | |
| NH | -- | 9.17 | |

#### Synthesis of 1-((3a*R*,3b*R*,4*R*,8*R*,8a*S*)-4-hydroxy-2,2-dimethyl-3a,3b,5,6,8,8a-hexahydro-4*H*-indeno[1,2-*d*][1,3]dioxol-8-yl)pyrimidine-2,4(1*H*,3*H*)-dione (16), 1-((3a*R*,3b*S*,5*R*,8a*S*)-5-hydroxy-2,2-dimethyl-3a,3b,5,6,8,8a-hexahydro-4*H*-indeno[1,2-*d*][1,3]dioxol-8-yl)pyrimidine-2,4(1*H*,3*H*)-dione (17) and 1-((3a*R*,3b*S*,5*S*,8a*S*)-5-hydroxy-2,2-dimethyl-3a,3b,5,6,8,8a-hexahydro-4*H*-indeno[1,2-*d*][1,3]dioxol-8-yl)pyrimidine-2,4(1*H*,3*H*)-dione (18)

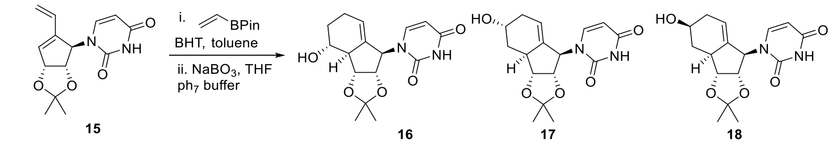

Chemical Formula: $C_{14}H_{16}N_2O_4$
Molecular Weight: 276.29

Chemical Formula: $C_{16}H_{20}N_2O_5$
Molecular Weight: 320.35

To a solution of diene **15** (227 mg, 0.82 mmol, 1.00 equiv.) in toluene (8 mL) was added BHT and vinyl pinacol borate (0.70 mL, 4.11 mmol, 5.00 equiv.). The reaction was stirred in a sealed tube for 4 d at 140°C (oil bath temperature). Then the volatiles were removed and the crude was dissolved in THF/ph 7 buffer (16 mL, 1:1). Then $NaBO_3$ $4H_2O$ (505 mg, 3.29 mmol, 4.00 equiv.) was added and the mixture was stirred for 1.5 h. The reaction was then quenched by sat. $Na_2S_2O_3$ solution (10 mL). After phase separation, the aqueous phase was extracted with EtOAc (3x15 mL). The combined organic phases were dried over $MgSO_4$ and concentrated *in vacuo*. Purification by column chromatography ($SiO_2$, DCM:EtOAc 1:1 to DCM:acetone 4:1 to 1:1) gave **16** (64.3 mg, 0.20 mmol, 24%), **17** (21.0 mg, 65.6 μmol, 8%) and **18** (79.0 mg, 0.25 mmol, 30%) as white powders.

**16:** $\mathbf{R_f}$ = 0.23 (DCM/acetone 3:1); **$^{1}$H-NMR** (600 MHz, 315 K, $CDCl_3$) δ [ppm] = 8.59 (s, 1H), 7.08 (brs, 1H), 5.73 (d, *J* = 6.0 Hz, 1H), 5.58 (brs, 0.3 H), 5.27 (s, 1H), 4.21-5.10 (brs, 2.7H), 3.66 (brs, 1H), 2.55 (brs, 1H), 2.13-2.31 (m, 2H), 1.91-2.03 (m, 2H), 1.57 (s, 3H), 1.34 (s, 3H), 1.25-1.29 (m, 1H); **$^{13}$C{$^{1}$H}-NMR** (150 MHz, 315 K, $CDCl_3$) δ [ppm] = 121.2, 114.8, 102.7, 83.1, 71.8, 53.5, 31.1, 27.4, 25.2, 24.8; **HRMS** (ESI, 3.5kV) calc for $[M+Na]^+$ $C_{16}H_{20}N_2O_5Na^+$ 343.12644, found 343.12623.

**17:** $\mathbf{R_f}$ = 0.19 (DCM/acetone 3:1); **$^{1}$H-NMR** (600 MHz, 315 K, $CDCl_3$) δ [ppm] = 8.97 (s, 1H), 7.10 (brs, 1H), 5.72 (d, *J* = 7.0 Hz, 1H), 5.23 (s, 1H), 4.30-5.10 (brs, 3H), 4.25 (s, 1H), 2.78 (brs, 1H), 2.29-2.38 (m, 2H), 2.09-2.19 (m, 1H), 1.55 (s, 3H), 1.43 (m, 1H), 1.33 (s, 3H); **$^{13}$C{$^{1}$H}-NMR** (150 MHz, 315 K, $CDCl_3$) δ [ppm] = 140.6, 139.0, 117.9, 113.4, 102.3, 83.8 (2x), 64.4, 39.7, 33.5, 33.0, 27.8, 25.6; **HRMS** (ESI, 3.5kV) calc for $[M+Na]^+$ $C_{16}H_{20}N_2O_5Na^+$ 343.12644, found 343.12638.

| **#** | **C [ppm]** | **H [ppm]** |
|---|---|---|
| 2 | *not found* | -- |
| 4 | *not found* | -- |
| 5 | 102.3 | 5.72 |
| 6 | 140.6 | 7.10 |
| 1’ | *not found* | |
| 2’ | 83.8 | 4.30-5.10 |
| 3’ | 83.8 | 4.30-5.10 |
| 4’ | 39.7 | 2.78 |
| 5’ | 33.5 | 2.29-2.38/1.43 |
| 6’ | 64.4 | 4.25 |
| 7’ | 33.0 | 2.29-2.38/2.09-2.19 |
| 8’ | 117.9 | 5.23 |
| 9’ | 139.0 | -- |
| 1” | 113.4 | -- |
| 2” | 27.8/25.6 | 1.55/1.33 |
| N3H | | 8.97 |

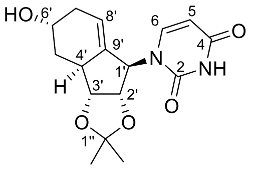

**17**

**18:** $\mathbf{R_f}$ = 0.16 (DCM/acetone 3:1); **$^{1}$H-NMR** (600 MHz, 315 K, $CDCl_3$) δ [ppm] = 9.18 (brs, 1H), 7.08 (brs, 1H), 5.74 (d, *J* = 7.4 Hz, 1H), 5.25 (s, 1H), 4.21-5.11 (brs, 3H), 3.97 (brs, 1H), 2.67 (brs, 1H), 2.35-2.50 (m, 2H), 1.93-2.01 (m, 2H), 1.54 (s, 3H), 1.40-1.47 (m, 1H), 1.31 (s, 3H); **$^{13}$C{$^{1}$H}-NMR** (150 MHz, 315 K, $CDCl_3$) δ [ppm] = 163.2, 150.7, 142.8, 139.2, 119.6, 113.8, 102.2, 83.3 (2x), 67.1, 64.5, 45.1, 36.5, 33.9, 27.5, 25.5; **HRMS** (ESI, 3.5kV) calc for $[M+Na]^+$ $C_{16}H_{20}N_2O_5Na^+$ 343.12644, found 343.12604.

| # | C [ppm] | H [ppm] |
|---|---|---|
| 2 | 150.7 | -- |
| N3H | -- | 9.18 |
| 4 | 163.2 | -- |
| 5 | 102.2 | 5.74 |
| 6 | 142.8 | 7.08 |
| 1’ | 64.5 | 4.21-5.11 |
| 2’ | 83.3 | 4.21-5.11 |
| 3’ | 83.3 | 4.21-5.11 |
| 4’ | 45.1 | 2.67 |
| 5’ | 36.5 | 2.39/1.42 |
| 6’ | 67.1 | 3.97 |
| 7’ | 33.9 | 2.43/1.97 |
| 8’ | 119.6 | 5.25 |
| 9’ | 139.2 | -- |
| 1” | 113.8 | -- |
| 2” | 27.5/25.5 | 1.54/1.31 |
| 6’-OH | | 1.99 |

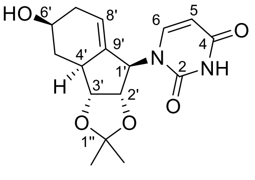

**18**

#### Synthesis of 1-((1*R*,2*S*,3*R*,3a*R*,4*R*)-2,3,4-trihydroxy-2,3,3a,4,5,6-hexahydro-1*H*-inden-1-yl)pyrimidine-2,4(1*H*,3*H*)-dione (19a)

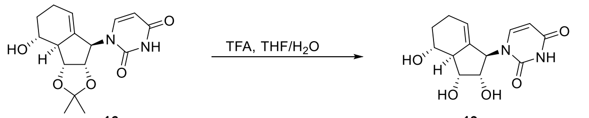

**16**
Chemical Formula: $C_{16}H_{20}N_2O_5$
Molecular Weight: 320.35

**19a**
Chemical Formula: $C_{13}H_{16}N_2O_5$
Molecular Weight: 280.28

Alcohol **16** (20.0 mg, 62.4 μmol, 1.00 equiv.) was dissolved in THF (0.5 ml) and TFA (1.0 mL, 10% in $H_2O$) was added. After 21 h, the solvent was removed *in vacuo*. Purification by column chromatography ($SiO_2$, EtOAc 5% MeOH) gave nucleoside analogue **19a** (17.0 mg, 60.7 μmol, 97%) as a white solid.

**$R_f$** = 0.23 (EtOAc 5% MeOH); $[\alpha]_D^{25}$ = -25.1 (c = 0.09, MeOH); **$^1$H-NMR** (400 MHz, MeOD) δ [ppm] = 7.47 (d, *J* = 8.1 Hz, 1H), 5.70 (d, *J* = 8.0 Hz, 1H), 5.30 (brs, 1H), 5.22 (m, 1H), 4.07 (brs, 1H), 3.97 (brs, 1H), 3.56 (ddd, *J* = 3.8, 9.5, 11.3 Hz, 1H), 2.41 (m, 1H), 2.23 (m, 2H), 1.96 (m, 1H), 1.58 (m, 1H); **$^{13}$C{$^1$H}-NMR** (100 MHz, MeOD) δ [ppm] = 166.2, 153.4, 135.4, 121.2, 102.6, 75.5, 73.8, 71.9, 53.5, 32.4, 26.0; **HRMS** (ESI, 3.5kV) calc for $[M+Na]^+$ $C_{13}H_{16}N_2O_5Na^+$ 303.09514, found 303.09520.

#### Synthesis of (1*R*,2*S*,3*R*,3a*S*,4*R*)-1-(2,4-dioxo-3,4-dihydropyrimidin-1(2*H*)-yl)-2,3-dihydroxy-2,3,3a,4,5,6-hexahydro-1*H*-inden-4-yl isobutyrate (19b)

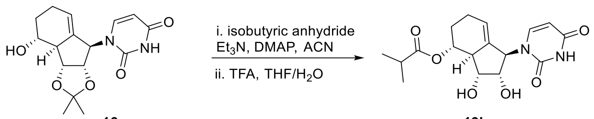

**16**
Chemical Formula: $C_{16}H_{20}N_2O_5$
Molecular Weight: 320.35

**19b**
Chemical Formula: $C_{17}H_{22}N_2O_6$
Molecular Weight: 350.37

Alcohol **16** (50 mg, 0.16 mmol, 1.00 equiv.) was dissolved in acetonitrile (3 mL) and isobutyric anhydride (52 µL, 0.31 mmol, 2.00 equiv.), $Et_3N$ (44 µL, 0.31 mmol, 2.00 equiv.) and DMAP (1.9 mg, 15.6 µmol, 10 mol%) were added. The reaction was stirred for 4 h, then sat. $NaHCO_3$ (10 mL) and EtOAc (10 mL) were added. After phase separation, the aqueous phase was extracted with EtOAc (3x15 mL). The combined organic phases were dried over $MgSO_4$ and concentrated *in vacuo*. Purification by column chromatography ($SiO_2$, DCM:EtOAc 1:1) gave the ester (58.0 mg, 0.15 mmol, 97%) as a white solid. For deprotection, an aliquote (15.2 mg, 38.9 µmol) was dissolved in THF (0.5 ml) and TFA (1.0 mL, 10% in $H_2O$) was added. After 16 h, the solvent was removed *in vacuo*. Purification by column chromatography ($SiO_2$, EtOAc 5% MeOH) gave nucleoside analogue **19b** (12.6 mg, 35.9 µmol, 92%, 90% over two steps) as a white solid.

**R$_f$** = 0.53 (EtOAc 5% MeOH); $[\alpha]_D^{25}$ = -44.0 (c = 0.11, MeOH); **$^1$H-NMR** (400 MHz, MeOD) δ [ppm] = 7.50 (d, *J* = 8.1 Hz, 1H), 5.71 (d, *J* = 8.0 Hz, 1H), 5.32 (m, 1H), 5.27 (brs, 1H), 4.86 (m, 1H), 3.99 (brs, 2H), 2.66 (m, 1H), 2.59 (sep, *J* = 7.0 Hz, 1H), 2.28 (m, 2H), 1.94-2.02 (m, 1H), 1.61-1.73 (m, 1H), 1.20 (d, *J* = 6.9 Hz, 3H), 1.18 (d, *J* = 7.0 Hz, 3H); **$^{13}$C{$^1$H}-NMR** (100 MHz, MeOD) δ [ppm] = 178.8, 165.0, 151.8, 144.9, 135.2, 121.5, 102.4, 75.7, 74.0, 73.8, 50.1, 35.4, 29.0, 25.6, 19.4 (2x); **HRMS** (ESI, 3.5kV) calc for $[M+Na]^+$ $C_{17}H_{22}N_2O_6Na^+$ 373.13701, found 373.13701.

| # | C [ppm] | H [ppm] |
|---|---|---|
| 2 | 151.8 | -- |
| 4 | 165.0 | -- |
| 5 | 102.4 | 5.71 |
| 6 | 144.9 | 7.50 |
| 1’ | *not found* | 5.27 |
| 2’ | 73.8 *or* 75.7 | 3.99 |
| 3’ | 73.8 *or* 75.7 | 3.99 |
| 4’ | 50.1 | 2.66 |
| 5’ | 74.0 | 4.86 |
| 6’ | 29.0 | 1.98/1.67 |
| 7’ | 25.6 | 2.28 |
| 8’ | 121.5 | 5.32 |
| 9’ | 135.2 | -- |
| 1’’ | 178.8 | -- |
| 2’’ | 35.4 | 2.59 |
| 2’’ Me | 19.4 | 1.20/1.18 |

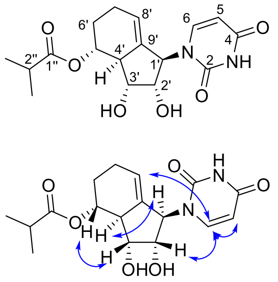

NOE correlations

#### Synthesis of 1-((1*R*,2*S*,3*R*,3a*S*,5*R*)-2,3,5-trihydroxy-2,3,3a,4,5,6-hexahydro-1*H*-inden-1-yl)pyrimidine-2,4(1*H*,3*H*)-dione (20a)

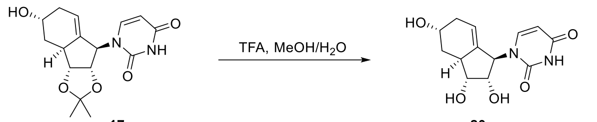

**17** — Chemical Formula: $C_{16}H_{20}N_2O_5$, Molecular Weight: 320.35

**20a** — Chemical Formula: $C_{13}H_{16}N_2O_5$, Molecular Weight: 280.28

Alcohol **17** (20.0 mg, 62.4 μmol, 1.00 equiv.) was dissolved in MeOH (0.5 ml) and TFA (0.5 mL, 10% in $H_2O$) was added. After 16 h, the solvent was removed *in vacuo*. Purification by column chromatography ($SiO_2$, EtOAc 5% MeOH) gave nucleoside analogue **20a** (16.0 mg, 60.7 μmol, 91%) as a white solid.

**$R_f$** = 0.15 (EtOAc 5% MeOH); $[\alpha]_D^{25}$ = -7.5 (c = 0.10, MeOH); **$^1$H-NMR** (600 MHz, MeOD, 315 K) δ [ppm] = 7.43 (d, *J* = 7.9 Hz, 1H), 5.69 (d, *J* = 7.9 Hz, 1H), 5.25 (m, 1H), 5.19 (brs, 1H), 4.17 (s, 1H), 4.05 (s, 1H), 3.84 (s, 1H), 2.71 (s, 1H), 2.30-2.37 (m, 1H), 2.25 (dt, *J* = 4.8, 12.9 Hz, 1H), 2.12 (d, *J* = 18.8 Hz, 1H), 1.40 (ddd, *J* = 2.0, 11.2, 15.1 Hz, 1H); **$^{13}$C{$^1$H}-NMR** (150 MHz, MeOD, 315K) δ [ppm] = 166.3, 152.9, 145.3, 137.6, 119.4, 102.6, 76.9, 76.4, 65.1, 59.3, 39.4, 34.2, 34.0; **HRMS** (ESI, 3.5kV) calc for $[M+Na]^+$ $C_{13}H_{16}N_2O_5Na^+$ 303.09514, found 303.09498.

| # | C [ppm] | H [ppm] |
|---|---|---|
| 2 | 152.9 | -- |
| 4 | 166.3 | -- |
| 5 | 102.6 | 5.69 |
| 6 | 145.3 | 7.43 |
| 1’ | 59.3 | 5.19 |
| 2’ | 76.4 | 4.05 |
| 3’ | 76.9 | 3.84 |
| 4’ | 39.4 | 2.71 |
| 5’ | 34.0 | 2.25 (down) 1.40 (up) |
| 6’ | 65.1 | 4.17 |
| 7’ | 34.2 | 2.34 (up) 2.12 (down) |
| 8’ | 119.4 | 5.25 |
| 9’ | 137.7 | -- |

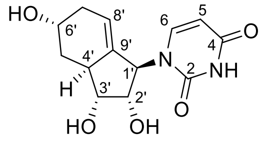

NOE correlations

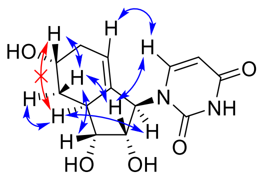

#### Synthesis of (1*R*,2*S*,3*R*,3a*S*,5*R)*-1-(2,4-dioxo-3,4-dihydropyrimidin-1(2*H*)-yl)-2,3-dihydroxy-2,3,3a,4,5,6-hexahydro-1*H*-inden-5-yl isobutyrate (20b)

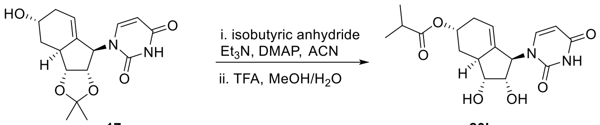

**17** Chemical Formula: $C_{16}H_{20}N_2O_5$ Molecular Weight: 320.35

**20b** Chemical Formula: $C_{17}H_{22}N_2O_6$ Molecular Weight: 350.37

Alcohol **17** (40.0 mg, 0.13 mmol, 1.00 equiv.) was dissolved in acetonitrile (2 mL) and isobutyric anhydride (41 µL, 0.25 mmol, 2.00 equiv.), $Et_3N$ (35 µL, 0.25 mmol, 2.00 equiv.) and DMAP (1.5 mg, 12.5 µmol, 10 mol%) were added. The reaction was stirred for 22 h, then sat. $NaHCO_3$ (10 mL) and EtOAc (10 mL) were added. After phase separation, the aqueous phase was extracted with EtOAc (3x15 mL). The combined organic phases were dried over $MgSO_4$ and concentrated *in vacuo*. Purification by column chromatography ($SiO_2$, DCM:EtOAc 1:1) gave the ester (27.9 mg, 71.5 µmol, 57%) as a white solid. For deprotection, an aliquote (20.0 mg, 51.2 µmol) was dissolved in MeOH (0.75 ml) and TFA (0.25 mL, 10% in $H_2O$) was added. After 44 h, the solvent was removed *in vacuo*. Purification by column chromatography ($SiO_2$, EtOAc 3% MeOH) gave nucleoside analogue **20b** (17.1 mg, 48.8 µmol, 96%, 55% over two steps) as a white solid.

**R$_f$** = 0.23 (EtOAc 3% MeOH); $[\alpha]_D^{25}$ = +7.7 (c = 0.11, MeOH); **$^1$H-NMR** (400 MHz, MeOD, 295K) δ [ppm] = 7.45 (d, *J* = 7.9 Hz, 1H), 5.69 (d, *J* = 7.9 Hz, 1H), 5.29 (m, 1H), 5.22 (s, 1H), 5.18 (brs, 1H), 4.08 (brs, 1H), 3.86 (brs, 1H), 2.66 (brs, 1H), 2.52 (sep, *J* = 7.0 Hz, 1H), 2.35-2.45 (m, 2H), 2.18 (d, *J* = 19.7 Hz, 1H), 1.47 (ddd, *J* = 2.0, 11.5, 15.2 Hz, 1H), 1.15 (d, *J* = 7.0 Hz, 3H), 1.14 (d, *J* = 7.0 Hz, 3H); **$^{13}$C{$^1$H}-NMR** (150 MHz, $CDCl_3$, 315 K) δ [ppm] = 178.3, 166.5, 153.1, 146.0, 137.9, 118.7, 102.7, 76.9, 76.4, 68.9, 62.9, 39.8, 35.4, 31.4 (2x), 19.3, 19.2; **HRMS** (ESI, 3.5kV) calc for $[M+Na]^+$ $C_{17}H_{22}N_2O_6Na^+$ 373.13701, found 373.13674.

| # | C [ppm] | H [ppm] |
|---|---|---|
| 2 | 153.1 | -- |
| 4 | 166.5 | -- |
| 5 | 102.7 | 5.69 |
| 6 | 146.0 | 7.45 |
| 1’ | 62.9 | 5.18 |
| 2’ | 76.4 | 4.08 |
| 3’ | 76.9 | 3.86 |
| 4’ | 39.8 | 2.66 |
| 5’ | 31.4 | 1.47 (up)/2.36 (down) |
| 6’ | 68.9 | 5.22 |
| 7’ | 31.4 | 2.18 (down)/2.42 (up) |
| 8’ | 118.7 | 5.29 |
| 9’ | 137.9 | -- |
| 1’’ | 178.3 | -- |
| 2’’ | 35.4 | 2.52 |
| 2’’ Me | 19.3/19.2 | 1.15/1.14 |

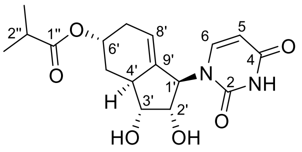

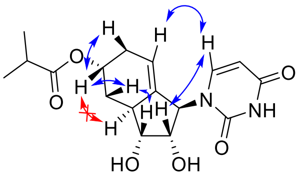

*key NOE Interactions*

#### Synthesis of 1-((1*R*,2*S*,3*R*,3a*S*,5*S*)-2,3,5-trihydroxy-2,3,3a,4,5,6-hexahydro-1*H*-inden-1-yl)pyrimidine-2,4(1*H*,3*H*)-dione (21a)

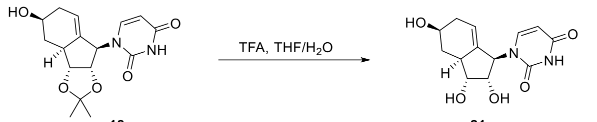

**18** Chemical Formula: $C_{16}H_{20}N_2O_5$ Molecular Weight: 320.35

**21a** Chemical Formula: $C_{13}H_{16}N_2O_5$ Molecular Weight: 280.28

Alcohol **18** (24.2 mg, 75.5 μmol, 1.00 equiv.) was dissolved in THF (0.5 ml) and TFA (1 mL, 10% in $H_2O$) was added. After 44 h, the solvent was removed *in vacuo*. Purification by column chromatography ($SiO_2$, EtOAc 7.5% MeOH) gave nucleoside analogue **21a** (19.9 mg, 71.0 μmol, 94%) as a white solid.

$\mathbf{R_f}$ = 0.15 (EtOAc 5% MeOH); $[\alpha]_D^{25}$ = -4.6 (c = 0.13, MeOH); **$^1$H-NMR** (600 MHz, MeOD, 315 K) δ [ppm] = 7.43 (d, *J* = 7.4 Hz, 1H), 5.69 (d, *J* = 7.8 Hz, 1H), 5.28 (s, 1H), 5.15 (brs, 1H), 4.05 (brs, 1H), 3.90 (m, 1H), 3.82 (brs, 1H), 2.61 (brs, 1H), 2.42 (d, *J* = 17.4 Hz, 1H), 2.32 (m, 1H), 1.93-1.97 (m, 1H), 1.32 (q, *J* = 11.6 Hz, 1H); **$^{13}$C{$^1$H}-NMR** (150 MHz, MeOD, 315 K) δ [ppm] = 166.4, 153.0, 145.5, 137.8, 120.6, 102.6, 76.8, 76.6, 68.2, 62.4, 45.3, 37.2, 35.4; **HRMS** (ESI, 3.5kV) calc for $[M+Na]^+$ $C_{13}H_{16}N_2O_5Na^+$ 303.09514, found 303.09489.

| # | C [ppm] | H [ppm] |
|---|---|---|
| 2 | 153.0 | -- |
| 4 | 166.4 | -- |
| 5 | 102.6 | 5.69 |
| 6 | 145.5 | 7.43 |
| 1’ | 62.4 | 5.15 |
| 2’ | 76.6 | 3.82 |
| 3’ | 76.8 | 4.05 |
| 4’ | 45.3 | 2.61 |
| 5’ | 37.2 | 2.32 (down) 1.32 (up) |
| 6’ | 68.2 | 3.90 |
| 7’ | 35.4 | 2.42 (down) 1.95 (up) |
| 8’ | 120.6 | 5.28 |
| 9’ | 137.8 | -- |

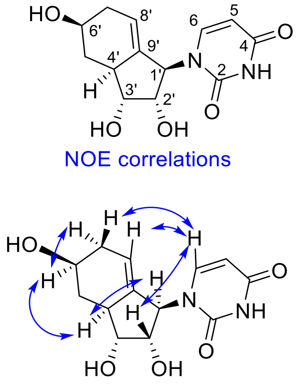

#### Synthesis of (1*R*,2*S*,3*R*,3a*S*,5*S*)-1-(2,4-dioxo-3,4-dihydropyrimidin-1(2*H*)-yl)-2,3-dihydroxy-2,3,3a,4,5,6-hexahydro-1*H*-inden-5-yl isobutyrate (21b)

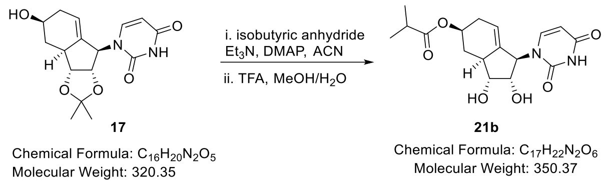

Alcohol **17** (40.0 mg, 0.13 mmol, 1.00 equiv.) was dissolved in acetonitrile (2 mL) and isobutyric anhydride (41 µL, 0.25 mmol, 2.00 equiv.), $Et_3N$ (35 µL, 0.25 mmol, 2.00 equiv.) and DMAP (1.5 mg, 12.5 µmol, 10 mol%) were added. The reaction was stirred for 24 h, then sat. $NaHCO_3$ (10 mL) and EtOAc (10 mL) were added. After phase separation, the aqueous phase was extracted with EtOAc (3x15 mL). The combined organic phases were dried over $MgSO_4$ and concentrated *in vacuo*. Purification by column chromatography ($SiO_2$, DCM:EtOAc 1:1) gave the ester (45.0 mg, 0.12 mmol, 92%) as a white solid. For deprotection, an aliquote (20.5 mg, 52.5 µmol) was dissolved in MeOH (0.75 ml) and TFA (0.25 mL, 10% in $H_2O$) was added. After 40 h, the solvent was removed *in vacuo*. Purification by column chromatography ($SiO_2$, EtOAc 2% MeOH) gave nucleoside analogue **21b** (18.4 mg, 52.5 µmol, quant., 92% over two steps) as a white solid.

**$R_f$** = 0.33 (EtOAc 5% MeOH); $[\alpha]_D^{25}$ = -11.0 (c = 0.13, MeOH); **$^1$H-NMR** (600 MHz, MeOD, 315 K) δ [ppm] = 7.43 (d, *J* = 7.40 Hz, 1H), 5.70 (d, *J* = 7.7 Hz, 1H), 5.31 (s, 1H), 5.12 (brs, 1H), 4.99 (m, 1H), 4.08 (brs, 1H), 3.87 (brs, 1H), 2.68 (brs, 1H), 2.54 (sep, *J* = 6.9 Hz, 1H), 2.50 (m, 1H), 2.33-2.38 (m, 1H), 2.05-2.12 (m, 1H), 1.46 (q, *J* = 11.6 Hz, 1H), 1.15 (d, *J* = 6.9 Hz, 3H), 1.14 (d, *J* = 6.9 Hz, 3H); **$^{13}$C{$^1$H}-NMR** (100 MHz, $CDCl_3$) δ [ppm] = 178.4, 166.3, 152.7, 145.3, 138.0, 119.8, 102.7, 76.8, 76.6, 71.2, 44.7, 35.3, 33.5, 31.8, 19.2 (2x); **HRMS** (ESI, 3.5kV) calc for $[M+Na]^+$ $C_{17}H_{22}N_2O_6Na^+$ 373.13701, found 373.13688.

| # | C [ppm] | H [ppm] |
|---|---|---|
| 2 | 152.7 | -- |
| 4 | 166.3 | -- |
| 5 | 102.7 | 5.70 |
| 6 | 145.3 | 7.43 |
| 1’ | *not found* | 5.12 |
| 2’ | 76.8 | 4.08 |
| 3’ | 76.6 | 3.87 |
| 4’ | 44.7 | 2.68 |
| 5’ | 33.5 | 2.37/1.46 |
| 6’ | 71.2 | 4.99 |
| 7’ | 31.8 | 2.50/2.08 |
| 8’ | 119.8 | 5.31 |
| 9’ | 138.0 | -- |
| 1’’ | 178.4 | -- |
| 2’’ | 35.3 | 2.54 |

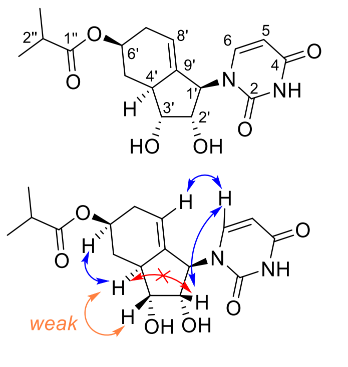

| | | |
|---|---|---|
| 2'' Me | 19.2 (2x) | 1.14/1.15 |

| # | **10a** | **10b** | **19a** |
|---|---|---|---|
| 1' | 5.25 brs | 6.28 s | 5.30 brs |
| 2' | 4.16 dd | 4.19 dd | 3.97 brs |
| 3' | 4.25 dd | 4.33 dd | 4.07 brs |
| 4' | **2.48 brs** | **2.51 brs** | **2.41 m** |
| 5' | **3.57 ddd** | **3.56 ddd** | **3.56 ddd** |
| 6' | **1.96 m**<br>**1.57 ddt** | **1.95 m**<br>**1.57 ddt** | **1.96 m**<br>**1.58 m** |
| 7' | **2.19 m** | **2.20 m**<br>**2.08 m** | **2.23 m** |
| 8' | 5.03 m | 4.91 m | 5.22 m |

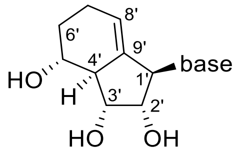

| # | **10c** | **10d** | **19b** |
|---|---|---|---|
| 1' | 5.25 s | 6.29 s | 5.27 brs |
| 2' | 4.08 dd | 4.14 dd | 3.99 brs |
| 3' | 4.22 dd | 4.27 dd | 3.99 brs |
| 4' | **2.74 brs** | **2.75 m** | **2.66 m** |
| 5' | **4.91 m** | **4.87 m** | **4.86 m** |
| 6' | **1.97 m**<br>**1.68 ddt** | **1.95 m**<br>**1.67 ddt** | **1.98 m**<br>**1.67 m** |
| 7' | **2.25 m** | **2.25/2.12** | **2.28 m** |
| 8' | 5.17 m | 5.04 m | 5.32 m |

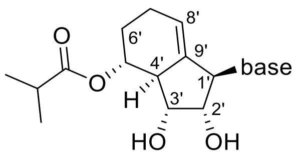

### X-Ray Data for 12b

Single-crystal X-ray data are collected on Bruker SMART CCD 6000 Diffractometer using a Cu Kα radiation (1.54178 Å) at 297 K. The crystal was kept at 297.00 K during data collection. Using Olex2,[7] the structure was solved with the SHELXS[8] structure solution program using Direct Methods and refined with the SHELXL[9] refinement package using Least Squares minimisation. The X-ray crystallographic data for the compound **12b** have been deposited at the Cambridge Crystallographic Data Centre (CCDC), under the deposition number CCDC 2211316. The data can be obtained free of charge from the Cambridge Crystallographic Data Center via https://www.ccdc.cam.ac.uk/structures/

| | |
|---|---|
| **Molecular Formula, Formula Weight** | $C_{12}H_{16}N_4O_4$, 280.28 |
| **Crystal system, Space group** | monoclinic, $P2_1$ |
| **a/Å** | 4.7807(4) |
| **b/Å** | 27.786(2) |
| **c/Å** | 9.8696(7) |
| **α/°** | 90 |
| **β/°** | 103.773(2) |
| **γ/°** | 90 |
| **V/Å$^3$** | 1273.34(17) |
| **Z** | 2 |
| **Density (calcd)/g cm$^{-3}$** | 1.462 |
| **μ/mm$^{-1}$** | 0.942 |
| **F(000)** | 592.0 |
| **Crystal size/ mm$^3$** | 0.4 × 0.2 × 0.1 |
| **Radiation type, Wavelength/Å** | CuKα (λ = 1.54178) |
| **Temperature/K** | 297 |
| **$Θ_{min}$, $Θ_{max}$/ °** | 9.226, 137.07 |
| **Index ranges** | -5 ≤ h ≤ 5, -33 ≤ k ≤ 33, -11 ≤ l ≤ 11 |
| **Collected, Independent reflections** | 18121, 4529 |
| **$R_{int}$** | 0.0619 |
| **Parameters, Restraints** | 368/1 |
| **Largest Peak/e Å$^3$, Deepest Hole/e Å$^3$** | 0.70, -0.65 |

| | |
|---|---|
| **Goodness-of-fit on $F^2$** | 1.165 |
| **$R_1$ [$F^2$ > 2σ( $F^2$)], $wR_2$ [$F^2$]** | 0.1030, 0.2976 |

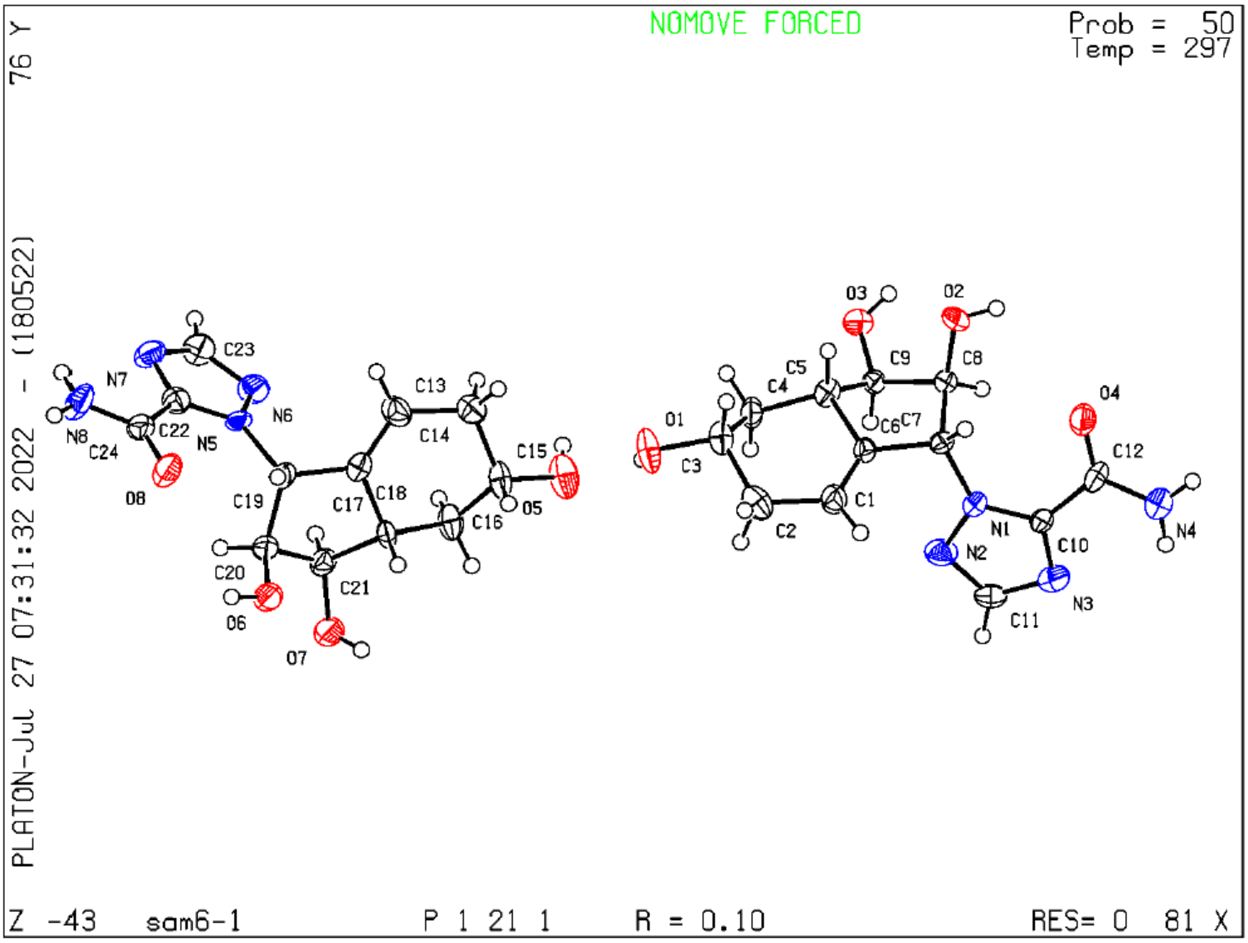

### NMR Spectra

-- starting from the next page –

#### NMR-Spectra for Compound S1

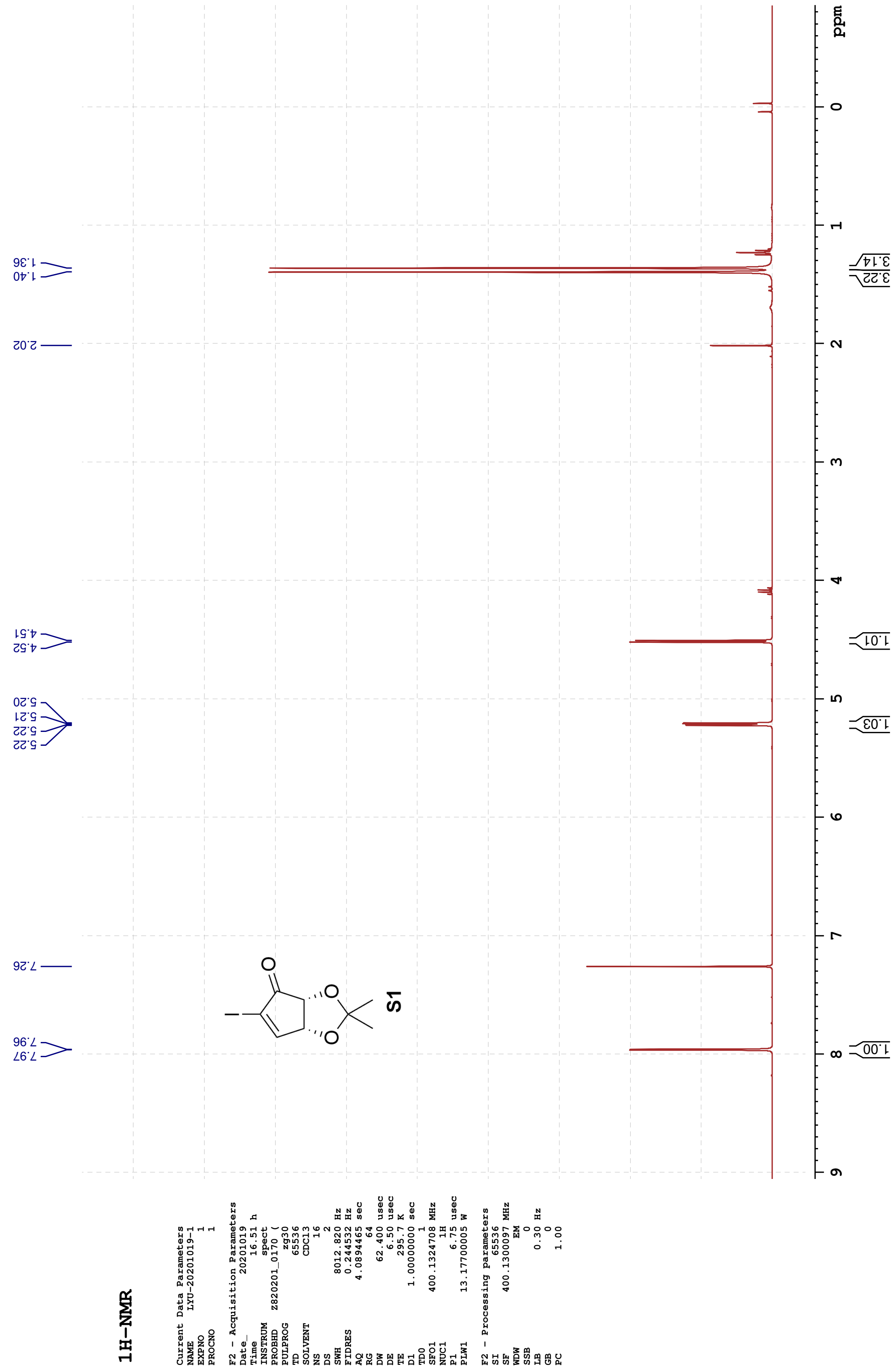

**13C-{1H}-NMR**

```
Current Data Parameters
NAME       LYU-20201019-1
EXPNO                   3
PROCNO                  1

F2 - Acquisition Parameters
Date_            20201019
Time                17.00 h
INSTRUM             spect
PROBHD     Z820201_0170 (
PULPROG           zgpg30
TD                 65536
SOLVENT            CDCl3
NS                    66
DS                     4
SWH            40760.871 Hz
FIDRES          1.243923 Hz
AQ             0.8039083 sec
RG                   203
DW                12.267 usec
DE                  6.50 usec
TE                 295.8 K
D1            2.00000000 sec
D11           0.03000000 sec
TD0                    1
SFO1         100.6228298 MHz
NUC1                 13C
P1                 28.00 usec
PLW1         14.80000019 W
SFO2         400.1316005 MHz
NUC2                  1H
CPDPRG[2         waltz16
PCPD2              90.00 usec
PLW2         13.17000008 W
PLW12         0.07408100 W
PLW13         0.03726200 W

F2 - Processing parameters
SI                 32768
SF           100.6127620 MHz
WDW                   EM
SSB                    0
LB                  1.00 Hz
GB                     0
PC                  1.40
```

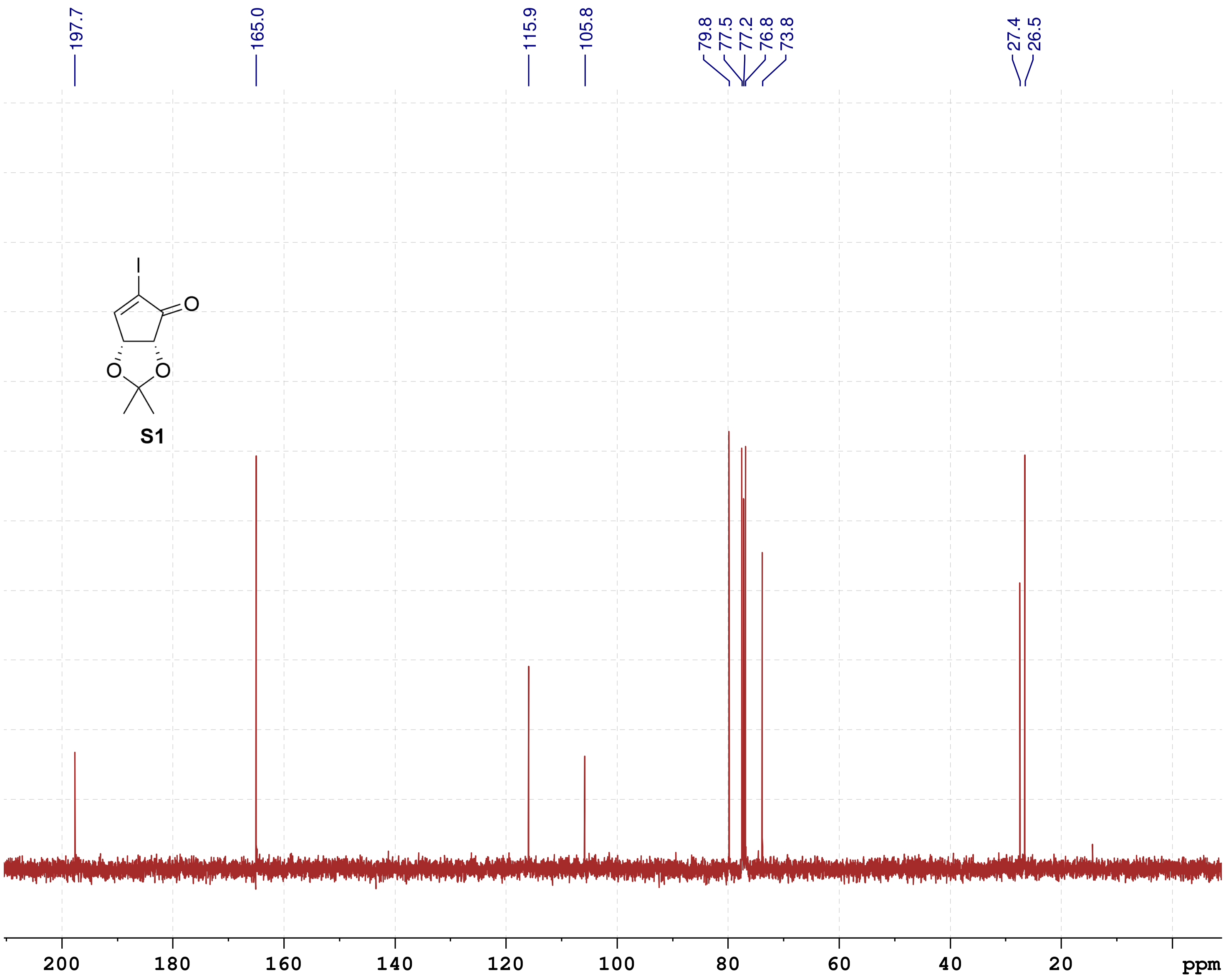

#### NMR-Spectra for Compound 3

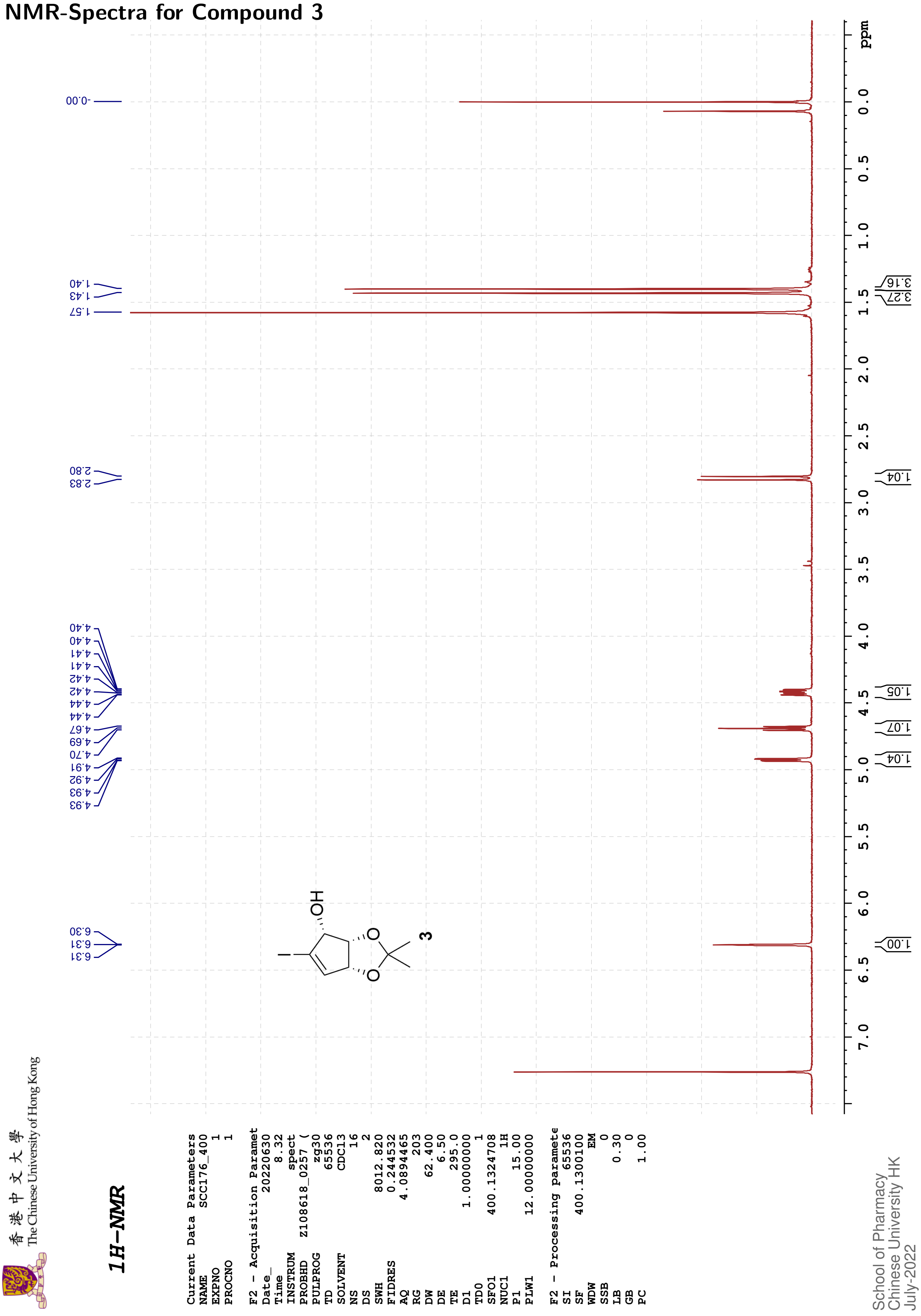

#### NMR-Spectra for Compound 5a

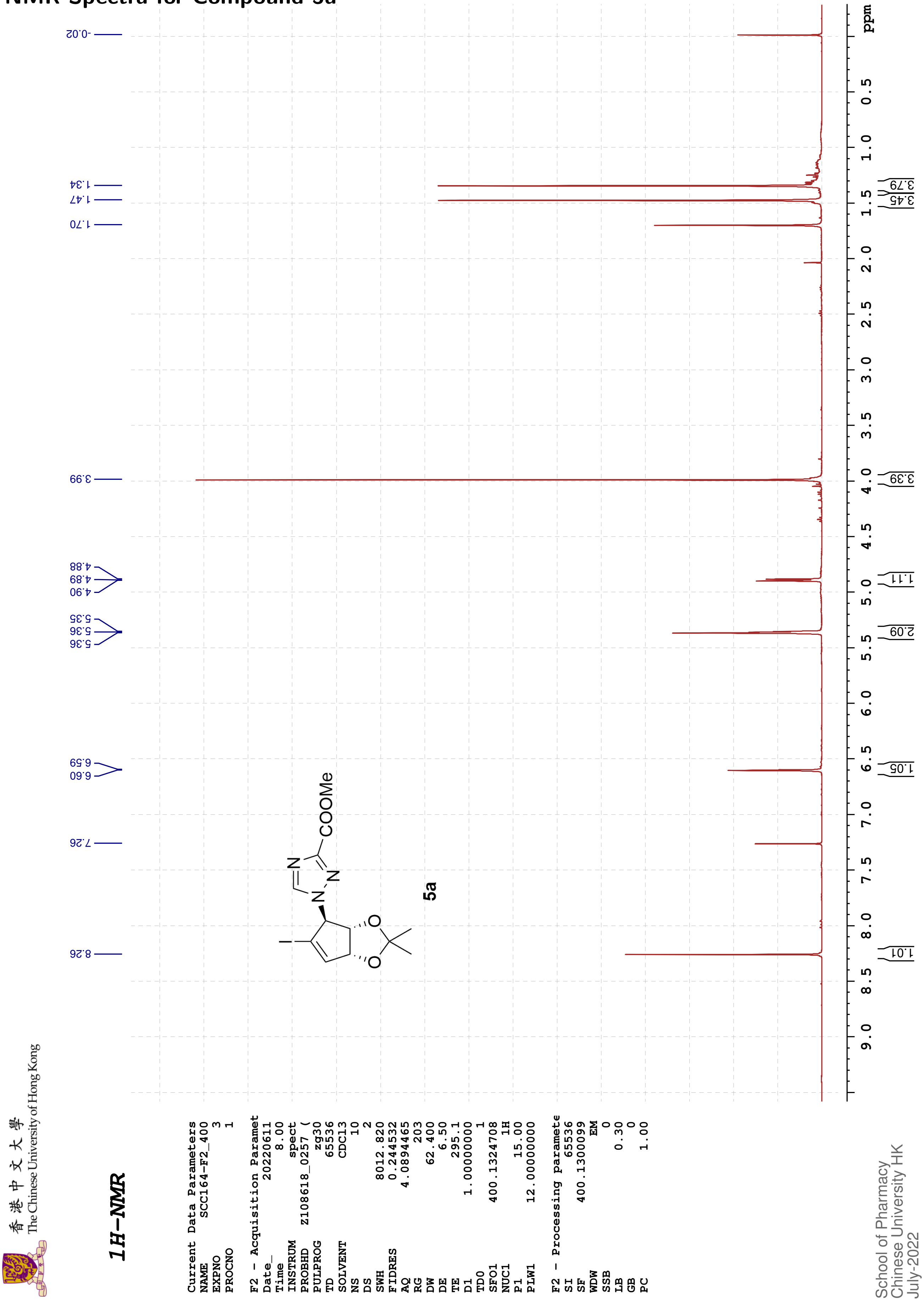

香港中文大學
The Chinese University of Hong Kong

#### 13C{1H}-NMR

| | |
|---|---|
| Current Data Parameters | |
| NAME | SCC164-F2_400 |
| EXPNO | 4 |
| PROCNO | 1 |
| F2 - Acquisition Paramet | |
| Date_ | 20220611 |
| Time | 8.12 |
| INSTRUM | spect |
| PROBHD | Z108618_0257 ( |
| PULPROG | zgpg30 |
| TD | 48074 |
| SOLVENT | CDCl3 |
| NS | 216 |
| DS | 4 |
| SWH | 24038.461 |
| FIDRES | 1.000061 |
| AQ | 0.9999392 |
| RG | 203 |
| DW | 20.800 |
| DE | 6.50 |
| TE | 295.3 |
| D1 | 2.00000000 |
| D11 | 0.03000000 |
| TD0 | 1 |
| SFO1 | 100.6228298 |
| NUC1 | 13C |
| P1 | 10.00 |
| PLW1 | 47.00000000 |
| SFO2 | 400.1316005 |
| NUC2 | 1H |
| CPDPRG[2 | waltz16 |
| PCPD2 | 90.00 |
| PLW2 | 12.00000000 |
| PLW12 | 0.33333001 |
| PLW13 | 0.16766000 |
| F2 - Processing paramete | |
| SI | 32768 |
| SF | 100.6127577 |
| WDW | EM |
| SSB | 0 |
| LB | 2.00 |
| GB | 0 |
| PC | 1.40 |

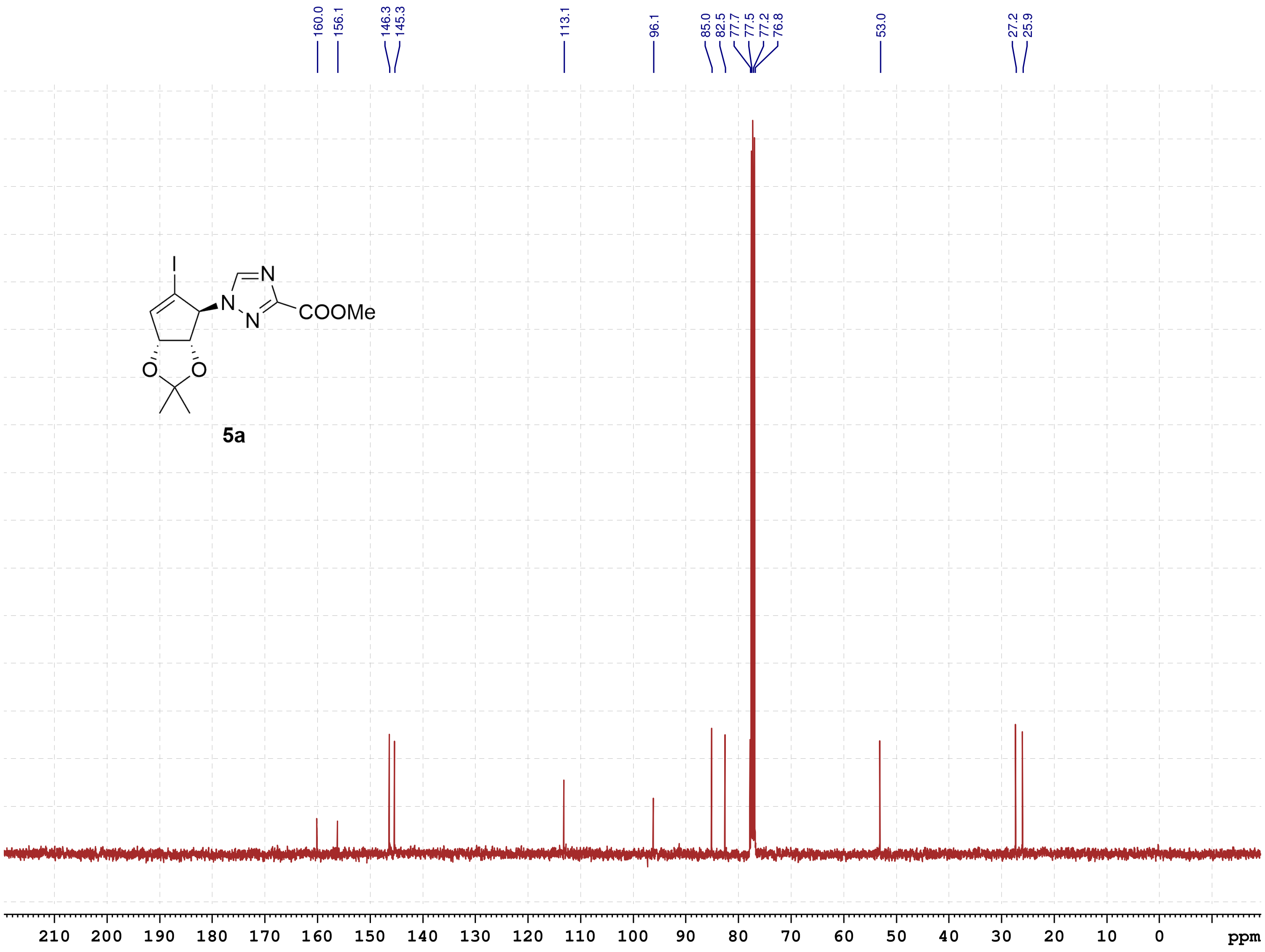

School of Pharmacy
Chinese University HK
July-2022

#### NMR-Spectra for Compound 6a

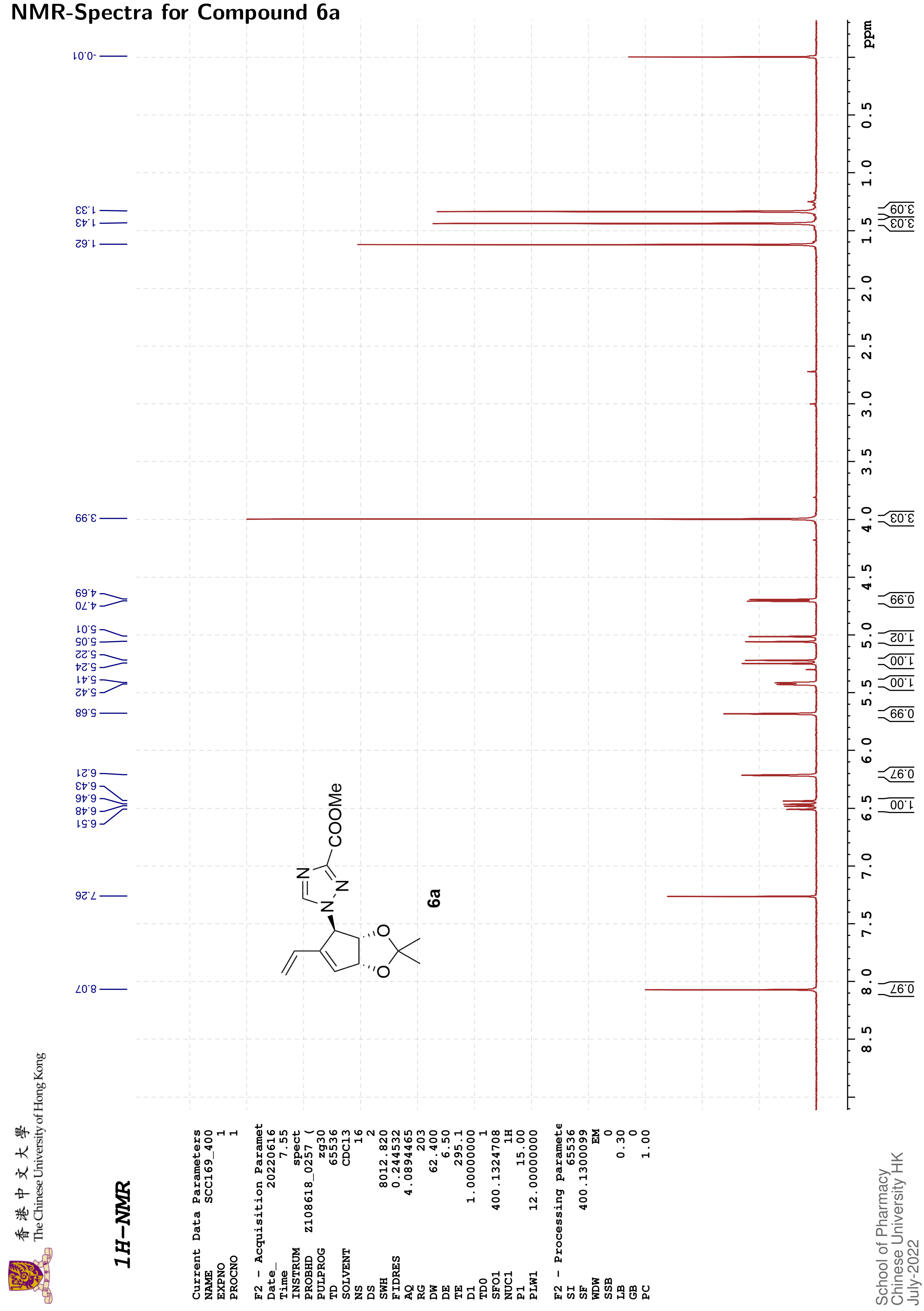

香港中文大學 The Chinese University of Hong Kong

13C{1H}-NMR

```
Current Data Parameters
NAME          SCC169_400
EXPNO                  2
PROCNO                 1

F2 - Acquisition Paramet
Date_           20220616
Time                8.13
INSTRUM            spect
PROBHD     Z108618_0257 (
PULPROG          zgpg30
TD                48074
SOLVENT           CDCl3
NS                  320
DS                    4
SWH           24038.461
FIDRES         1.000061
AQ            0.9999392
RG                  203
DW               20.800
DE                 6.50
TE                295.8
D1           2.00000000
D11          0.03000000
TD0                   1
SFO1        100.6228298
NUC1                13C
P1                10.00
PLW1        47.00000000
SFO2        400.1316005
NUC2                 1H
CPDPRG[2        waltz16
PCPD2             90.00
PLW2        12.00000000
PLW12        0.33333001
PLW13        0.16766000

F2 - Processing paramete
SI                32768
SF          100.6127559
WDW                  EM
SSB                   0
LB                 0.40
GB                    0
PC                 1.40
```

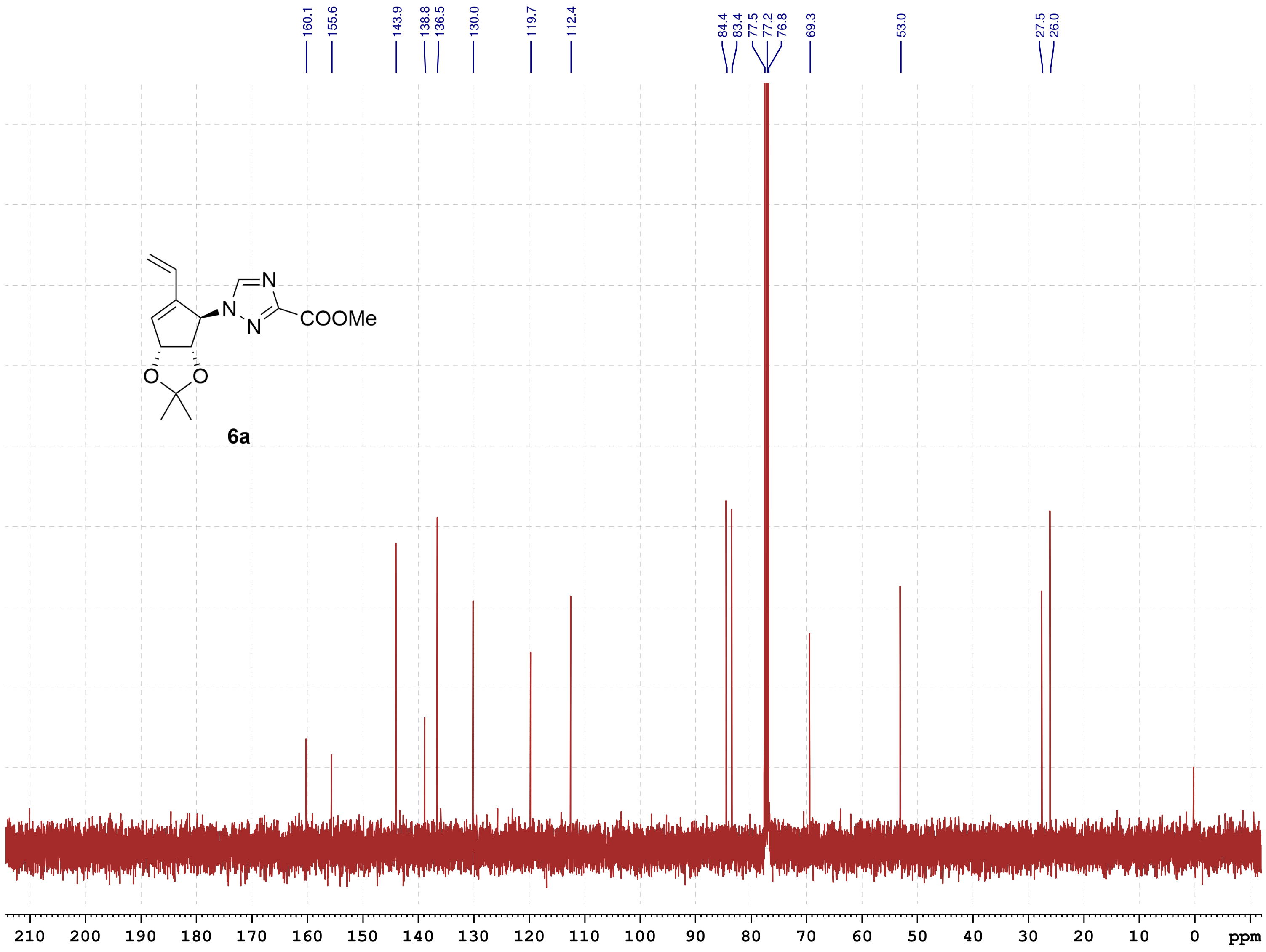

School of Pharmacy
Chinese University HK
September-2022

#### NMR-Spectra for Compound 7a

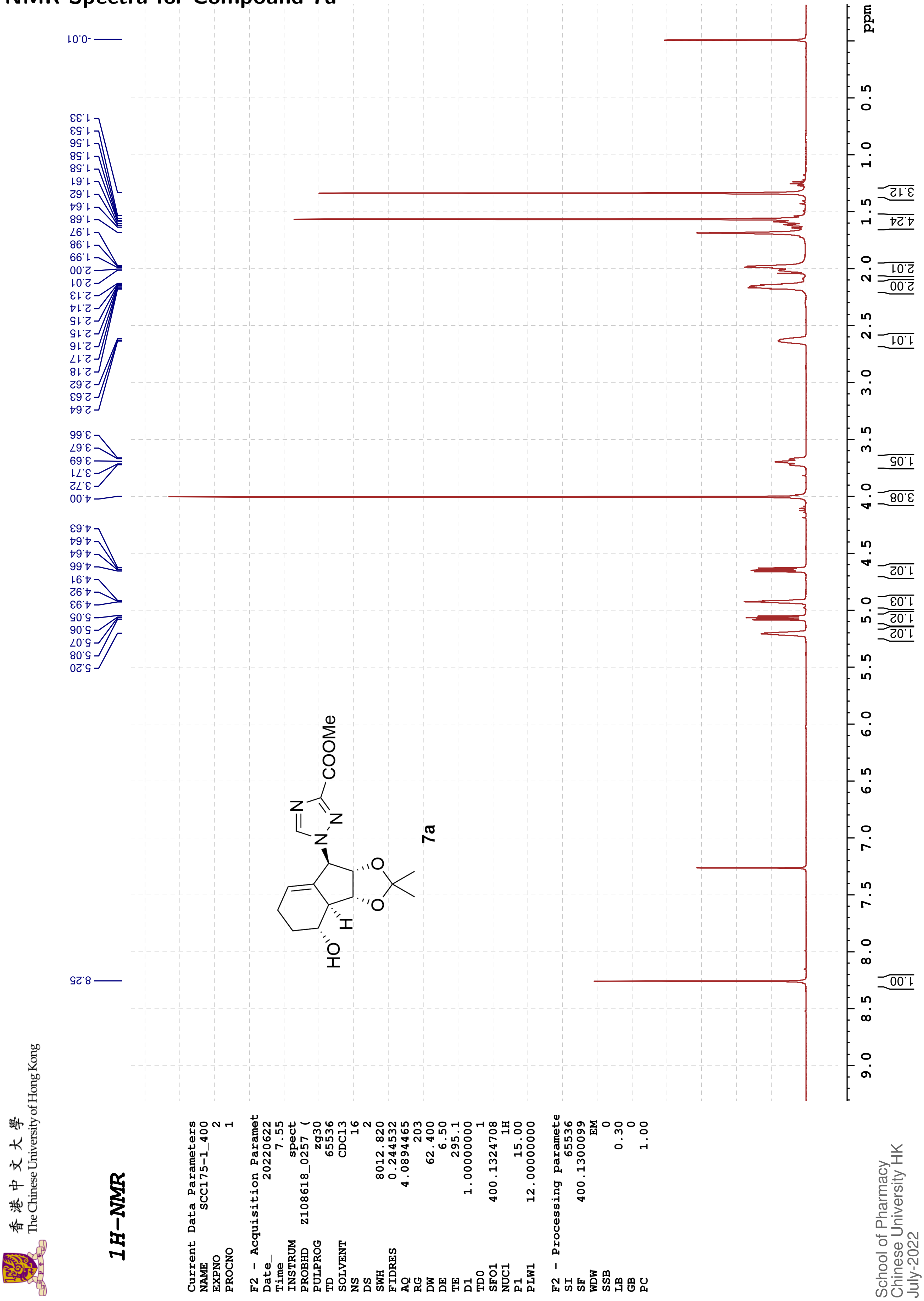

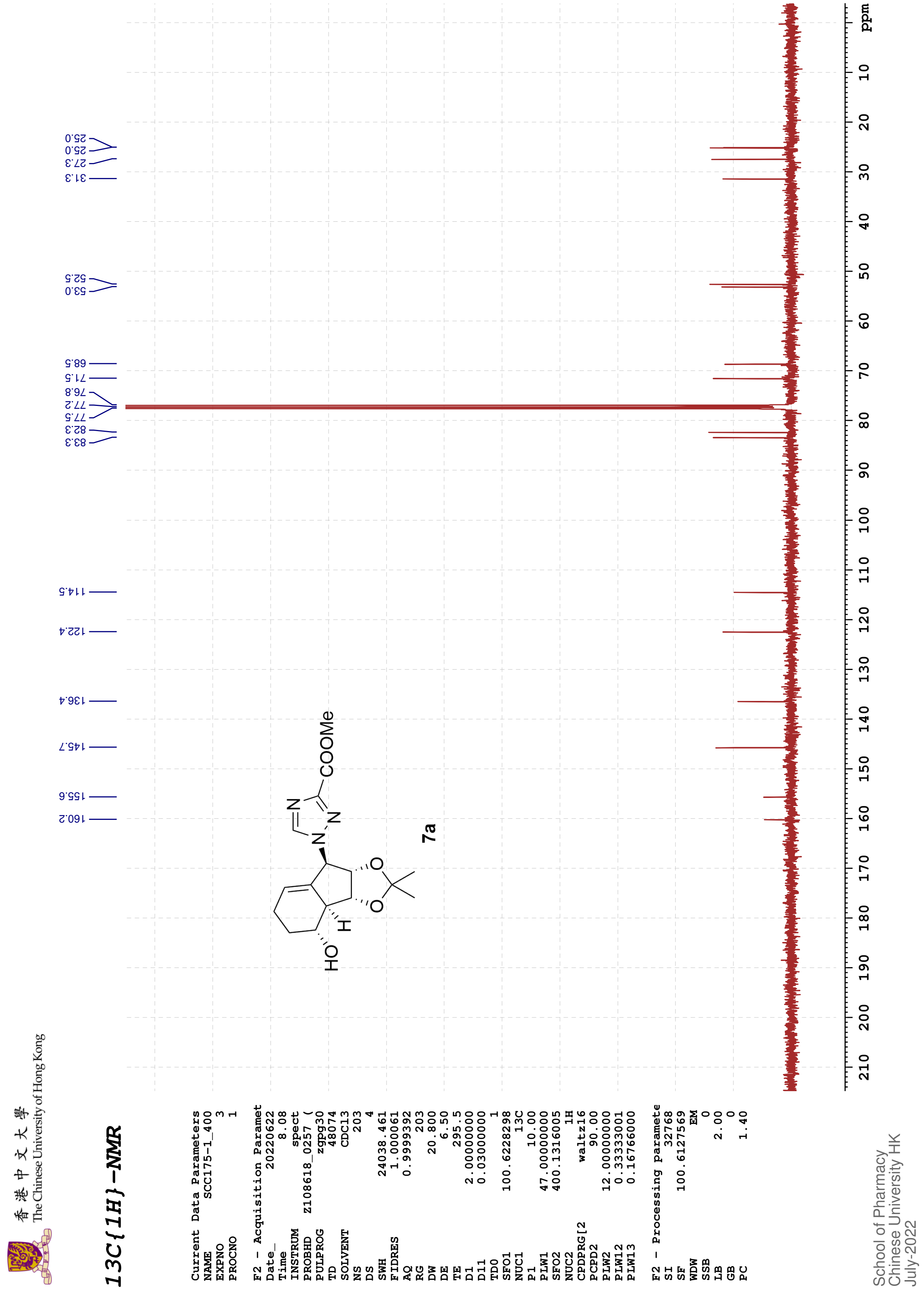
13C{1H}-NMR
The Chinese University of Hong Kong
Current Data Parameters
NAME SCC175-1_400
EXPNO 3
PROCNO 1
F2 - Acquisition Paramet
Date_ 20220622
Time 8.08
INSTRUM spect
PROBHD Z108618_0257 (
PULPROG zgpg30
TD 48074
SOLVENT CDCl3
NS 203
DS 4
SWH 24038.461
FIDRES 1.000061
AQ 0.9999392
RG 203
DW 20.800
DE 6.50
TE 295.5
D1 2.00000000
D11 0.03000000
TD0 1
SFO1 100.6228298
NUC1 13C
P1 10.00
PLW1 47.00000000
SFO2 400.1316005
NUC2 1H
CPDPRG[2 waltz16
PCPD2 90.00
PLW2 12.00000000
PLW12 0.33333001
PLW13 0.16766000
F2 - Processing paramete
SI 32768
SF 100.6127569
WDW EM
SSB 0
LB 2.00
GB 0
PC 1.40
School of Pharmacy
Chinese University HK
July-2022
160.2
155.6
145.7
136.4
122.4
114.5
83.3
82.3
77.5
77.2
76.8
71.5
68.5
53.0
52.5
31.3
27.3
25.0
25.0
COOMe
N
N
N
HO
H
O
O
7a
ppm

#### NMR-Spectra for Compound 10a

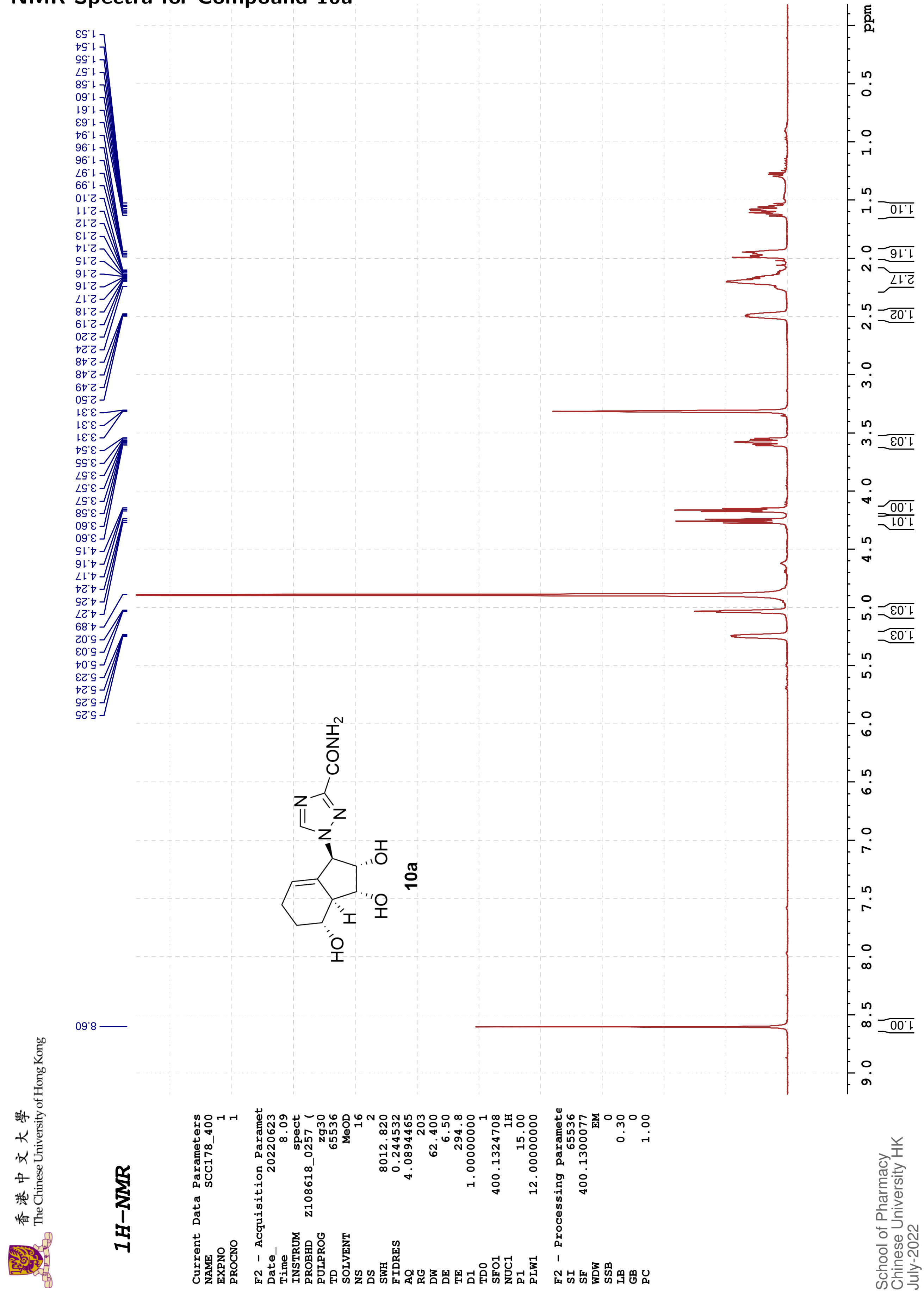

香港中文大學
The Chinese University of Hong Kong

#### 13C{1H}-NMR

```
Current Data Parameters
NAME            SCC178_400
EXPNO                    2
PROCNO                   1

F2 - Acquisition Paramet
Date_             20220623
Time                  8.25
INSTRUM              spect
PROBHD    Z108618_0257 (
PULPROG            zgpg30
TD                   48074
SOLVENT               MeOD
NS                     289
DS                       4
SWH              24038.461
FIDRES            1.000061
AQ               0.9999392
RG                     203
DW                  20.800
DE                    6.50
TE                   295.6
D1              2.00000000
D11             0.03000000
TD0                      1
SFO1           100.6228298
NUC1                   13C
P1                   10.00
PLW1           47.00000000
SFO2           400.1316005
NUC2                    1H
CPDPRG[2           waltz16
PCPD2                90.00
PLW2           12.00000000
PLW12           0.33333001
PLW13           0.16766000

F2 - Processing paramete
SI                   32768
SF             100.6126282
WDW                     EM
SSB                      0
LB                    2.00
GB                       0
PC                    1.40
```

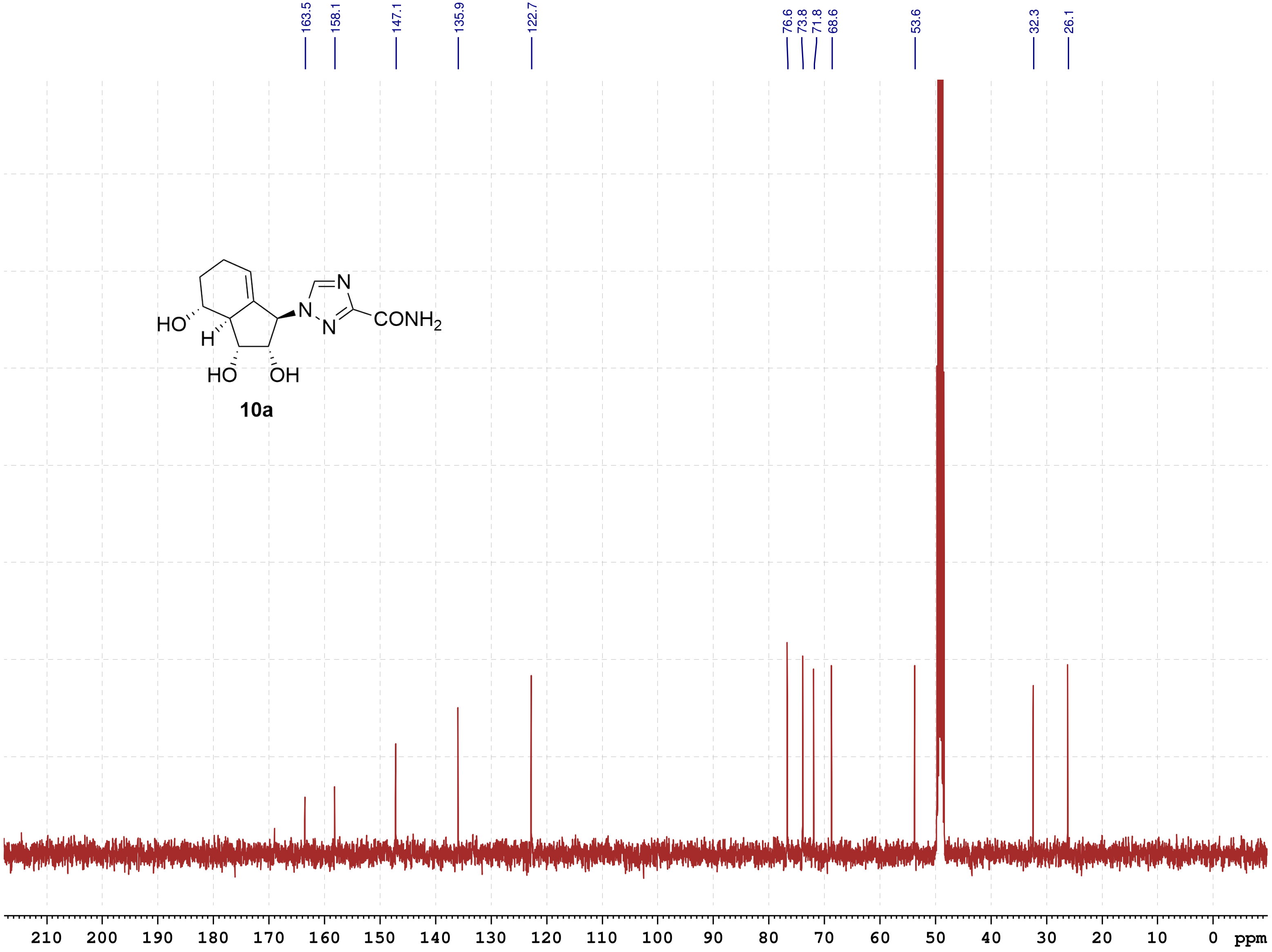

School of Pharmacy
Chinese University HK
July-2022

**COSY**

```
Current Data Parameters
NAME          SCC178_400
EXPNO                  4
PROCNO                 1

F2 - Acquisition Parameters
Date_           20220623
Time                8.41 h
INSTRUM            spect
PROBHD   Z108618_0257 (
PULPROG         cosygpqf
TD                  2048
SOLVENT             MeOD
NS                     2
DS                     8
SWH             4000.000 Hz
FIDRES          3.906250 Hz
AQ             0.2560000 sec
RG                   203
DW               125.000 usec
DE                  6.50 usec
TE                 294.9 K
D0            0.00000300 sec
D1            1.48689198 sec
D13           0.00000400 sec
D16           0.00020000 sec
IN0           0.00025000 sec
TDav                   1
SFO1         400.1320006 MHz
NUC1                  1H
P0                 15.00 usec
P1                 15.00 usec
PLW1         12.00000000 W
GPNAM[1]        SINE.100
GPZ1               10.00 %
P16              1000.00 usec

F1 - Acquisition parameters
TD                   128
SFO1             400.132 MHz
FIDRES         62.500000 Hz
SW                 9.997 ppm
FnMODE                QF

F2 - Processing parameters
SI                  1024
SF           400.1300081 MHz
WDW                 SINE
SSB                    0
LB                     0 Hz
GB                     0
PC                  1.40

F1 - Processing parameters
SI                  1024
MC2                   QF
SF           400.1300078 MHz
WDW                 SINE
SSB                    0
LB                     0 Hz
GB                     0
```

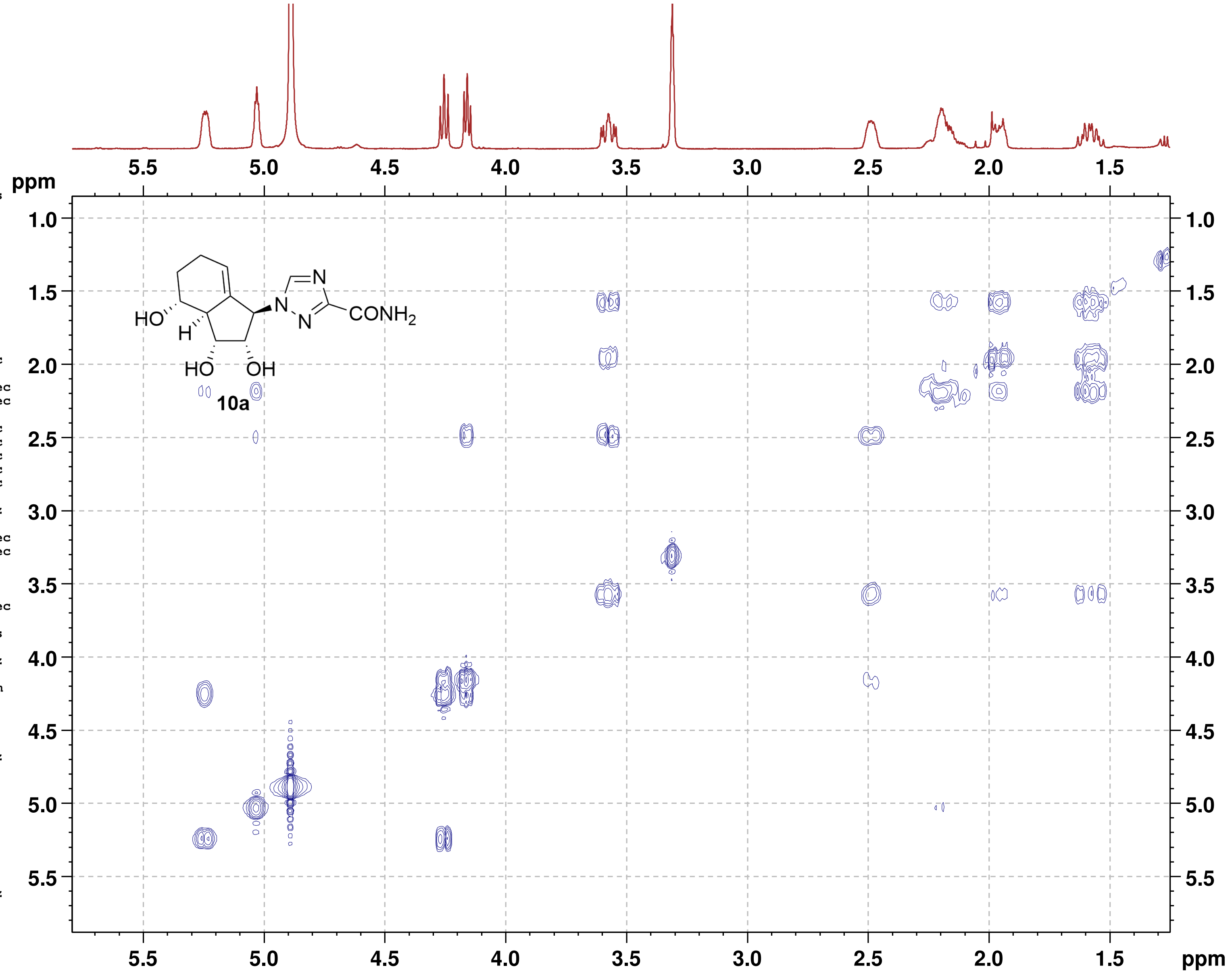

**NOESY**

```
Current Data Parameters
NAME          SCC178_400
EXPNO                  3
PROCNO                 1

F2 - Acquisition Parameters
Date_           20220623
Time                8.27 h
INSTRUM            spect
PROBHD     Z108618_0257 (
PULPROG      noesygpphpp
TD                  2048
SOLVENT             MeOD
NS                     2
DS                    16
SWH             4000.000 Hz
FIDRES          3.906250 Hz
AQ             0.2560000 sec
RG                   203
DW               125.000 usec
DE                  6.50 usec
TE                 295.1 K
D0            0.00010590 sec
D1            2.00000000 sec
D8            0.80000001 sec
D11           0.03000000 sec
D12           0.00002000 sec
D16           0.00020000 sec
IN0           0.00025000 sec
TDav                   1
SFO1         400.1318806 MHz
NUC1                  1H
P1                 15.00 usec
P2                 30.00 usec
P17              2500.00 usec
PLW1         12.00000000 W
PLW10         3.00000000 W
GPNAM[1]       SMSQ10.100
GPZ1               40.00 %
P16              1000.00 usec

F1 - Acquisition parameters
TD                   128
SFO1            400.1319 MHz
FIDRES         62.500000 Hz
SW                 9.997 ppm
FnMODE       States-TPPI

F2 - Processing parameters
SI                  1024
SF           400.1300080 MHz
WDW                QSINE
SSB                    2
LB                     0 Hz
GB                     0
PC                  1.00

F1 - Processing parameters
SI                  1024
MC2          States-TPPI
SF           400.1300072 MHz
WDW                QSINE
SSB                    2
LB                     0 Hz
GB                     0
```

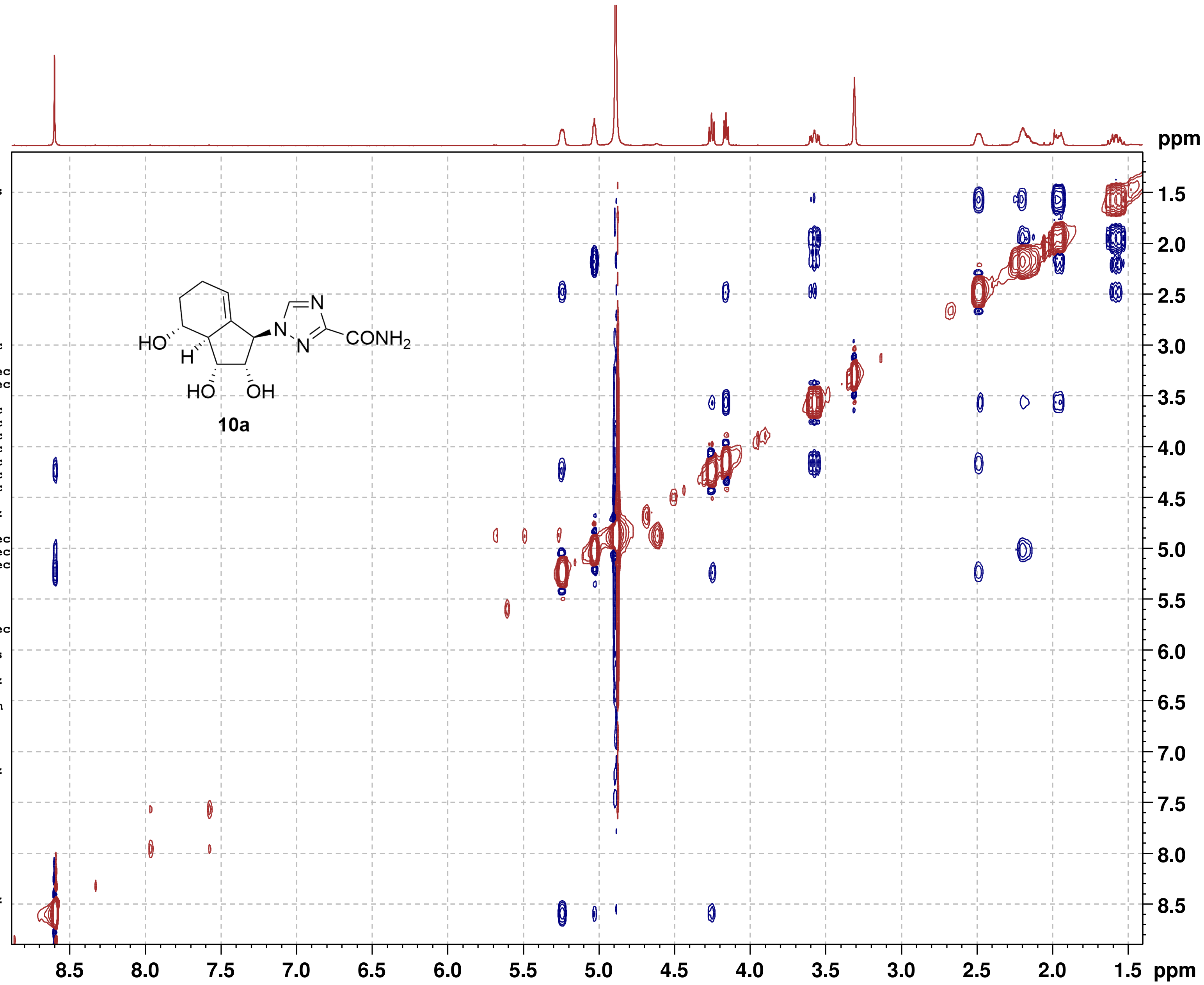

#### NMR-Spectra for Compound 10c

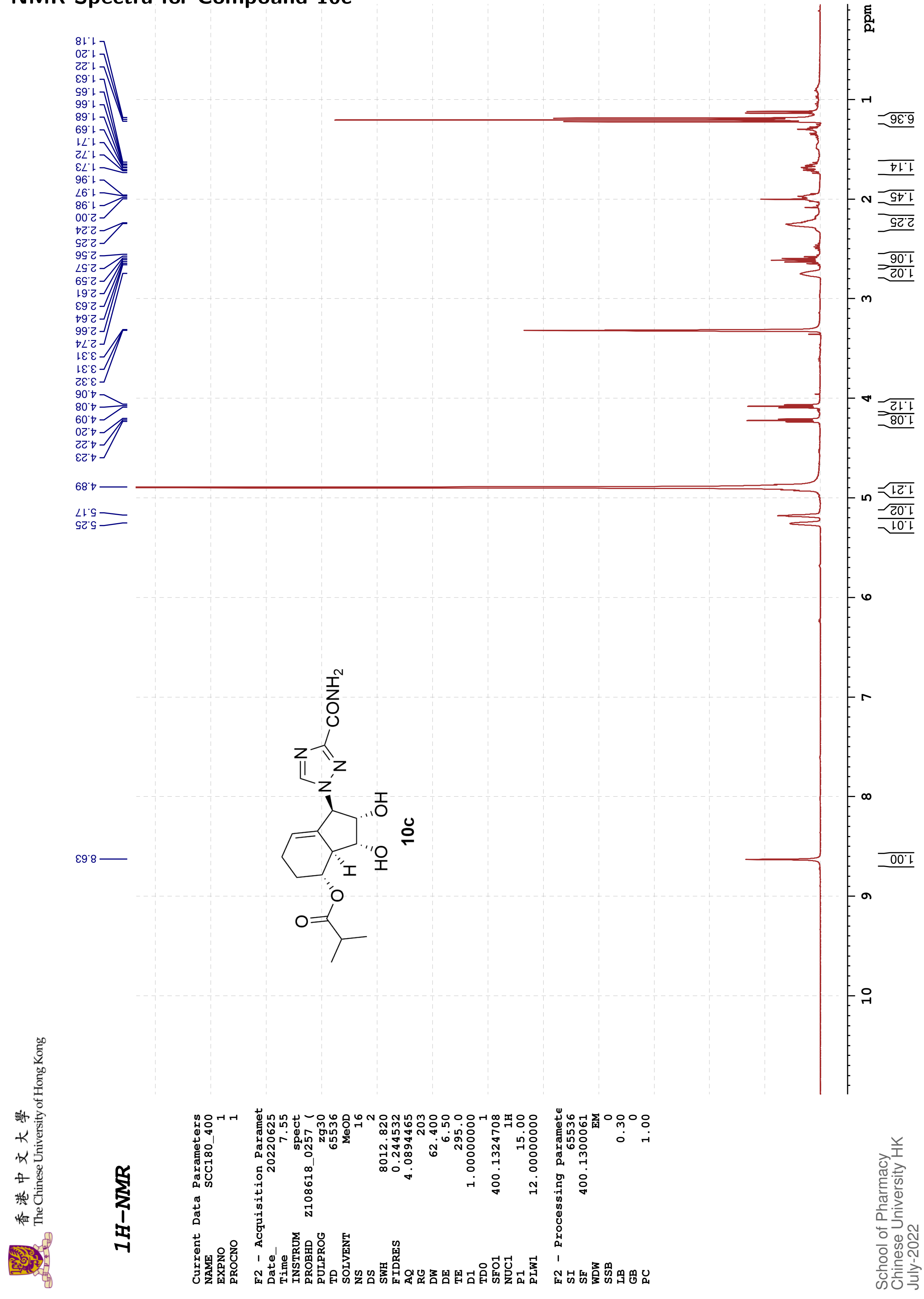

香港中文大學 The Chinese University of Hong Kong

**13C{1H}-NMR**

```
Current Data Parameters
NAME          SCC180_400
EXPNO                  2
PROCNO                 1

F2 - Acquisition Paramet
Date_           20220625
Time                8.21
INSTRUM            spect
PROBHD    Z108618_0257 (
PULPROG          zgpg30
TD                48074
SOLVENT             MeOD
NS                   487
DS                     4
SWH            24038.461
FIDRES          1.000061
AQ             0.9999392
RG                   203
DW                20.800
DE                  6.50
TE                 295.6
D1            2.00000000
D11           0.03000000
TD0                    1
SFO1         100.6228298
NUC1                 13C
P1                 10.00
PLW1         47.00000000
SFO2         400.1316005
NUC2                  1H
CPDPRG[2         waltz16
PCPD2              90.00
PLW2         12.00000000
PLW12         0.33333001
PLW13         0.16766000

F2 - Processing paramete
SI                 32768
SF           100.6126282
WDW                   EM
SSB                    0
LB                  0.40
GB                     0
PC                  1.40
```

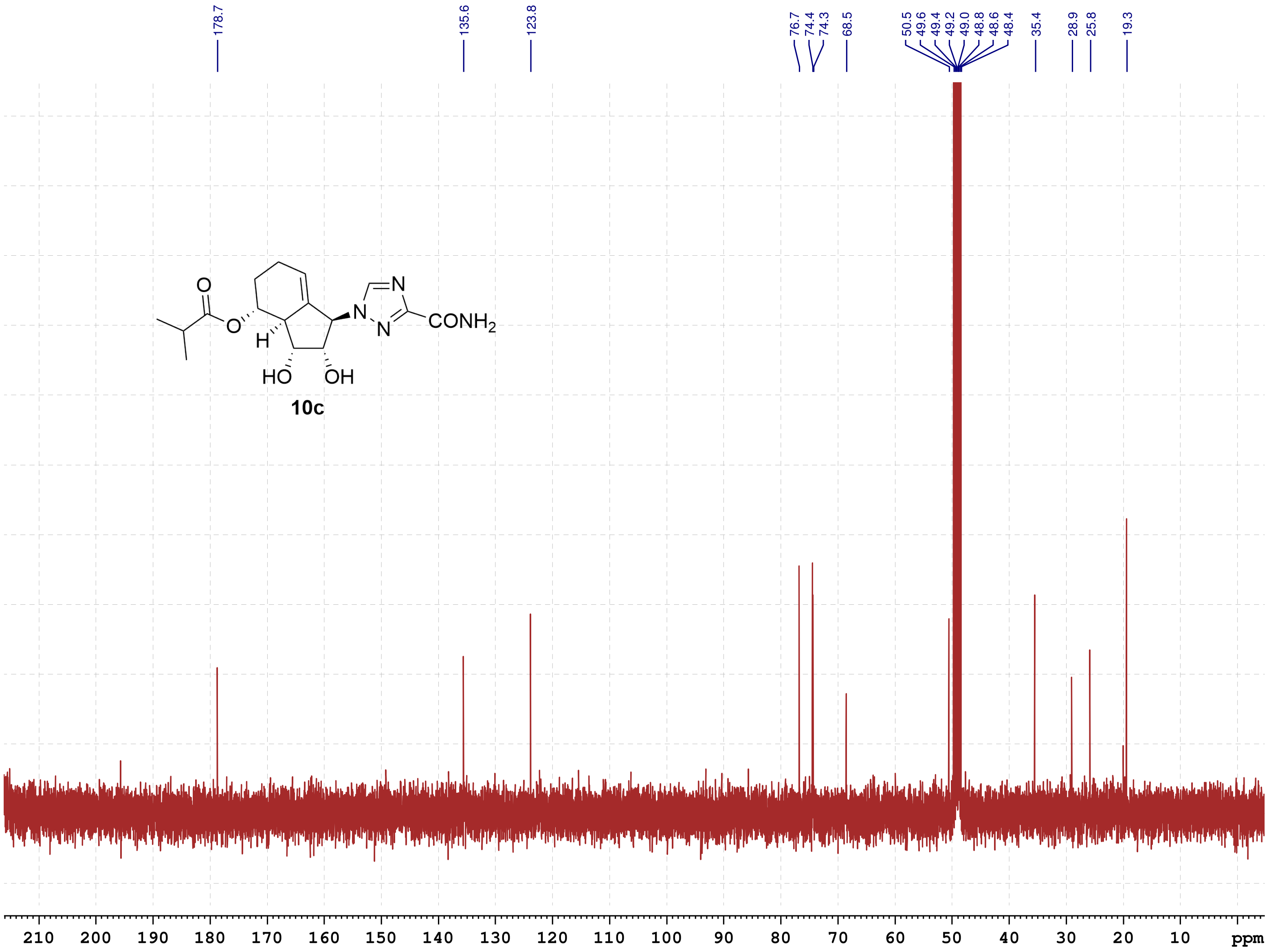

School of Pharmacy
Chinese University HK
July-2022

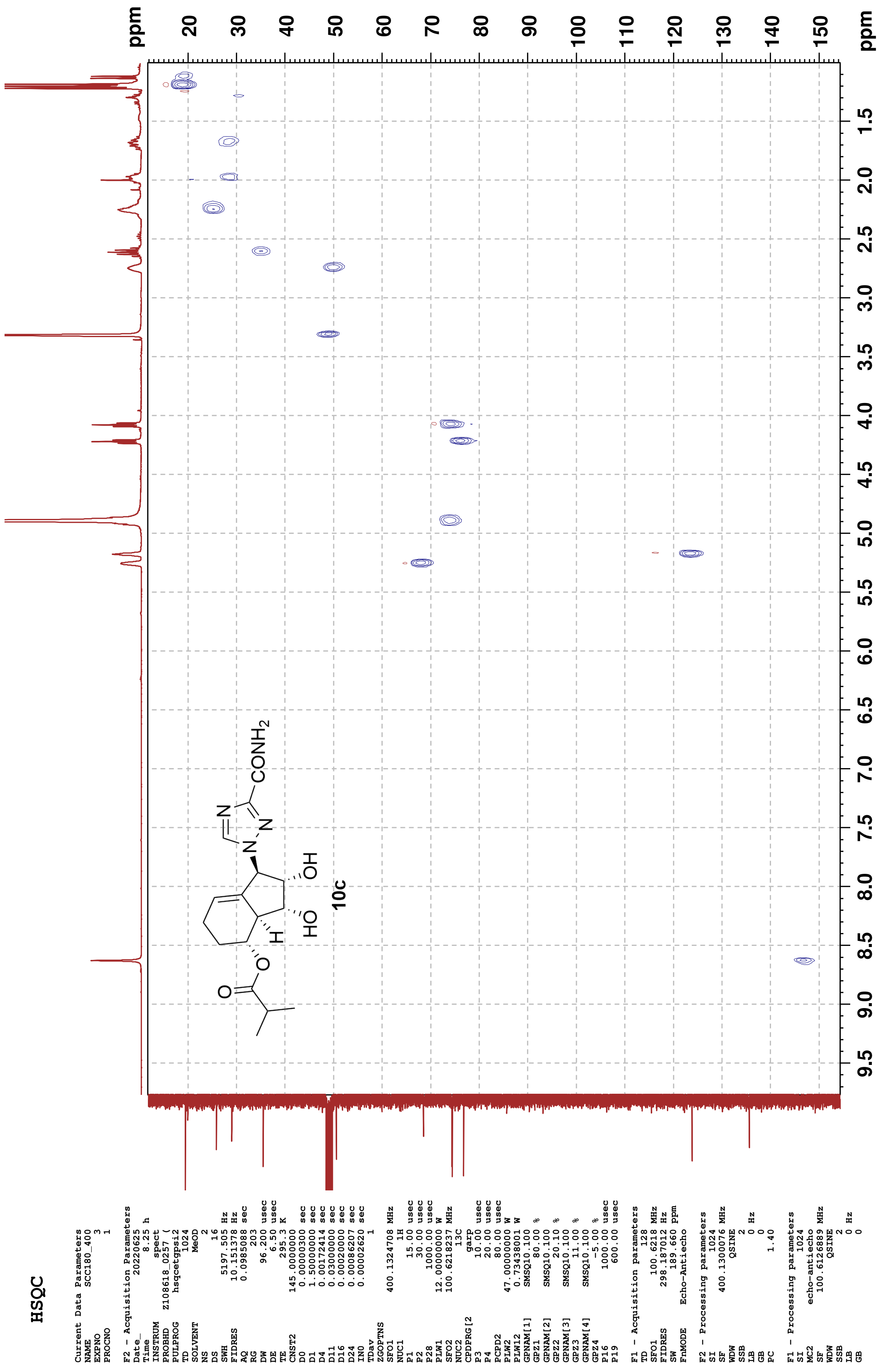
HSQC
Current Data Parameters
NAME SCC180_400
EXPNO 3
PROCNO 1
F2 - Acquisition Parameters
Date_ 20220625
Time 8.25 h
INSTRUM spect
PROBHD Z108618_0257 (
PULPROG hsqcetgpsi2
TD 1024
SOLVENT MeOD
NS 2
DS 16
SWH 5197.505 Hz
FIDRES 10.151378 Hz
AQ 0.0985088 sec
RG 203
DW 96.200 usec
DE 6.50 usec
TE 295.3 K
CNST2 145.0000000
D0 0.00000300 sec
D1 1.50000000 sec
D4 0.00172414 sec
D11 0.03000000 sec
D16 0.00020000 sec
D24 0.00086207 sec
IN0 0.00002620 sec
TDav 1
ZGOPTNS
SFO1 400.1324708 MHz
NUC1 1H
P1 15.00 usec
P2 30.00 usec
P28 1000.00 usec
PLW1 12.00000000 W
SFO2 100.6218237 MHz
NUC2 13C
CPDPRG[2 garp
P3 10.00 usec
P4 20.00 usec
PCPD2 80.00 usec
PLW2 47.00000000 W
PLW12 0.73438001 W
GPNAM[1] SMSQ10.100
GPZ1 80.00 %
GPNAM[2] SMSQ10.100
GPZ2 20.10 %
GPNAM[3] SMSQ10.100
GPZ3 11.00 %
GPNAM[4] SMSQ10.100
GPZ4 -5.00 %
P16 1000.00 usec
P19 600.00 usec
F1 - Acquisition parameters
TD 128
SFO1 100.6218 MHz
FIDRES 298.187012 Hz
SW 189.660 ppm
FnMODE Echo-Antiecho
F2 - Processing parameters
SI 1024
SF 400.1300076 MHz
WDW QSINE
SSB 2
LB 0 Hz
GB 0
PC 1.40
F1 - Processing parameters
SI 1024
MC2 echo-antiecho
SF 100.6126889 MHz
WDW QSINE
SSB 2
LB 0 Hz
GB 0
O
O
H
HO
OH
N
N
N
CONH2
10c
ppm
20
30
40
50
60
70
80
90
100
110
120
130
140
150
9.5
9.0
8.5
8.0
7.5
7.0
6.5
6.0
5.5
5.0
4.5
4.0
3.5
3.0
2.5
2.0
1.5
ppm

**HMBC**

```
Current Data Parameters
NAME          SCC180_400
EXPNO                  4
PROCNO                 1

F2 - Acquisition Parameters
Date_           20220625
Time                8.32 h
INSTRUM            spect
PROBHD   Z108618_0257 (
PULPROG     hmbcgplpndqf
TD                  2048
SOLVENT             MeOD
NS                     6
DS                    16
SWH             5197.505 Hz
FIDRES          5.075689 Hz
AQ             0.1970176 sec
RG                   203
DW                96.200 usec
DE                  6.50 usec
TE                 295.2 K
CNST2        145.0000000
CNST13        10.0000000
D0            0.00000300 sec
D1            1.50000000 sec
D2            0.00344828 sec
D6            0.05000000 sec
D16           0.00020000 sec
IN0           0.00002260 sec
TDav                   1
SFO1         400.1325208 MHz
NUC1                  1H
P1                 15.00 usec
P2                 30.00 usec
PLW1         12.00000000 W
SFO2         100.6228298 MHz
NUC2                 13C
P3                 10.00 usec
PLW2         47.00000000 W
GPNAM[1]      SMSQ10.100
GPZ1               50.00 %
GPNAM[2]      SMSQ10.100
GPZ2               30.00 %
GPNAM[3]      SMSQ10.100
GPZ3               40.10 %
P16              1000.00 usec

F1 - Acquisition parameters
TD                    64
SFO1            100.6228 MHz
FIDRES        691.371704 Hz
SW               219.870 ppm
FnMODE                QF

F2 - Processing parameters
SI                  2048
SF           400.1300060 MHz
WDW                 SINE
SSB                    0
LB                     0 Hz
GB                     0
PC                  1.40

F1 - Processing parameters
SI                  1024
MC2                   QF
SF           100.6126810 MHz
WDW                 SINE
SSB                    0
LB                     0 Hz
GB                     0
```

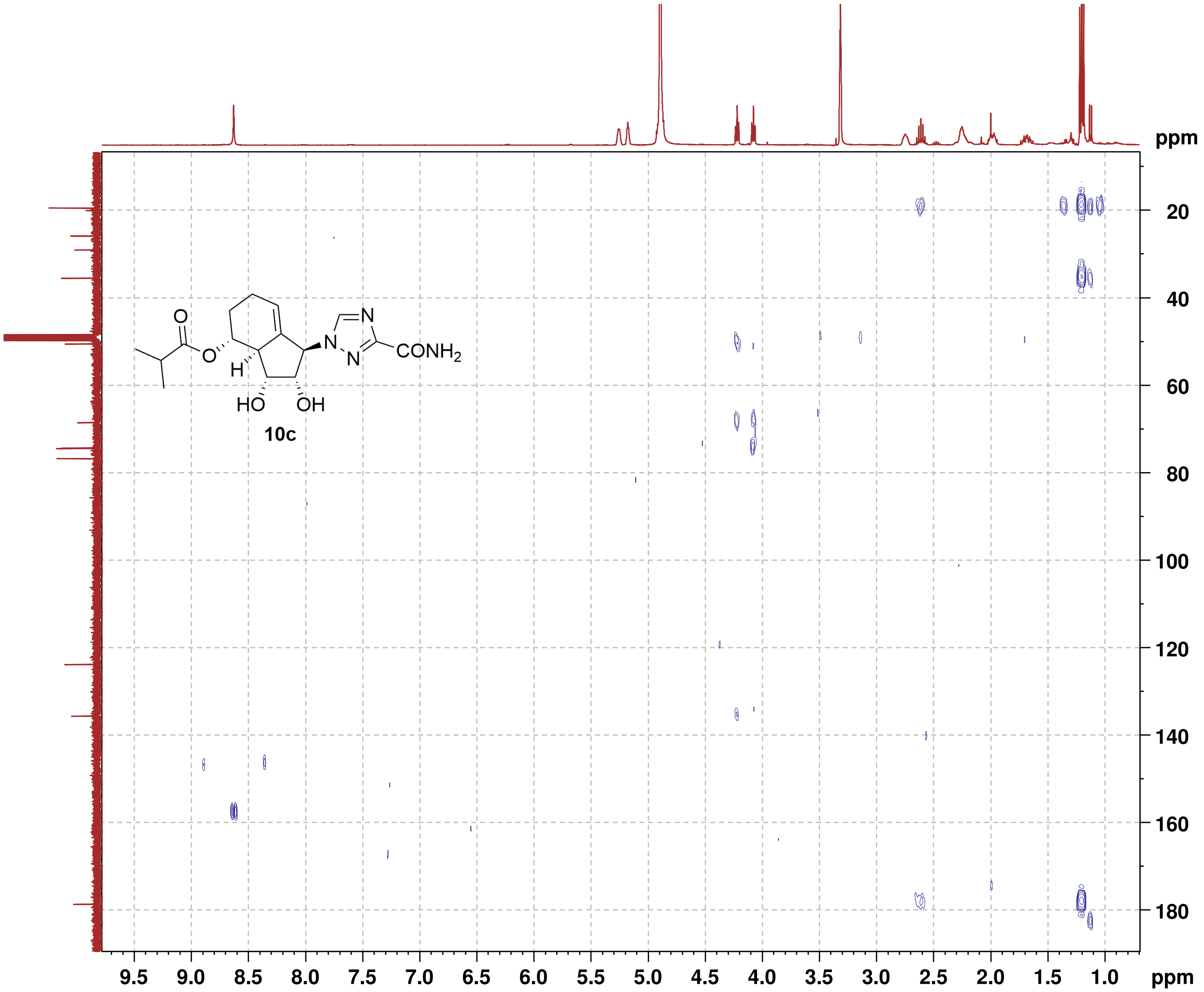

#### NMR-Spectra for Compound 8a

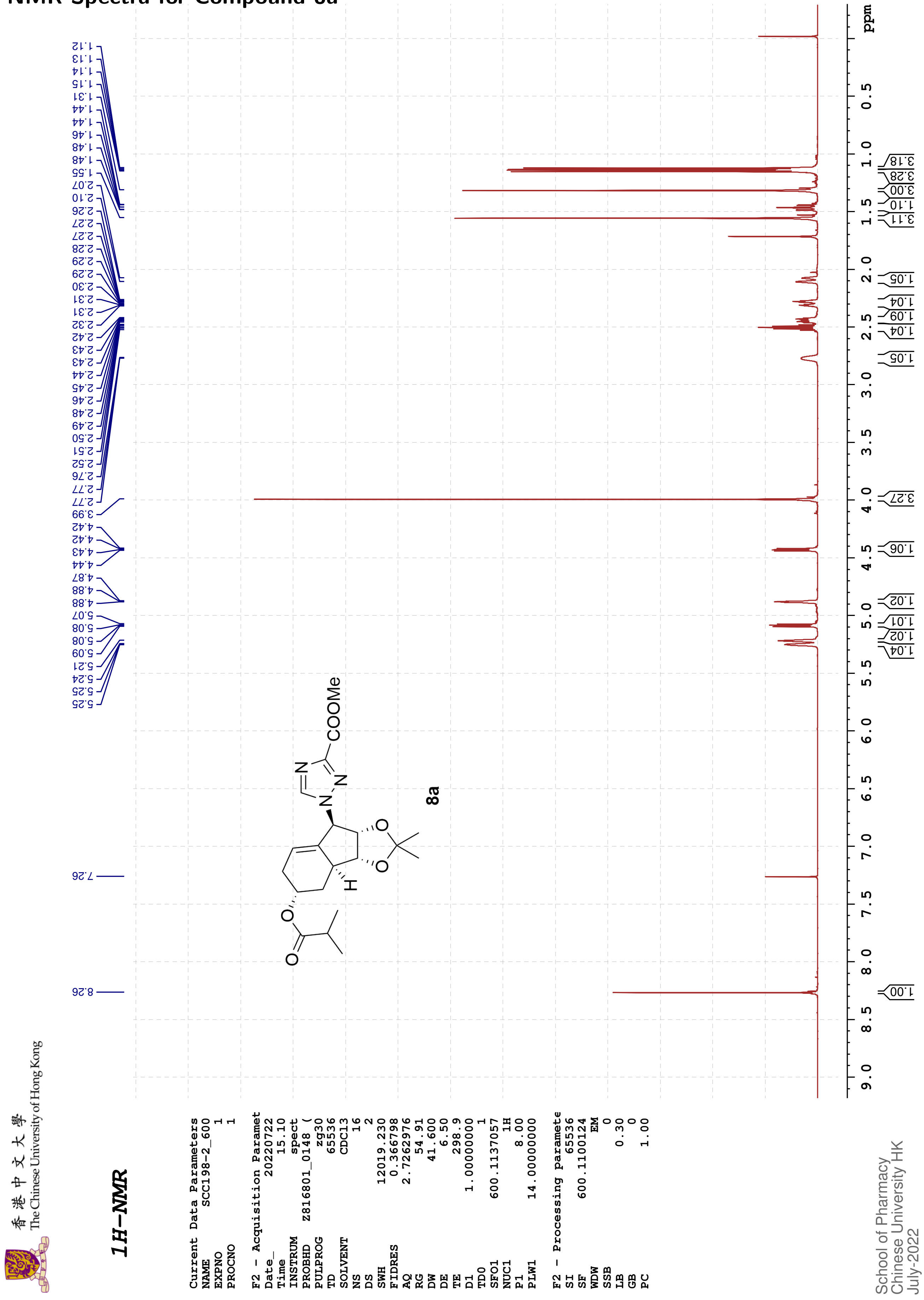

香港中文大學
The Chinese University of Hong Kong

#### 13C{1H}-NMR

```
Current Data Parameters
NAME          SCC198-2_600
EXPNO                    2
PROCNO                   1

F2 - Acquisition Paramet
Date_             20220722
Time                 15.17
INSTRUM              spect
PROBHD     Z816801_0148 (
PULPROG            zgpg30
TD                   65536
SOLVENT              CDCl3
NS                     103
DS                       4
SWH              36231.883
FIDRES            1.105709
AQ               0.9043968
RG                   60.39
DW                  13.800
DE                    6.50
TE                   299.1
D1              2.00000000
D11             0.03000000
TD0                      1
SFO1           150.9128693
NUC1                   13C
P1                   12.00
PLW1          183.00000000
SFO2           600.1124004
NUC2                    1H
CPDPRG[2           waltz16
PCPD2                70.00
PLW2           14.00000000
PLW12           0.18286000
PLW13           0.09197600

F2 - Processing paramete
SI                   32768
SF             150.8977618
WDW                     EM
SSB                      0
LB                    2.00
GB                       0
PC                    1.40
```

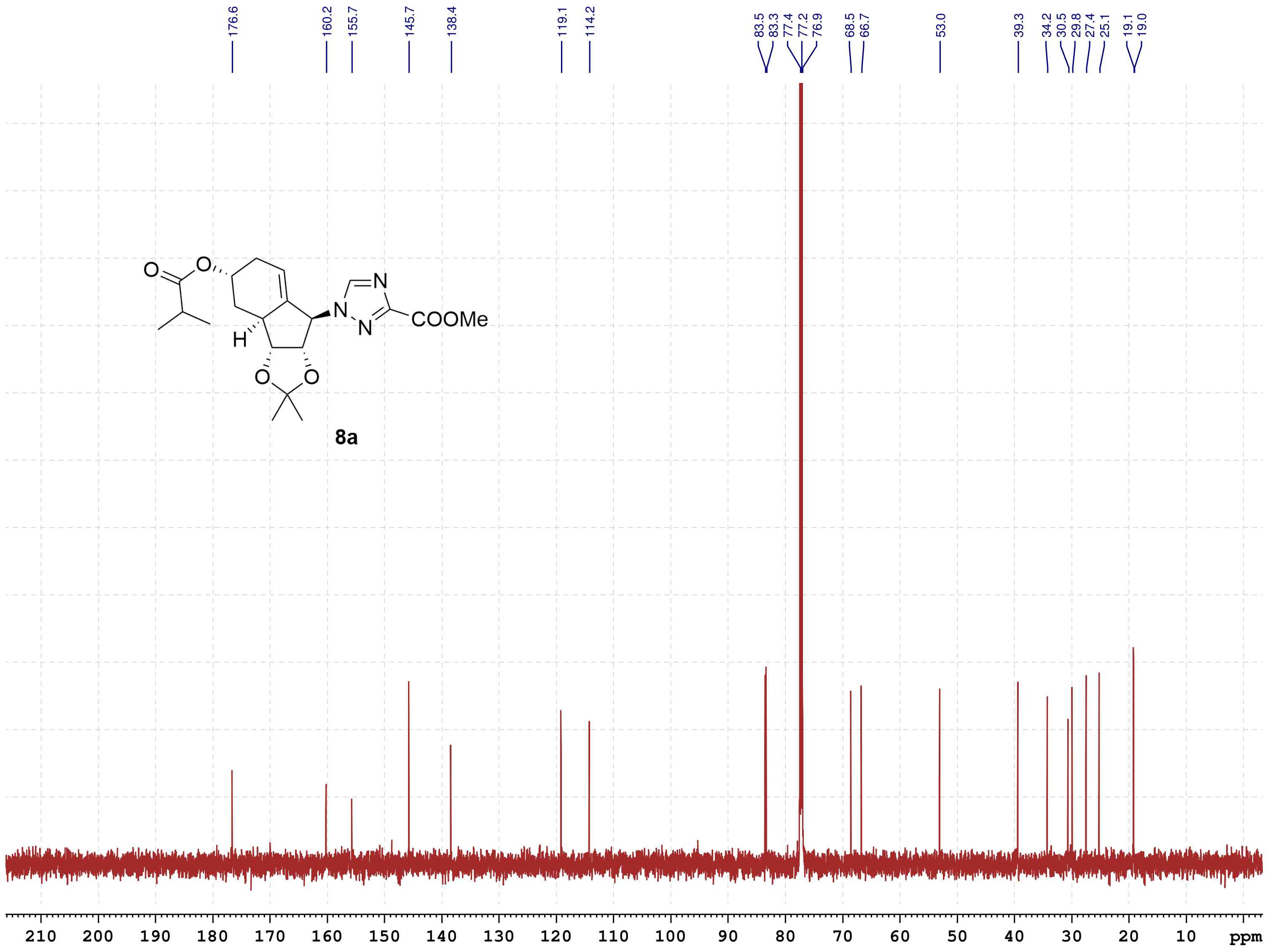

School of Pharmacy
Chinese University HK
July-2022

#### NMR-Spectra for Compound 11a

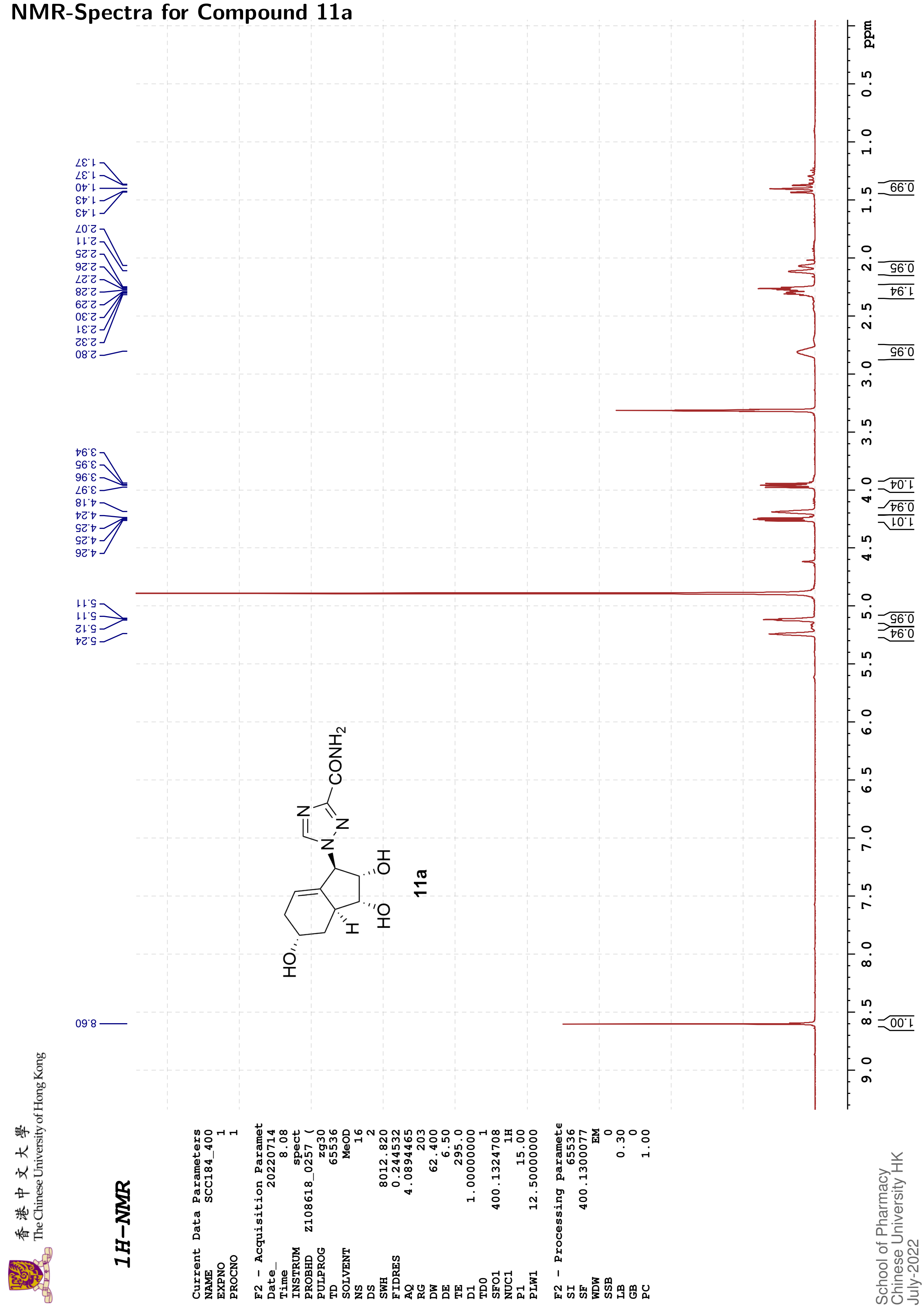

香港中文大學
The Chinese University of Hong Kong

#### 13C{1H}-NMR

```
Current Data Parameters
NAME          SCC184_400
EXPNO                  2
PROCNO                 1

F2 - Acquisition Paramet
Date_           20220714
Time                8.20
INSTRUM            spect
PROBHD    Z108618_0257 (
PULPROG          zgpg30
TD                48074
SOLVENT            MeOD
NS                  200
DS                    4
SWH           24038.461
FIDRES         1.000061
AQ            0.9999392
RG                  203
DW               20.800
DE                 6.50
TE                295.5
D1           2.00000000
D11          0.03000000
TD0                   1
SFO1        100.6228298
NUC1                13C
P1                10.00
PLW1        51.00000000
SFO2        400.1316005
NUC2                 1H
CPDPRG[2        waltz16
PCPD2             90.00
PLW2        12.50000000
PLW12        0.34722000
PLW13        0.17465000

F2 - Processing paramete
SI                32768
SF          100.6126282
WDW                  EM
SSB                   0
LB                 0.75
GB                    0
PC                 1.40
```

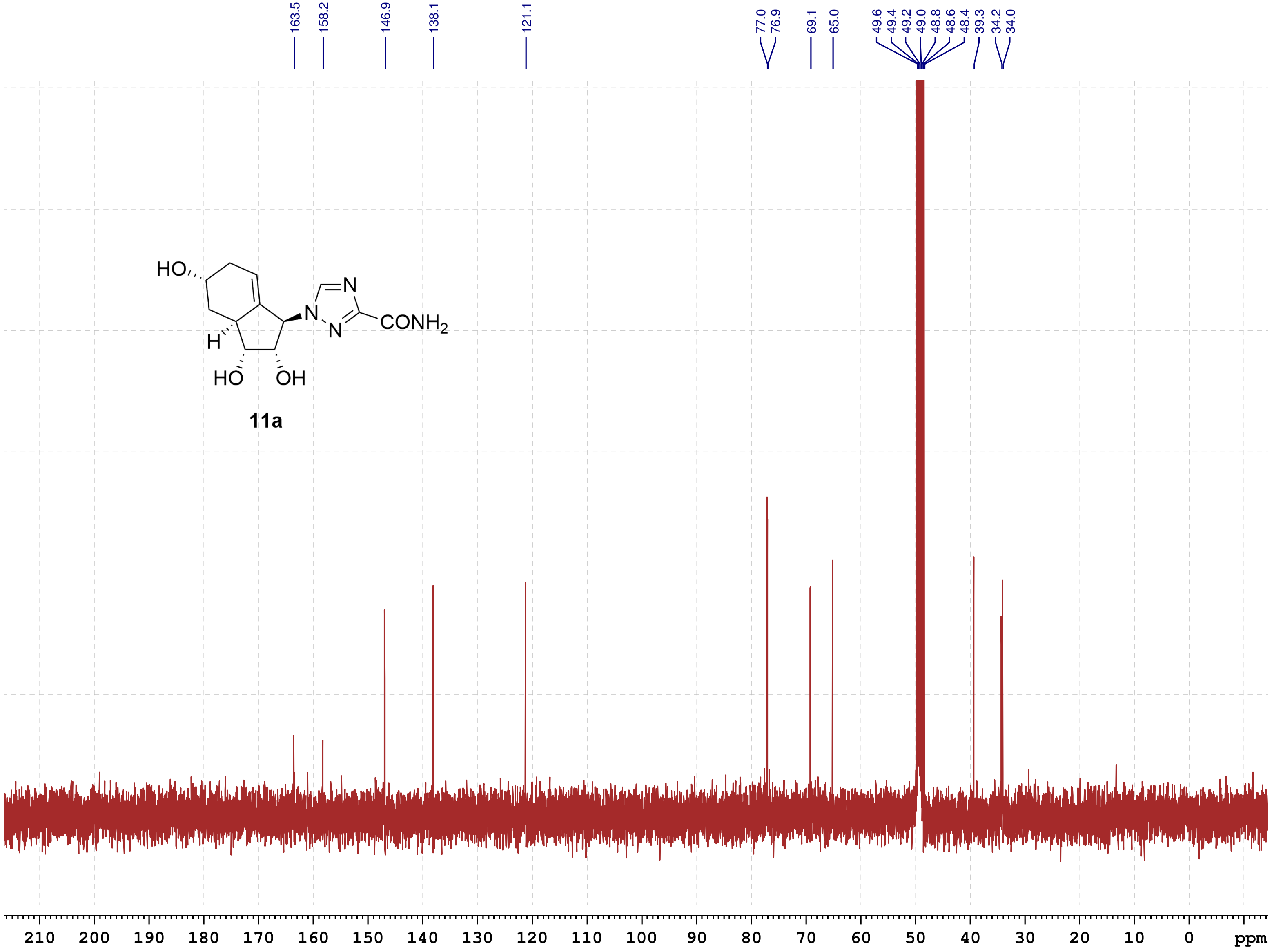

School of Pharmacy
Chinese University HK
July-2022

香港中文大學 The Chinese University of Hong Kong

**COSY**

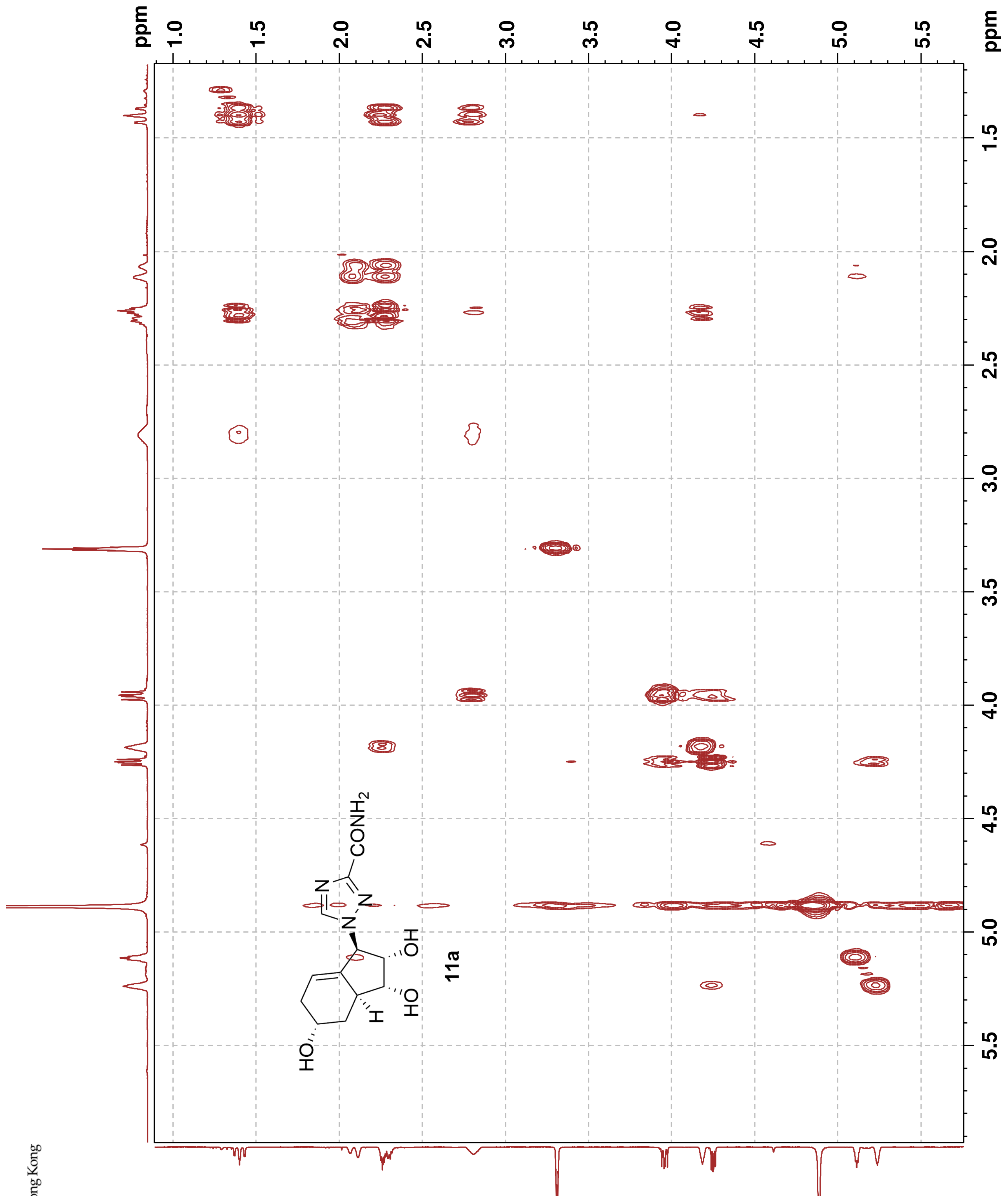

Current Data Parameters
NAME SCC184_400
EXPNO 3
PROCNO 1

F2 - Acquisition Parameters
Date_ 20220714
Time 8.21 h
INSTRUM spect
PROBHD Z108618_0257 (
PULPROG cosygpqf
TD 2048
SOLVENT MeOD
NS 2
DS 4
SWH 4000.000 Hz
FIDRES 3.906250 Hz
AQ 0.2560000 sec
RG 203
DW 125.000 usec
DE 6.50 usec
TE 295.4 K
D0 0.00000300 sec
D1 1.48689198 sec
D13 0.00000400 sec
D16 0.00020000 sec
IN0 0.00025000 sec
TDav 1
SFO1 400.1320006 MHz
NUC1 1H
P0 15.00 usec
P1 15.00 usec
PLW1 12.50000000 W
GPNAM[1] SINE.100
GPZ1 10.00 %
P16 1000.00 usec

F1 - Acquisition parameters
TD 100
SFO1 400.132 MHz
FIDRES 80.000000 Hz
SW 9.997 ppm
FnMODE QF

F2 - Processing parameters
SI 1024
SF 400.1300097 MHz
WDW SINE
SSB 0
LB 0 Hz
GB 0
PC 1.40

F1 - Processing parameters
SI 1024
MC2 QF
SF 400.1300081 MHz
WDW SINE
SSB 0
LB 0 Hz
GB 0

香港中文大學
The Chinese University of Hong Kong

**NOESY**

Current Data Parameters

| | |
|---|---|
| NAME | SCC184_400 |
| EXPNO | 4 |
| PROCNO | 1 |

F2 - Acquisition Parameters

| | |
|---|---|
| Date_ | 20220714 |
| Time | 8.27 h |
| INSTRUM | spect |
| PROBHD | Z108618_0257 ( |
| PULPROG | noesygpphpp |
| TD | 2048 |
| SOLVENT | MeOD |
| NS | 2 |
| DS | 4 |
| SWH | 4000.000 Hz |
| FIDRES | 3.906250 Hz |
| AQ | 0.2560000 sec |
| RG | 203 |
| DW | 125.000 usec |
| DE | 6.50 usec |
| TE | 295.1 K |
| D0 | 0.00010590 sec |
| D1 | 2.00000000 sec |
| D8 | 0.85000002 sec |
| D11 | 0.03000000 sec |
| D12 | 0.00002000 sec |
| D16 | 0.00020000 sec |
| IN0 | 0.00025000 sec |
| TDav | 1 |
| SFO1 | 400.1318806 MHz |
| NUC1 | 1H |
| P1 | 15.00 usec |
| P2 | 30.00 usec |
| P17 | 2500.00 usec |
| PLW1 | 12.50000000 W |
| PLW10 | 3.12500000 W |
| GPNAM[1] | SMSQ10.100 |
| GPZ1 | 40.00 % |
| P16 | 1000.00 usec |

F1 - Acquisition parameters

| | |
|---|---|
| TD | 128 |
| SFO1 | 400.1319 MHz |
| FIDRES | 62.500000 Hz |
| SW | 9.997 ppm |
| FnMODE | States-TPPI |

F2 - Processing parameters

| | |
|---|---|
| SI | 1024 |
| SF | 400.1300105 MHz |
| WDW | QSINE |
| SSB | 2 |
| LB | 0 Hz |
| GB | 0 |
| PC | 1.00 |

F1 - Processing parameters

| | |
|---|---|
| SI | 1024 |
| MC2 | States-TPPI |
| SF | 400.1300093 MHz |
| WDW | QSINE |
| SSB | 2 |
| LB | 0 Hz |
| GB | 0 |

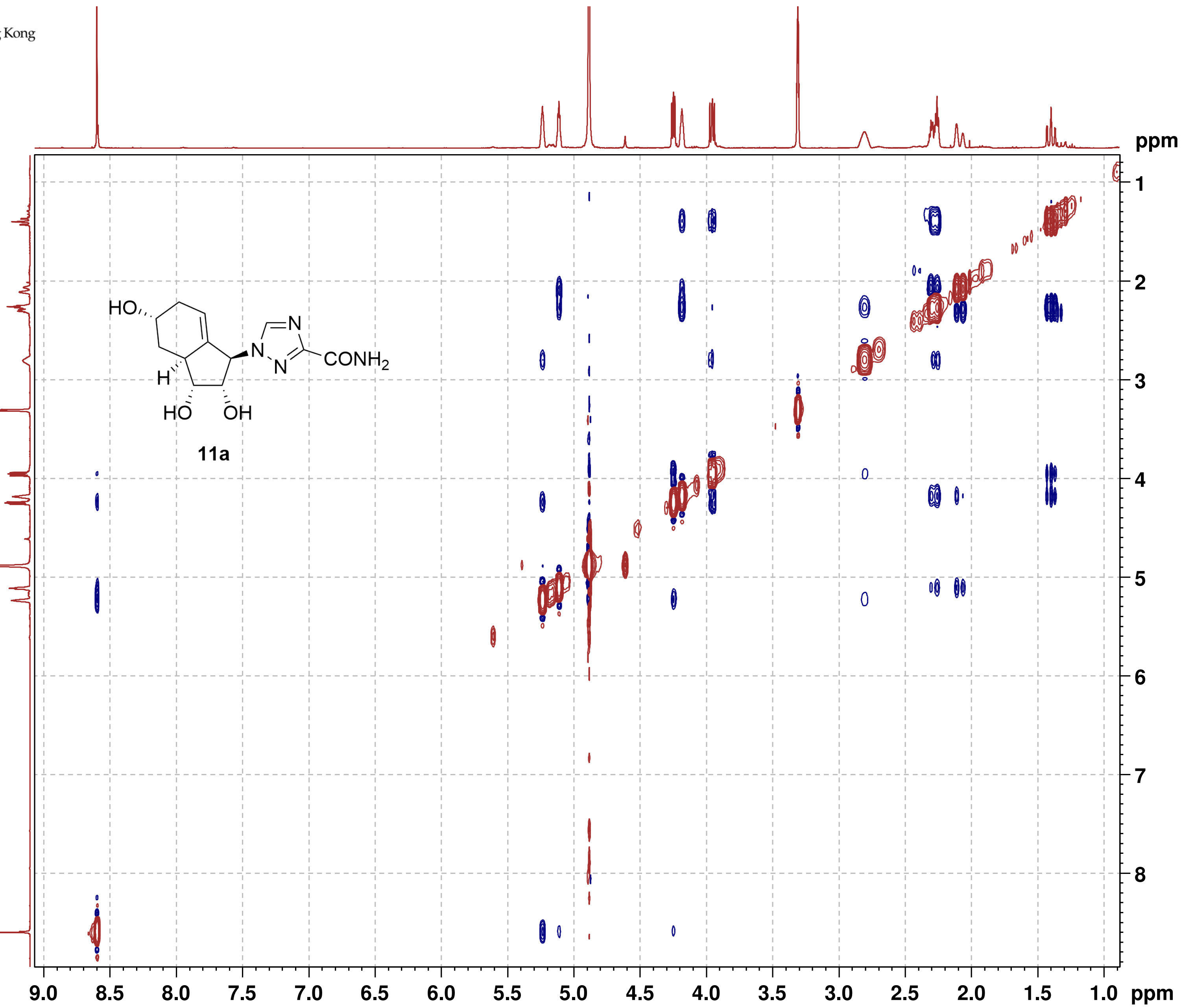

11a

#### NMR-Spectra for Compound 11c

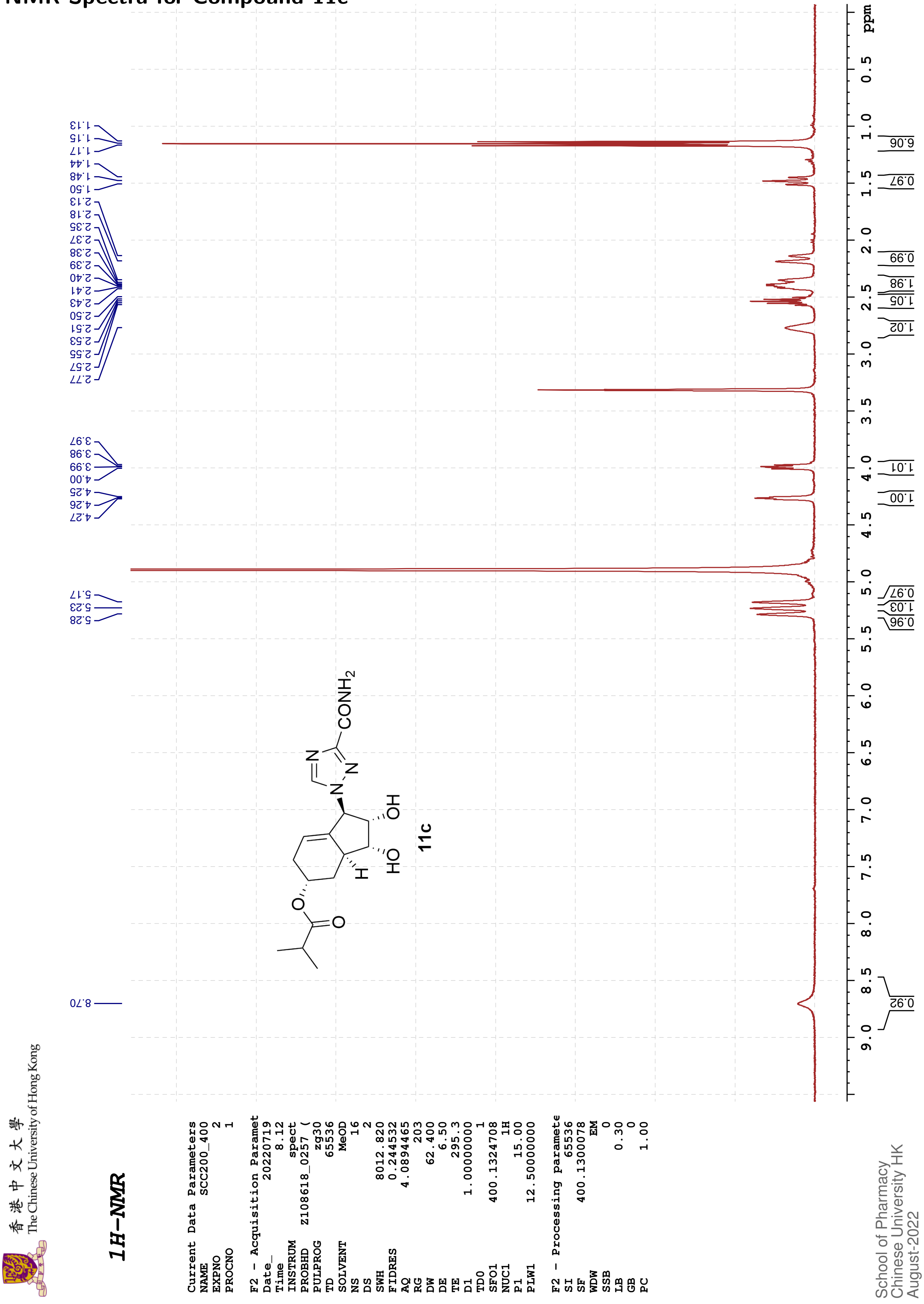

香港中文大學 The Chinese University of Hong Kong

#### 13C{1H}-NMR

```
Current Data Parameters
NAME              SCC200_400
EXPNO                      1
PROCNO                     1

F2 - Acquisition Paramet
Date_               20220719
Time                    8.25
INSTRUM                spect
PROBHD       Z108618_0257 (
PULPROG              zgpg30
TD                     48074
SOLVENT                 MeOD
NS                       224
DS                         4
SWH                24038.461
FIDRES              1.000061
AQ                 0.9999392
RG                       203
DW                    20.800
DE                      6.50
TE                     295.5
D1               2.00000000
D11              0.03000000
TD0                        1
SFO1             100.6228298
NUC1                     13C
P1                     10.00
PLW1             51.00000000
SFO2             400.1316005
NUC2                      1H
CPDPRG[2             waltz16
PCPD2                  90.00
PLW2             12.50000000
PLW12             0.34722000
PLW13             0.17465000

F2 - Processing paramete
SI                     32768
SF               100.6126288
WDW                       EM
SSB                        0
LB                      1.00
GB                         0
PC                      1.40
```

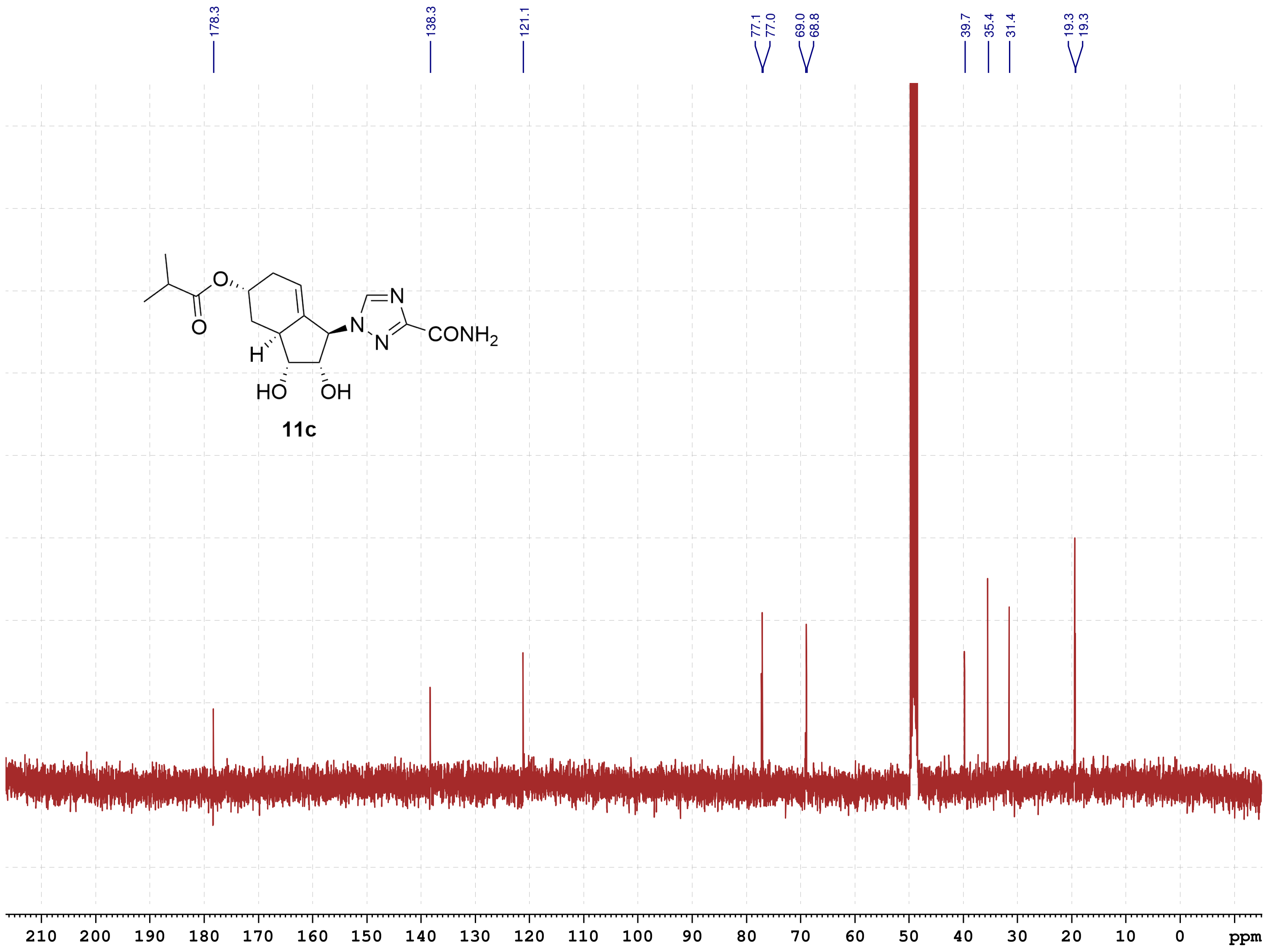

School of Pharmacy
Chinese University HK
August-2022

#### NMR-Spectra for Compound 9a

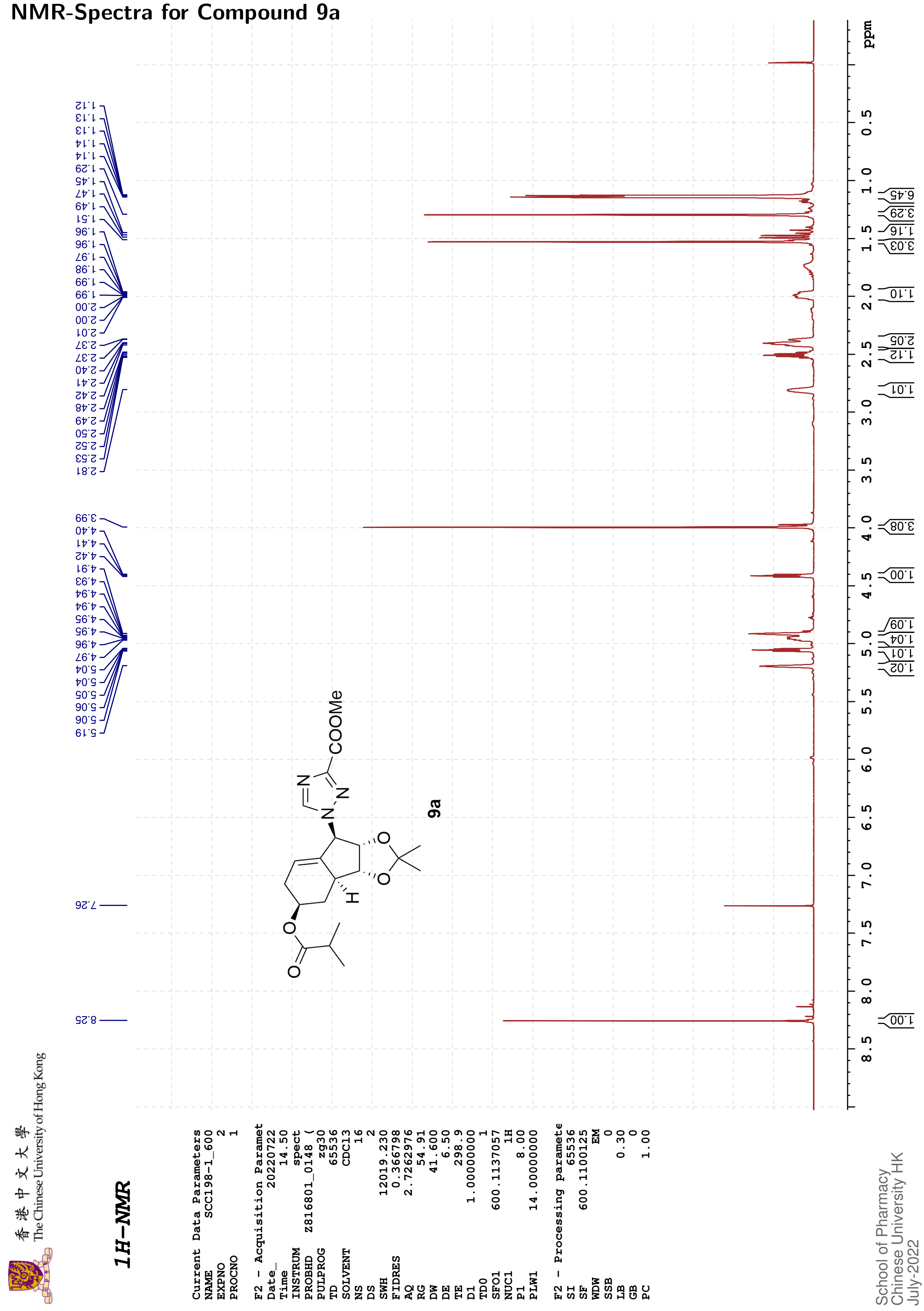

The Chinese University of Hong Kong

香港中文大學

#### 13C{1H}-NMR

| | |
|---|---|
| Current Data Parameters | |
| NAME | SCC198-1_600 |
| EXPNO | 3 |
| PROCNO | 1 |
| F2 - Acquisition Paramet | |
| Date_ | 20220722 |
| Time | 15.01 |
| INSTRUM | spect |
| PROBHD | Z816801_0148 ( |
| PULPROG | zgpg30 |
| TD | 65536 |
| SOLVENT | CDCl3 |
| NS | 147 |
| DS | 4 |
| SWH | 36231.883 |
| FIDRES | 1.105709 |
| AQ | 0.9043968 |
| RG | 60.39 |
| DW | 13.800 |
| DE | 6.50 |
| TE | 299.1 |
| D1 | 2.00000000 |
| D11 | 0.03000000 |
| TD0 | 1 |
| SFO1 | 150.9128693 |
| NUC1 | 13C |
| P1 | 12.00 |
| PLW1 | 183.00000000 |
| SFO2 | 600.1124004 |
| NUC2 | 1H |
| CPDPRG[2 | waltz16 |
| PCPD2 | 70.00 |
| PLW2 | 14.00000000 |
| PLW12 | 0.18286000 |
| PLW13 | 0.09197600 |
| F2 - Processing paramete | |
| SI | 32768 |
| SF | 150.8977618 |
| WDW | EM |
| SSB | 0 |
| LB | 2.00 |
| GB | 0 |
| PC | 1.40 |

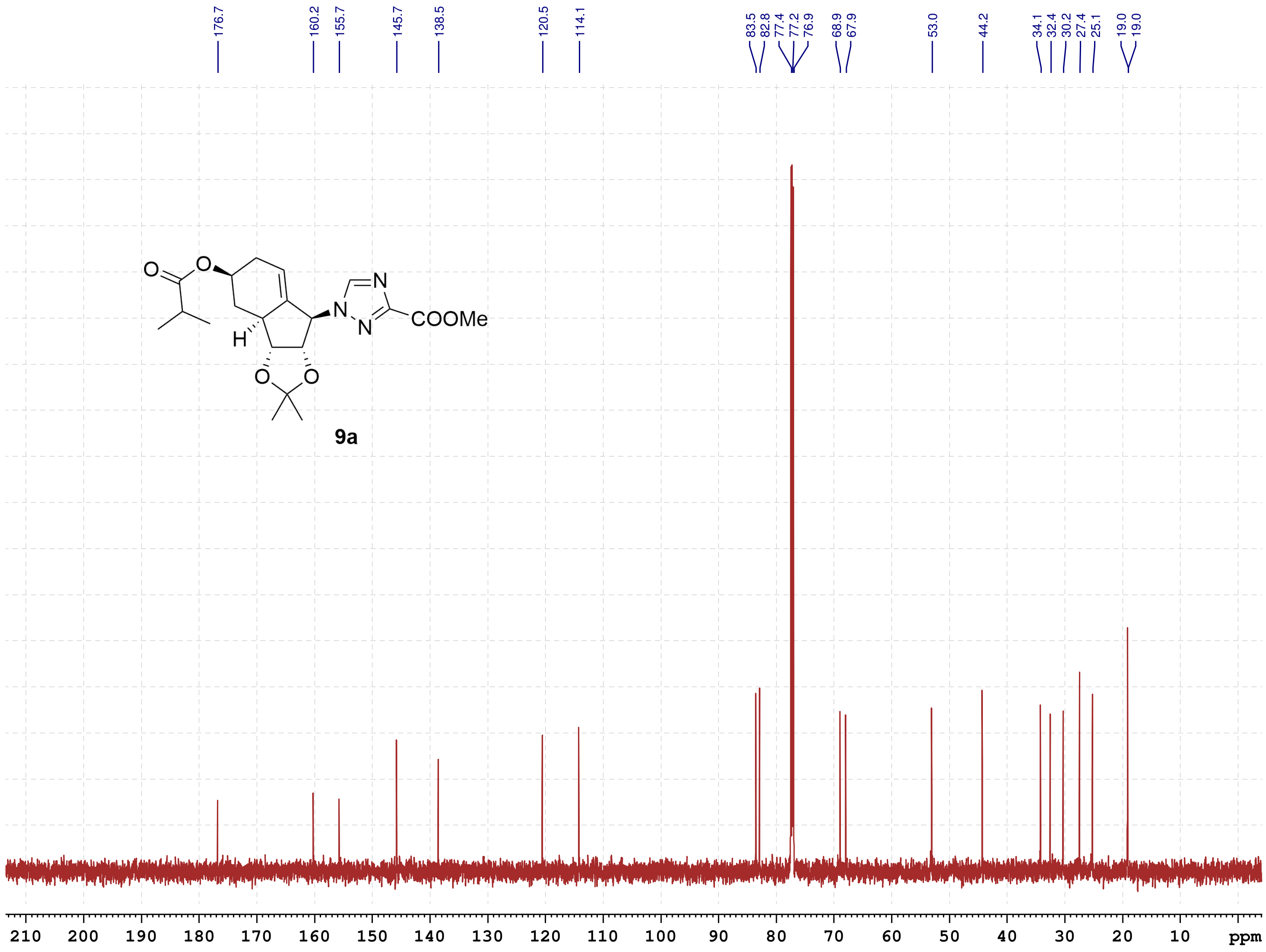

School of Pharmacy
Chinese University HK
July-2022

#### NMR-Spectra for Compound 12a

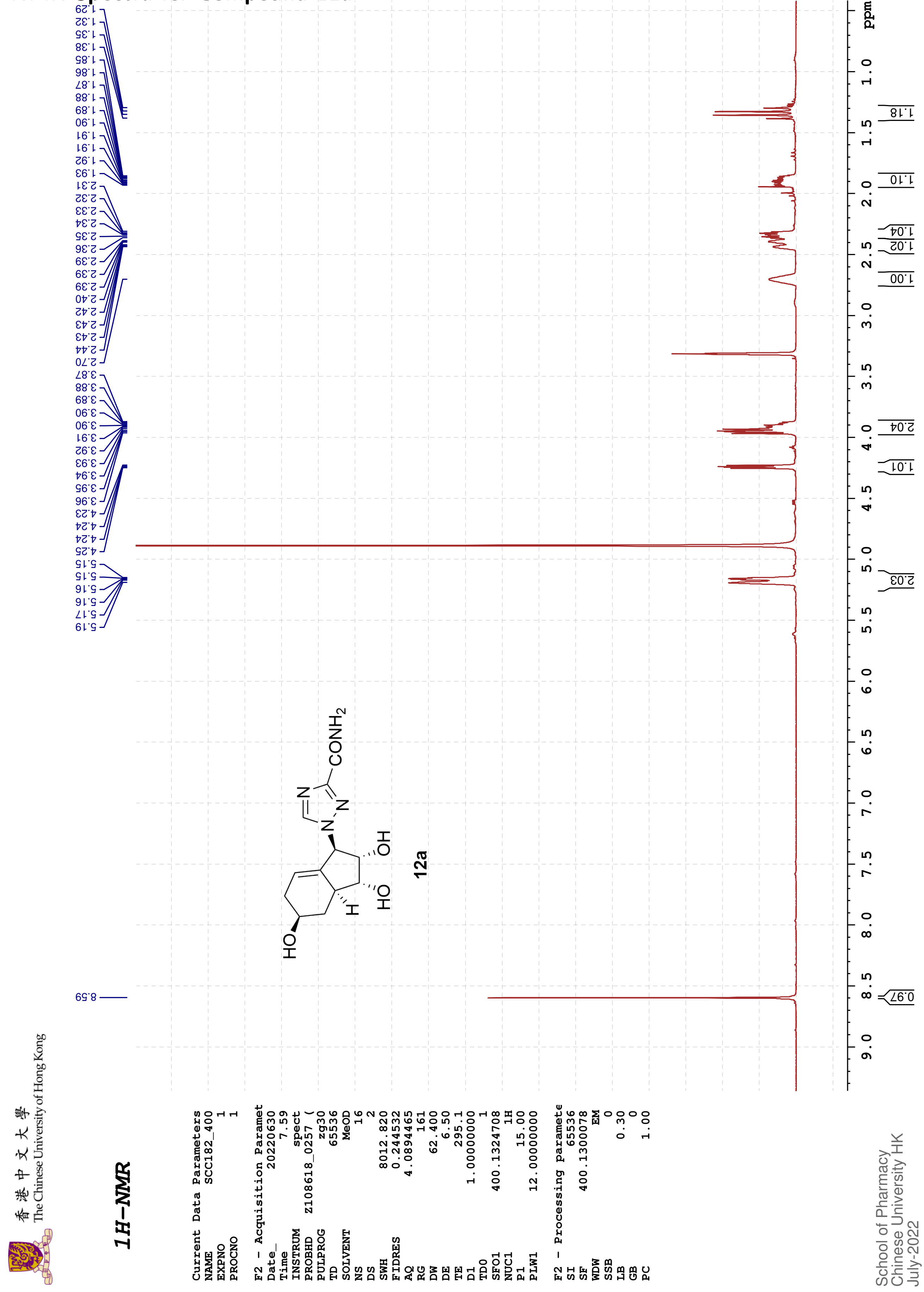

The Chinese University of Hong Kong 香港中文大學

#### 13C{1H}-NMR

Current Data Parameters
NAME SCC182_400
EXPNO 4
PROCNO 1

F2 - Acquisition Paramet
Date_ 20220630
Time 8.07
INSTRUM spect
PROBHD Z108618_0257 (
PULPROG zgpg30
TD 48074
SOLVENT MeOD
NS 115
DS 4
SWH 24038.461
FIDRES 1.000061
AQ 0.9999392
RG 203
DW 20.800
DE 6.50
TE 295.4
D1 2.00000000
D11 0.03000000
TD0 1
SFO1 100.6228298
NUC1 13C
P1 10.00
PLW1 47.00000000
SFO2 400.1316005
NUC2 1H
CPDPRG[2 waltz16
PCPD2 90.00
PLW2 12.00000000
PLW12 0.33333001
PLW13 0.16766000

F2 - Processing paramete
SI 32768
SF 100.6126289
WDW EM
SSB 0
LB 1.00
GB 0
PC 1.40

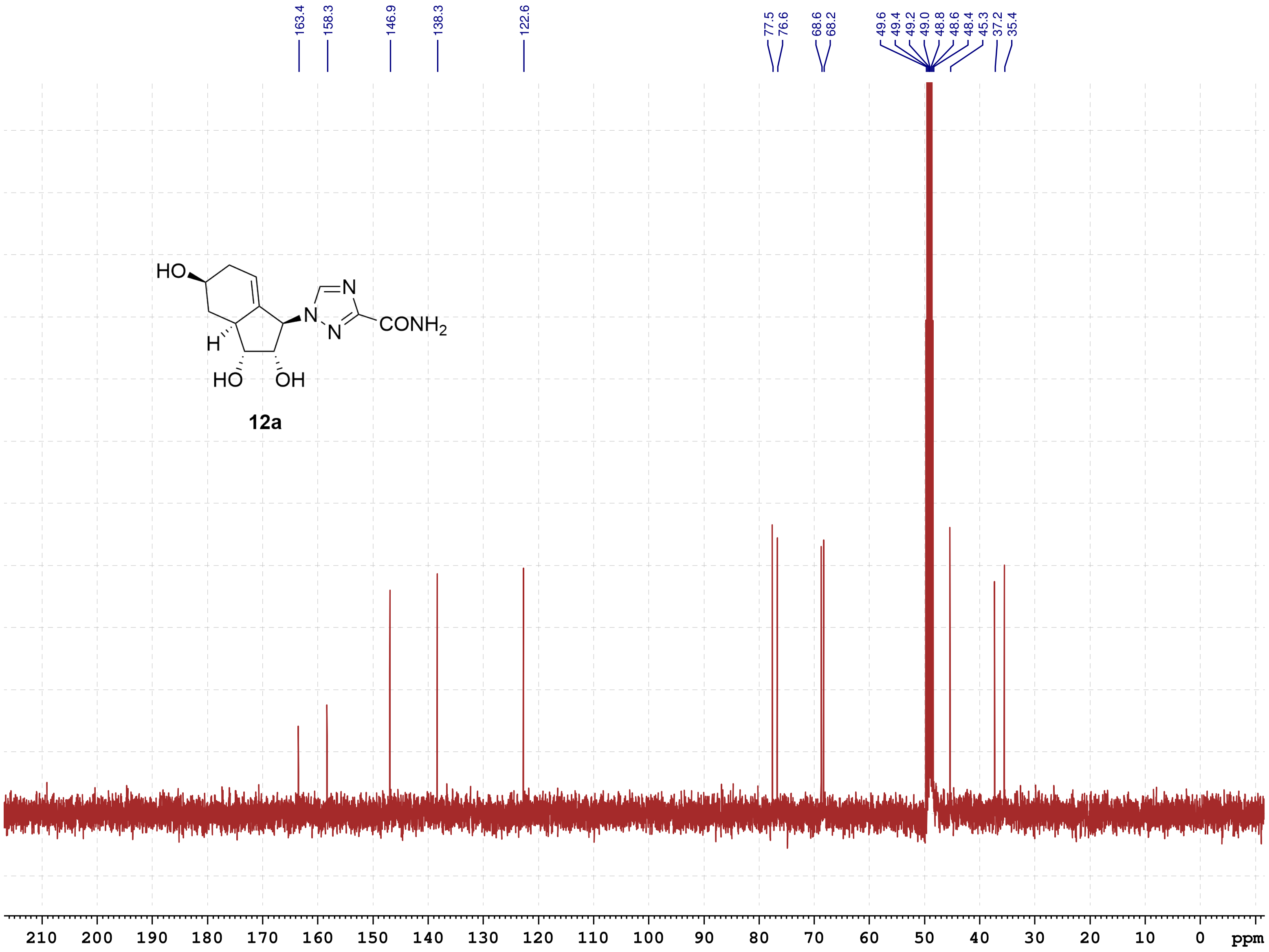

School of Pharmacy
Chinese University HK
July-2022

香港中文大學
The Chinese University of Hong Kong

**COSY**

```
Current Data Parameters
NAME          SCC182_400
EXPNO                  2
PROCNO                 1

F2 - Acquisition Parameters
Date_           20220630
Time                8.08 h
INSTRUM            spect
PROBHD     Z108618_0257 (
PULPROG         cosygpqf
TD                  2048
SOLVENT             MeOD
NS                     2
DS                     4
SWH             4000.000 Hz
FIDRES          3.906250 Hz
AQ             0.2560000 sec
RG                   203
DW               125.000 usec
DE                  6.50 usec
TE                 295.4 K
D0            0.00000300 sec
D1            1.48689198 sec
D13           0.00000400 sec
D16           0.00020000 sec
IN0           0.00025000 sec
TDav                   1
SFO1         400.1320006 MHz
NUC1                  1H
P0                 15.00 usec
P1                 15.00 usec
PLW1         12.00000000 W
GPNAM[1]        SINE.100
GPZ1               10.00 %
P16              1000.00 usec

F1 - Acquisition parameters
TD                   100
SFO1             400.132 MHz
FIDRES         80.000000 Hz
SW                 9.997 ppm
FnMODE                QF

F2 - Processing parameters
SI                  1024
SF           400.1300097 MHz
WDW                 SINE
SSB                    0
LB                     0 Hz
GB                     0
PC                  1.40

F1 - Processing parameters
SI                  1024
MC2                   QF
SF           400.1300081 MHz
WDW                 SINE
SSB                    0
LB                     0 Hz
GB                     0
```

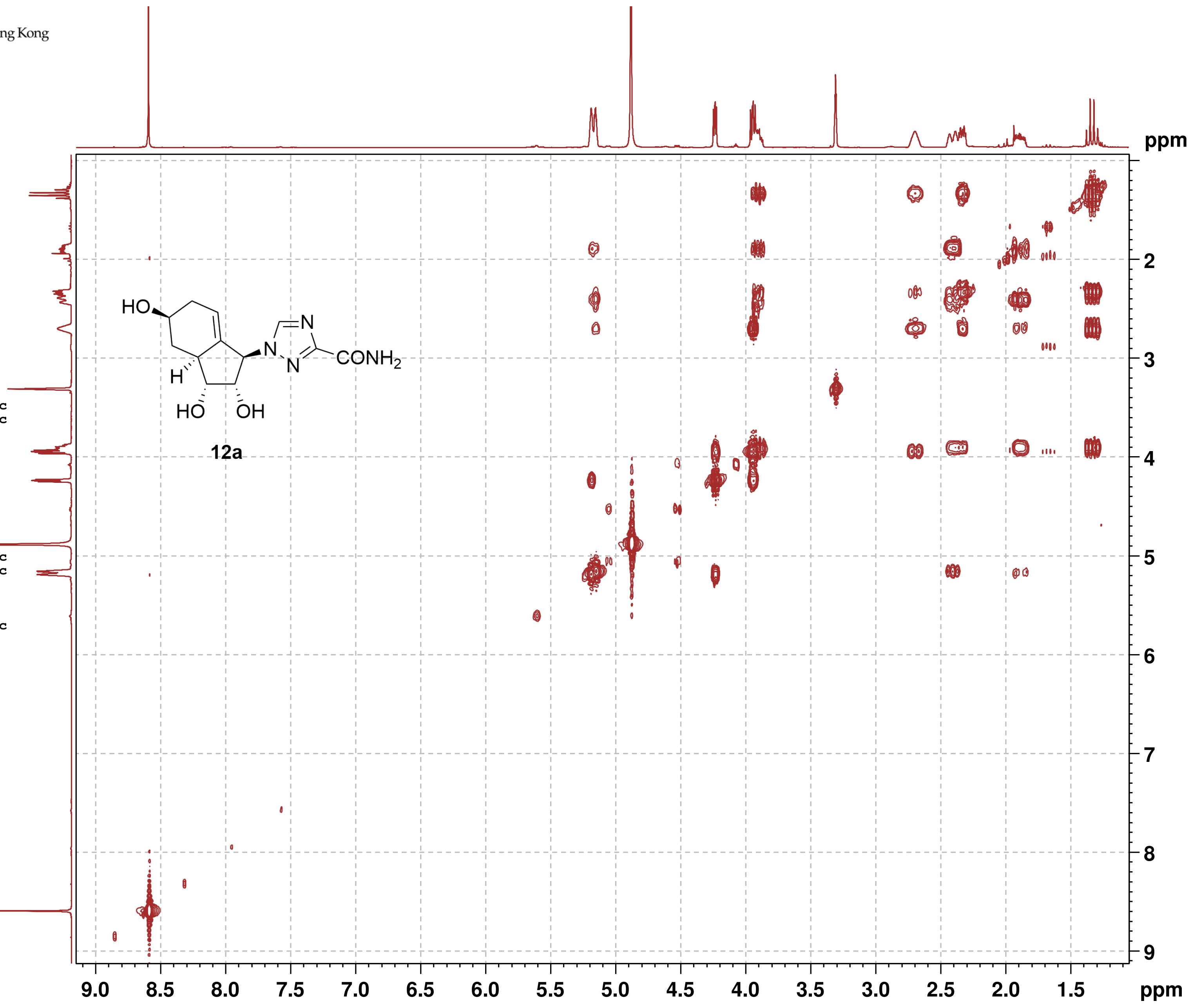

**12a**

香港中文大學
The Chinese University of Hong Kong

#### NOESY

Current Data Parameters

| | |
|---|---|
| NAME | SCC182_400 |
| EXPNO | 3 |
| PROCNO | 1 |

F2 - Acquisition Parameters

| | |
|---|---|
| Date_ | 20220630 |
| Time | 8.14 h |
| INSTRUM | spect |
| PROBHD | Z108618_0257 ( |
| PULPROG | noesygpphpp |
| TD | 2048 |
| SOLVENT | MeOD |
| NS | 2 |
| DS | 4 |
| SWH | 4000.000 Hz |
| FIDRES | 3.906250 Hz |
| AQ | 0.2560000 sec |
| RG | 203 |
| DW | 125.000 usec |
| DE | 6.50 usec |
| TE | 295.2 K |
| D0 | 0.00010590 sec |
| D1 | 2.00000000 sec |
| D8 | 0.75000000 sec |
| D11 | 0.03000000 sec |
| D12 | 0.00002000 sec |
| D16 | 0.00020000 sec |
| IN0 | 0.00025000 sec |
| TDav | 1 |
| SFO1 | 400.1318806 MHz |
| NUC1 | 1H |
| P1 | 15.00 usec |
| P2 | 30.00 usec |
| P17 | 2500.00 usec |
| PLW1 | 12.00000000 W |
| PLW10 | 3.00000000 W |
| GPNAM[1] | SMSQ10.100 |
| GPZ1 | 40.00 % |
| P16 | 1000.00 usec |

F1 - Acquisition parameters

| | |
|---|---|
| TD | 128 |
| SFO1 | 400.1319 MHz |
| FIDRES | 62.500000 Hz |
| SW | 9.997 ppm |
| FnMODE | States-TPPI |

F2 - Processing parameters

| | |
|---|---|
| SI | 1024 |
| SF | 400.1300076 MHz |
| WDW | QSINE |
| SSB | 2 |
| LB | 0 Hz |
| GB | 0 |
| PC | 1.00 |

F1 - Processing parameters

| | |
|---|---|
| SI | 1024 |
| MC2 | States-TPPI |
| SF | 400.1300067 MHz |
| WDW | QSINE |
| SSB | 2 |
| LB | 0 Hz |
| GB | 0 |

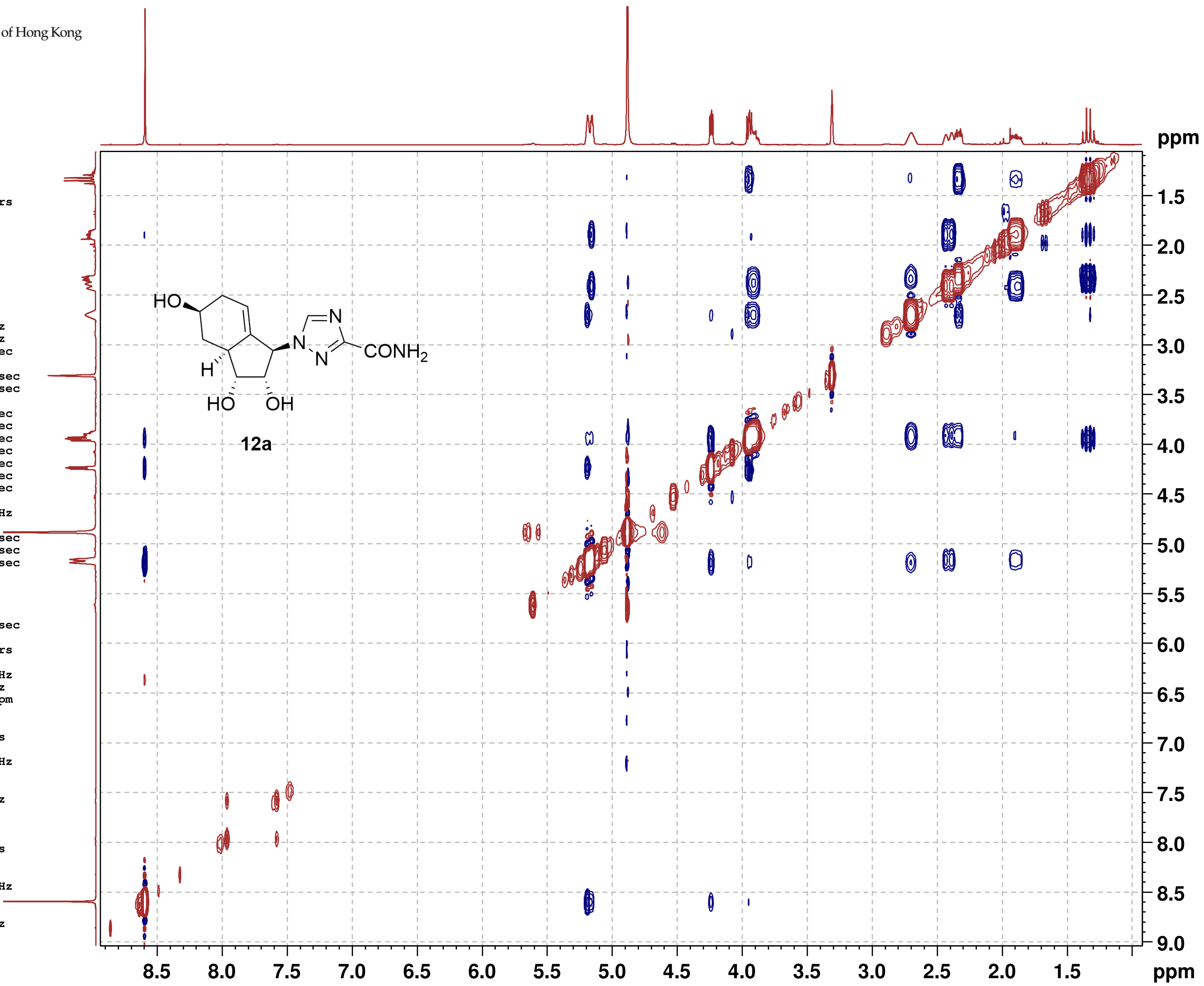

**12a**

#### NMR-Spectra for Compound 12c

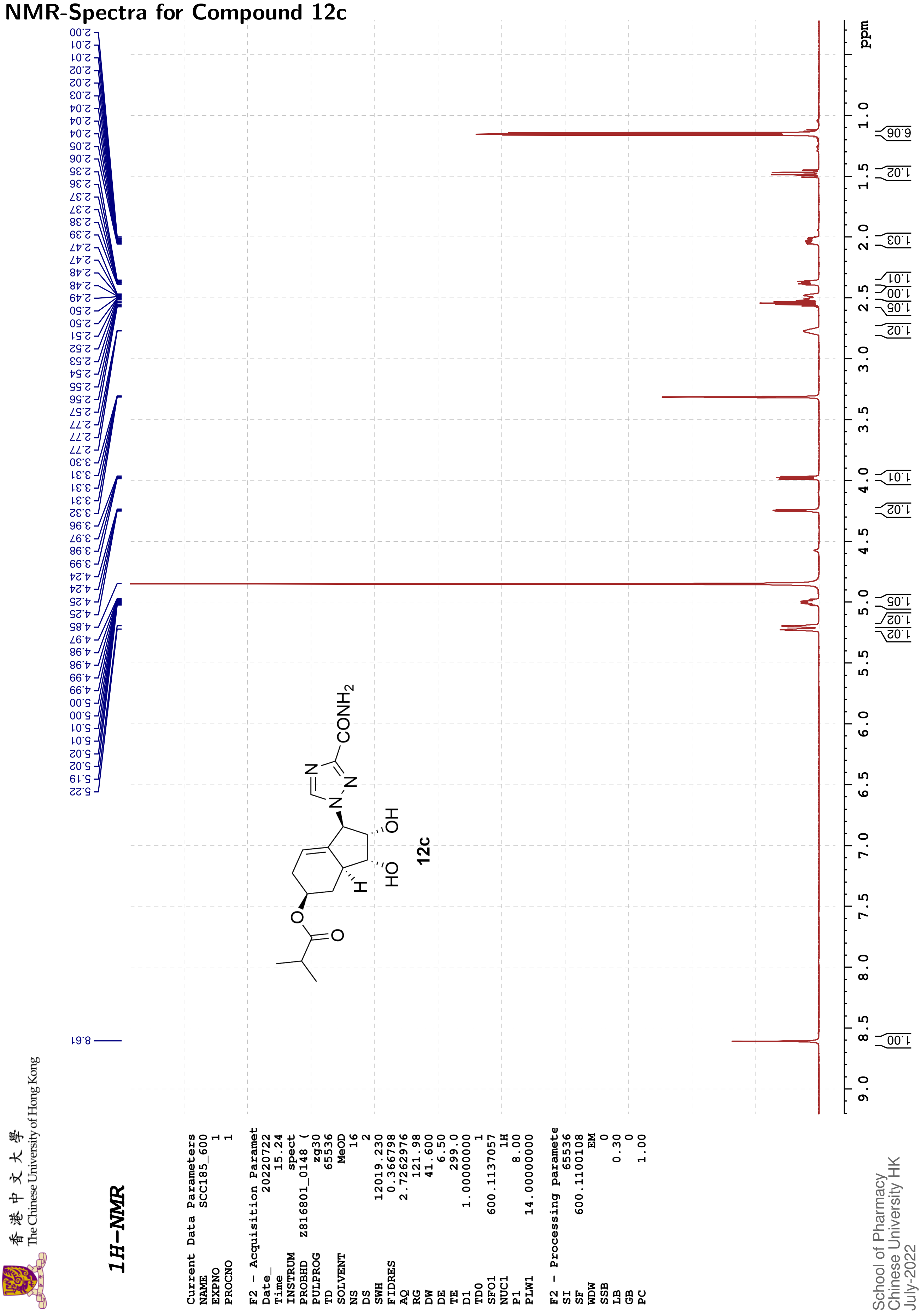

香港中文大學 The Chinese University of Hong Kong

#### 13C{1H}-NMR

| | |
|---|---|
| Current Data Parameters | |
| NAME | SCC185_600 |
| EXPNO | 4 |
| PROCNO | 1 |
| F2 - Acquisition Paramet | |
| Date_ | 20220722 |
| Time | 16.05 |
| INSTRUM | spect |
| PROBHD | Z816801_0148 ( |
| PULPROG | zgpg30 |
| TD | 65536 |
| SOLVENT | MeOD |
| NS | 308 |
| DS | 4 |
| SWH | 36231.883 |
| FIDRES | 1.105709 |
| AQ | 0.9043968 |
| RG | 60.39 |
| DW | 13.800 |
| DE | 6.50 |
| TE | 299.2 |
| D1 | 2.00000000 |
| D11 | 0.03000000 |
| TD0 | 1 |
| SFO1 | 150.9128693 |
| NUC1 | 13C |
| P1 | 12.00 |
| PLW1 | 183.00000000 |
| SFO2 | 600.1124004 |
| NUC2 | 1H |
| CPDPRG[2 | waltz16 |
| PCPD2 | 70.00 |
| PLW2 | 14.00000000 |
| PLW12 | 0.18286000 |
| PLW13 | 0.09197600 |
| F2 - Processing paramete | |
| SI | 32768 |
| SF | 150.8975689 |
| WDW | EM |
| SSB | 0 |
| LB | 3.00 |
| GB | 0 |
| PC | 1.40 |

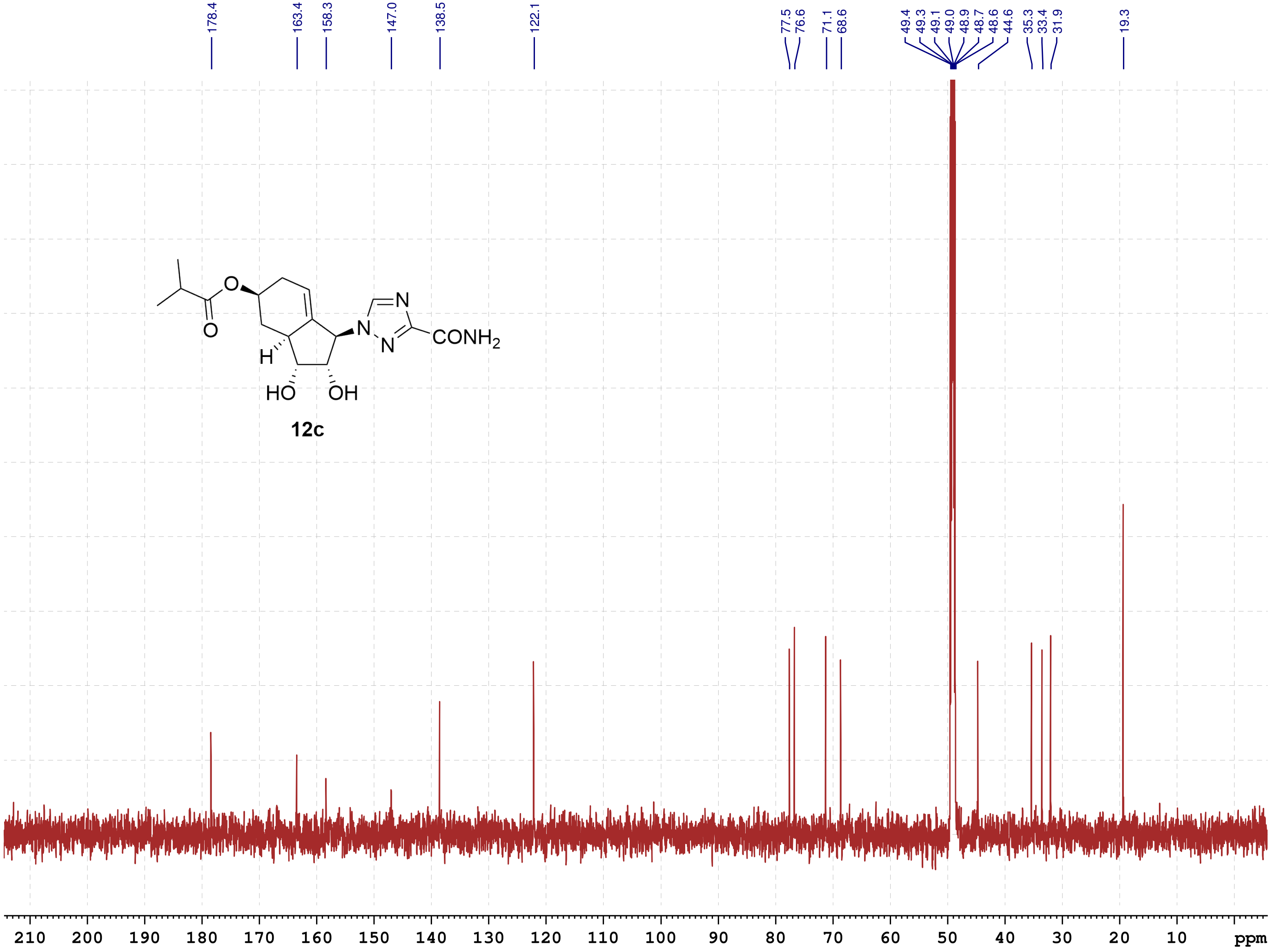

12c

School of Pharmacy
Chinese University HK
July-2022

香港中文大學
The Chinese University of Hong Kong

COSY

```
Current Data Parameters
NAME          SCC185_600
EXPNO                  2
PROCNO                 1

F2 - Acquisition Parameters
Date_           20220722
Time               15.25 h
INSTRUM            spect
PROBHD    Z816801_0148 (
PULPROG       cosygpppqf
TD                  2048
SOLVENT             MeOD
NS                     2
DS                     8
SWH             7812.500 Hz
FIDRES          7.629395 Hz
AQ             0.1310720 sec
RG                189.92
DW                64.000 usec
DE                  6.50 usec
TE                 299.0 K
D0            0.00000300 sec
D1            2.00000000 sec
D11           0.03000000 sec
D12           0.00002000 sec
D13           0.00000400 sec
D16           0.00020000 sec
IN0           0.00012800 sec
TDav                   1
SFO1         600.1136007 MHz
NUC1                  1H
P0                  8.00 usec
P1                  8.00 usec
P17              2500.00 usec
PLW1         14.00000000 W
PLW10         1.43359995 W
GPNAM[1]        SMSQ10.100
GPZ1               10.00 %
P16              1000.00 usec

F1 - Acquisition parameters
TD                   128
SFO1            600.1136 MHz
FIDRES        122.070312 Hz
SW                13.018 ppm
FnMODE                QF

F2 - Processing parameters
SI                  1024
SF           600.1100000 MHz
WDW                QSINE
SSB                    0
LB                     0 Hz
GB                     0
PC                  1.40

F1 - Processing parameters
SI                  1024
MC2                   QF
SF           600.1100000 MHz
WDW                QSINE
SSB                    0
LB                     0 Hz
GB                     0
```

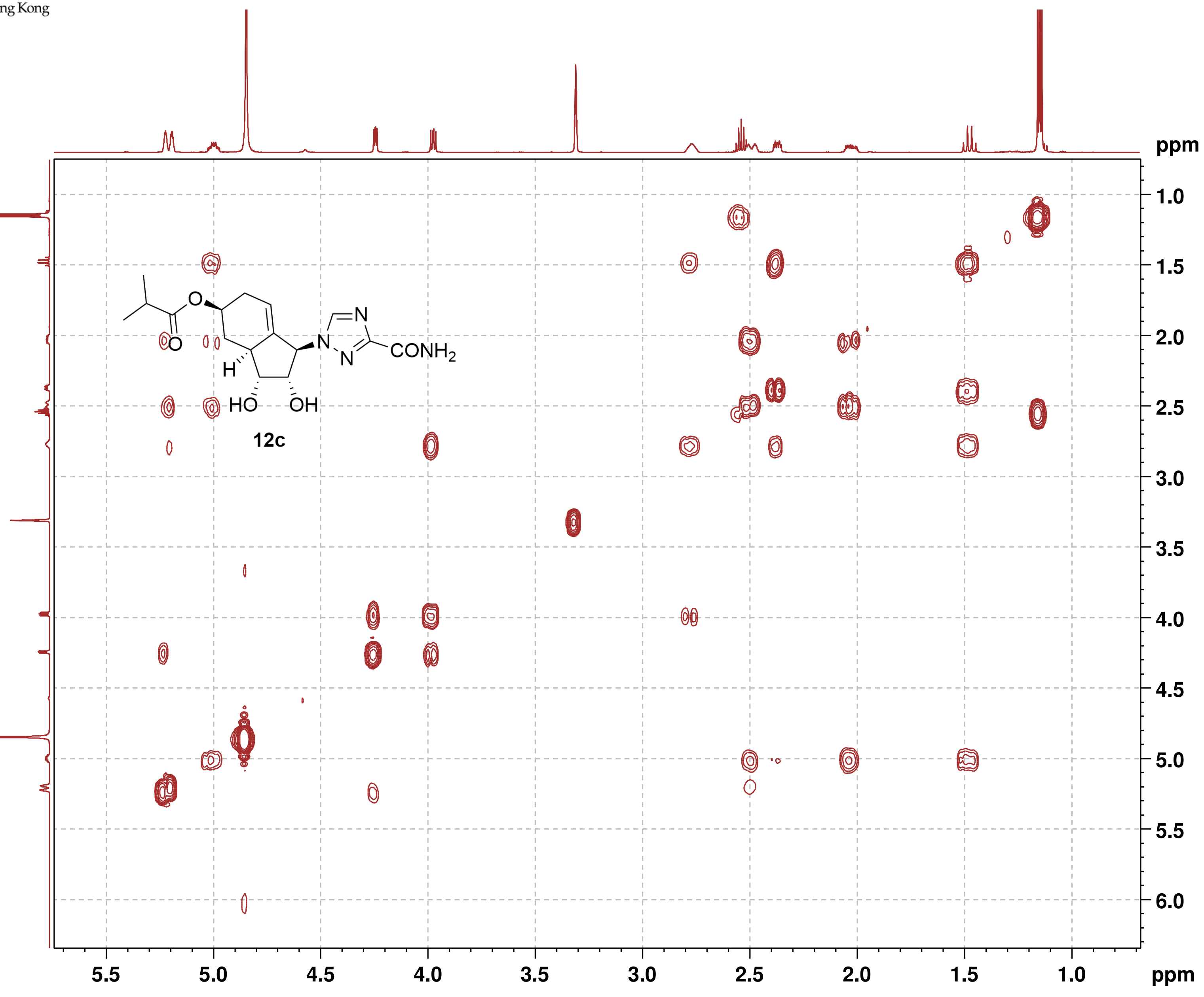

12c

香 港 中 文 大 學
The Chinese University of Hong Kong

**NOESY**

```
Current Data Parameters
NAME          SCC185_600
EXPNO                  3
PROCNO                 1

F2 - Acquisition Parameters
Date_           20220722
Time               15.36 h
INSTRUM            spect
PROBHD     Z816801_0148 (
PULPROG      noesygpphpp
TD                  2048
SOLVENT             MeOD
NS                     2
DS                    16
SWH             6009.615 Hz
FIDRES          5.868765 Hz
AQ             0.1703936 sec
RG                189.92
DW                83.200 usec
DE                  6.50 usec
TE                 299.0 K
D0            0.00007301 sec
D1            2.00000000 sec
D8            0.75000000 sec
D11           0.03000000 sec
D12           0.00002000 sec
D16           0.00020000 sec
IN0           0.00016640 sec
TDav                   1
SFO1         600.1128205 MHz
NUC1                  1H
P1                  8.00 usec
P2                 16.00 usec
P17              2500.00 usec
PLW1         14.00000000 W
PLW10         1.43359995 W
GPNAM[1]       SMSQ10.100
GPZ1               40.00 %
P16              1000.00 usec

F1 - Acquisition parameters
TD                   128
SFO1            600.1128 MHz
FIDRES         93.900238 Hz
SW                10.014 ppm
FnMODE       States-TPPI

F2 - Processing parameters
SI                  1024
SF           600.1100000 MHz
WDW                QSINE
SSB                    2
LB                     0 Hz
GB                     0
PC                  1.00

F1 - Processing parameters
SI                  1024
MC2          States-TPPI
SF           600.1100000 MHz
WDW                QSINE
SSB                    2
LB                     0 Hz
GB                     0
```

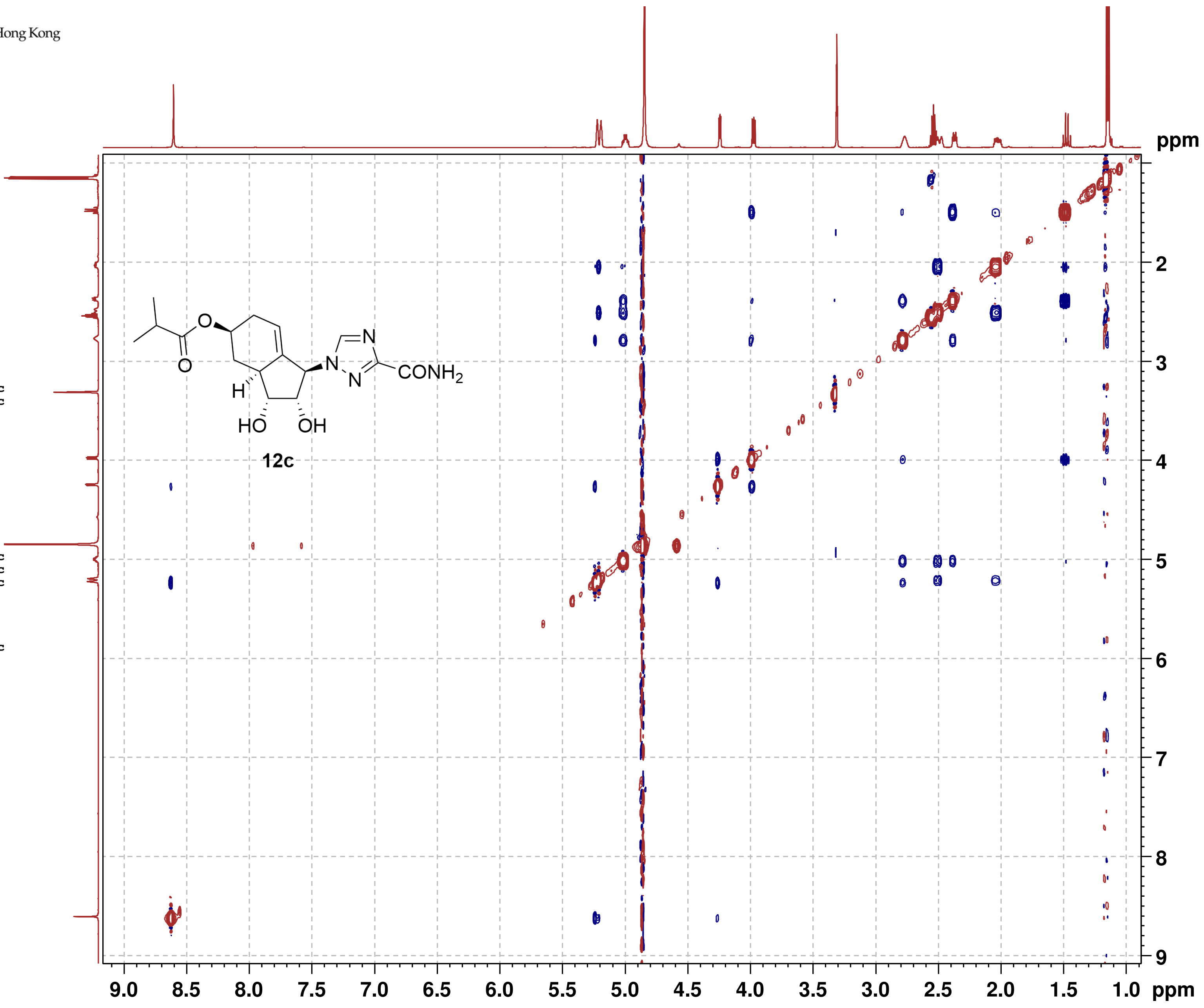

#### NMR-Spectra for Compound 5b

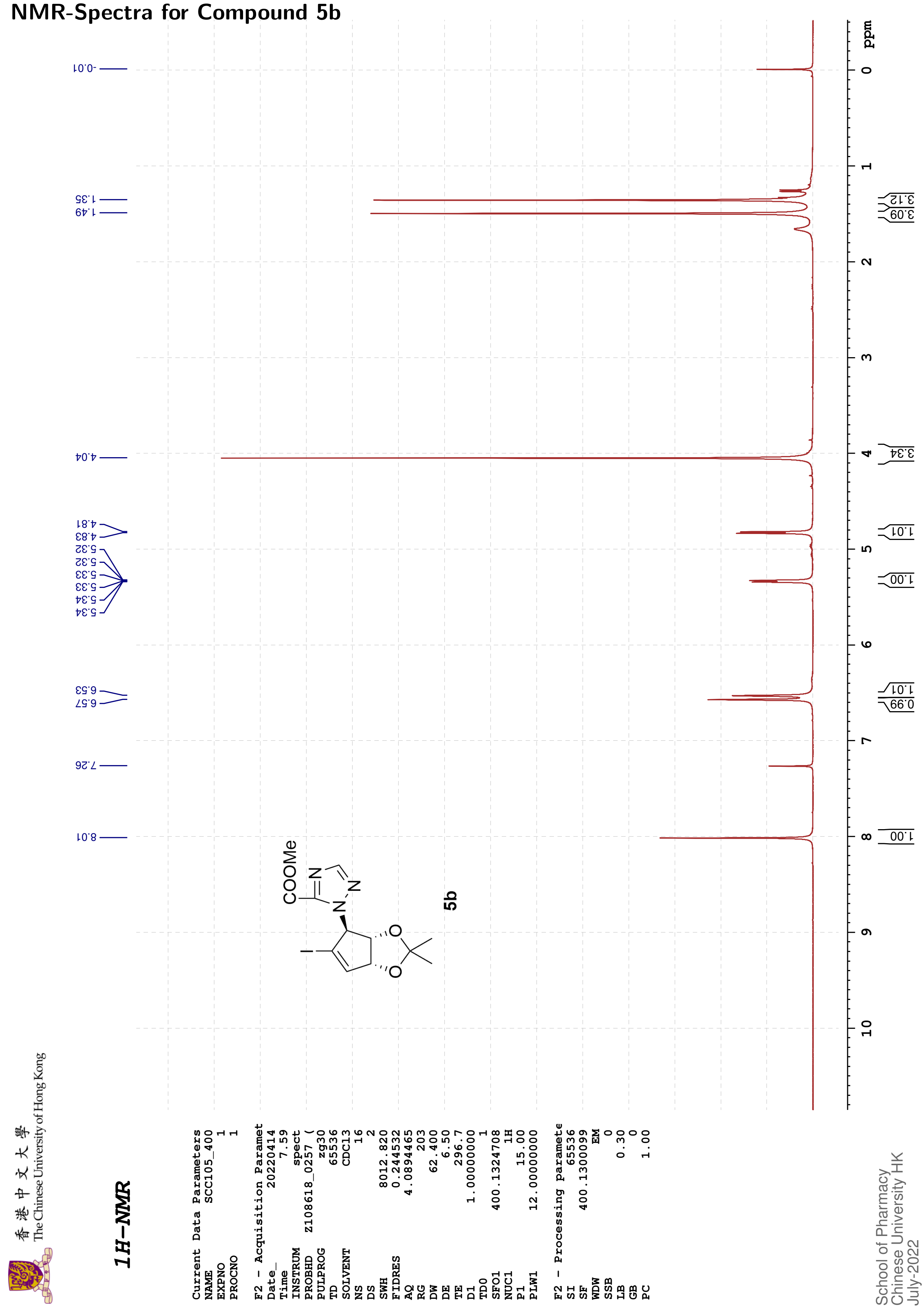

香港中文大學 The Chinese University of Hong Kong

#### 13C{1H}-NMR

```
Current Data Parameters
NAME              SCC105_400
EXPNO                      2
PROCNO                     1

F2 - Acquisition Paramet
Date_               20220414
Time                    8.19
INSTRUM                spect
PROBHD       Z108618_0257 (
PULPROG              zgpg30
TD                     48074
SOLVENT                CDCl3
NS                       353
DS                         4
SWH                24038.461
FIDRES              1.000061
AQ                 0.9999392
RG                       203
DW                    20.800
DE                      6.50
TE                     297.3
D1                2.00000000
D11               0.03000000
TD0                        1
SFO1             100.6228298
NUC1                     13C
P1                     10.00
PLW1             47.00000000
SFO2             400.1316005
NUC2                      1H
CPDPRG[2            waltz16
PCPD2                  90.00
PLW2             12.00000000
PLW12             0.33333001
PLW13             0.16766000

F2 - Processing paramete
SI                     32768
SF               100.6127571
WDW                       EM
SSB                        0
LB                      2.00
GB                         0
PC                      1.40
```

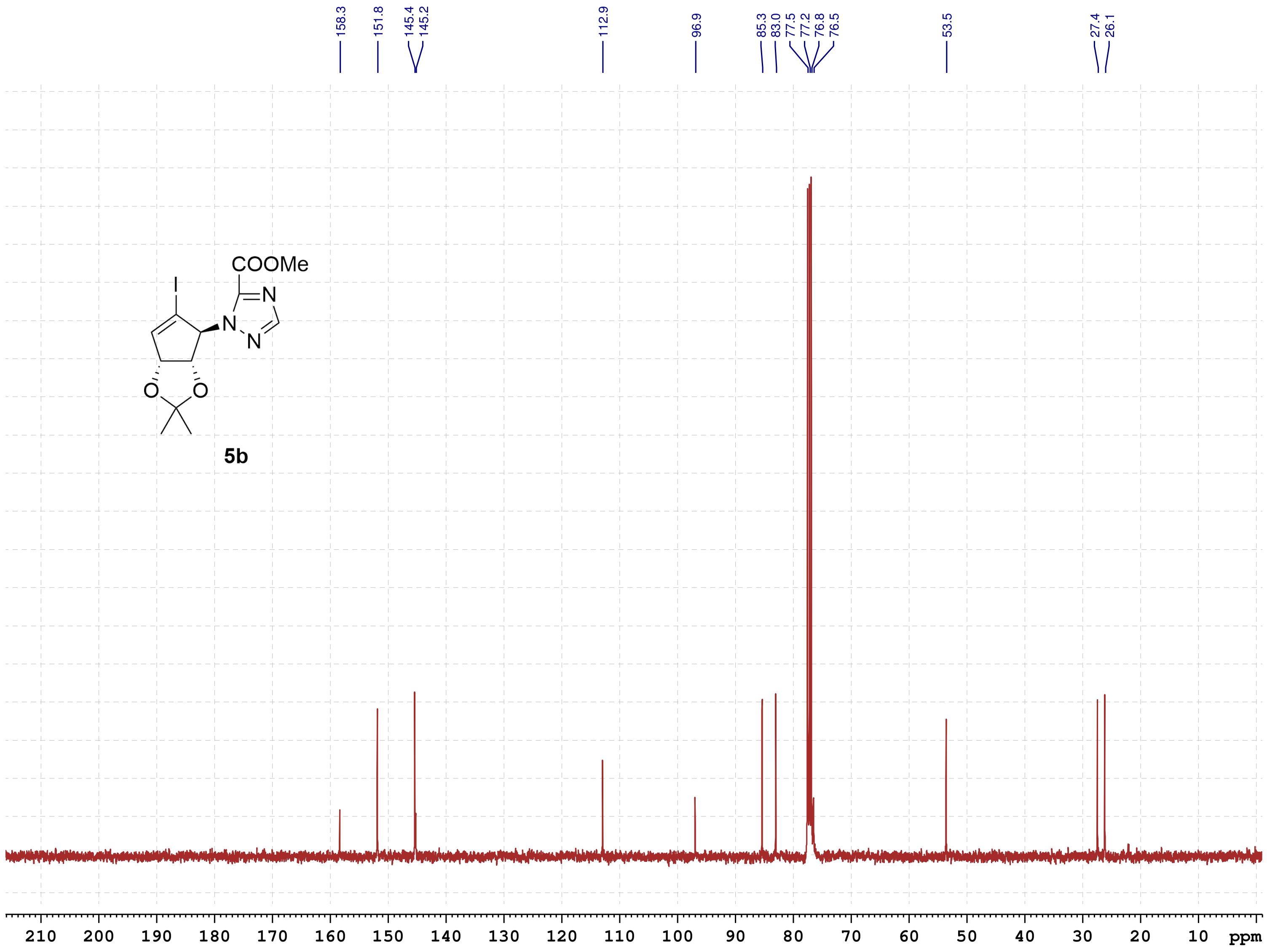

School of Pharmacy
Chinese University HK
July-2022

#### NMR-Spectra for Compound 6b

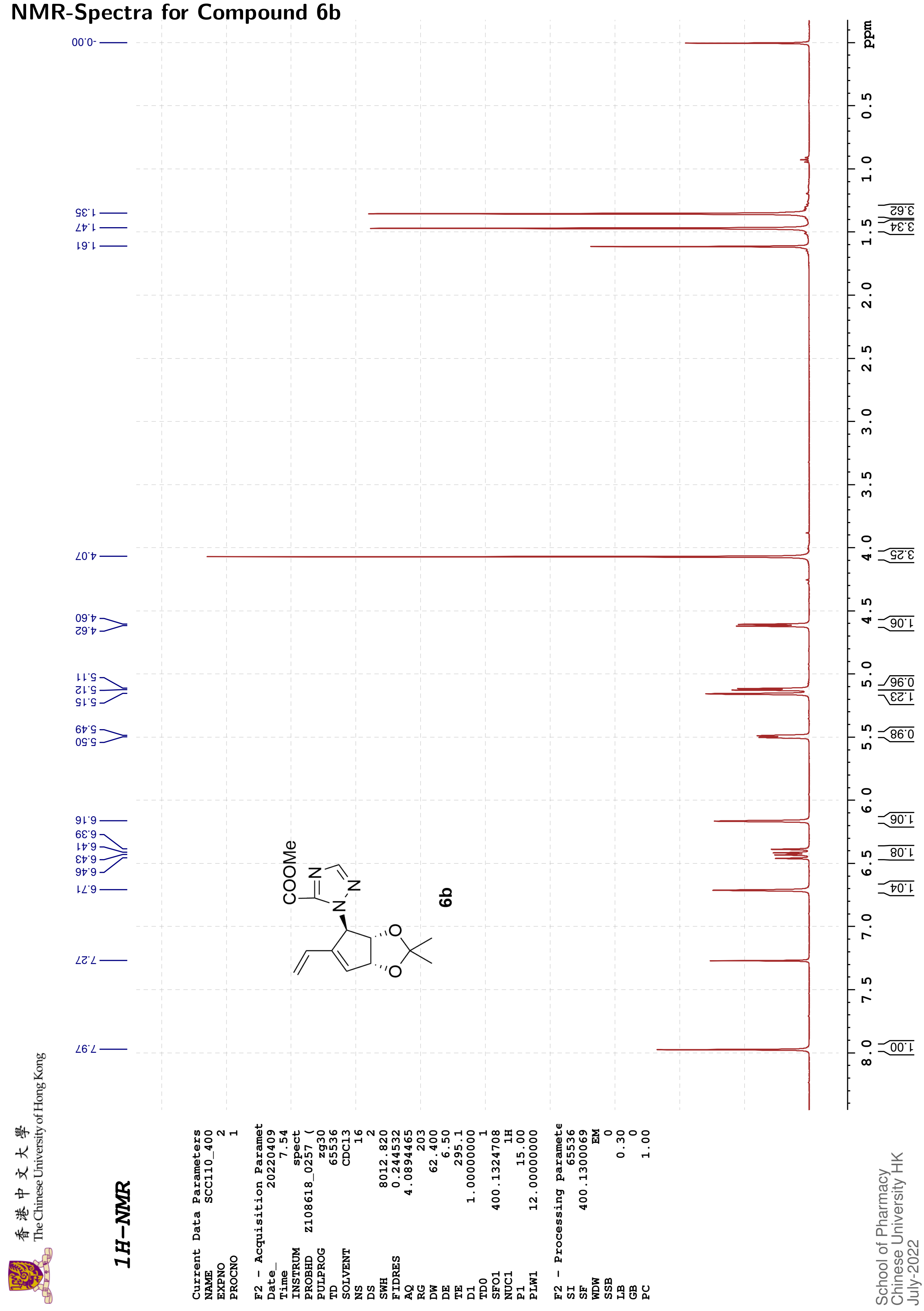

香港中文大學 The Chinese University of Hong Kong

#### 13C{1H}-NMR

| | |
|---|---|
| Current Data Parameters | |
| NAME | SCC110_400 |
| EXPNO | 3 |
| PROCNO | 1 |
| F2 - Acquisition Paramet | |
| Date_ | 20220409 |
| Time | 8.22 |
| INSTRUM | spect |
| PROBHD | Z108618_0257 ( |
| PULPROG | zgpg30 |
| TD | 48074 |
| SOLVENT | CDCl3 |
| NS | 506 |
| DS | 4 |
| SWH | 24038.461 |
| FIDRES | 1.000061 |
| AQ | 0.9999392 |
| RG | 203 |
| DW | 20.800 |
| DE | 6.50 |
| TE | 295.7 |
| D1 | 2.00000000 |
| D11 | 0.03000000 |
| TD0 | 1 |
| SFO1 | 100.6228298 |
| NUC1 | 13C |
| P1 | 10.00 |
| PLW1 | 47.00000000 |
| SFO2 | 400.1316005 |
| NUC2 | 1H |
| CPDPRG[2 | waltz16 |
| PCPD2 | 90.00 |
| PLW2 | 12.00000000 |
| PLW12 | 0.33333001 |
| PLW13 | 0.16766000 |
| F2 - Processing paramete | |
| SI | 32768 |
| SF | 100.6127558 |
| WDW | EM |
| SSB | 0 |
| LB | 2.00 |
| GB | 0 |
| PC | 1.40 |

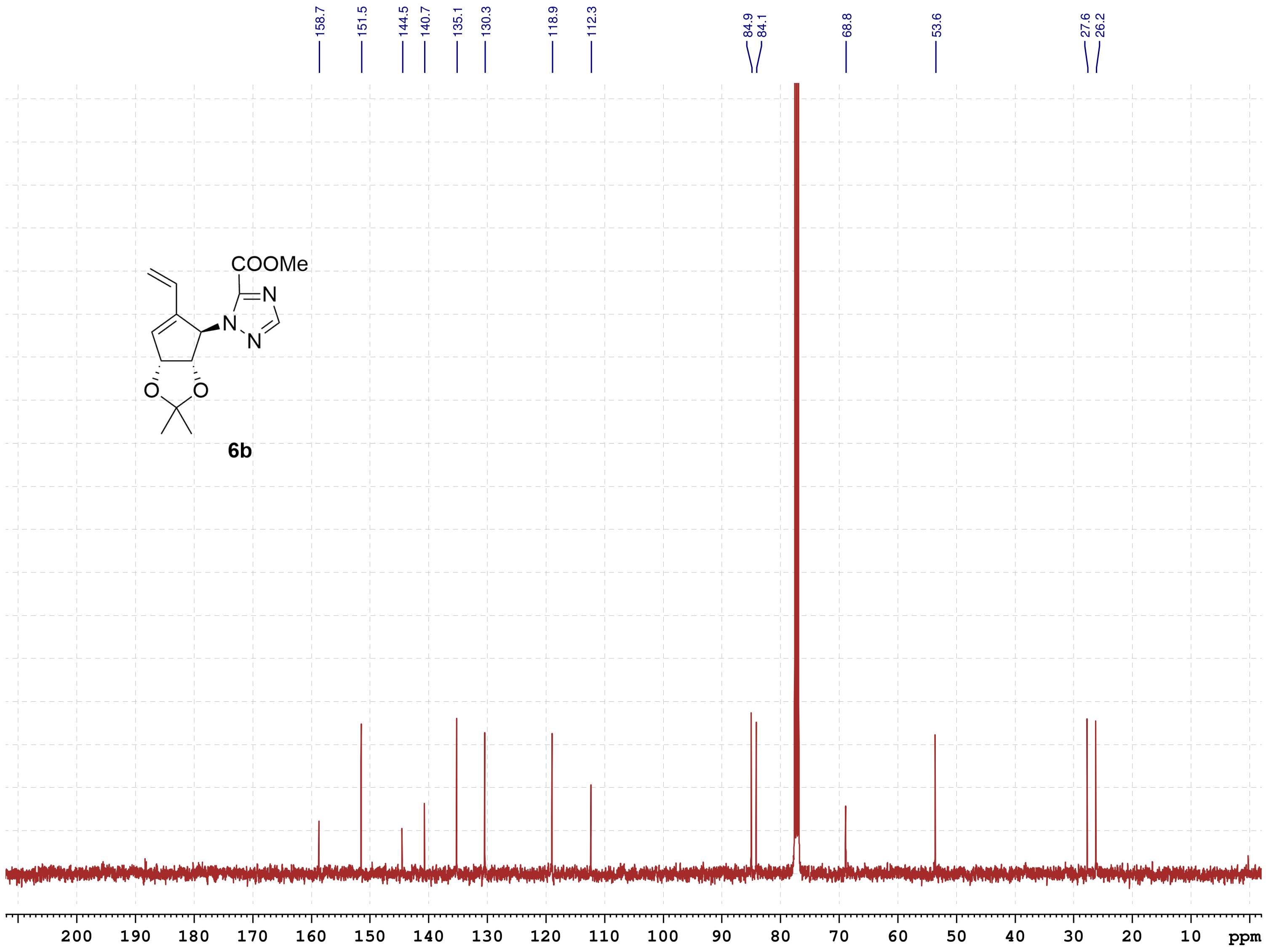

School of Pharmacy
Chinese University HK
July-2022

#### NMR-Spectra for Compound 7b

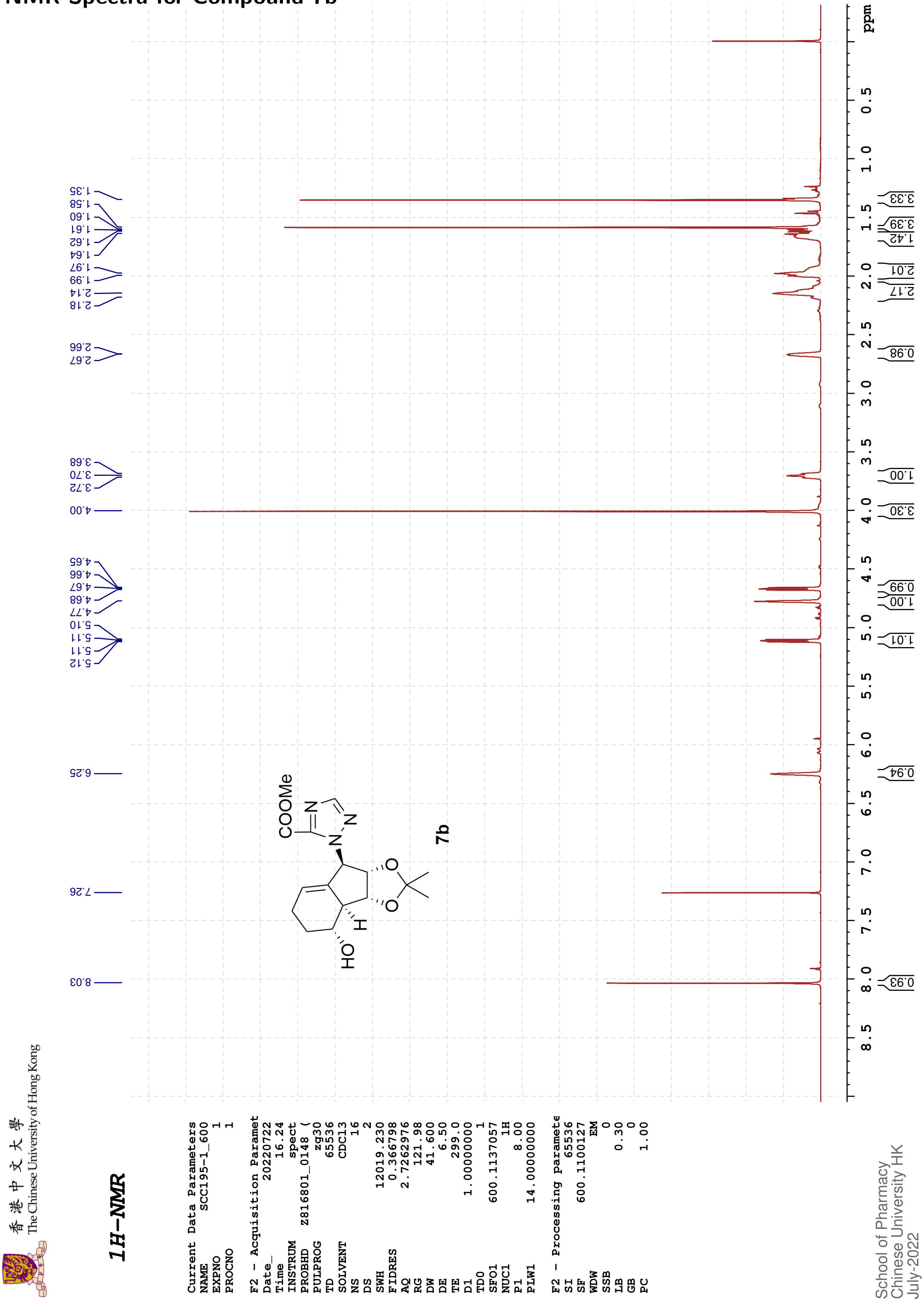

The Chinese University of Hong Kong

香港中文大學

#### 13C{1H}-NMR

```
Current Data Parameters
NAME        SCC195-1_600
EXPNO                  2
PROCNO                 1

F2 - Acquisition Paramet
Date_           20220722
Time               16.53
INSTRUM            spect
PROBHD    Z816801_0148 (
PULPROG          zgpg30
TD                 65536
SOLVENT            CDCl3
NS                   365
DS                     4
SWH            36231.883
FIDRES          1.105709
AQ             0.9043968
RG                 60.39
DW                13.800
DE                  6.50
TE                 299.3
D1            2.00000000
D11           0.03000000
TD0                    1
SFO1         150.9128693
NUC1                 13C
P1                 12.00
PLW1        183.00000000
SFO2         600.1124004
NUC2                  1H
CPDPRG[2         waltz16
PCPD2              70.00
PLW2         14.00000000
PLW12         0.18286000
PLW13         0.09197600

F2 - Processing paramete
SI                 32768
SF           150.8977593
WDW                   EM
SSB                    0
LB                  2.00
GB                     0
PC                  1.40
```

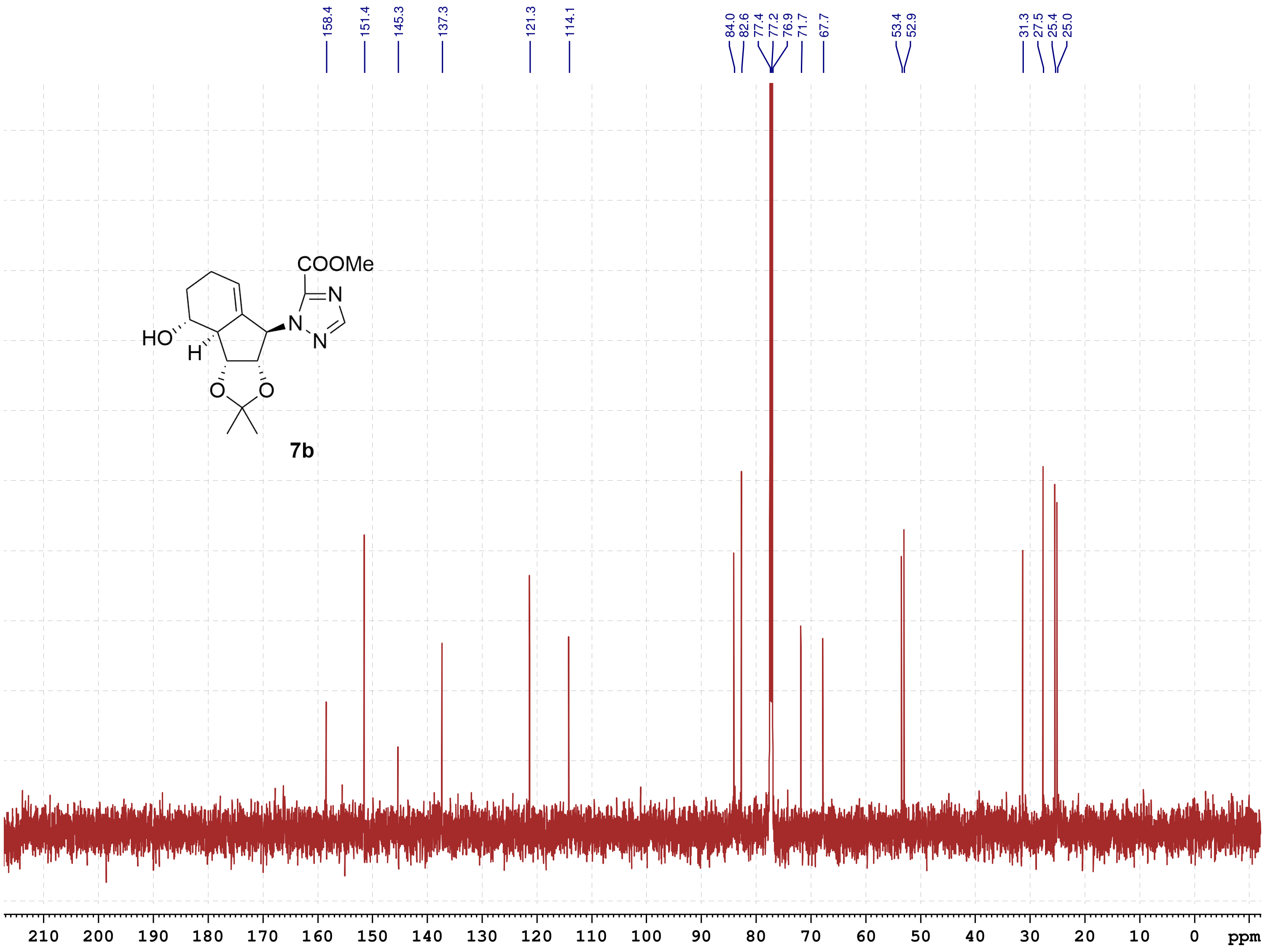

**7b**

School of Pharmacy
Chinese University HK
July-2022

#### NMR-Spectra for Compound 10b

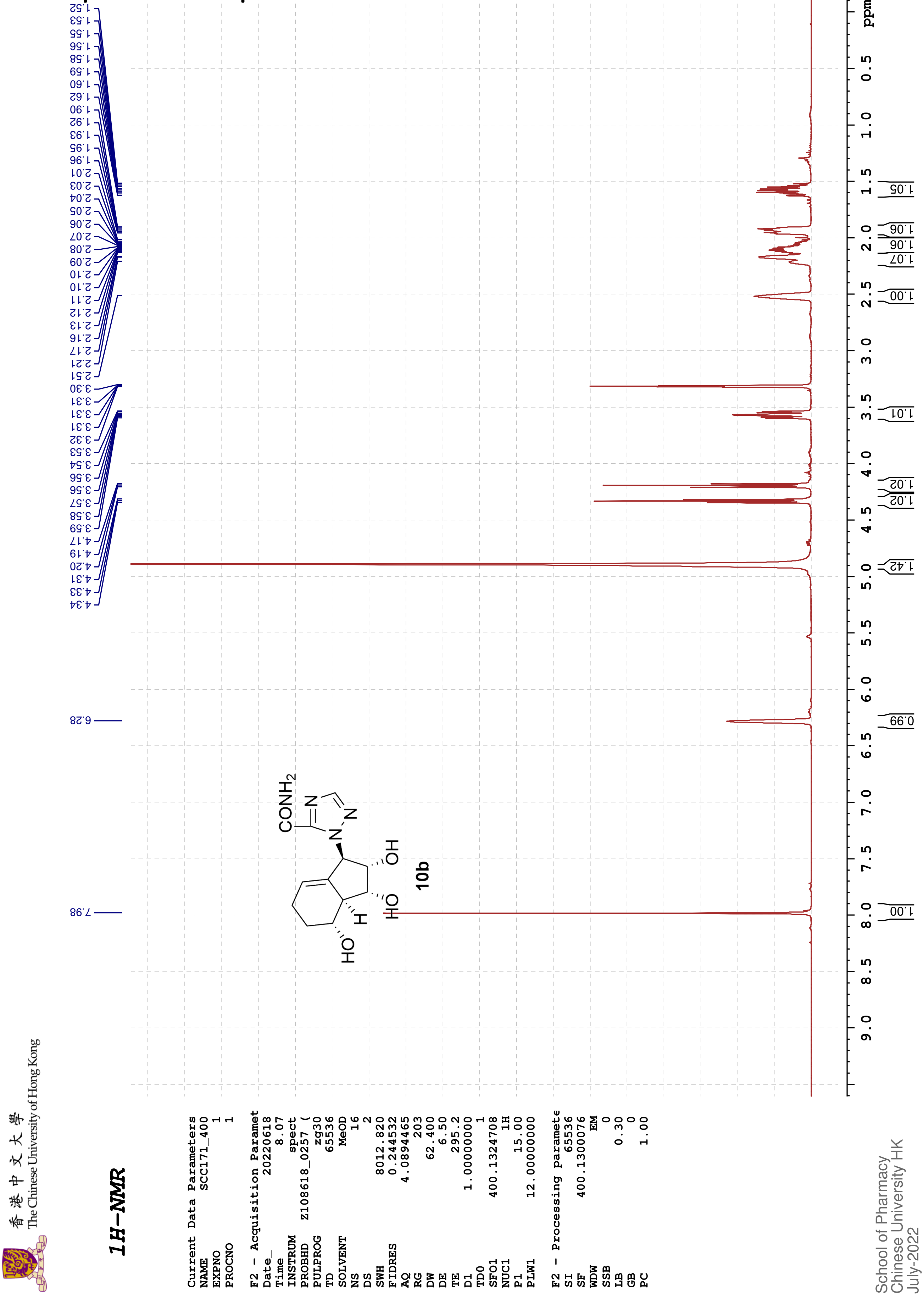

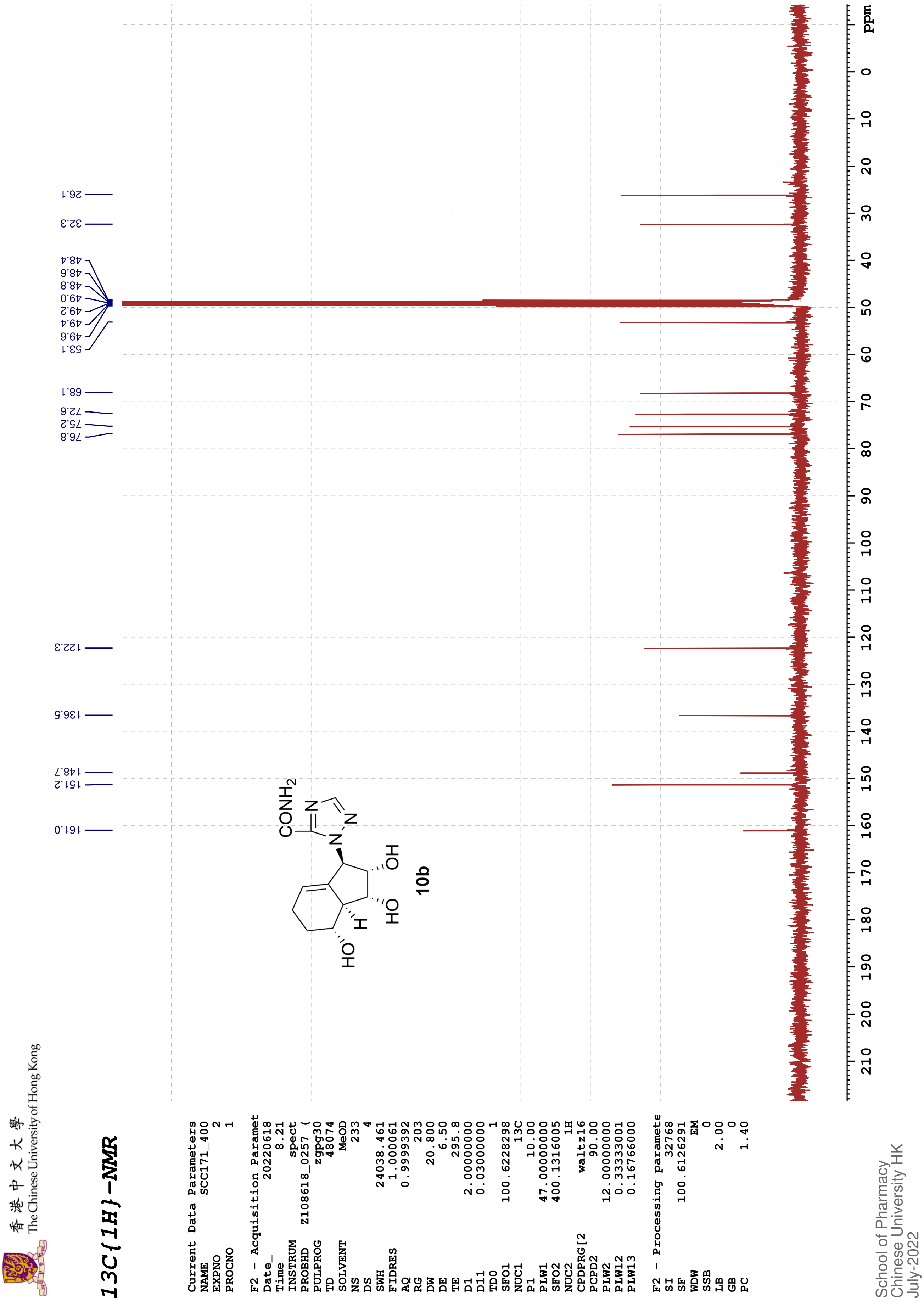
The Chinese University of Hong Kong
13C{1H}-NMR
Current Data Parameters
NAME SCC171_400
EXPNO 2
PROCNO 1
F2 - Acquisition Paramet
Date_ 20220618
Time 8.21
INSTRUM spect
PROBHD Z108618_0257 (
PULPROG zgpg30
TD 48074
SOLVENT MeOD
NS 233
DS 4
SWH 24038.461
FIDRES 1.000061
AQ 0.9999392
RG 203
DW 20.800
DE 6.50
TE 295.8
D1 2.00000000
D11 0.03000000
TD0 1
SFO1 100.6228298
NUC1 13C
P1 10.00
PLW1 47.00000000
SFO2 400.1316005
NUC2 1H
CPDPRG[2 waltz16
PCPD2 90.00
PLW2 12.00000000
PLW12 0.33333001
PLW13 0.16766000
F2 - Processing paramete
SI 32768
SF 100.6126291
WDW EM
SSB 0
LB 2.00
GB 0
PC 1.40
161.0
151.2
148.7
136.5
122.3
76.8
75.2
72.6
68.1
53.1
49.6
49.4
49.2
49.0
48.8
48.6
48.4
32.3
26.1
CONH2
N
N
N
OH
HO
HO
H
10b
ppm
School of Pharmacy
Chinese University HK
July-2022

COSY

```
Current Data Parameters
NAME          SCC171_400
EXPNO                  3
PROCNO                 1

F2 - Acquisition Parameters
Date_           20220618
Time                8.23 h
INSTRUM            spect
PROBHD   Z108618_0257 (
PULPROG         cosygpqf
TD                  2048
SOLVENT             MeOD
NS                     2
DS                     8
SWH             4000.000 Hz
FIDRES          3.906250 Hz
AQ             0.2560000 sec
RG                   203
DW               125.000 usec
DE                  6.50 usec
TE                 295.4 K
D0            0.00000300 sec
D1            1.48689198 sec
D13           0.00000400 sec
D16           0.00020000 sec
IN0           0.00025000 sec
TDav                   1
SFO1         400.1320006 MHz
NUC1                  1H
P0                 15.00 usec
P1                 15.00 usec
PLW1         12.00000000 W
GPNAM[1]        SINE.100
GPZ1               10.00 %
P16              1000.00 usec

F1 - Acquisition parameters
TD                   128
SFO1             400.132 MHz
FIDRES         62.500000 Hz
SW                 9.997 ppm
FnMODE                QF

F2 - Processing parameters
SI                  1024
SF           400.1300081 MHz
WDW                 SINE
SSB                    0
LB                     0 Hz
GB                     0
PC                  1.40

F1 - Processing parameters
SI                  1024
MC2                   QF
SF           400.1300079 MHz
WDW                 SINE
SSB                    0
LB                     0 Hz
GB                     0
```

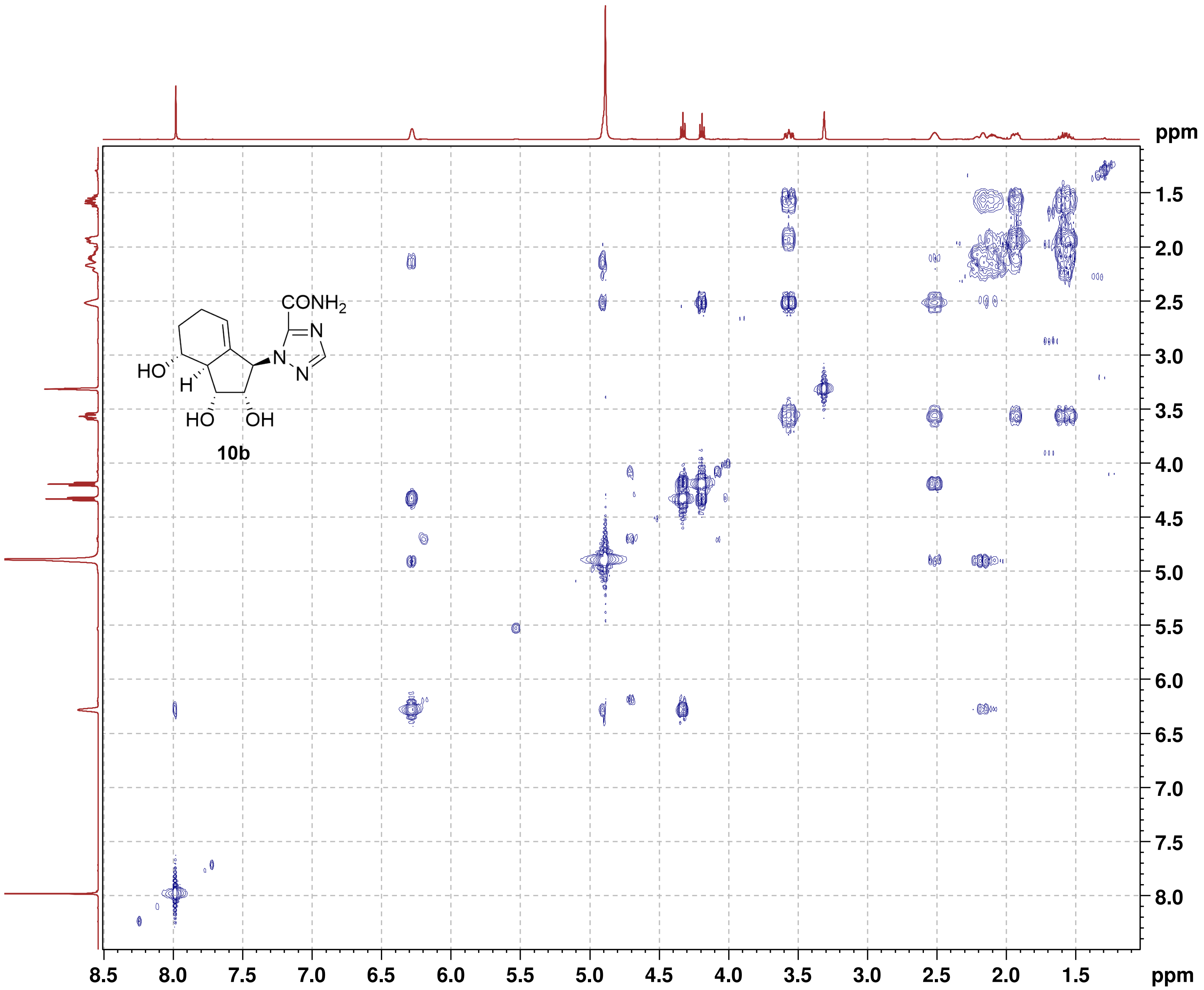

**NOESY**

```
Current Data Parameters
NAME          SCC171_400
EXPNO                  4
PROCNO                 1

F2 - Acquisition Parameters
Date_           20220618
Time                8.33 h
INSTRUM            spect
PROBHD   Z108618_0257 (
PULPROG      noesygpphpp
TD                  2048
SOLVENT             MeOD
NS                     2
DS                    16
SWH             4000.000 Hz
FIDRES          3.906250 Hz
AQ             0.2560000 sec
RG                   203
DW               125.000 usec
DE                  6.50 usec
TE                 295.2 K
D0            0.00010590 sec
D1            2.00000000 sec
D8            0.80000001 sec
D11           0.03000000 sec
D12           0.00002000 sec
D16           0.00020000 sec
IN0           0.00025000 sec
TDav                   1
SFO1         400.1318806 MHz
NUC1                  1H
P1                 15.00 usec
P2                 30.00 usec
P17              2500.00 usec
PLW1         12.00000000 W
PLW10         3.00000000 W
GPNAM[1]      SMSQ10.100
GPZ1               40.00 %
P16              1000.00 usec

F1 - Acquisition parameters
TD                   128
SFO1            400.1319 MHz
FIDRES         62.500000 Hz
SW                 9.997 ppm
FnMODE       States-TPPI

F2 - Processing parameters
SI                  1024
SF           400.1300088 MHz
WDW                QSINE
SSB                    2
LB                     0 Hz
GB                     0
PC                  1.00

F1 - Processing parameters
SI                  1024
MC2          States-TPPI
SF           400.1300079 MHz
WDW                QSINE
SSB                    2
LB                     0 Hz
GB                     0
```

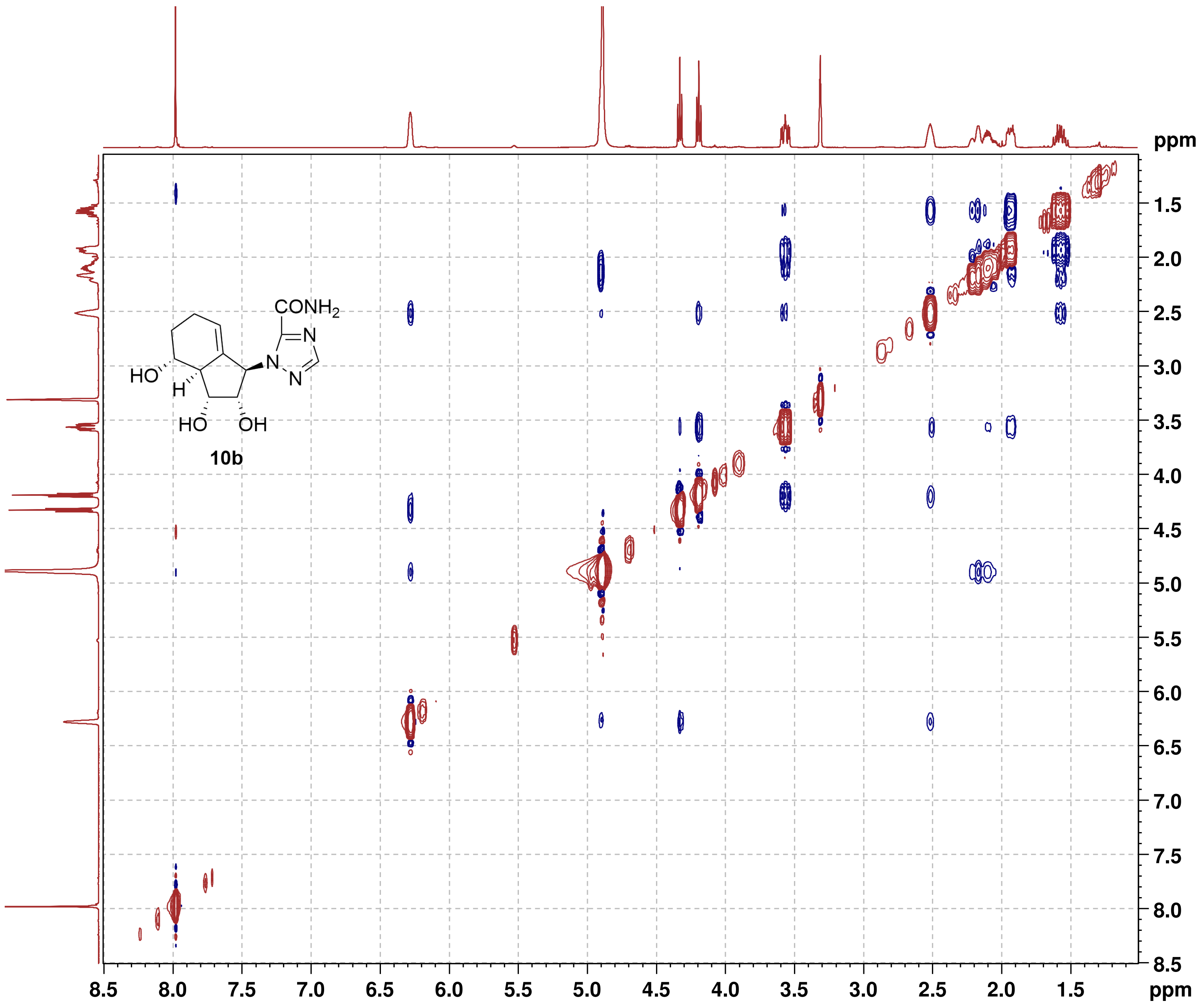

#### NMR-Spectra for Compound 10d

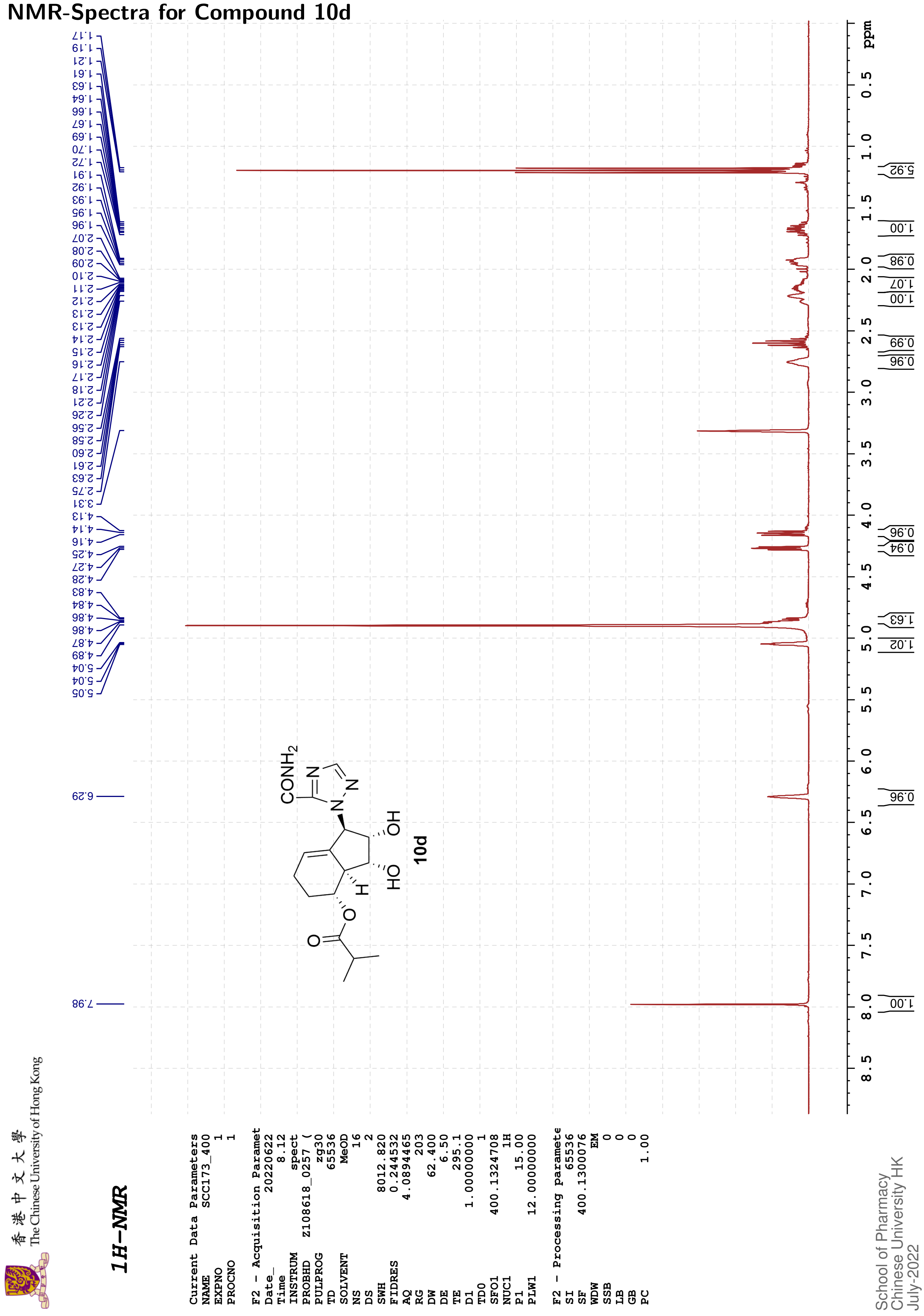

The Chinese University of Hong Kong 香港中文大學

#### 13C{1H}-NMR

| | |
|---|---|
| Current Data Parameters | |
| NAME | SCC173_400 |
| EXPNO | 2 |
| PROCNO | 1 |
| F2 - Acquisition Paramet | |
| Date_ | 20220622 |
| Time | 8.26 |
| INSTRUM | spect |
| PROBHD | Z108618_0257 ( |
| PULPROG | zgpg30 |
| TD | 48074 |
| SOLVENT | MeOD |
| NS | 240 |
| DS | 4 |
| SWH | 24038.461 |
| FIDRES | 1.000061 |
| AQ | 0.9999392 |
| RG | 203 |
| DW | 20.800 |
| DE | 6.50 |
| TE | 295.4 |
| D1 | 2.00000000 |
| D11 | 0.03000000 |
| TD0 | 1 |
| SFO1 | 100.6228298 |
| NUC1 | 13C |
| P1 | 10.00 |
| PLW1 | 47.00000000 |
| SFO2 | 400.1316005 |
| NUC2 | 1H |
| CPDPRG[2 | waltz16 |
| PCPD2 | 90.00 |
| PLW2 | 12.00000000 |
| PLW12 | 0.33333001 |
| PLW13 | 0.16766000 |
| F2 - Processing paramete | |
| SI | 32768 |
| SF | 100.6126292 |
| WDW | EM |
| SSB | 0 |
| LB | 1.00 |
| GB | 0 |
| PC | 1.40 |

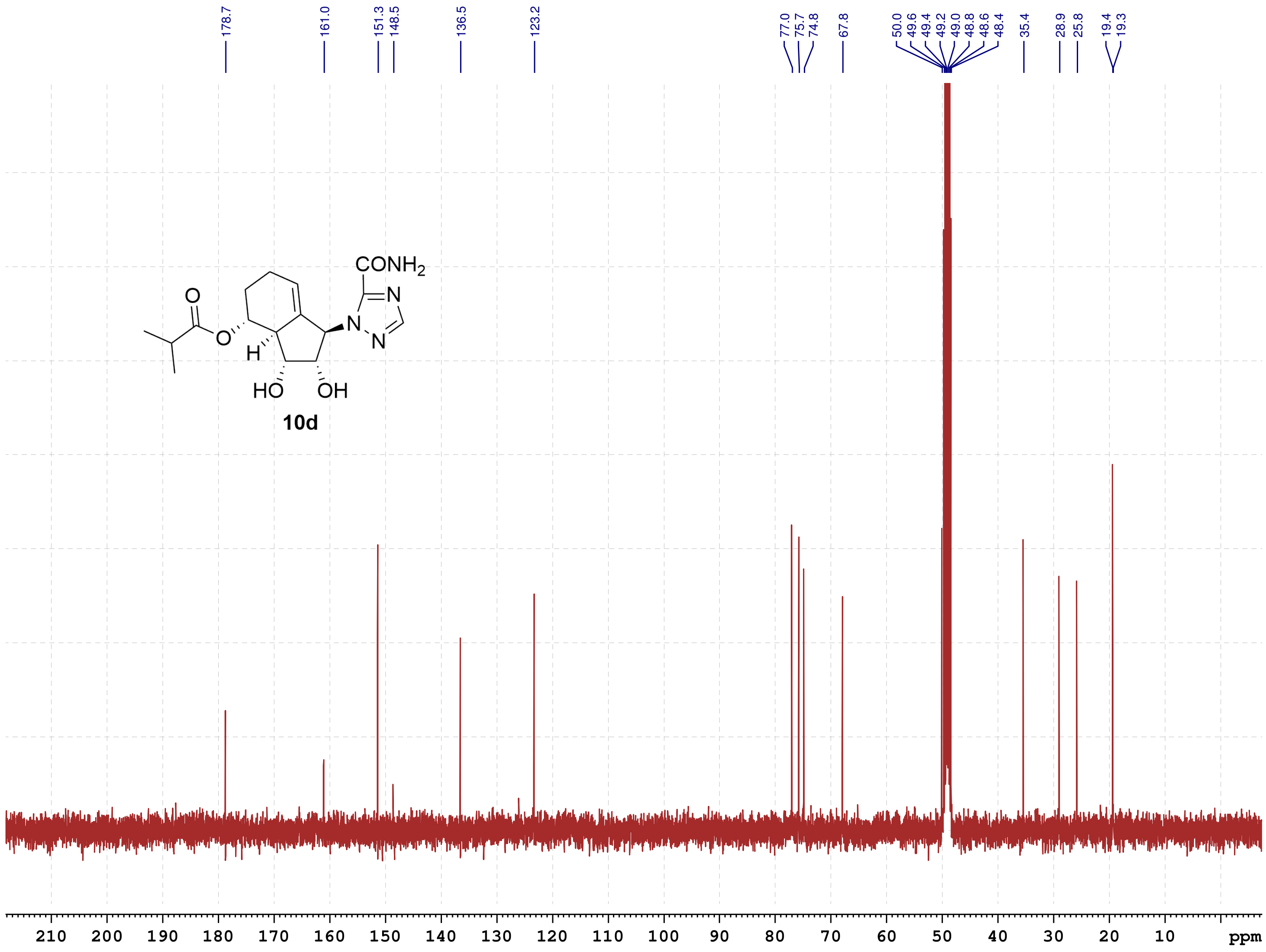

School of Pharmacy
Chinese University HK
July-2022

#### NMR-Spectra for Compound 8b

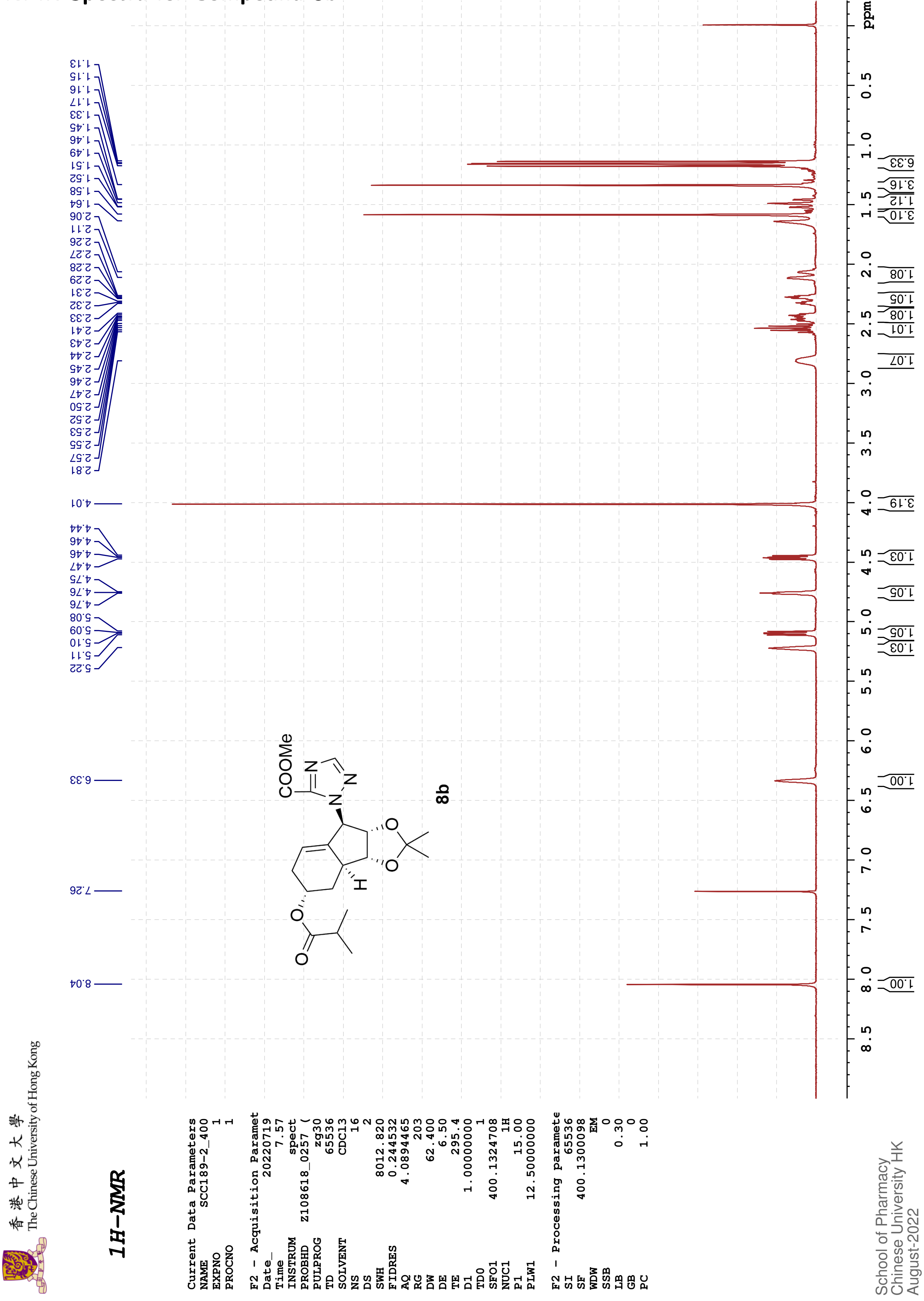

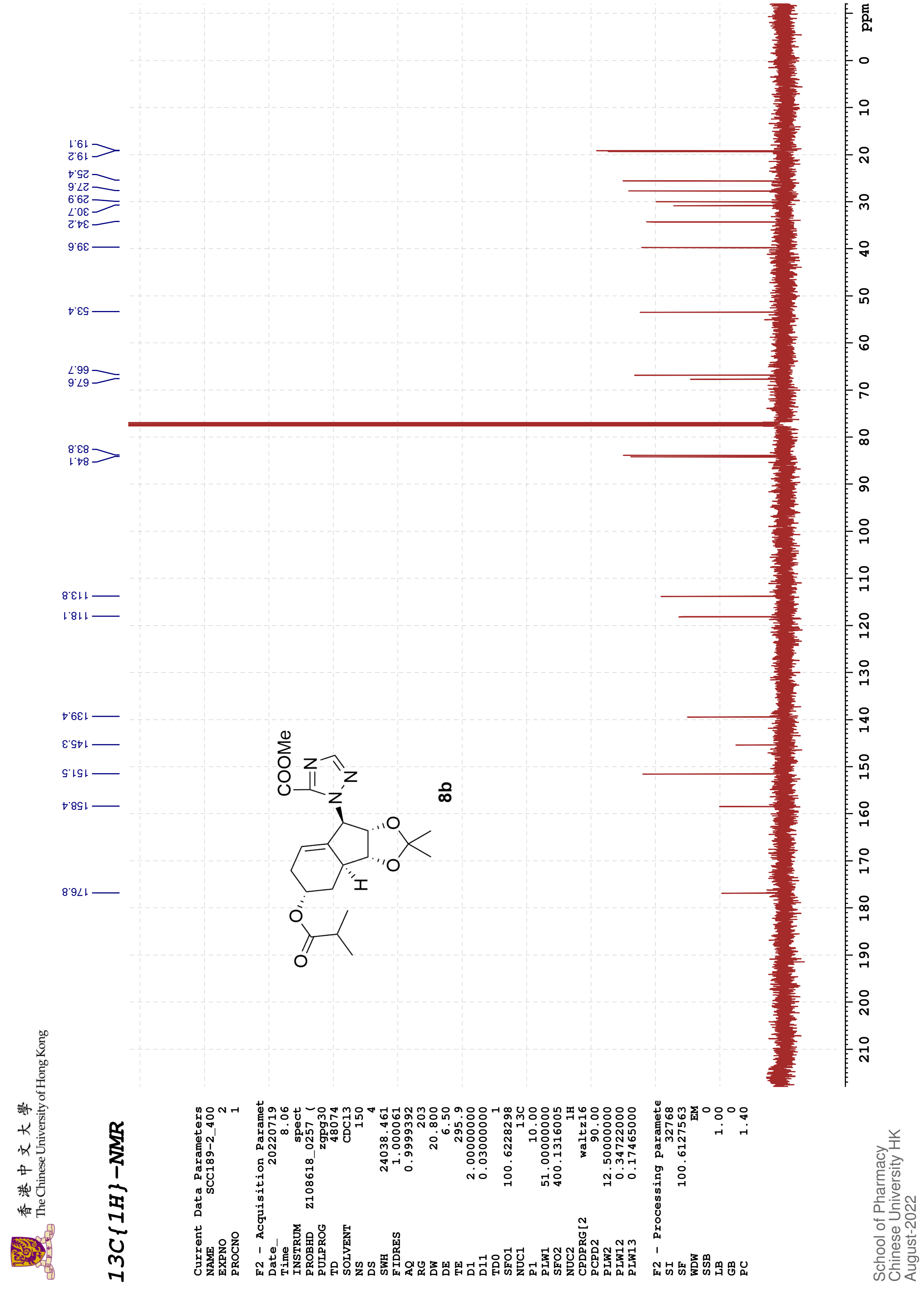

香港中文大學 The Chinese University of Hong Kong
13C{1H}-NMR
Current Data Parameters
NAME SCC189-2_400
EXPNO 2
PROCNO 1
F2 - Acquisition Paramet
Date_ 20220719
Time 8.06
INSTRUM spect
PROBHD Z108618_0257 (
PULPROG zgpg30
TD 48074
SOLVENT CDCl3
NS 150
DS 4
SWH 24038.461
FIDRES 1.000061
AQ 0.9999392
RG 203
DW 20.800
DE 6.50
TE 295.9
D1 2.00000000
D11 0.03000000
TD0 1
SFO1 100.6228298
NUC1 13C
P1 10.00
PLW1 51.00000000
SFO2 400.1316005
NUC2 1H
CPDPRG[2 waltz16
PCPD2 90.00
PLW2 12.50000000
PLW12 0.34722000
PLW13 0.17465000
F2 - Processing paramete
SI 32768
SF 100.6127563
WDW EM
SSB 0
LB 1.00
GB 0
PC 1.40
176.8
158.4
151.5
145.3
139.4
118.1
113.8
84.1
83.8
67.6
66.7
53.4
39.6
34.2
30.7
29.9
27.6
25.4
19.2
19.1
COOMe
N
N
N
O
O
O
O
H
8b
ppm
School of Pharmacy
Chinese University HK
August-2022

#### NMR-Spectra for Compound 11b

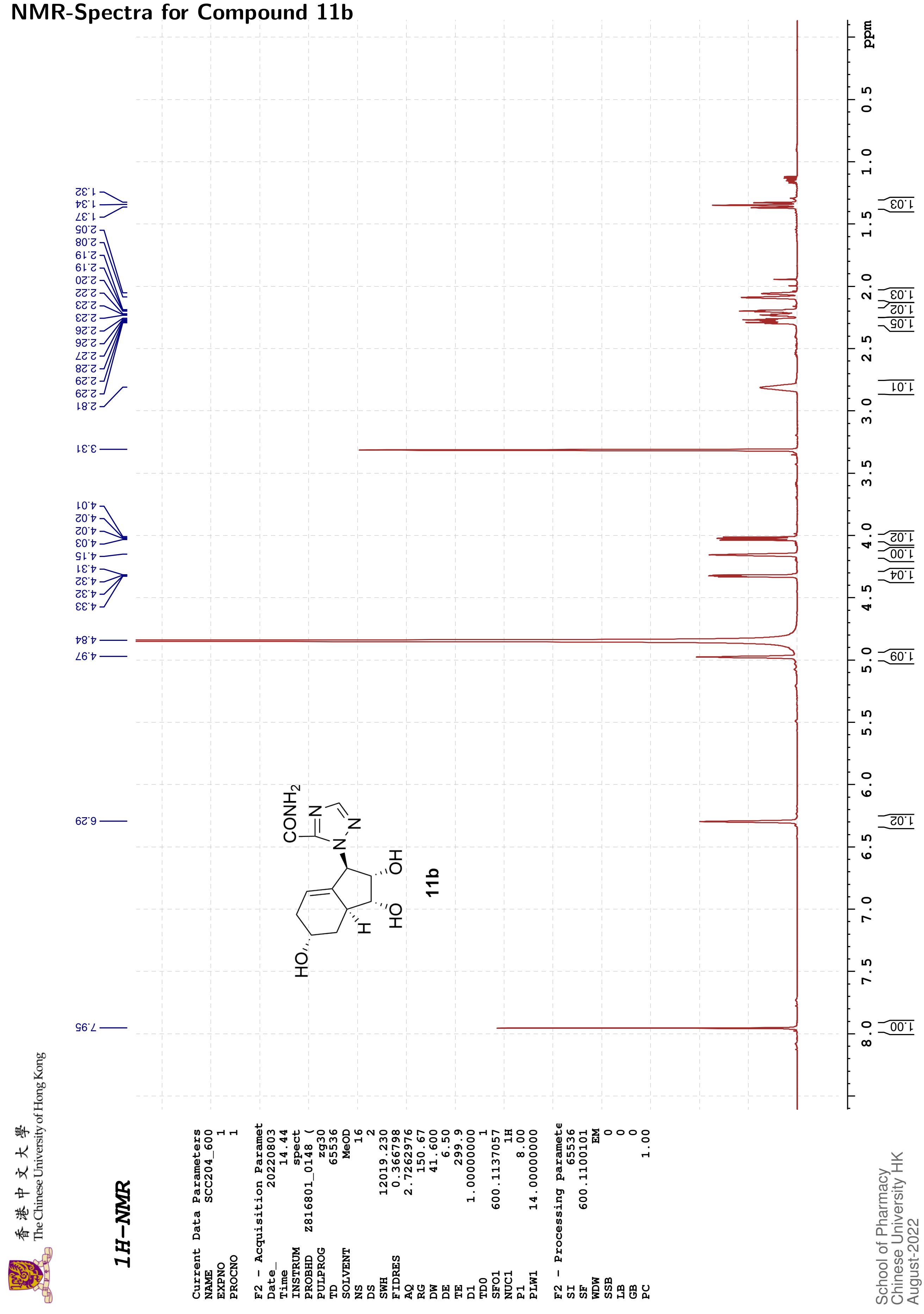

香港中文大學 The Chinese University of Hong Kong

#### 13C{1H}-NMR

```
Current Data Parameters
NAME          SCC204_600
EXPNO                  2
PROCNO                 1

F2 - Acquisition Paramet
Date_           20220803
Time               15.14
INSTRUM            spect
PROBHD    Z816801_0148 (
PULPROG          zgpg30
TD                 65536
SOLVENT             MeOD
NS                   569
DS                     4
SWH            36231.883
FIDRES          1.105709
AQ             0.9043968
RG                 98.37
DW                13.800
DE                  6.50
TE                 300.0
D1            2.00000000
D11           0.03000000
TD0                    1
SFO1         150.9128693
NUC1                 13C
P1                 12.00
PLW1        183.00000000
SFO2         600.1124004
NUC2                  1H
CPDPRG[2         waltz16
PCPD2              70.00
PLW2         14.00000000
PLW12         0.18286000
PLW13         0.09197600

F2 - Processing paramete
SI                 32768
SF           150.8975685
WDW                   EM
SSB                    0
LB                  1.00
GB                     0
PC                  1.40
```

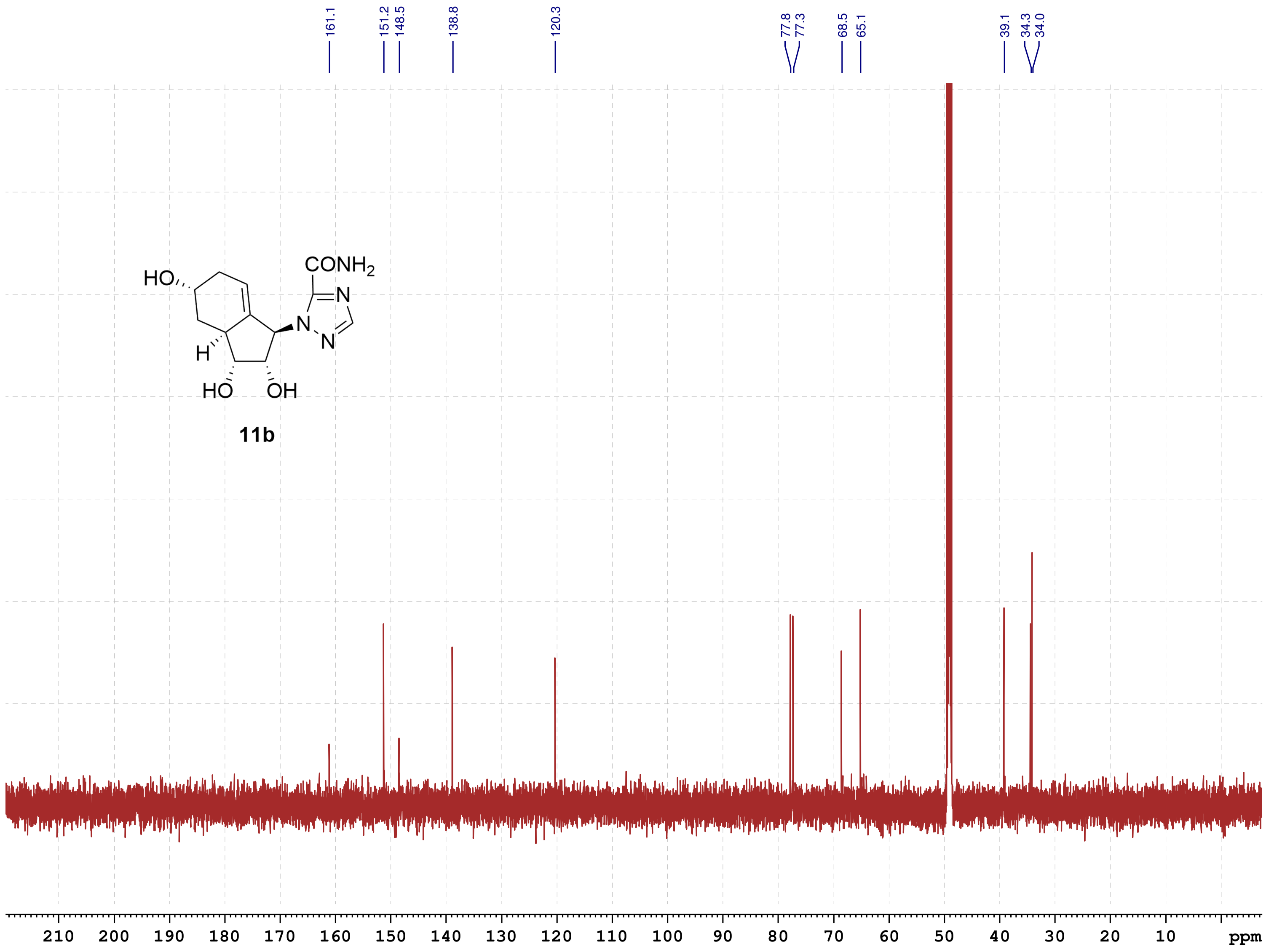

School of Pharmacy
Chinese University HK
August-2022

#### NMR-Spectra for Compound 11d

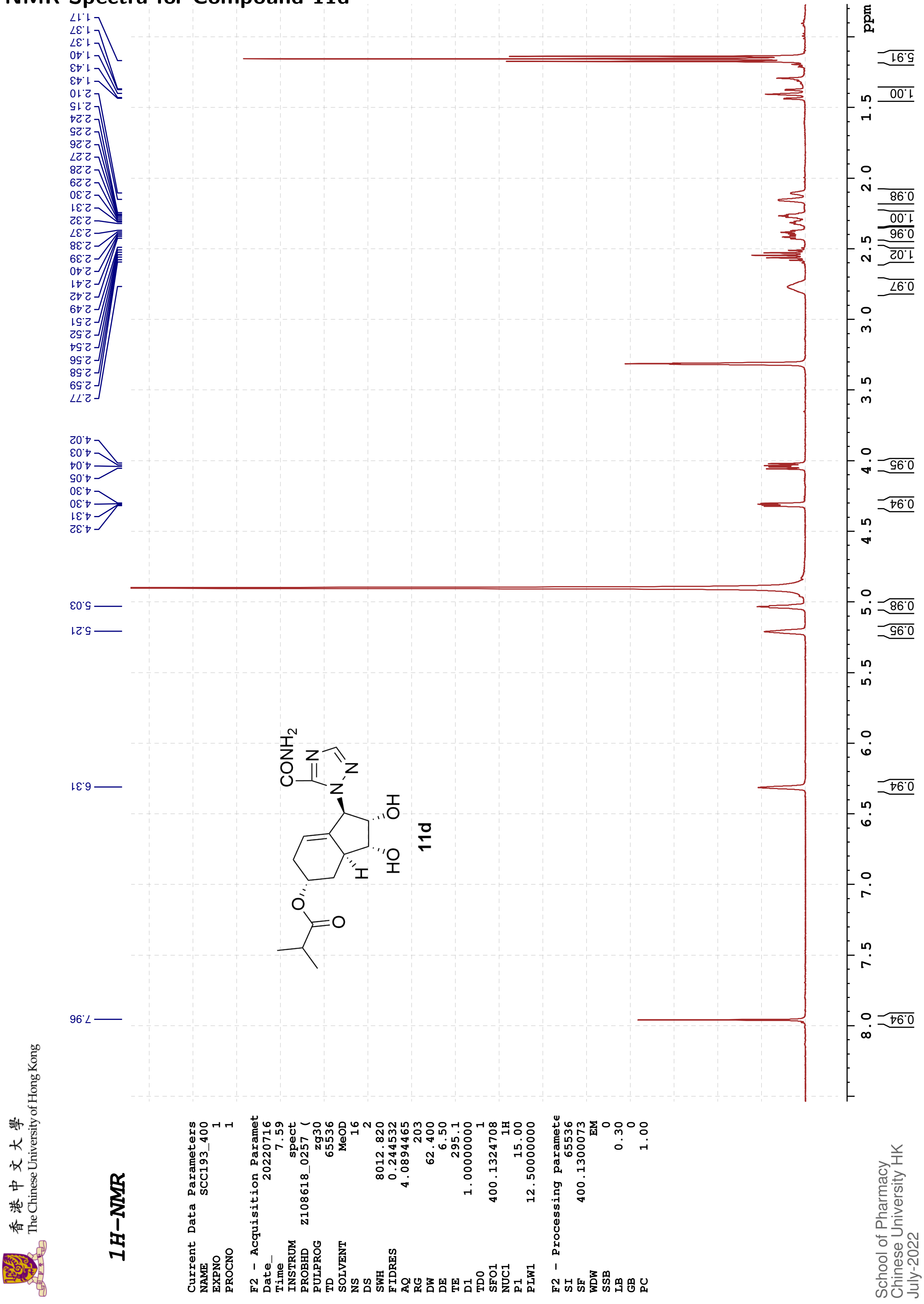

香港中文大學
The Chinese University of Hong Kong

*13C{1H}-NMR*

```
Current Data Parameters
NAME          SCC193_400
EXPNO                  2
PROCNO                 1

F2 - Acquisition Paramet
Date_           20220716
Time                8.11
INSTRUM            spect
PROBHD   Z108618_0257 (
PULPROG          zgpg30
TD                48074
SOLVENT             MeOD
NS                  206
DS                    4
SWH            24038.461
FIDRES          1.000061
AQ             0.9999392
RG                   203
DW                20.800
DE                  6.50
TE                 295.2
D1            2.00000000
D11           0.03000000
TD0                    1
SFO1         100.6228298
NUC1                 13C
P1                 10.00
PLW1         51.00000000
SFO2         400.1316005
NUC2                  1H
CPDPRG[2         waltz16
PCPD2              90.00
PLW2         12.50000000
PLW12         0.34722000
PLW13         0.17465000

F2 - Processing paramete
SI                 32768
SF           100.6127559
WDW                   EM
SSB                    0
LB                  2.00
GB                     0
PC                  1.40
```

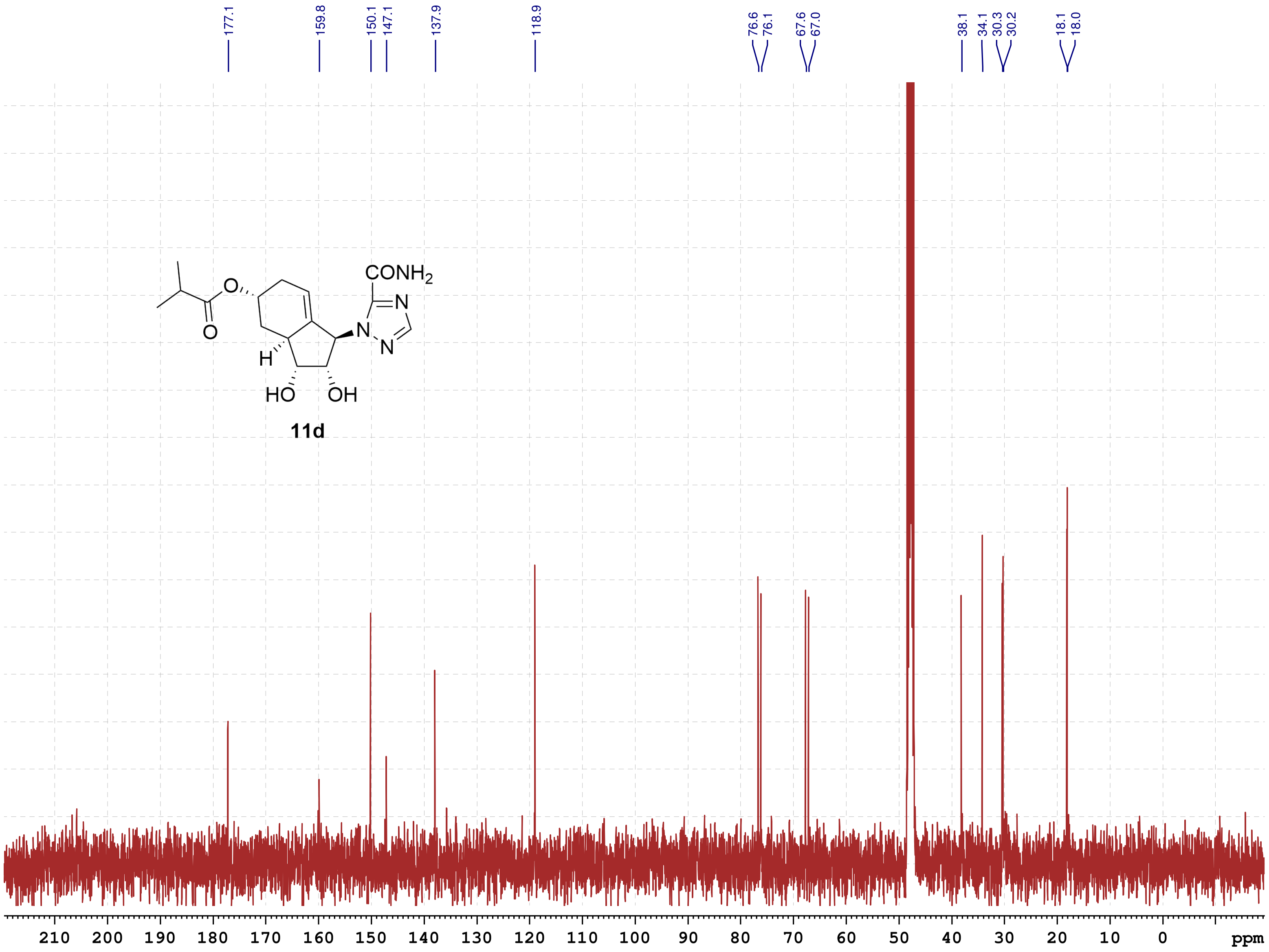

School of Pharmacy
Chinese University HK
July-2022

香港中文大學 The Chinese University of Hong Kong

#### COSY

Current Data Parameters
NAME SCC193_400
EXPNO 3
PROCNO 1

F2 - Acquisition Parameters
Date_ 20220716
Time 8.12 h
INSTRUM spect
PROBHD Z108618_0257 (
PULPROG cosygpqf
TD 2048
SOLVENT MeOD
NS 2
DS 4
SWH 4000.000 Hz
FIDRES 3.906250 Hz
AQ 0.2560000 sec
RG 203
DW 125.000 usec
DE 6.50 usec
TE 295.4 K
D0 0.00000300 sec
D1 1.48689198 sec
D13 0.00000400 sec
D16 0.00020000 sec
IN0 0.00025000 sec
TDav 1
SFO1 400.1320006 MHz
NUC1 1H
P0 15.00 usec
P1 15.00 usec
PLW1 12.50000000 W
GPNAM[1] SINE.100
GPZ1 10.00 %
P16 1000.00 usec

F1 - Acquisition parameters
TD 128
SFO1 400.132 MHz
FIDRES 62.500000 Hz
SW 9.997 ppm
FnMODE QF

F2 - Processing parameters
SI 1024
SF 400.1300097 MHz
WDW SINE
SSB 0
LB 0 Hz
GB 0
PC 1.40

F1 - Processing parameters
SI 1024
MC2 QF
SF 400.1300081 MHz
WDW SINE
SSB 0
LB 0 Hz
GB 0

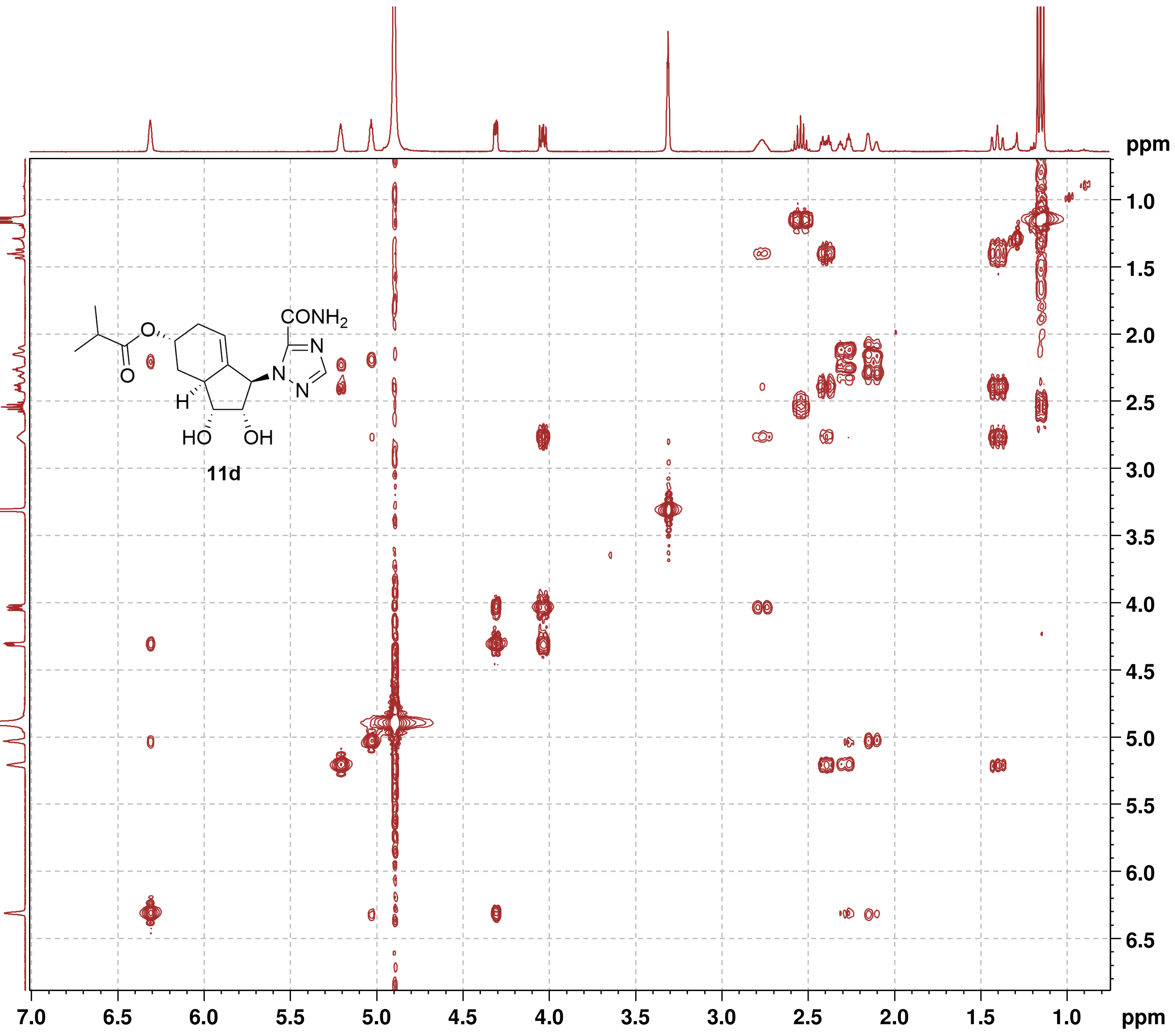

香港中文大學
The Chinese University of Hong Kong

**NOESY**

```
Current Data Parameters
NAME            SCC193_400
EXPNO                    4
PROCNO                   1

F2 - Acquisition Parameters
Date_             20220716
Time                  8.21 h
INSTRUM              spect
PROBHD      Z108618_0257 (
PULPROG        noesygpphpp
TD                    2048
SOLVENT               MeOD
NS                       2
DS                       4
SWH               4000.000 Hz
FIDRES            3.906250 Hz
AQ               0.2560000 sec
RG                     203
DW                 125.000 usec
DE                    6.50 usec
TE                   295.1 K
D0              0.00010590 sec
D1              2.00000000 sec
D8              0.85000002 sec
D11             0.03000000 sec
D12             0.00002000 sec
D16             0.00020000 sec
IN0             0.00025000 sec
TDav                     1
SFO1           400.1318806 MHz
NUC1                    1H
P1                   15.00 usec
P2                   30.00 usec
P17                2500.00 usec
PLW1           12.50000000 W
PLW10           3.12500000 W
GPNAM[1]         SMSQ10.100
GPZ1                 40.00 %
P16                1000.00 usec

F1 - Acquisition parameters
TD                     128
SFO1              400.1319 MHz
FIDRES           62.500000 Hz
SW                   9.997 ppm
FnMODE         States-TPPI

F2 - Processing parameters
SI                    1024
SF             400.1300072 MHz
WDW                  QSINE
SSB                      2
LB                       0 Hz
GB                       0
PC                    1.00

F1 - Processing parameters
SI                    1024
MC2            States-TPPI
SF             400.1300086 MHz
WDW                  QSINE
SSB                      2
LB                       0 Hz
GB                       0
```

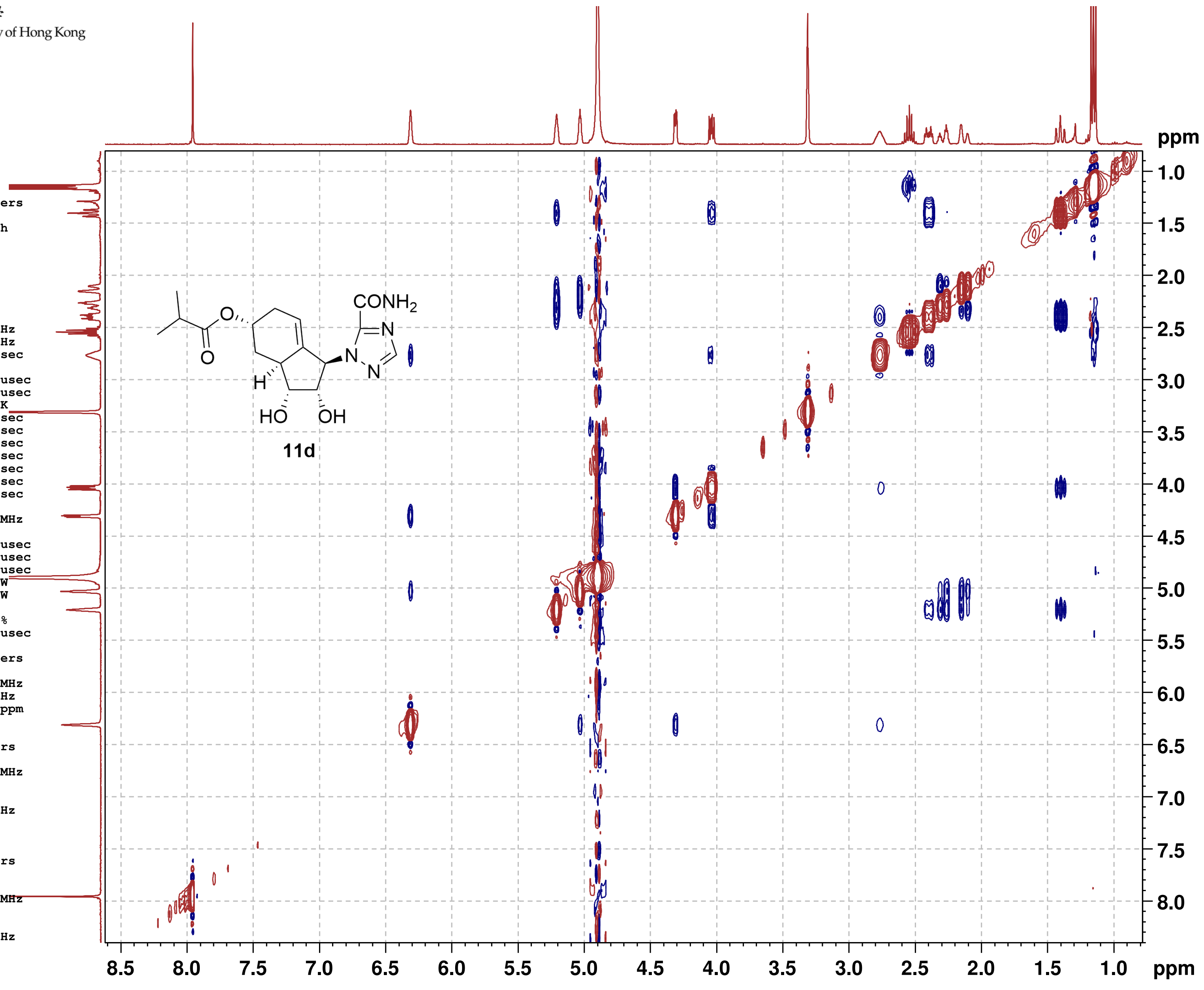

#### NMR-Spectra for Compound 9b

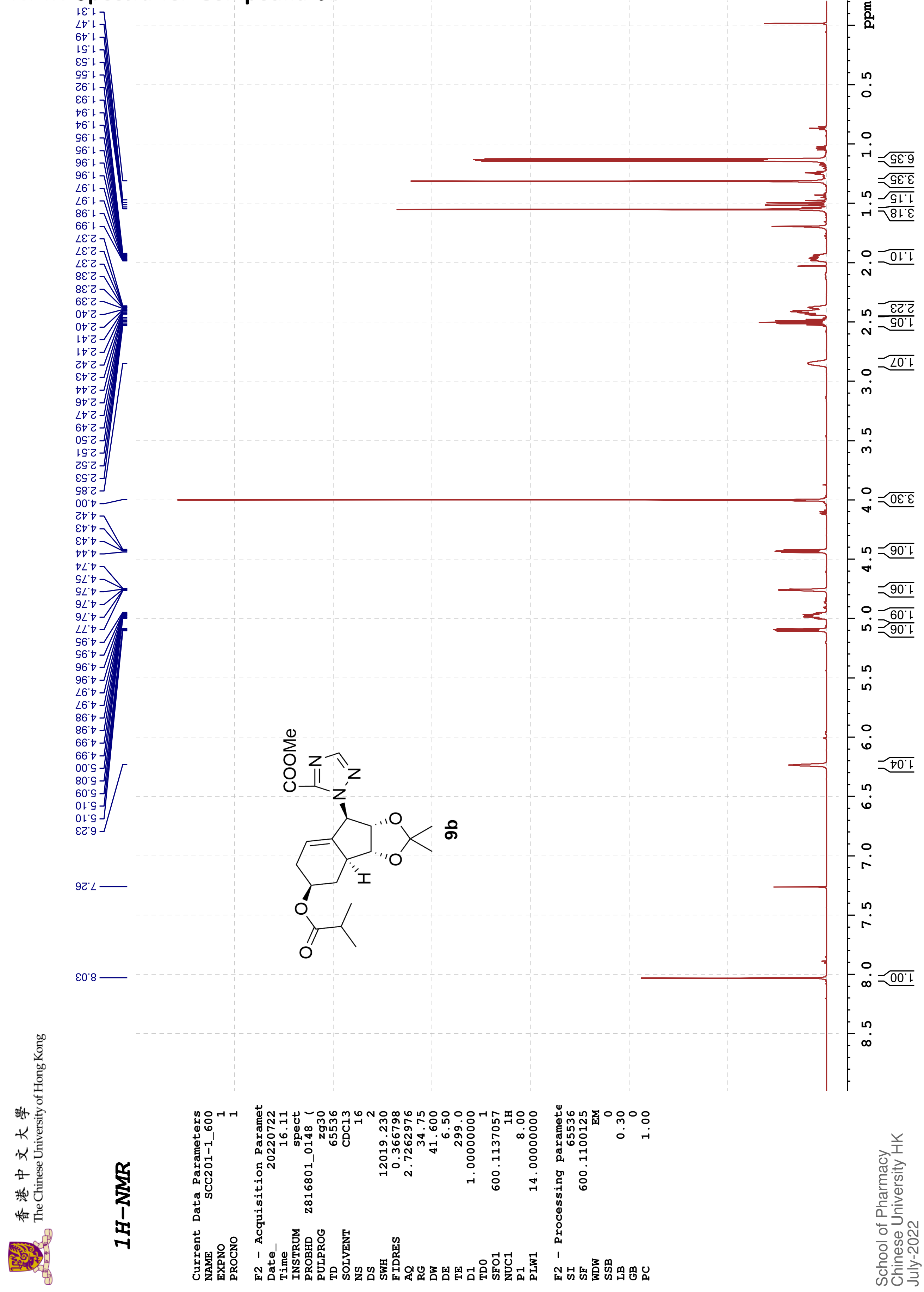

香港中文大學 The Chinese University of Hong Kong

#### 13C{1H}-NMR

| | |
|---|---|
| Current Data Parameters | |
| NAME | SCC201-1_600 |
| EXPNO | 2 |
| PROCNO | 1 |
| F2 - Acquisition Paramet | |
| Date_ | 20220722 |
| Time | 16.18 |
| INSTRUM | spect |
| PROBHD | Z816801_0148 ( |
| PULPROG | zgpg30 |
| TD | 65536 |
| SOLVENT | CDCl3 |
| NS | 125 |
| DS | 4 |
| SWH | 36231.883 |
| FIDRES | 1.105709 |
| AQ | 0.9043968 |
| RG | 60.39 |
| DW | 13.800 |
| DE | 6.50 |
| TE | 299.2 |
| D1 | 2.00000000 |
| D11 | 0.03000000 |
| TD0 | 1 |
| SFO1 | 150.9128693 |
| NUC1 | 13C |
| P1 | 12.00 |
| PLW1 | 183.00000000 |
| SFO2 | 600.1124004 |
| NUC2 | 1H |
| CPDPRG[2 | waltz16 |
| PCPD2 | 70.00 |
| PLW2 | 14.00000000 |
| PLW12 | 0.18286000 |
| PLW13 | 0.09197600 |
| F2 - Processing paramete | |
| SI | 32768 |
| SF | 150.8977608 |
| WDW | EM |
| SSB | 0 |
| LB | 2.00 |
| GB | 0 |
| PC | 1.40 |

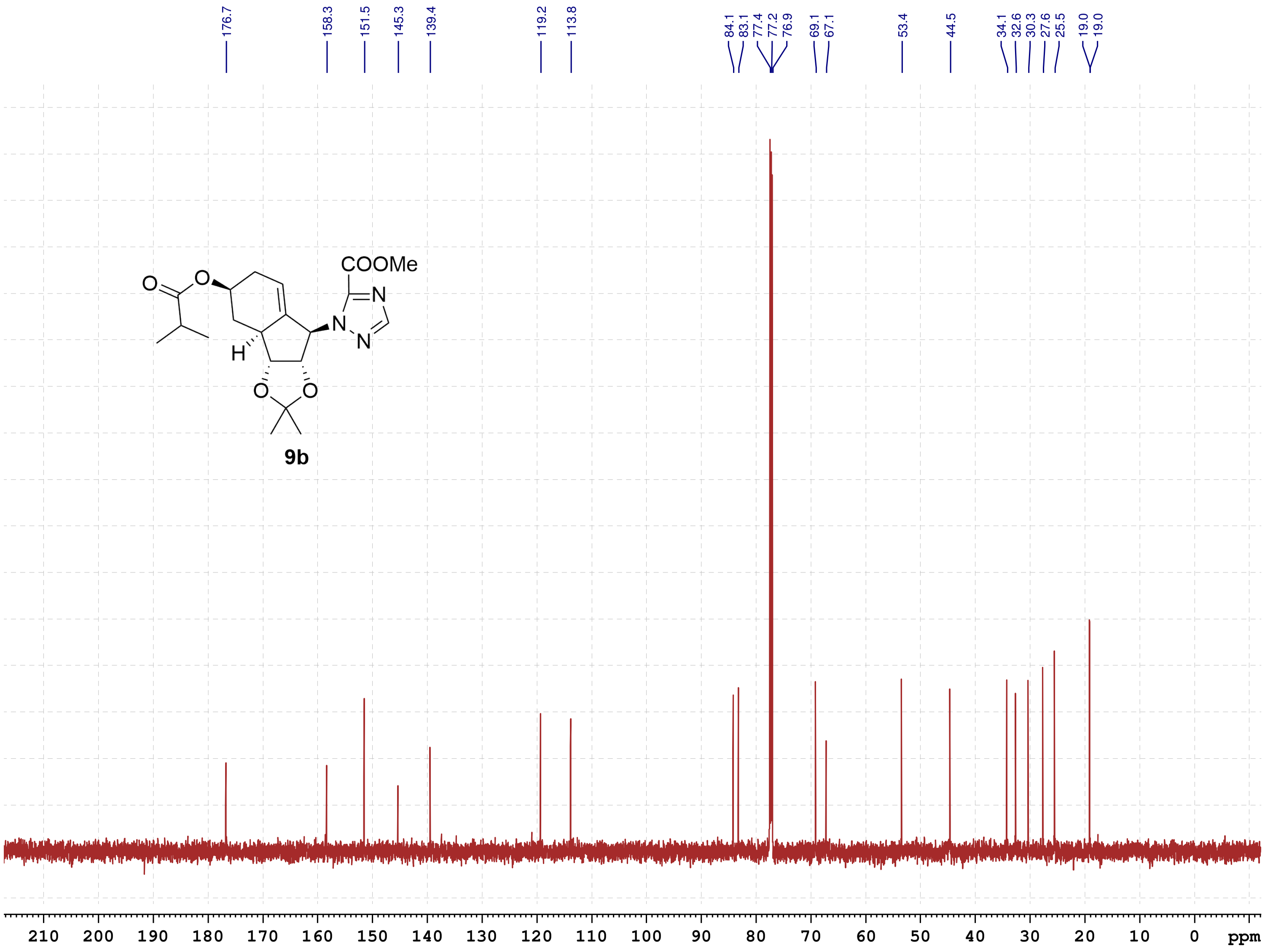

School of Pharmacy
Chinese University HK
July-2022

#### NMR-Spectra for Compound 12b

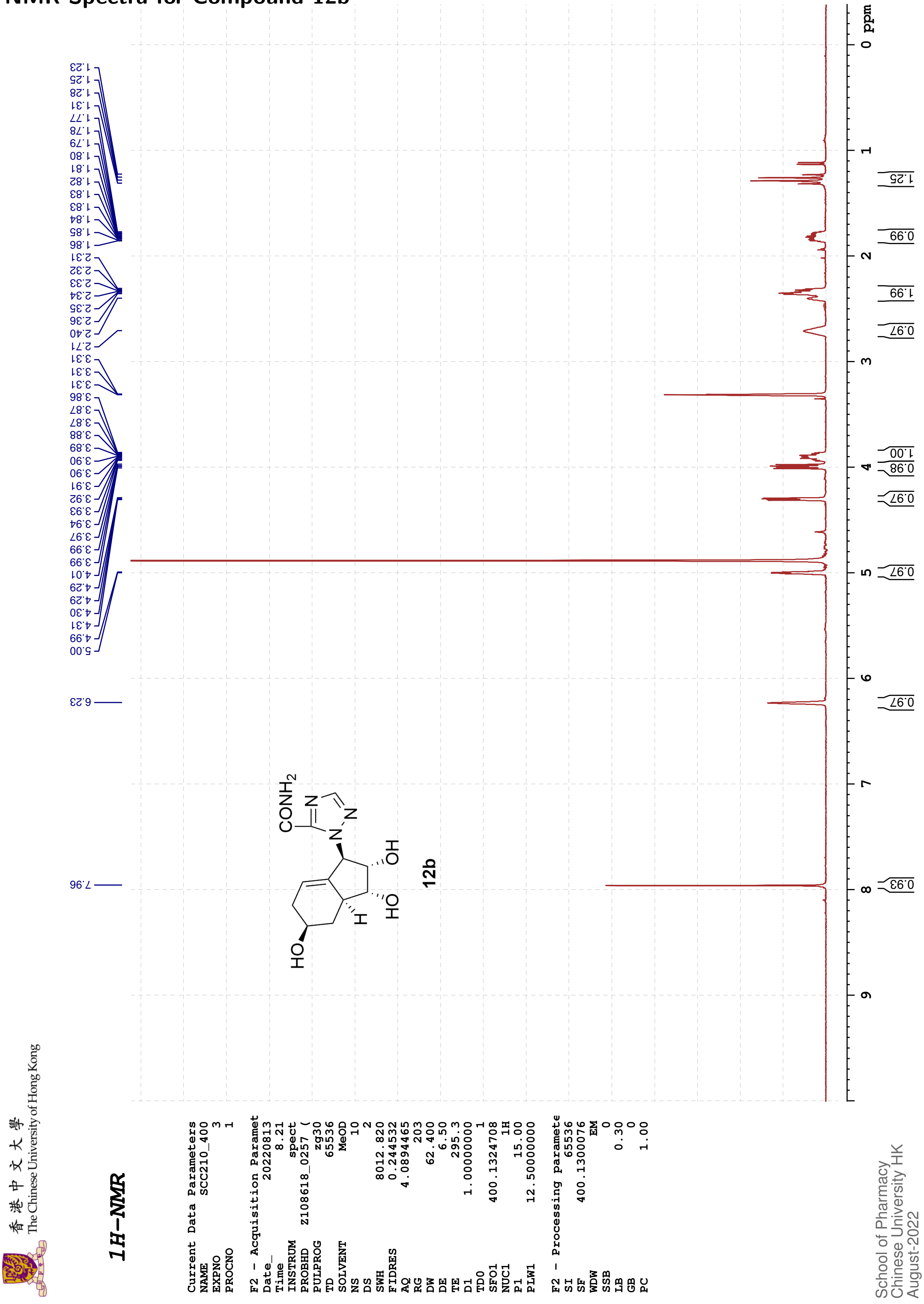

The Chinese University of Hong Kong

#### 13C{1H}-NMR

Current Data Parameters
NAME SCC210_400
EXPNO 4
PROCNO 1

F2 - Acquisition Paramet
Date_ 20220813
Time 8.31
INSTRUM spect
PROBHD Z108618_0257 (
PULPROG zgpg30
TD 48074
SOLVENT MeOD
NS 170
DS 4
SWH 24038.461
FIDRES 1.000061
AQ 0.9999392
RG 203
DW 20.800
DE 6.50
TE 295.6
D1 2.00000000
D11 0.03000000
TD0 1
SFO1 100.6228298
NUC1 13C
P1 10.00
PLW1 51.00000000
SFO2 400.1316005
NUC2 1H
CPDPRG[2 waltz16
PCPD2 90.00
PLW2 12.50000000
PLW12 0.34722000
PLW13 0.17465000

F2 - Processing paramete
SI 32768
SF 100.6126282
WDW EM
SSB 0
LB 1.00
GB 0
PC 1.40

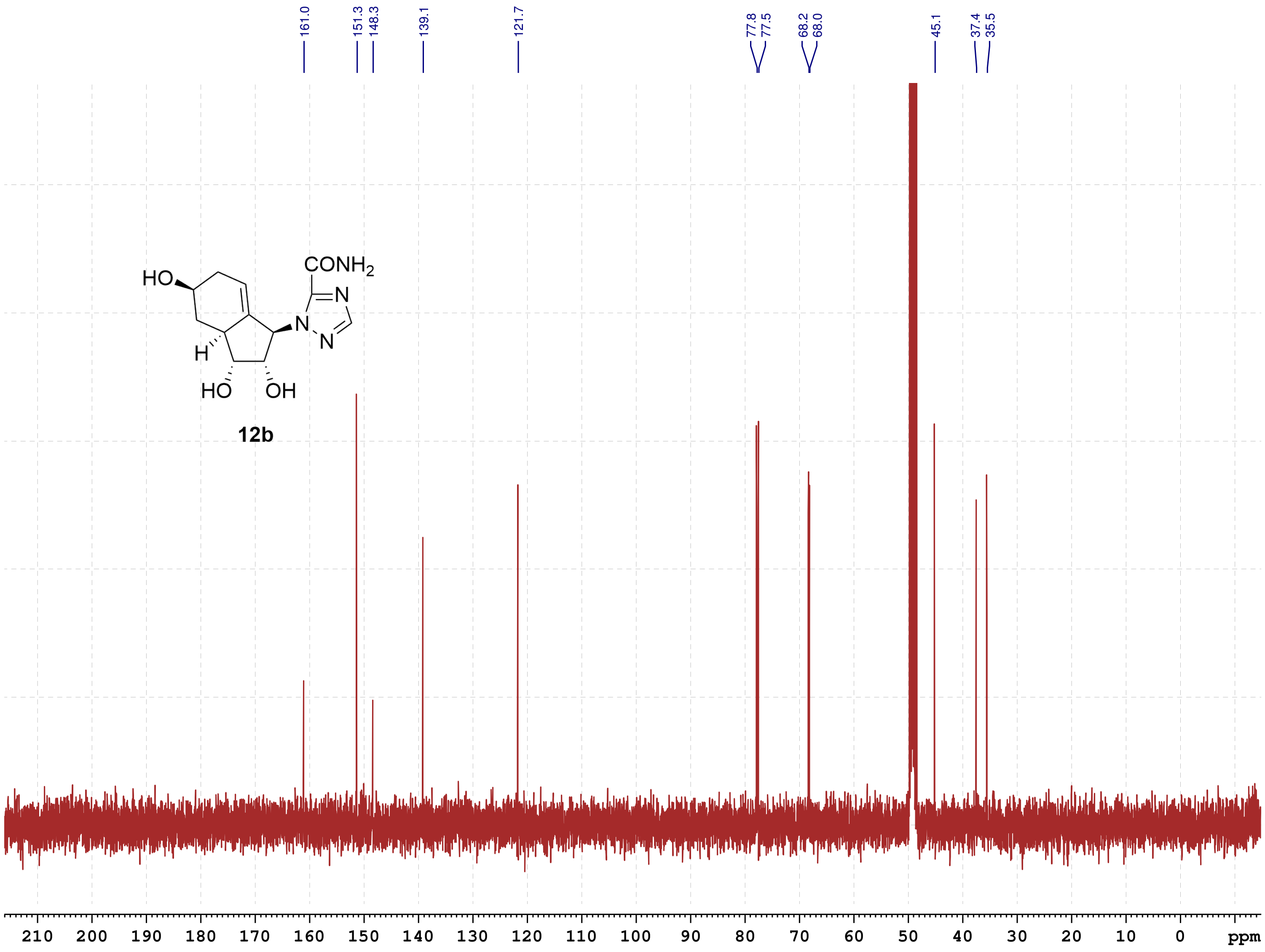

12b

School of Pharmacy
Chinese University HK
August-2022

#### NMR-Spectra for Compound 12d

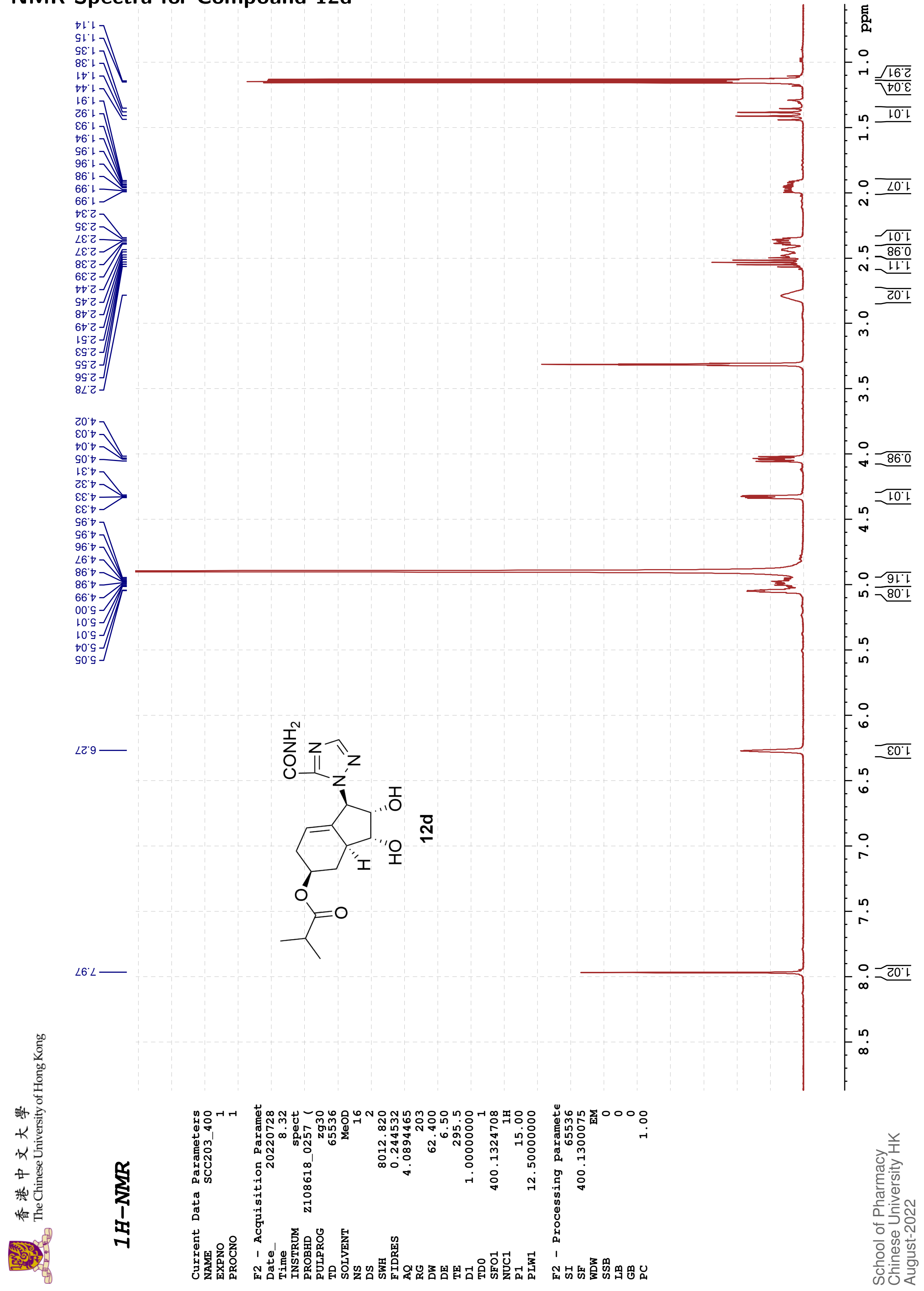

香港中文大學 The Chinese University of Hong Kong

#### 13C{1H}-NMR

```
Current Data Parameters
NAME              SCC203_400
EXPNO                      2
PROCNO                     1

F2 - Acquisition Paramet
Date_               20220728
Time                    8.57
INSTRUM                spect
PROBHD   Z108618_0257 (
PULPROG              zgpg30
TD                     48074
SOLVENT                 MeOD
NS                       473
DS                         4
SWH                24038.461
FIDRES              1.000061
AQ                 0.9999392
RG                       203
DW                    20.800
DE                      6.50
TE                     295.9
D1                2.00000000
D11               0.03000000
TD0                        1
SFO1             100.6228298
NUC1                     13C
P1                     10.00
PLW1             51.00000000
SFO2             400.1316005
NUC2                      1H
CPDPRG[2             waltz16
PCPD2                  90.00
PLW2             12.50000000
PLW12             0.34722000
PLW13             0.17465000

F2 - Processing paramete
SI                     32768
SF               100.6126289
WDW                       EM
SSB                        0
LB                      3.00
GB                         0
PC                      1.40
```

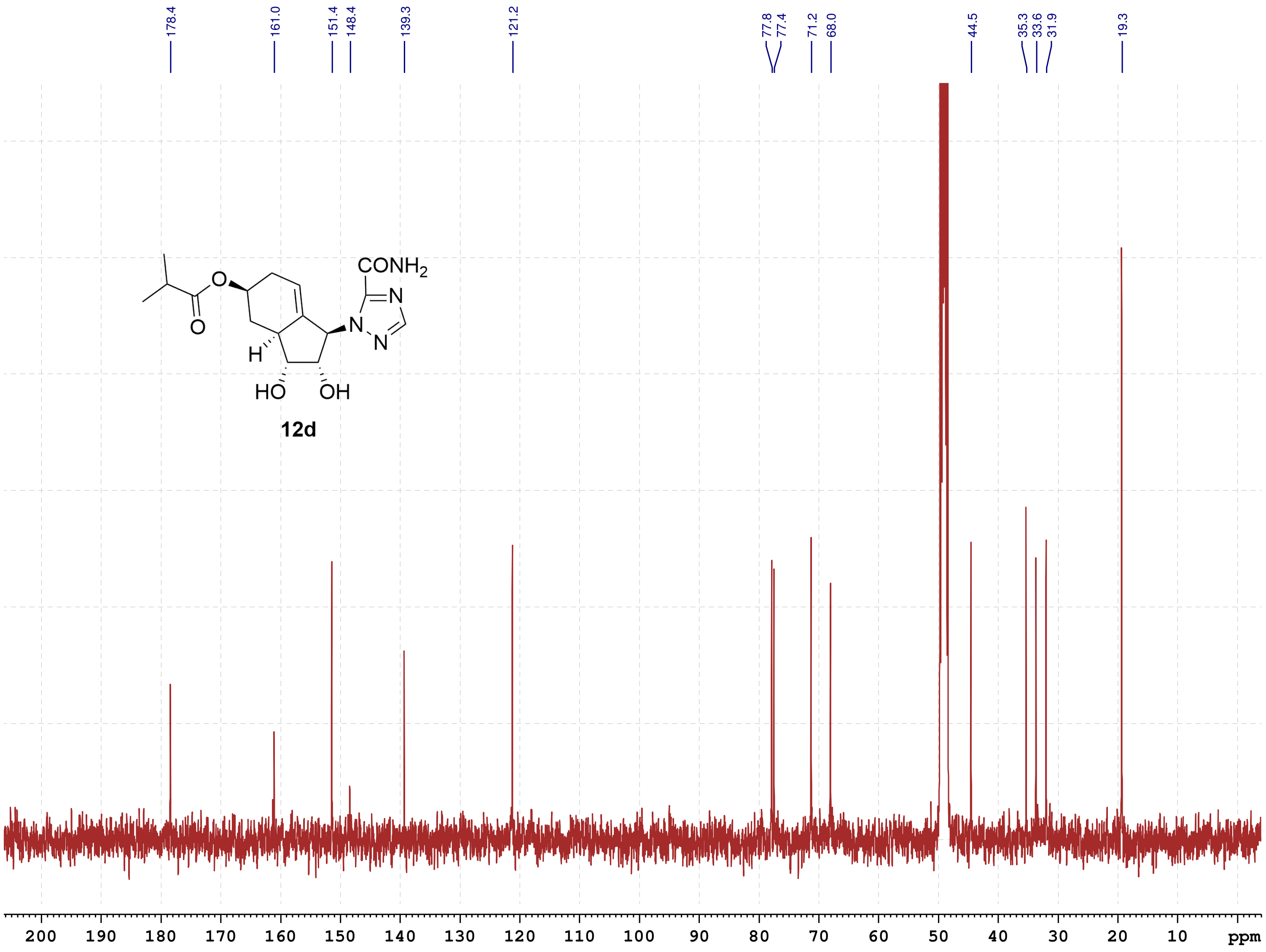

12d

School of Pharmacy
Chinese University HK
August-2022

#### NMR-Spectra for Compound 14

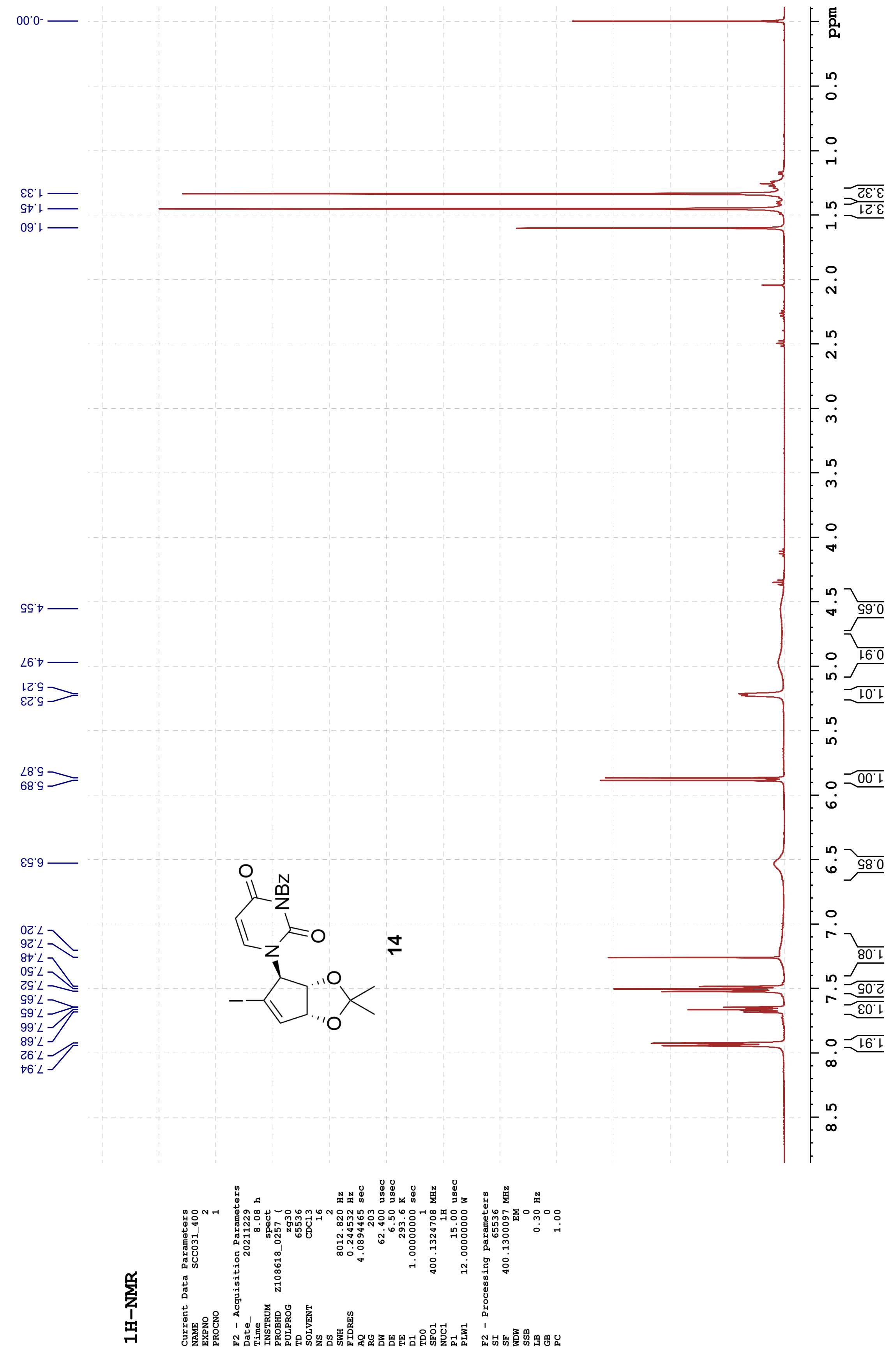

**13C-{1H}-NMR**

```
Current Data Parameters
NAME          SCC031_400
EXPNO                  7
PROCNO                 1

F2 - Acquisition Parameters
Date_           20211229
Time                9.44 h
INSTRUM            spect
PROBHD    Z108618_0257 (
PULPROG          zgpg30
TD                 65536
SOLVENT            CDCl3
NS                   442
DS                     4
SWH            24038.461 Hz
FIDRES          0.733596 Hz
AQ             1.3631488 sec
RG                   203
DW                20.800 usec
DE                  6.50 usec
TE                 293.8 K
D1            2.00000000 sec
D11           0.03000000 sec
TD0                    1
SFO1         100.6228298 MHz
NUC1                 13C
P1                 10.00 usec
PLW1         47.00000000 W
SFO2         400.1316005 MHz
NUC2                  1H
CPDPRG[2         waltz16
PCPD2              90.00 usec
PLW2         12.00000000 W
PLW12         0.33333001 W
PLW13         0.16766000 W

F2 - Processing parameters
SI                 32768
SF           100.6127572 MHz
WDW                   EM
SSB                    0
LB                  3.00 Hz
GB                     0
PC                  1.40
```

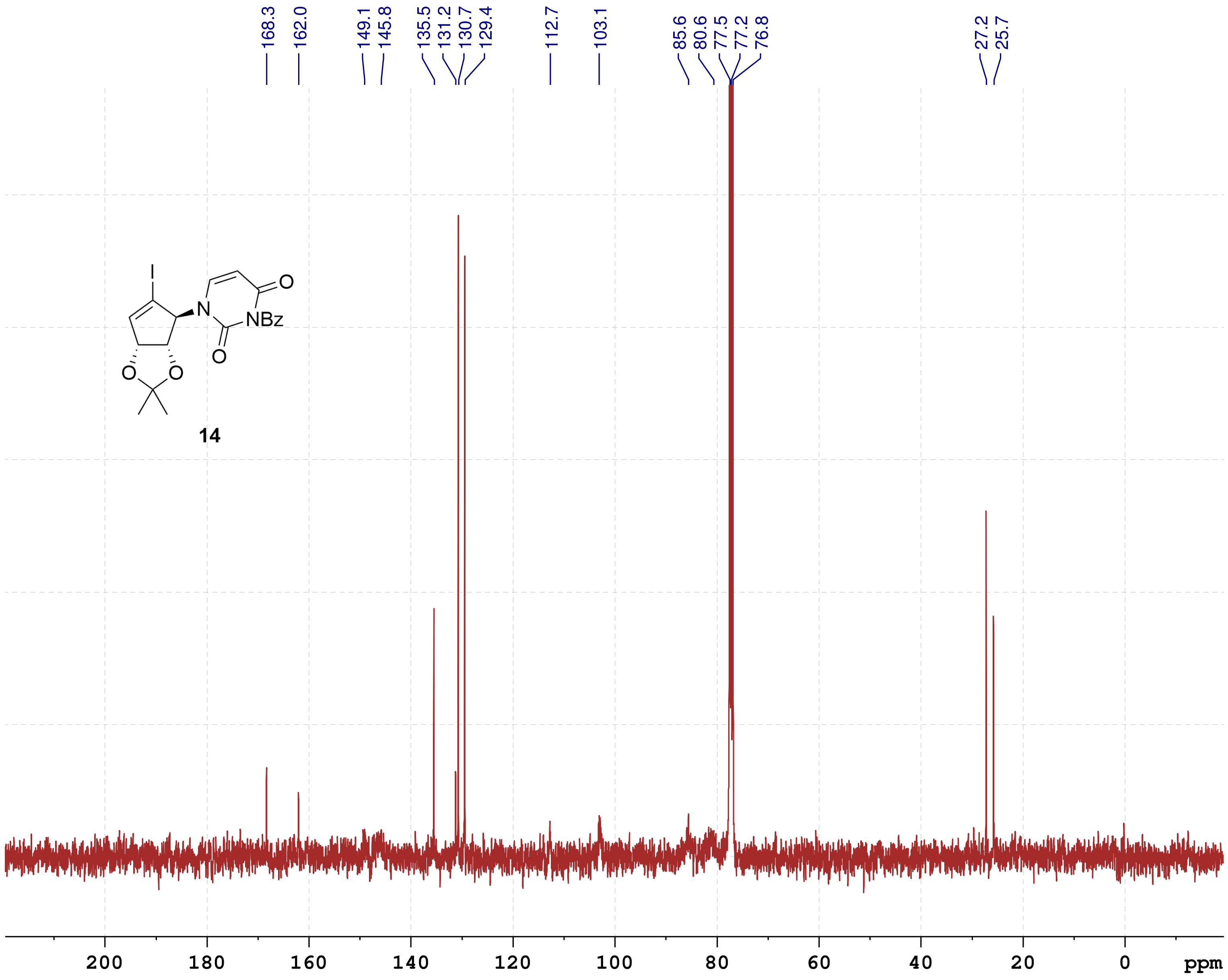

#### HSQC

```
Current Data Parameters
NAME          SCC031_400
EXPNO                  5
PROCNO                 1

F2 - Acquisition Parameters
Date_           20211229
Time                9.19 h
INSTRUM            spect
PROBHD   Z108618_0257 (
PULPROG      hsqcetgpsi2
TD                  1024
SOLVENT            CDCl3
NS                     2
DS                    16
SWH             5197.505 Hz
FIDRES         10.151378 Hz
AQ             0.0985088 sec
RG                   203
DW                96.200 usec
DE                  6.50 usec
TE                 293.7 K
CNST2        145.0000000
D0            0.00000300 sec
D1            1.50000000 sec
D4            0.00172414 sec
D11           0.03000000 sec
D16           0.00020000 sec
D24           0.00086207 sec
IN0           0.00002620 sec
TDav                   1
ZGOPTNS
SFO1         400.1324708 MHz
NUC1                  1H
P1                 15.00 usec
P2                 30.00 usec
P28              1000.00 usec
PLW1         12.00000000 W
SFO2         100.6218237 MHz
NUC2                 13C
CPDPRG[2            garp
P3                 10.00 usec
P4                 20.00 usec
PCPD2              80.00 usec
PLW2         47.00000000 W
PLW12         0.73438001 W
GPNAM[1]      SMSQ10.100
GPZ1               80.00 %
GPNAM[2]      SMSQ10.100
GPZ2               20.10 %
GPNAM[3]      SMSQ10.100
GPZ3               11.00 %
GPNAM[4]      SMSQ10.100
GPZ4               -5.00 %
P16              1000.00 usec
P19               600.00 usec

F1 - Acquisition parameters
TD                   256
SFO1            100.6218 MHz
FIDRES        149.093506 Hz
SW               189.660 ppm
FnMODE      Echo-Antiecho

F2 - Processing parameters
SI                  1024
SF           400.1300094 MHz
WDW                QSINE
SSB                    2
LB                     0 Hz
GB                     0
PC                  1.40

F1 - Processing parameters
SI                  1024
MC2         echo-antiecho
SF           100.6127761 MHz
WDW                QSINE
SSB                    2
LB                     0 Hz
GB                     0
```

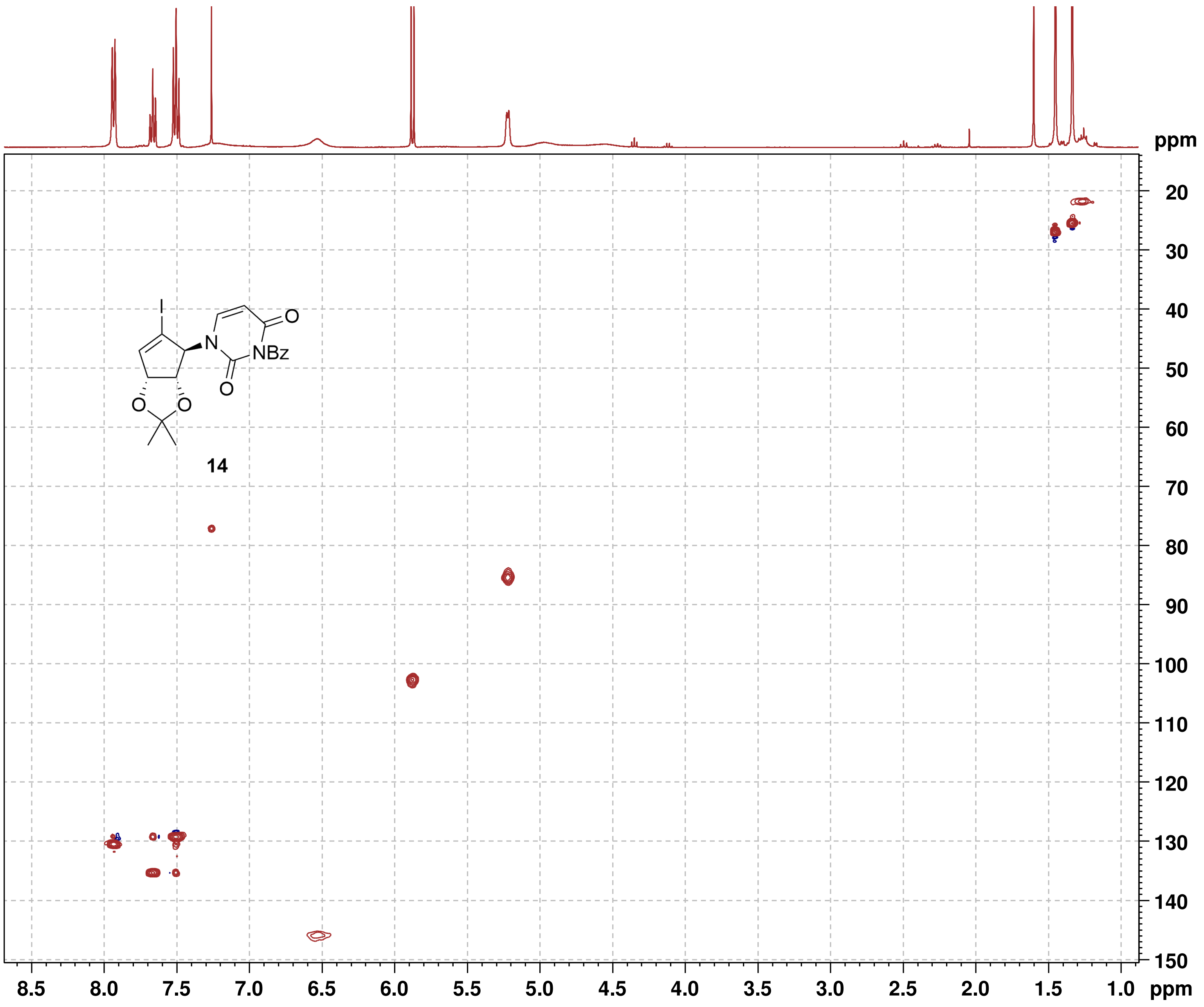

**HMBC**

```
Current Data Parameters
NAME          SCC031_400
EXPNO                  4
PROCNO                 1

F2 - Acquisition Parameters
Date_           20211229
Time                9.06 h
INSTRUM            spect
PROBHD   Z108618_0257 (
PULPROG     hmbcgplpndqf
TD                  2048
SOLVENT            CDCl3
NS                     3
DS                    16
SWH             5197.505 Hz
FIDRES          5.075689 Hz
AQ             0.1970176 sec
RG                   203
DW                96.200 usec
DE                  6.50 usec
TE                 293.7 K
CNST2        145.0000000
CNST13        10.0000000
D0            0.00000300 sec
D1            1.50000000 sec
D2            0.00344828 sec
D6            0.05000000 sec
D16           0.00020000 sec
IN0           0.00002260 sec
TDav                   1
SFO1         400.1325208 MHz
NUC1                  1H
P1                 15.00 usec
P2                 30.00 usec
PLW1         12.00000000 W
SFO2         100.6228298 MHz
NUC2                 13C
P3                 10.00 usec
PLW2         47.00000000 W
GPNAM[1]      SMSQ10.100
GPZ1               50.00 %
GPNAM[2]      SMSQ10.100
GPZ2               30.00 %
GPNAM[3]      SMSQ10.100
GPZ3               40.10 %
P16              1000.00 usec

F1 - Acquisition parameters
TD                   128
SFO1            100.6228 MHz
FIDRES        345.685852 Hz
SW               219.870 ppm
FnMODE                QF

F2 - Processing parameters
SI                  2048
SF           400.1300095 MHz
WDW                 SINE
SSB                    0
LB                     0 Hz
GB                     0
PC                  1.40

F1 - Processing parameters
SI                  1024
MC2                   QF
SF           100.6127437 MHz
WDW                 SINE
SSB                    0
LB                     0 Hz
GB                     0
```

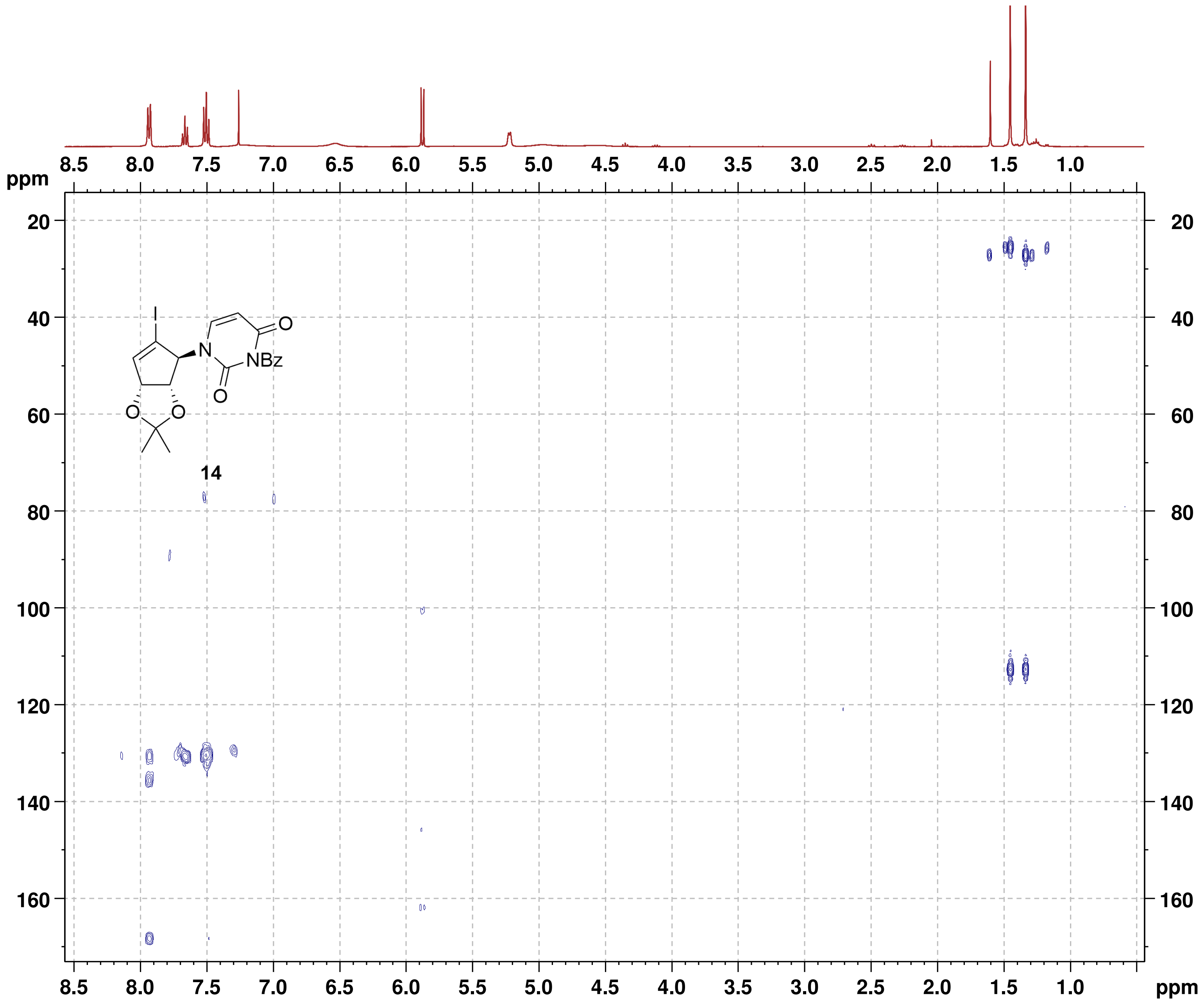

#### NMR-Spectra for Compound S4

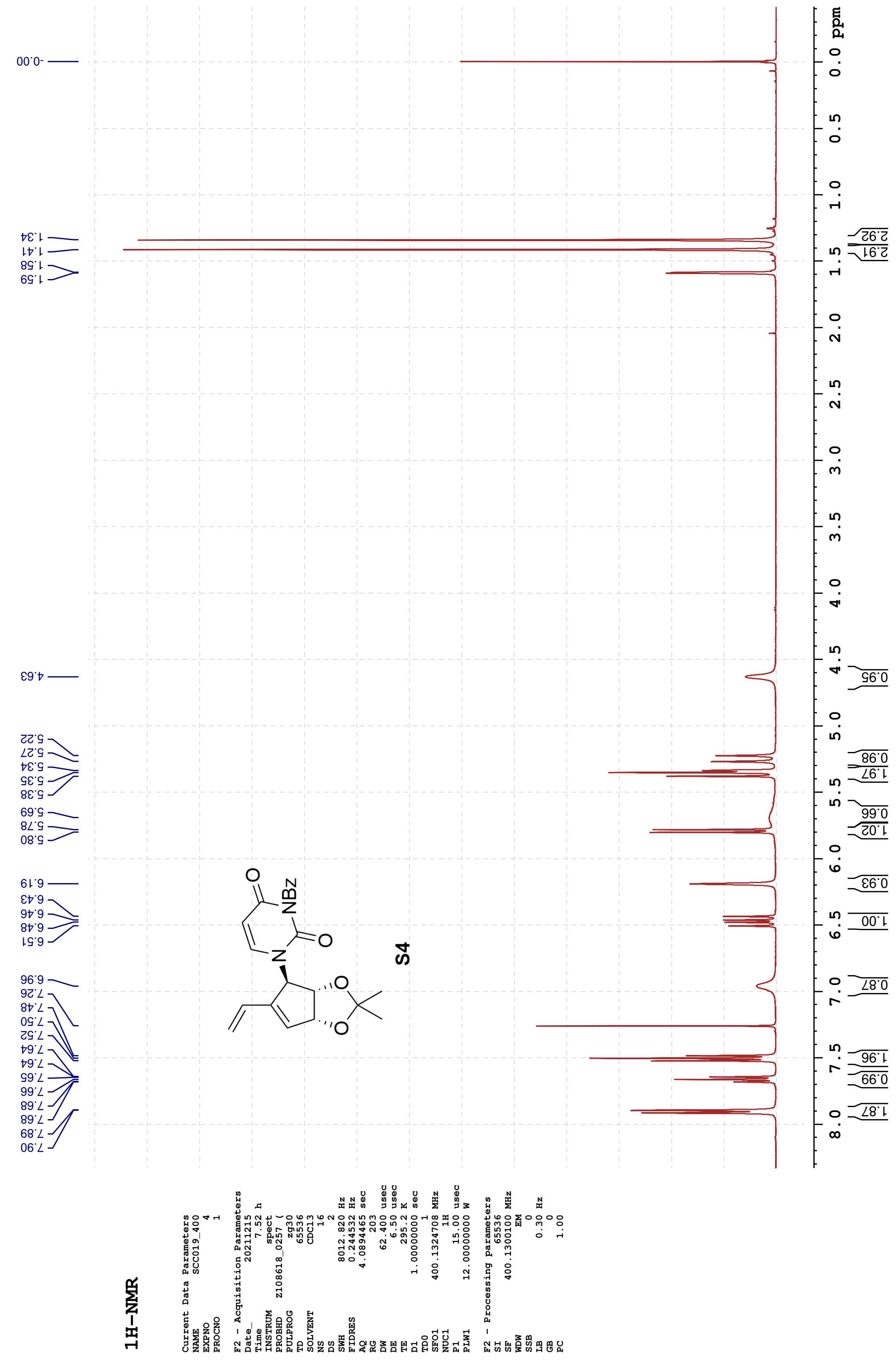

#### 13C-{1H}-NMR

Current Data Parameters

| | |
|---|---|
| NAME | SCC019_400 |
| EXPNO | 5 |
| PROCNO | 1 |

F2 - Acquisition Parameters

| | |
|---|---|
| Date_ | 20211215 |
| Time | 9.03 h |
| INSTRUM | spect |
| PROBHD | Z108618_0257 ( |
| PULPROG | zgpg30 |
| TD | 65536 |
| SOLVENT | CDCl3 |
| NS | 1024 |
| DS | 4 |
| SWH | 24038.461 Hz |
| FIDRES | 0.733596 Hz |
| AQ | 1.3631488 sec |
| RG | 203 |
| DW | 20.800 usec |
| DE | 6.50 usec |
| TE | 295.3 K |
| D1 | 2.00000000 sec |
| D11 | 0.03000000 sec |
| TD0 | 1 |
| SFO1 | 100.6228298 MHz |
| NUC1 | 13C |
| P1 | 10.00 usec |
| PLW1 | 47.00000000 W |
| SFO2 | 400.1316005 MHz |
| NUC2 | 1H |
| CPDPRG[2 | waltz16 |
| PCPD2 | 90.00 usec |
| PLW2 | 12.00000000 W |
| PLW12 | 0.33333001 W |
| PLW13 | 0.16766000 W |

F2 - Processing parameters

| | |
|---|---|
| SI | 32768 |
| SF | 100.6127564 MHz |
| WDW | EM |
| SSB | 0 |
| LB | 2.00 Hz |
| GB | 0 |
| PC | 1.40 |

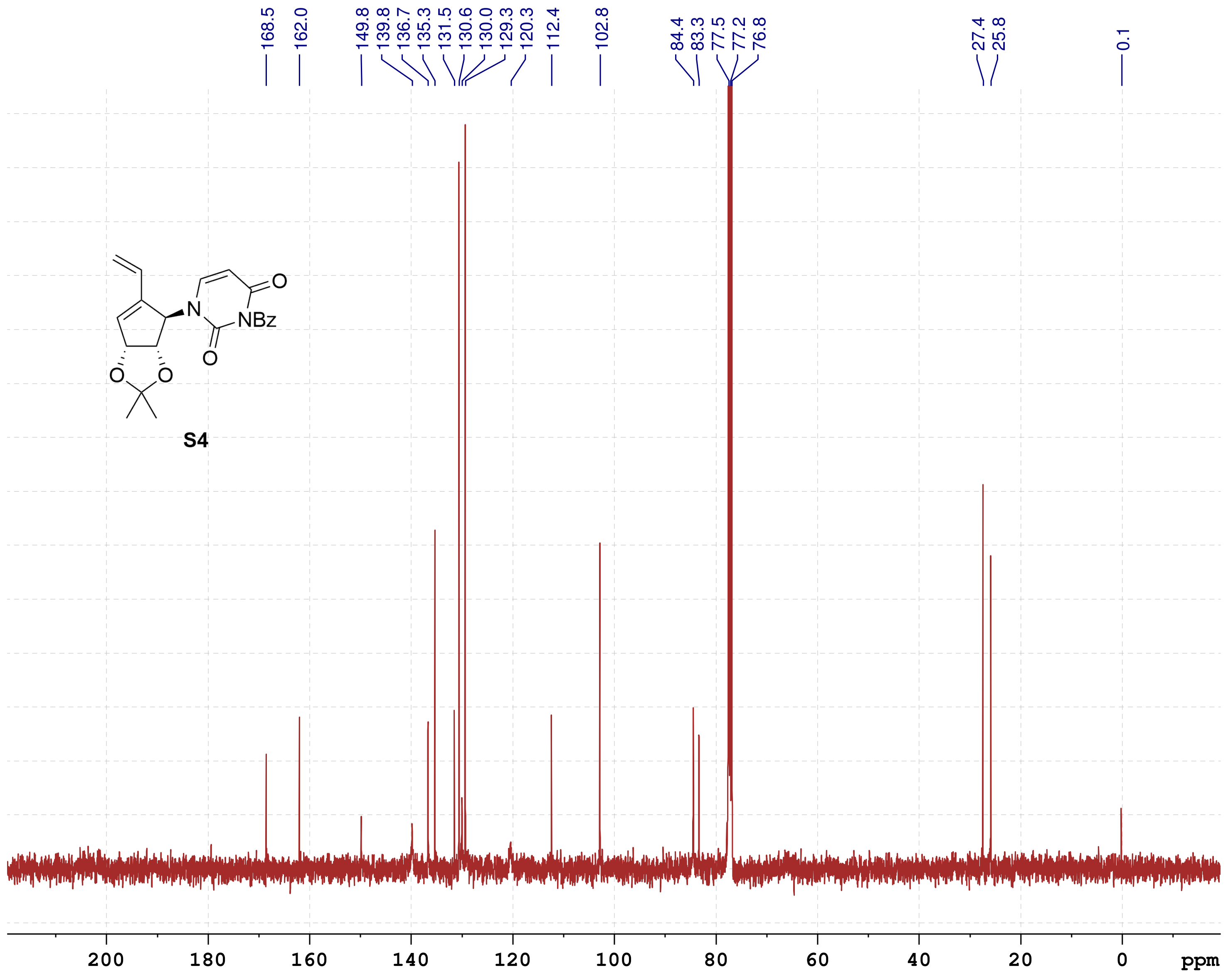

S4

#### NMR-Spectra for Compound 15

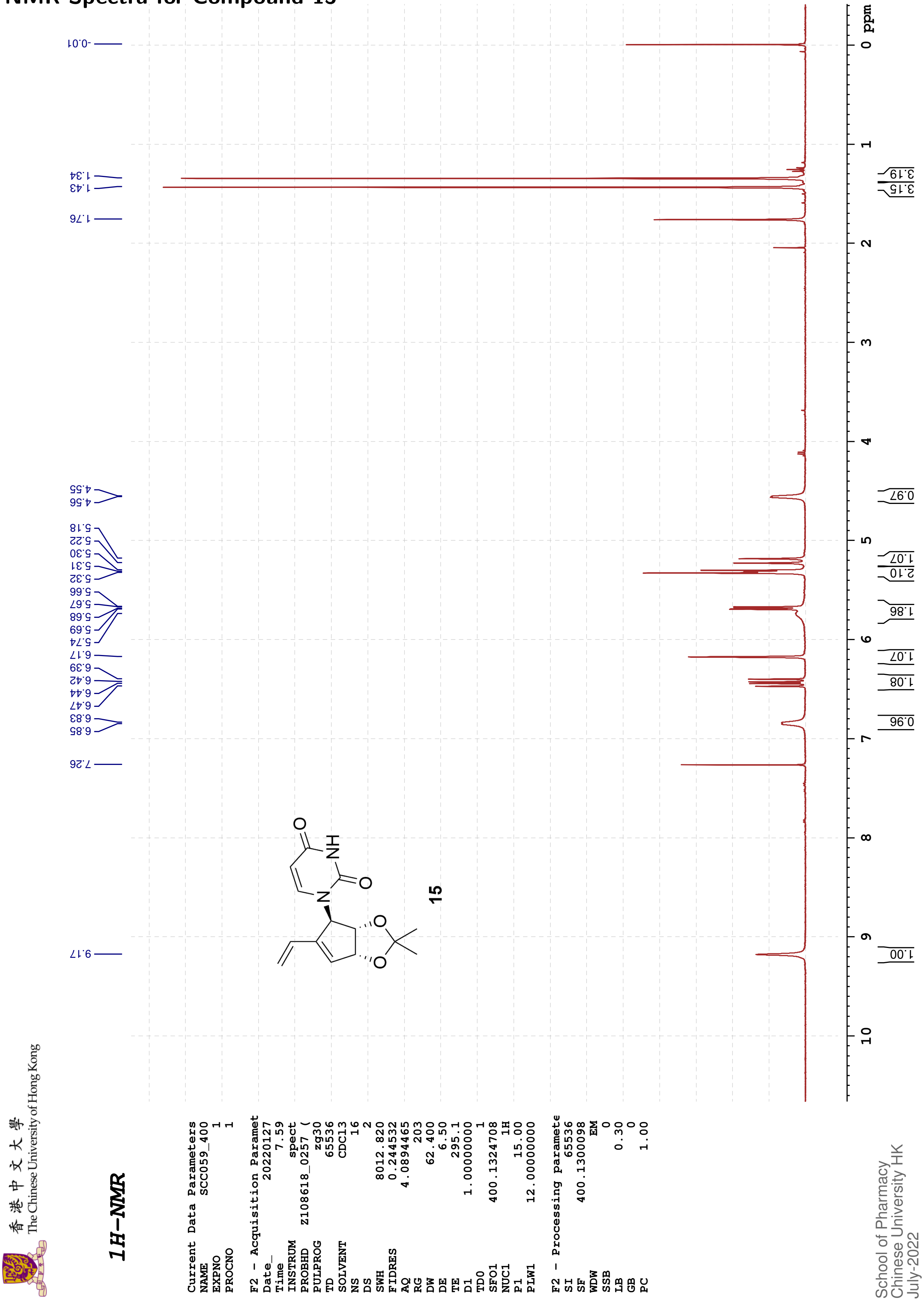

香港中文大學
The Chinese University of Hong Kong

*13C{1H}-NMR*

| | |
|---|---|
| Current Data Parameters | |
| NAME | SCC059_400 |
| EXPNO | 98 |
| PROCNO | 1 |
| F2 - Acquisition Paramet | |
| Date_ | 20220127 |
| Time | 8.39 |
| INSTRUM | spect |
| PROBHD | Z108618_0257 ( |
| PULPROG | zgpg30 |
| TD | 65536 |
| SOLVENT | CDCl3 |
| NS | 361 |
| DS | 4 |
| SWH | 24038.461 |
| FIDRES | 0.733596 |
| AQ | 1.3631488 |
| RG | 203 |
| DW | 20.800 |
| DE | 6.50 |
| TE | 295.6 |
| D1 | 2.00000000 |
| D11 | 0.03000000 |
| TD0 | 1 |
| SFO1 | 100.6228298 |
| NUC1 | 13C |
| P1 | 10.00 |
| PLW1 | 47.00000000 |
| SFO2 | 400.1316005 |
| NUC2 | 1H |
| CPDPRG[2 | waltz16 |
| PCPD2 | 90.00 |
| PLW2 | 12.00000000 |
| PLW12 | 0.33333001 |
| PLW13 | 0.16766000 |
| F2 - Processing paramete | |
| SI | 32768 |
| SF | 100.6127573 |
| WDW | EM |
| SSB | 0 |
| LB | 3.00 |
| GB | 0 |
| PC | 1.40 |

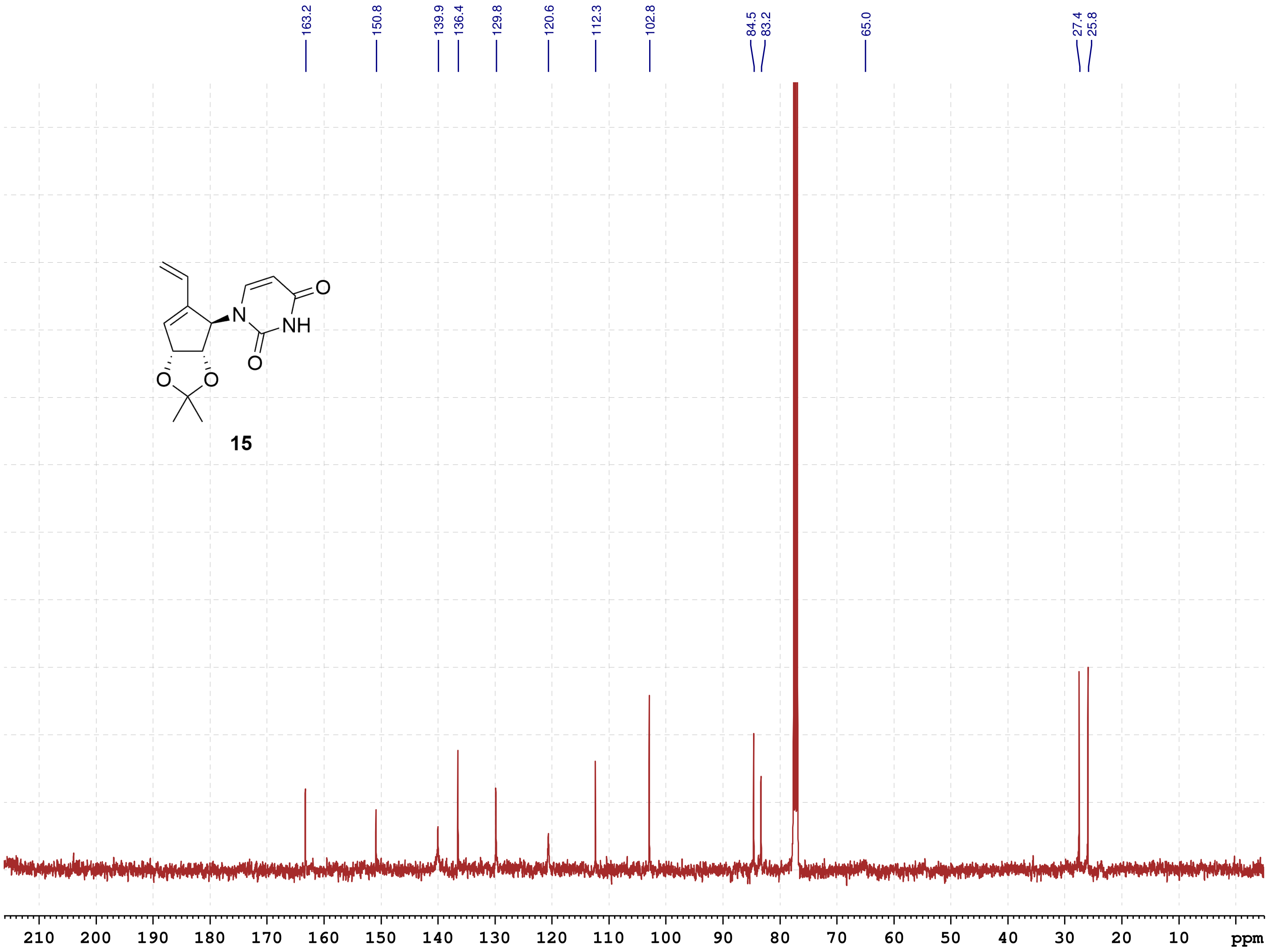

School of Pharmacy
Chinese University HK
July-2022

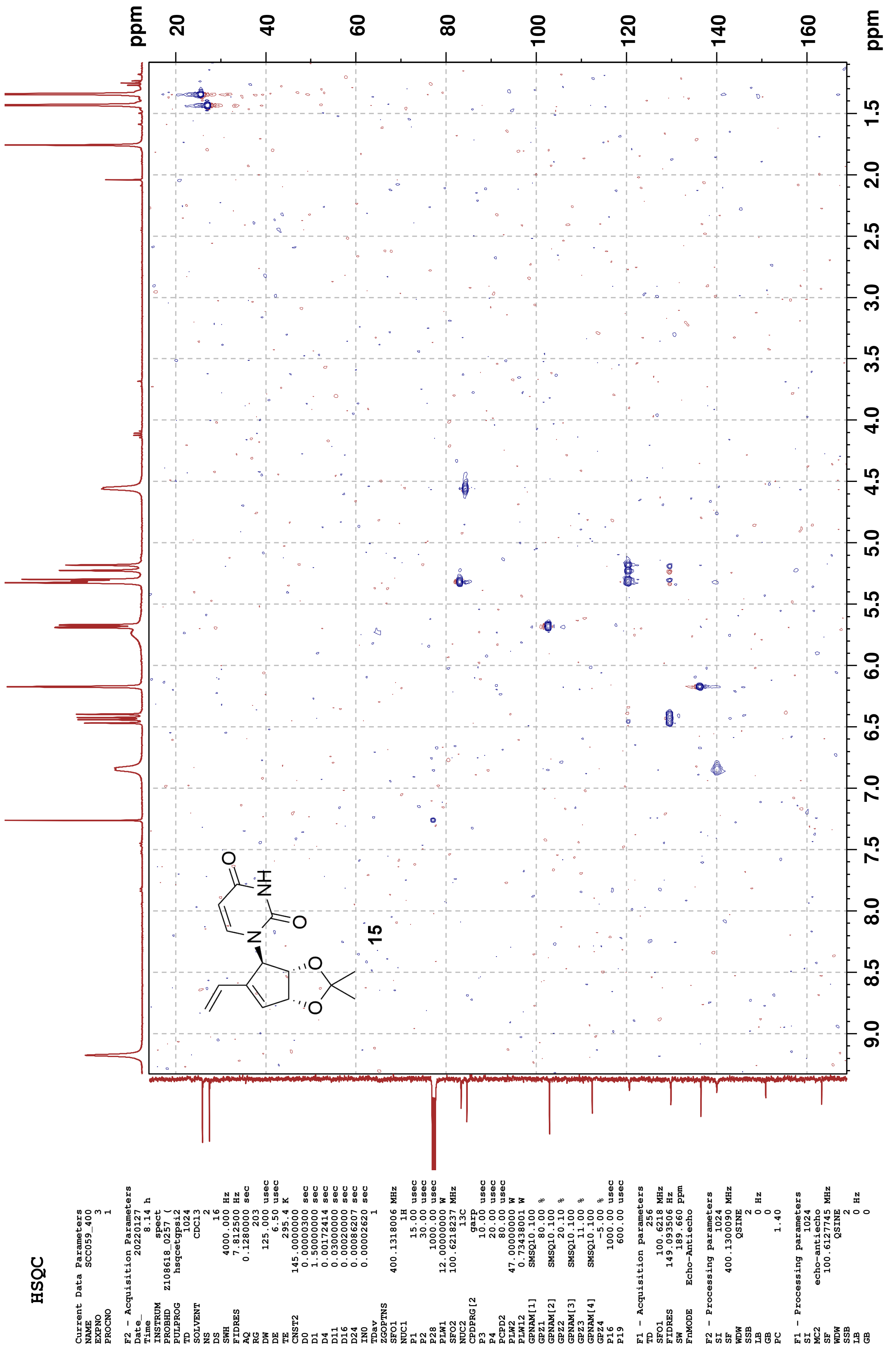
HSQC
Current Data Parameters
NAME SCC059_400
EXPNO 3
PROCNO 1
F2 - Acquisition Parameters
Date_ 20220127
Time 8.14 h
INSTRUM spect
PROBHD Z108618_0257 (
PULPROG hsqcetgpsi2
TD 1024
SOLVENT CDCl3
NS 2
DS 16
SWH 4000.000 Hz
FIDRES 7.812500 Hz
AQ 0.1280000 sec
RG 203
DW 125.000 usec
DE 6.50 usec
TE 295.4 K
CNST2 145.0000000
D0 0.00000300 sec
D1 1.50000000 sec
D4 0.00172414 sec
D11 0.03000000 sec
D16 0.00020000 sec
D24 0.00086207 sec
IN0 0.00002620 sec
TDav 1
ZGOPTNS
SFO1 400.1318006 MHz
NUC1 1H
P1 15.00 usec
P2 30.00 usec
P28 1000.00 usec
PLW1 12.00000000 W
SFO2 100.6218237 MHz
NUC2 13C
CPDPRG[2 garp
P3 10.00 usec
P4 20.00 usec
PCPD2 80.00 usec
PLW2 47.00000000 W
PLW12 0.73438001 W
GPNAM[1] SMSQ10.100
GPZ1 80.00 %
GPNAM[2] SMSQ10.100
GPZ2 20.10 %
GPNAM[3] SMSQ10.100
GPZ3 11.00 %
GPNAM[4] SMSQ10.100
GPZ4 -5.00 %
P16 1000.00 usec
P19 600.00 usec
F1 - Acquisition parameters
TD 256
SFO1 100.6218 MHz
FIDRES 149.093506 Hz
SW 189.660 ppm
FnMODE Echo-Antiecho
F2 - Processing parameters
SI 1024
SF 400.1300090 MHz
WDW QSINE
SSB 2
LB 0 Hz
GB 0
PC 1.40
F1 - Processing parameters
SI 1024
MC2 echo-antiecho
SF 100.6127745 MHz
WDW QSINE
SSB 2
LB 0 Hz
GB 0
15
ppm
20
40
60
80
100
120
140
160
ppm
9.0
8.5
8.0
7.5
7.0
6.5
6.0
5.5
5.0
4.5
4.0
3.5
3.0
2.5
2.0
1.5

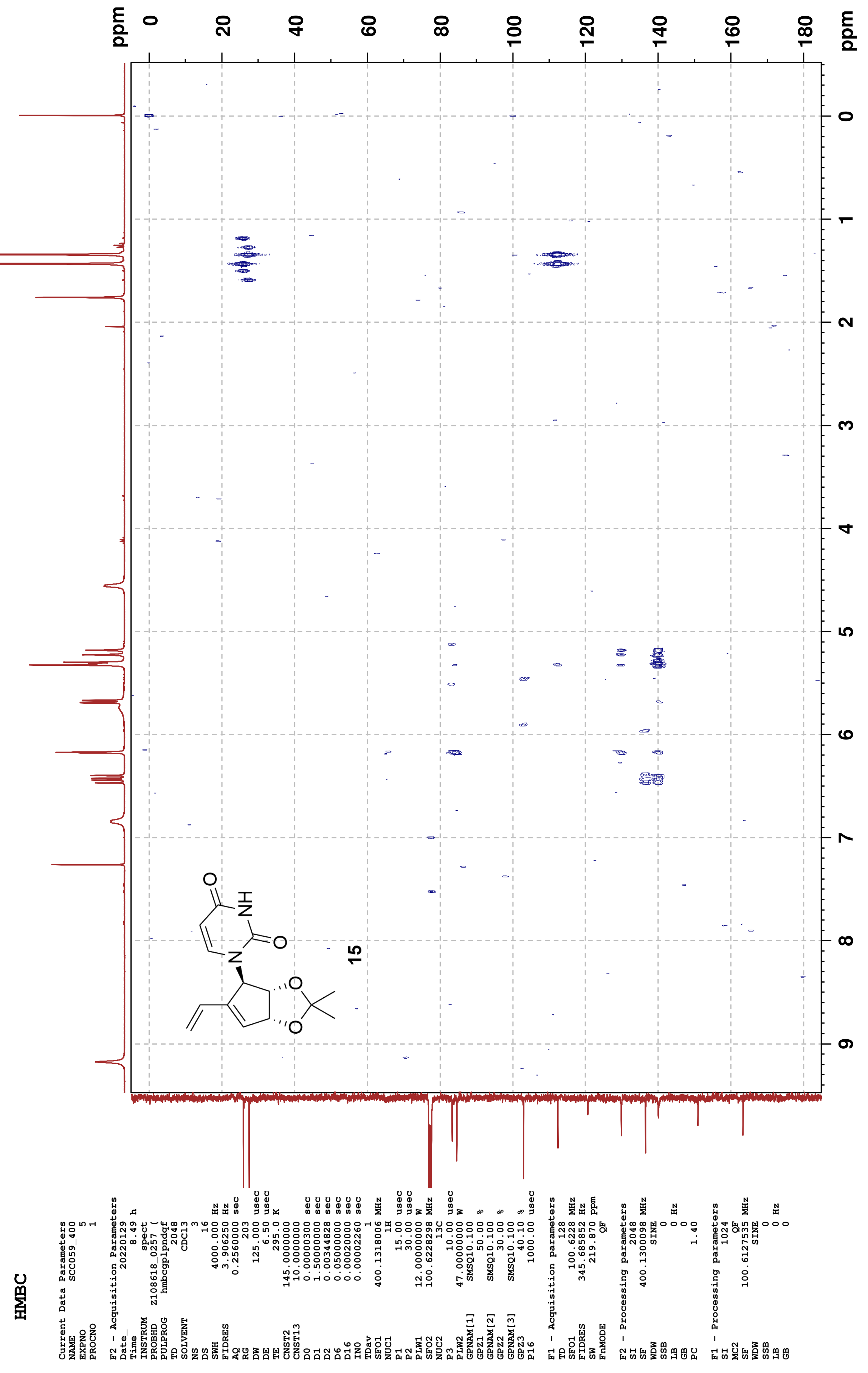

HMBC
15
ppm
Current Data Parameters
NAME SCC059_400
EXPNO 5
PROCNO 1
F2 - Acquisition Parameters
Date_ 20220129
Time 8.49 h
INSTRUM spect
PROBHD Z108618_0257 (
PULPROG hmbcgplpndqf
TD 2048
SOLVENT CDCl3
NS 3
DS 16
SWH 4000.000 Hz
FIDRES 3.906250 Hz
AQ 0.2560000 sec
RG 203
DW 125.000 usec
DE 6.50 usec
TE 295.0 K
CNST2 145.0000000
CNST13 10.0000000
D0 0.00000300 sec
D1 1.50000000 sec
D2 0.00344828 sec
D6 0.05000000 sec
D16 0.00020000 sec
IN0 0.00002260 sec
TDav 1
SFO1 400.1318006 MHz
NUC1 1H
P1 15.00 usec
P2 30.00 usec
PLW1 12.00000000 W
SFO2 100.6228298 MHz
NUC2 13C
P3 10.00 usec
PLW2 47.00000000 W
GPNAM[1] SMSQ10.100
GPZ1 50.00 %
GPNAM[2] SMSQ10.100
GPZ2 30.00 %
GPNAM[3] SMSQ10.100
GPZ3 40.10 %
P16 1000.00 usec
F1 - Acquisition parameters
TD 128
SFO1 100.6228 MHz
FIDRES 345.685852 Hz
SW 219.870 ppm
FnMODE QF
F2 - Processing parameters
SI 2048
SF 400.1300098 MHz
WDW SINE
SSB 0
LB 0 Hz
GB 0
PC 1.40
F1 - Processing parameters
SI 1024
MC2 QF
SF 100.6127535 MHz
WDW SINE
SSB 0
LB 0 Hz
GB 0

#### NMR-Spectra for Compound 16

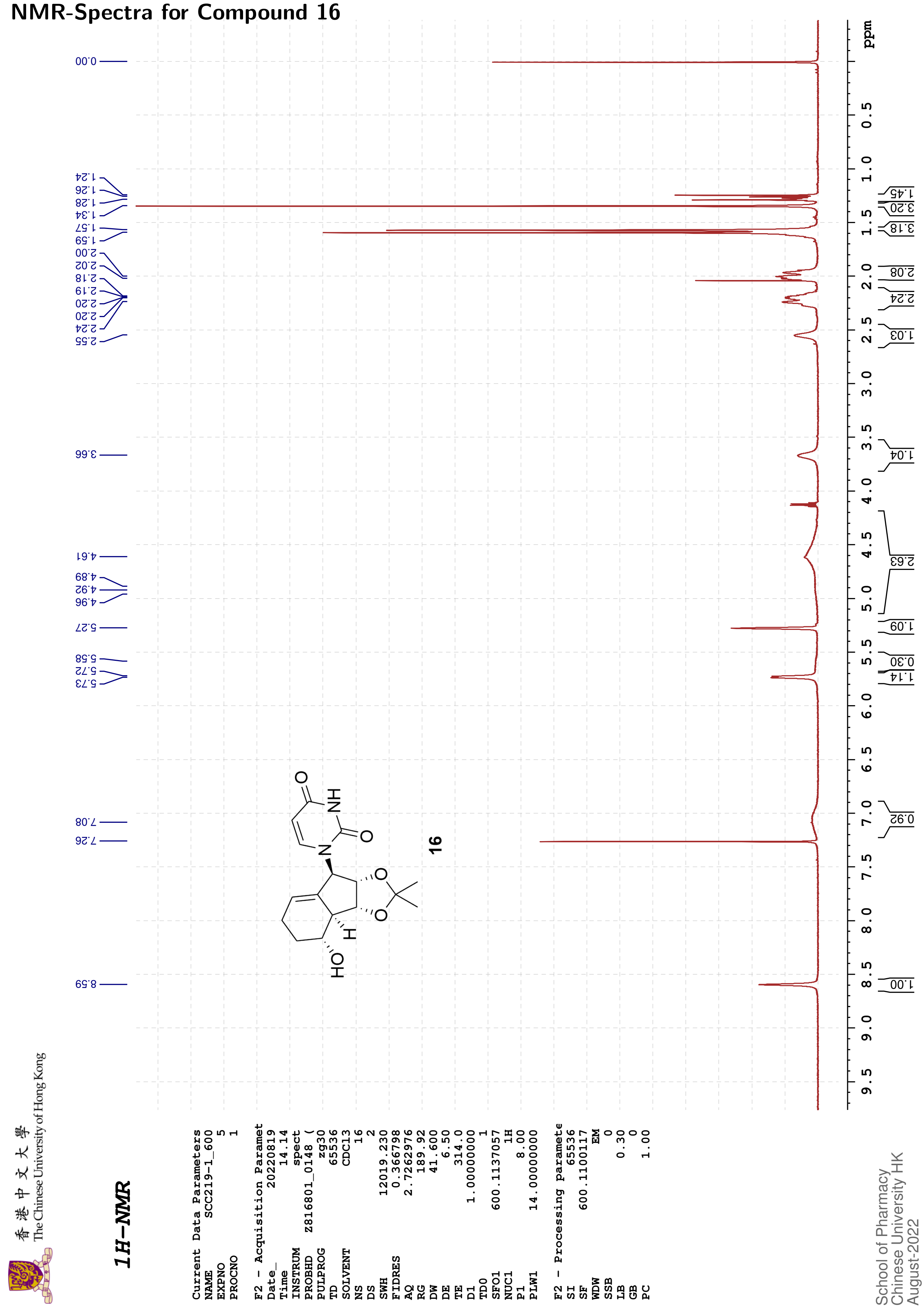

### HSQC

```
Current Data Parameters
NAME        SCC219-1_600
EXPNO                  6
PROCNO                 1

F2 - Acquisition Parameters
Date_           20220819
Time               14.24 h
INSTRUM            spect
PROBHD    Z816801_0148 (
PULPROG      hsqcetgpsi2
TD                  1024
SOLVENT            CDCl3
NS                     2
DS                    16
SWH             7812.500 Hz
FIDRES         15.258789 Hz
AQ             0.0655360 sec
RG                189.92
DW                64.000 usec
DE                  6.50 usec
TE                 316.8 K
CNST2        145.0000000
D0            0.00000300 sec
D1            1.50000000 sec
D4            0.00172414 sec
D11           0.03000000 sec
D16           0.00020000 sec
D24           0.00086207 sec
IN0           0.00001660 sec
TDav                   1
ZGOPTNS
SFO1         600.1137057 MHz
NUC1                  1H
P1                  8.00 usec
P2                 16.00 usec
P28              1000.00 usec
PLW1         14.00000000 W
SFO2         150.9128693 MHz
NUC2                 13C
CPDPRG[2            garp
P3                 12.00 usec
P4                 24.00 usec
PCPD2              60.00 usec
PLW2        183.00000000 W
PLW12         7.32000017 W
GPNAM[1]      SMSQ10.100
GPZ1               80.00 %
GPNAM[2]      SMSQ10.100
GPZ2               20.10 %
GPNAM[3]      SMSQ10.100
GPZ3               11.00 %
GPNAM[4]      SMSQ10.100
GPZ4               -5.00 %
P16              1000.00 usec
P19               600.00 usec

F1 - Acquisition parameters
TD                   256
SFO1            150.9129 MHz
FIDRES        235.316269 Hz
SW               199.589 ppm
FnMODE      Echo-Antiecho

F2 - Processing parameters
SI                  1024
SF           600.1100101 MHz
WDW                QSINE
SSB                    2
LB                     0 Hz
GB                     0
PC                  1.40

F1 - Processing parameters
SI                  1024
MC2         echo-antiecho
SF           150.8977841 MHz
WDW                QSINE
SSB                    2
LB                     0 Hz
GB                     0
```

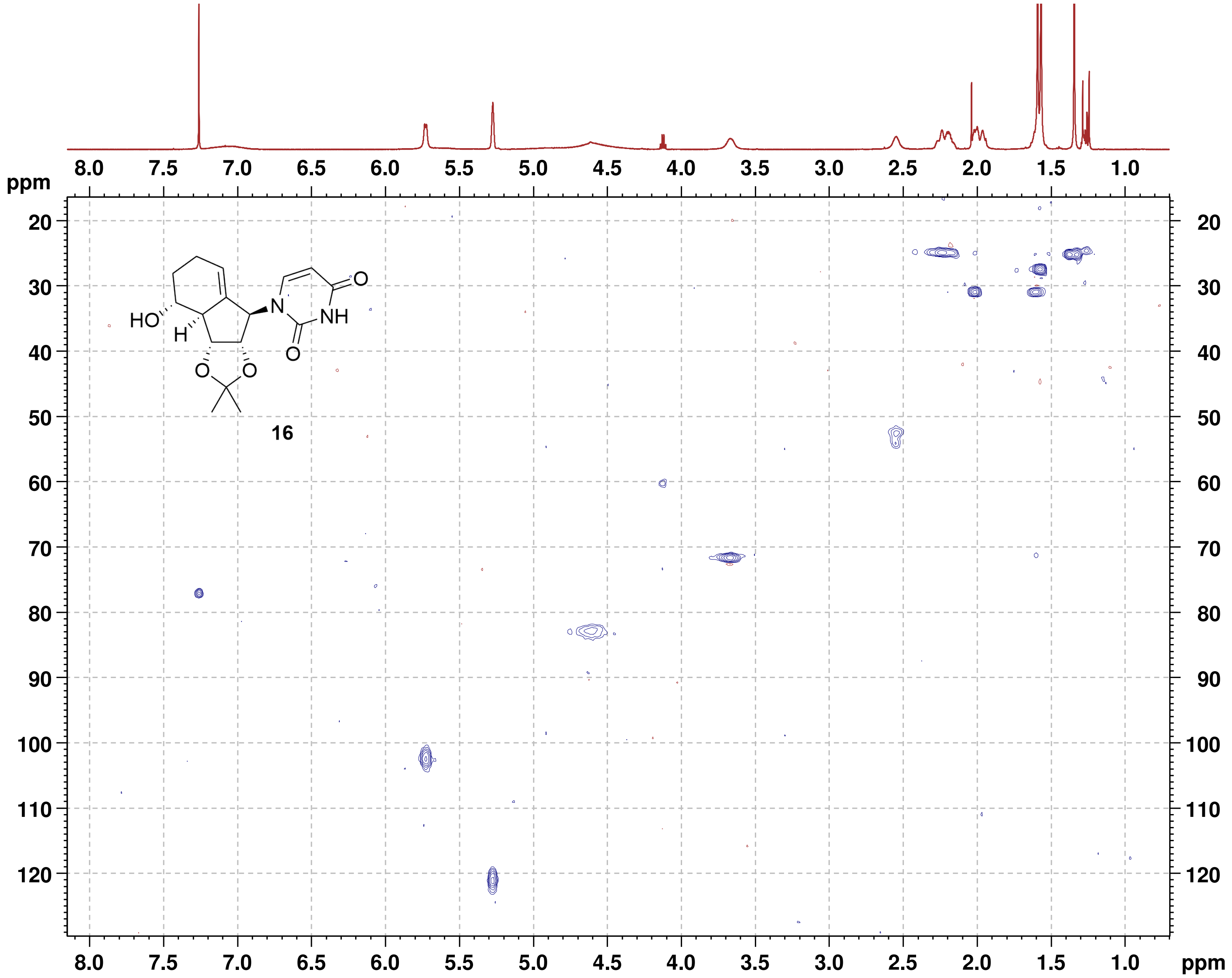

### HMBC

```
Current Data Parameters
NAME          SCC219-1_600
EXPNO                    8
PROCNO                   1

F2 - Acquisition Parameters
Date_             20220819
Time                 14.42 h
INSTRUM              spect
PROBHD   Z816801_0148 (
PULPROG      hmbcgplpndqf
TD                    2048
SOLVENT              CDCl3
NS                       8
DS                      16
SWH               7812.500 Hz
FIDRES            7.629395 Hz
AQ               0.1310720 sec
RG                  189.92
DW                  64.000 usec
DE                    6.50 usec
TE                   315.0 K
CNST2          145.0000000
CNST13          10.0000000
D0              0.00000300 sec
D1              1.50000000 sec
D2              0.00344828 sec
D6              0.05000000 sec
D16             0.00020000 sec
IN0             0.00001510 sec
TDav                     1
SFO1           600.1137807 MHz
NUC1                    1H
P1                    8.00 usec
P2                   16.00 usec
PLW1           14.00000000 W
SFO2           150.9128693 MHz
NUC2                   13C
P3                   12.00 usec
PLW2          183.00000000 W
GPNAM[1]        SMSQ10.100
GPZ1                 50.00 %
GPNAM[2]        SMSQ10.100
GPZ2                 30.00 %
GPNAM[3]        SMSQ10.100
GPZ3                 40.10 %
P16                1000.00 usec

F1 - Acquisition parameters
TD                     128
SFO1              150.9129 MHz
FIDRES          517.384094 Hz
SW                 219.415 ppm
FnMODE                  QF

F2 - Processing parameters
SI                    2048
SF             600.1100083 MHz
WDW                   SINE
SSB                      0
LB                       0 Hz
GB                       0
PC                    1.40

F1 - Processing parameters
SI                    1024
MC2                     QF
SF             150.8977757 MHz
WDW                   SINE
SSB                      0
LB                       0 Hz
GB                       0
```

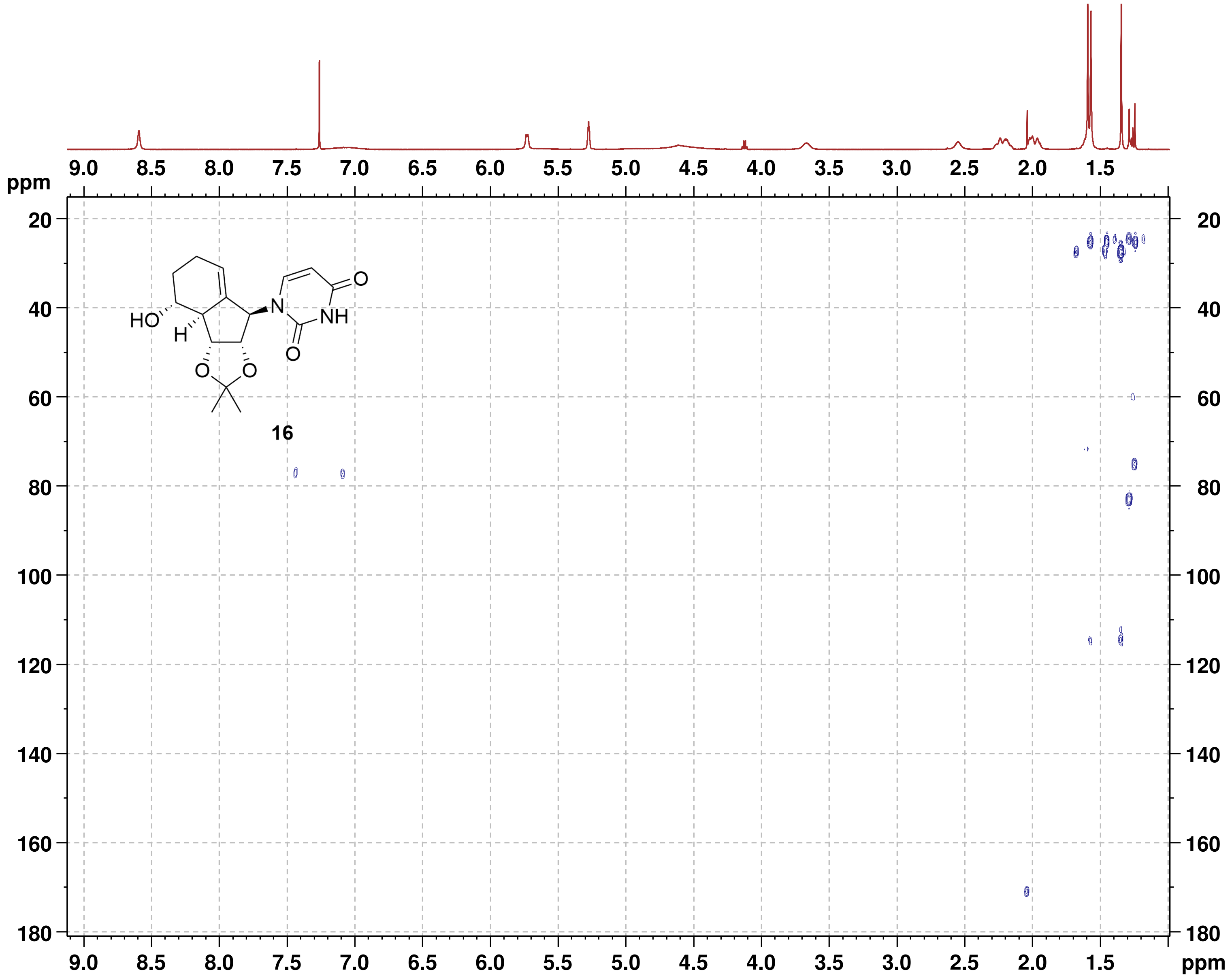

#### NMR-Spectra for Compound 17

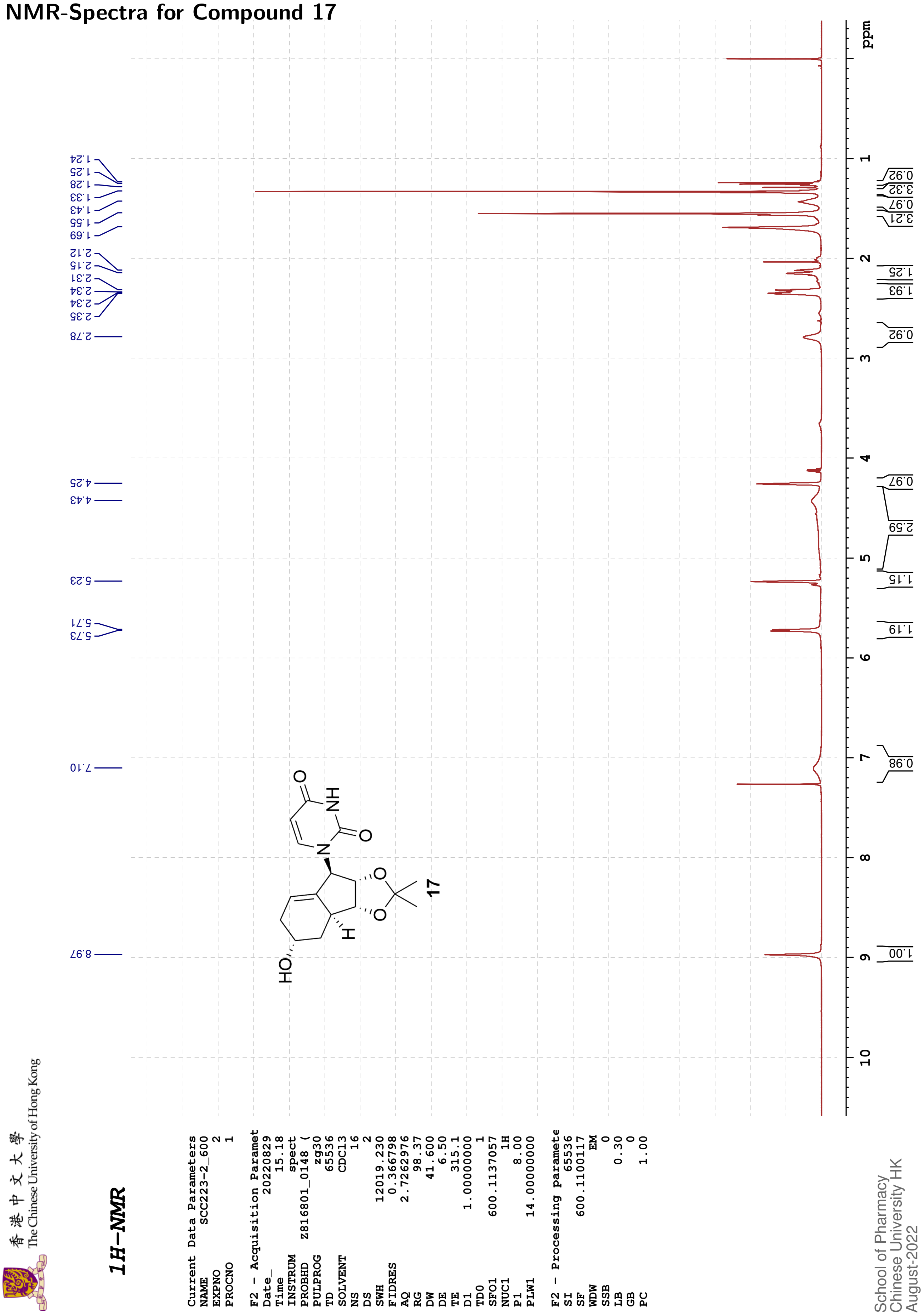

### 13C{1H}-NMR

```
Current Data Parameters
NAME        SCC223-2_600
EXPNO                  3
PROCNO                 1

F2 - Acquisition Paramet
Date_           20220829
Time               15.27
INSTRUM            spect
PROBHD    Z816801_0148 (
PULPROG          zgpg30
TD                 65536
SOLVENT            CDCl3
NS                   147
DS                     4
SWH            36231.883
FIDRES          1.105709
AQ             0.9043968
RG                 98.37
DW                13.800
DE                  6.50
TE                 315.0
D1            2.00000000
D11           0.03000000
TD0                    1
SFO1         150.9128693
NUC1                 13C
P1                 12.00
PLW1        183.00000000
SFO2         600.1124004
NUC2                  1H
CPDPRG[2         waltz16
PCPD2              70.00
PLW2         14.00000000
PLW12         0.18286000
PLW13         0.09197600

F2 - Processing paramete
SI                 32768
SF           150.8977520
WDW                   EM
SSB                    0
LB                  1.00
GB                     0
PC                  1.40
```

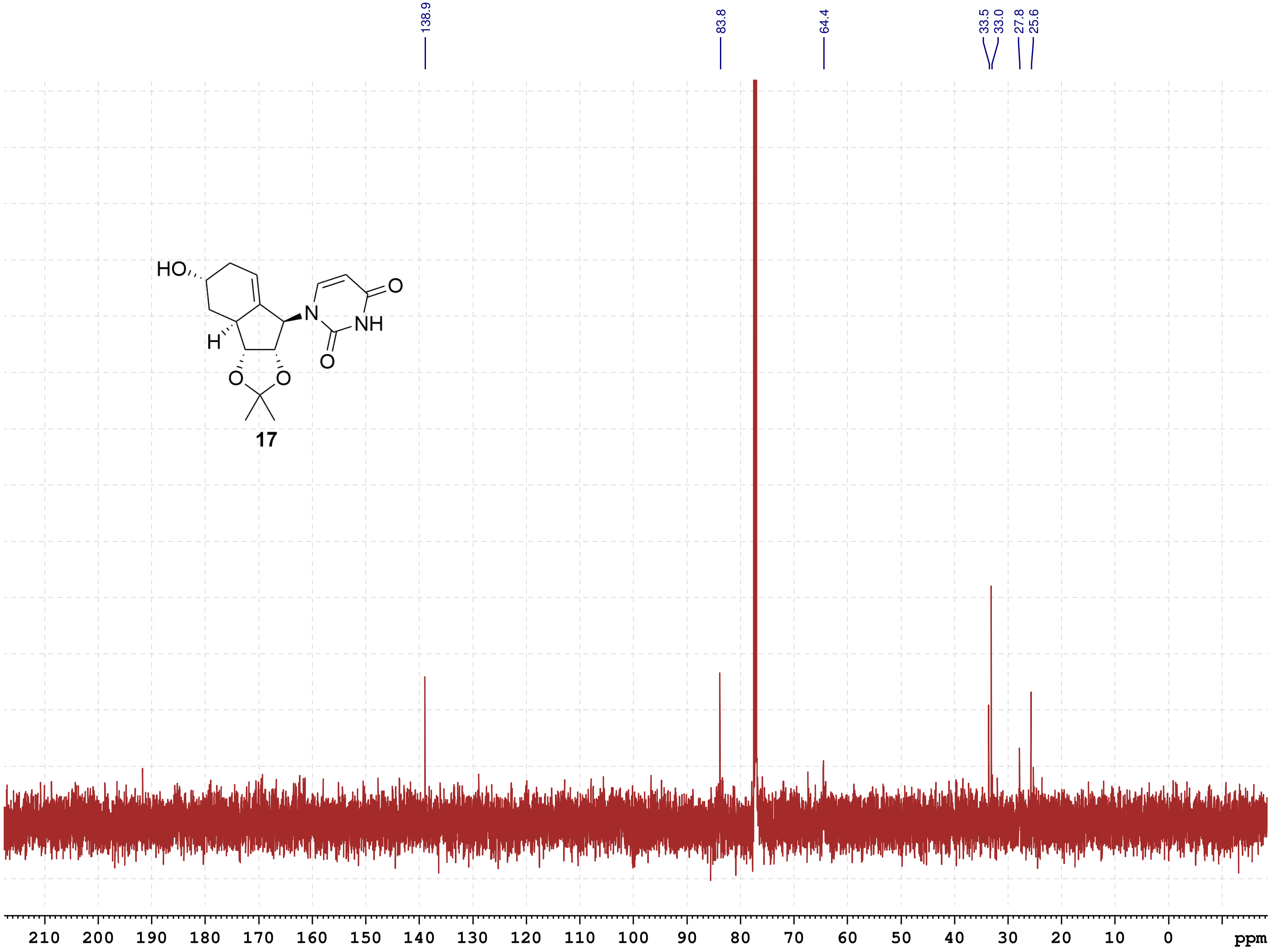

School of Pharmacy
Chinese University HK
August-2022

### HSQC

```
Current Data Parameters
NAME          SCC223-2_600
EXPNO                    5
PROCNO                   1

F2 - Acquisition Parameters
Date_             20220829
Time                 15.30 h
INSTRUM              spect
PROBHD    Z816801_0148 (
PULPROG      hsqcetgpsi2
TD                    1024
SOLVENT              CDCl3
NS                       4
DS                      16
SWH               7812.500 Hz
FIDRES           15.258789 Hz
AQ               0.0655360 sec
RG                  189.92
DW                  64.000 usec
DE                    6.50 usec
TE                   315.2 K
CNST2          145.0000000
D0              0.00000300 sec
D1              1.50000000 sec
D4              0.00172414 sec
D11             0.03000000 sec
D16             0.00020000 sec
D24             0.00086207 sec
IN0             0.00001660 sec
TDav                     1
ZGOPTNS
SFO1           600.1137057 MHz
NUC1                    1H
P1                    8.00 usec
P2                   16.00 usec
P28                1000.00 usec
PLW1           14.00000000 W
SFO2           150.9128693 MHz
NUC2                   13C
CPDPRG[2              garp
P3                   12.00 usec
P4                   24.00 usec
PCPD2                60.00 usec
PLW2          183.00000000 W
PLW12           7.32000017 W
GPNAM[1]        SMSQ10.100
GPZ1                 80.00 %
GPNAM[2]        SMSQ10.100
GPZ2                 20.10 %
GPNAM[3]        SMSQ10.100
GPZ3                 11.00 %
GPNAM[4]        SMSQ10.100
GPZ4                 -5.00 %
P16                1000.00 usec
P19                 600.00 usec

F1 - Acquisition parameters
TD                     128
SFO1              150.9129 MHz
FIDRES          470.632538 Hz
SW                 199.589 ppm
FnMODE        Echo-Antiecho

F2 - Processing parameters
SI                    1024
SF             600.1100094 MHz
WDW                  QSINE
SSB                      2
LB                       0 Hz
GB                       0
PC                    1.40

F1 - Processing parameters
SI                    1024
MC2            echo-antiecho
SF             150.8977724 MHz
WDW                  QSINE
SSB                      2
LB                       0 Hz
GB                       0
```

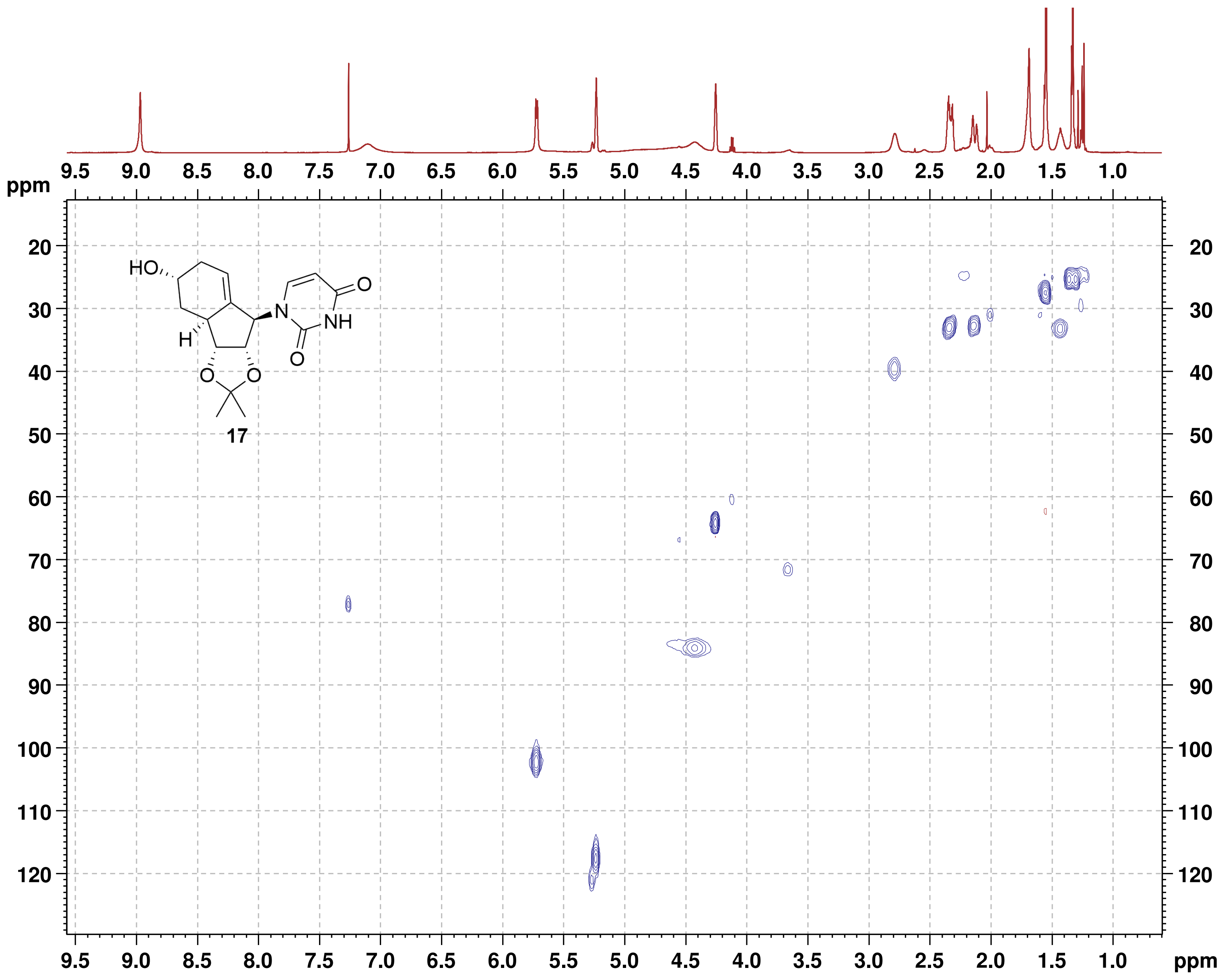

### HMBC

```
Current Data Parameters
NAME        SCC223-2_600
EXPNO                  6
PROCNO                 1

F2 - Acquisition Parameters
Date_           20220829
Time               15.45 h
INSTRUM            spect
PROBHD   Z816801_0148 (
PULPROG    hmbcgplpndqf
TD                  2048
SOLVENT            CDCl3
NS                     6
DS                    16
SWH             6009.615 Hz
FIDRES          5.868765 Hz
AQ             0.1703936 sec
RG                189.92
DW                83.200 usec
DE                  6.50 usec
TE                 315.0 K
CNST2        145.0000000
CNST13        10.0000000
D0            0.00000300 sec
D1            1.50000000 sec
D2            0.00344828 sec
D6            0.05000000 sec
D16           0.00020000 sec
IN0           0.00001510 sec
TDav                   1
SFO1        600.1127005 MHz
NUC1                  1H
P1                  8.00 usec
P2                 16.00 usec
PLW1         14.00000000 W
SFO2        150.9128693 MHz
NUC2                 13C
P3                 12.00 usec
PLW2        183.00000000 W
GPNAM[1]      SMSQ10.100
GPZ1               50.00 %
GPNAM[2]      SMSQ10.100
GPZ2               30.00 %
GPNAM[3]      SMSQ10.100
GPZ3               40.10 %
P16              1000.00 usec

F1 - Acquisition parameters
TD                   128
SFO1            150.9129 MHz
FIDRES        517.384094 Hz
SW               219.415 ppm
FnMODE                QF

F2 - Processing parameters
SI                  2048
SF           600.1100073 MHz
WDW                 SINE
SSB                    0
LB                     0 Hz
GB                     0
PC                  1.40

F1 - Processing parameters
SI                  1024
MC2                   QF
SF           150.8977736 MHz
WDW                 SINE
SSB                    0
LB                     0 Hz
GB                     0
```

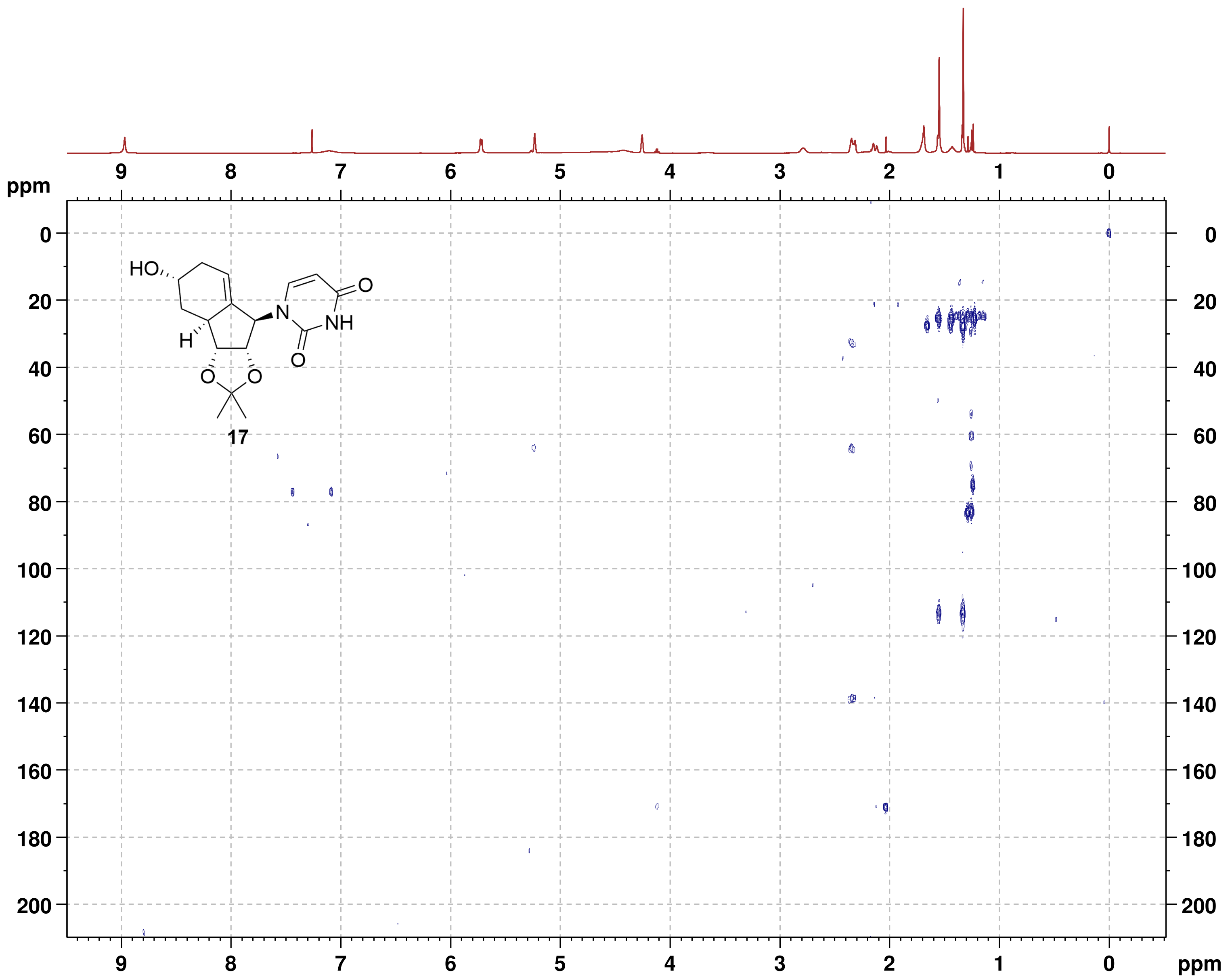

#### NMR-Spectra for Compound 18

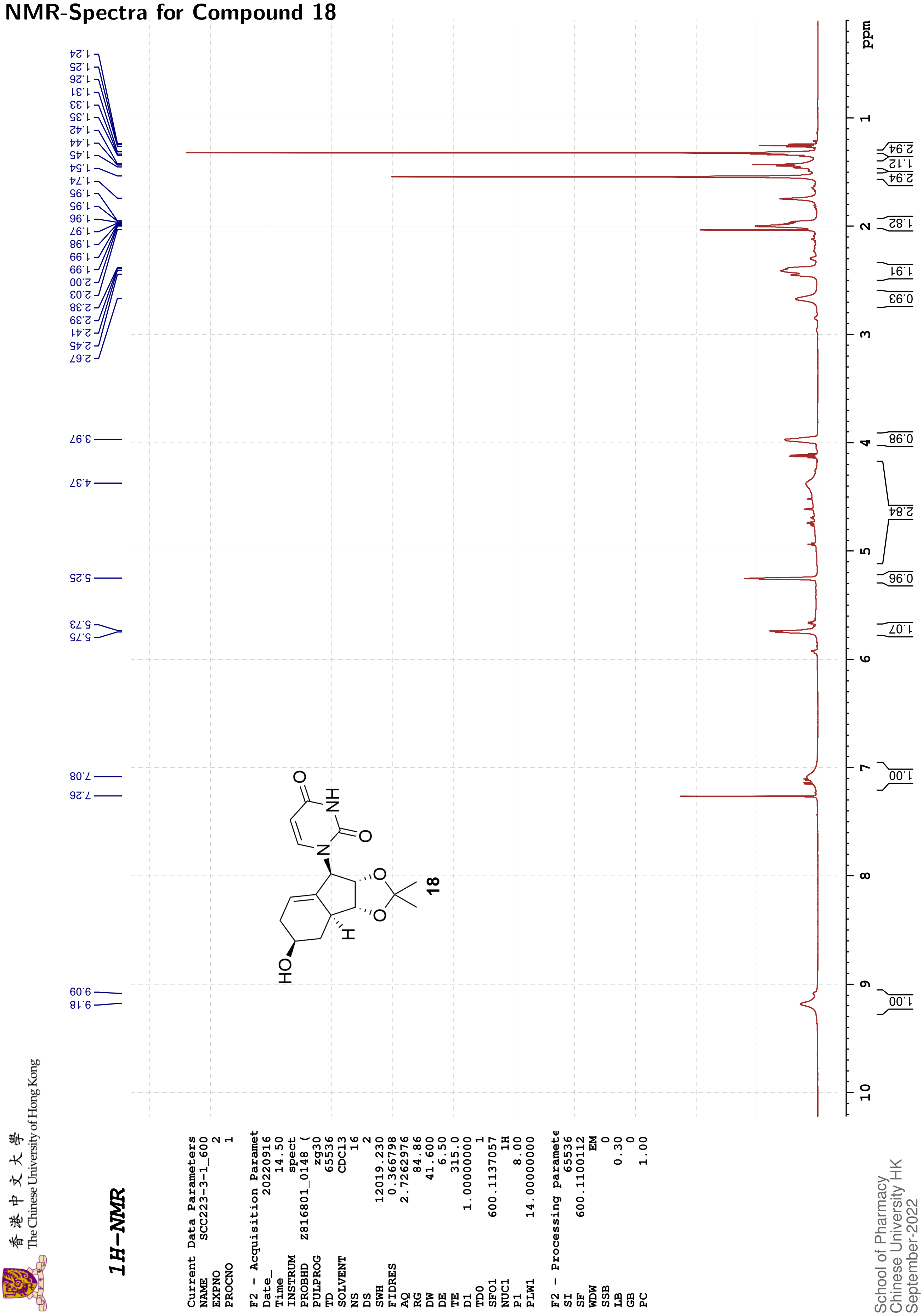

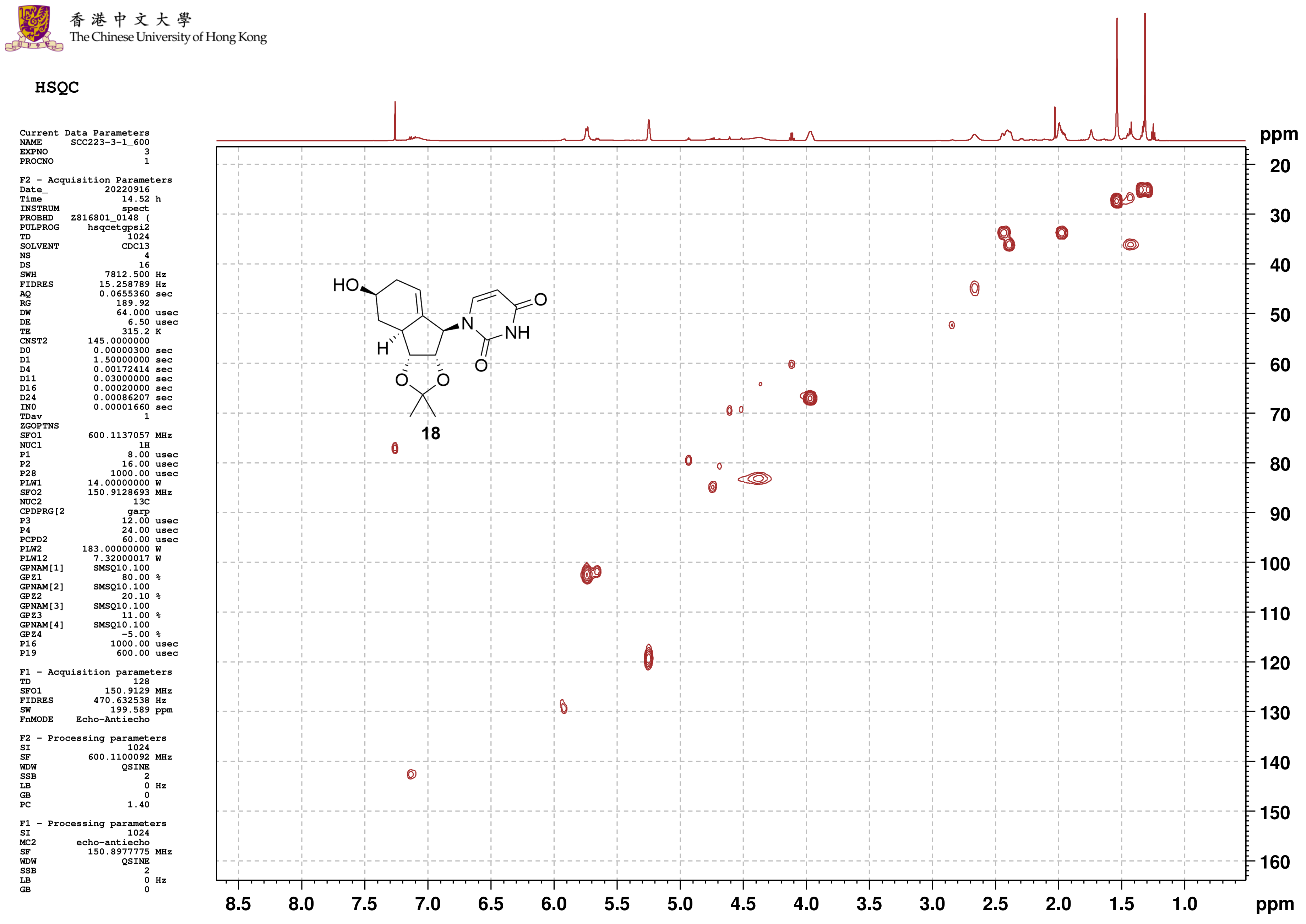
香港中文大學
The Chinese University of Hong Kong
HSQC
Current Data Parameters
NAME SCC223-3-1_600
EXPNO 3
PROCNO 1
F2 - Acquisition Parameters
Date_ 20220916
Time 14.52 h
INSTRUM spect
PROBHD Z816801_0148 (
PULPROG hsqcetgpsi2
TD 1024
SOLVENT CDCl3
NS 4
DS 16
SWH 7812.500 Hz
FIDRES 15.258789 Hz
AQ 0.0655360 sec
RG 189.92
DW 64.000 usec
DE 6.50 usec
TE 315.2 K
CNST2 145.0000000
D0 0.00000300 sec
D1 1.50000000 sec
D4 0.00172414 sec
D11 0.03000000 sec
D16 0.00020000 sec
D24 0.00086207 sec
IN0 0.00001660 sec
TDav 1
ZGOPTNS
SFO1 600.1137057 MHz
NUC1 1H
P1 8.00 usec
P2 16.00 usec
P28 1000.00 usec
PLW1 14.00000000 W
SFO2 150.9128693 MHz
NUC2 13C
CPDPRG[2 garp
P3 12.00 usec
P4 24.00 usec
PCPD2 60.00 usec
PLW2 183.00000000 W
PLW12 7.32000017 W
GPNAM[1] SMSQ10.100
GPZ1 80.00 %
GPNAM[2] SMSQ10.100
GPZ2 20.10 %
GPNAM[3] SMSQ10.100
GPZ3 11.00 %
GPNAM[4] SMSQ10.100
GPZ4 -5.00 %
P16 1000.00 usec
P19 600.00 usec
F1 - Acquisition parameters
TD 128
SFO1 150.9129 MHz
FIDRES 470.632538 Hz
SW 199.589 ppm
FnMODE Echo-Antiecho
F2 - Processing parameters
SI 1024
SF 600.1100092 MHz
WDW QSINE
SSB 2
LB 0 Hz
GB 0
PC 1.40
F1 - Processing parameters
SI 1024
MC2 echo-antiecho
SF 150.8977775 MHz
WDW QSINE
SSB 2
LB 0 Hz
GB 0
HO
H
N
NH
O
O
O
O
18
ppm
20
30
40
50
60
70
80
90
100
110
120
130
140
150
160
8.5
8.0
7.5
7.0
6.5
6.0
5.5
5.0
4.5
4.0
3.5
3.0
2.5
2.0
1.5
1.0
ppm

香港中文大學
The Chinese University of Hong Kong

#### HMBC

```
Current Data Parameters
NAME      SCC223-3-1_600
EXPNO                  4
PROCNO                 1

F2 - Acquisition Parameters
Date_           20220916
Time               15.06 h
INSTRUM            spect
PROBHD   Z816801_0148 (
PULPROG     hmbcgplpndqf
TD                  2048
SOLVENT            CDCl3
NS                     6
DS                    16
SWH             6009.615 Hz
FIDRES          5.868765 Hz
AQ             0.1703936 sec
RG                189.92
DW                83.200 usec
DE                  6.50 usec
TE                 314.9 K
CNST2        145.0000000
CNST13        10.0000000
D0            0.00000300 sec
D1            1.50000000 sec
D2            0.00344828 sec
D6            0.05000000 sec
D16           0.00020000 sec
IN0           0.00001510 sec
TDav                   1
SFO1         600.1127005 MHz
NUC1                  1H
P1                  8.00 usec
P2                 16.00 usec
PLW1         14.00000000 W
SFO2         150.9128693 MHz
NUC2                 13C
P3                 12.00 usec
PLW2        183.00000000 W
GPNAM[1]       SMSQ10.100
GPZ1               50.00 %
GPNAM[2]       SMSQ10.100
GPZ2               30.00 %
GPNAM[3]       SMSQ10.100
GPZ3               40.10 %
P16              1000.00 usec

F1 - Acquisition parameters
TD                   128
SFO1            150.9129 MHz
FIDRES        517.384094 Hz
SW               219.415 ppm
FnMODE                QF

F2 - Processing parameters
SI                  2048
SF           600.1100081 MHz
WDW                 SINE
SSB                    0
LB                     0 Hz
GB                     0
PC                  1.40

F1 - Processing parameters
SI                  1024
MC2                   QF
SF           150.8977883 MHz
WDW                 SINE
SSB                    0
LB                     0 Hz
GB                     0
```

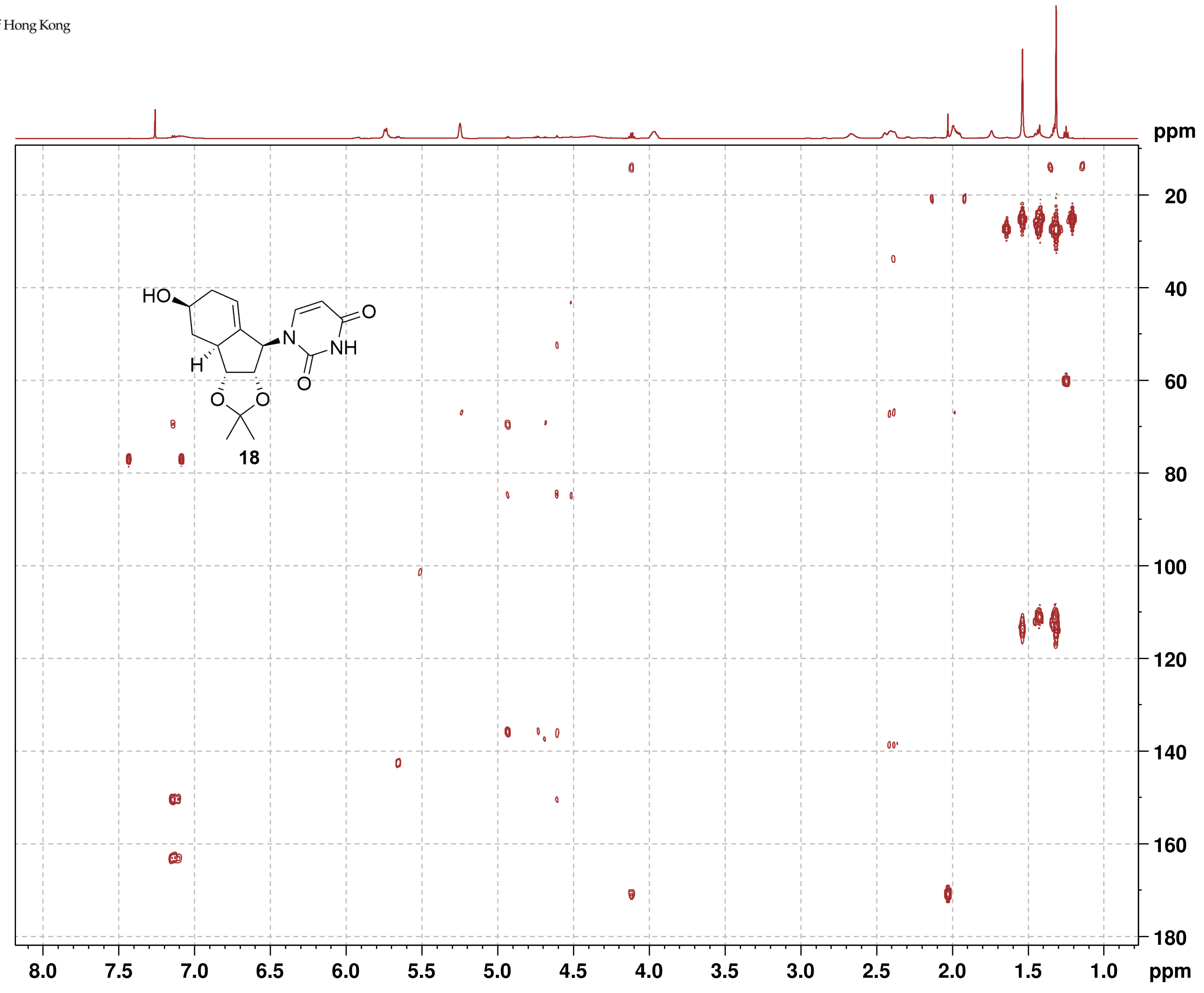

#### NMR-Spectra for Compound 19a

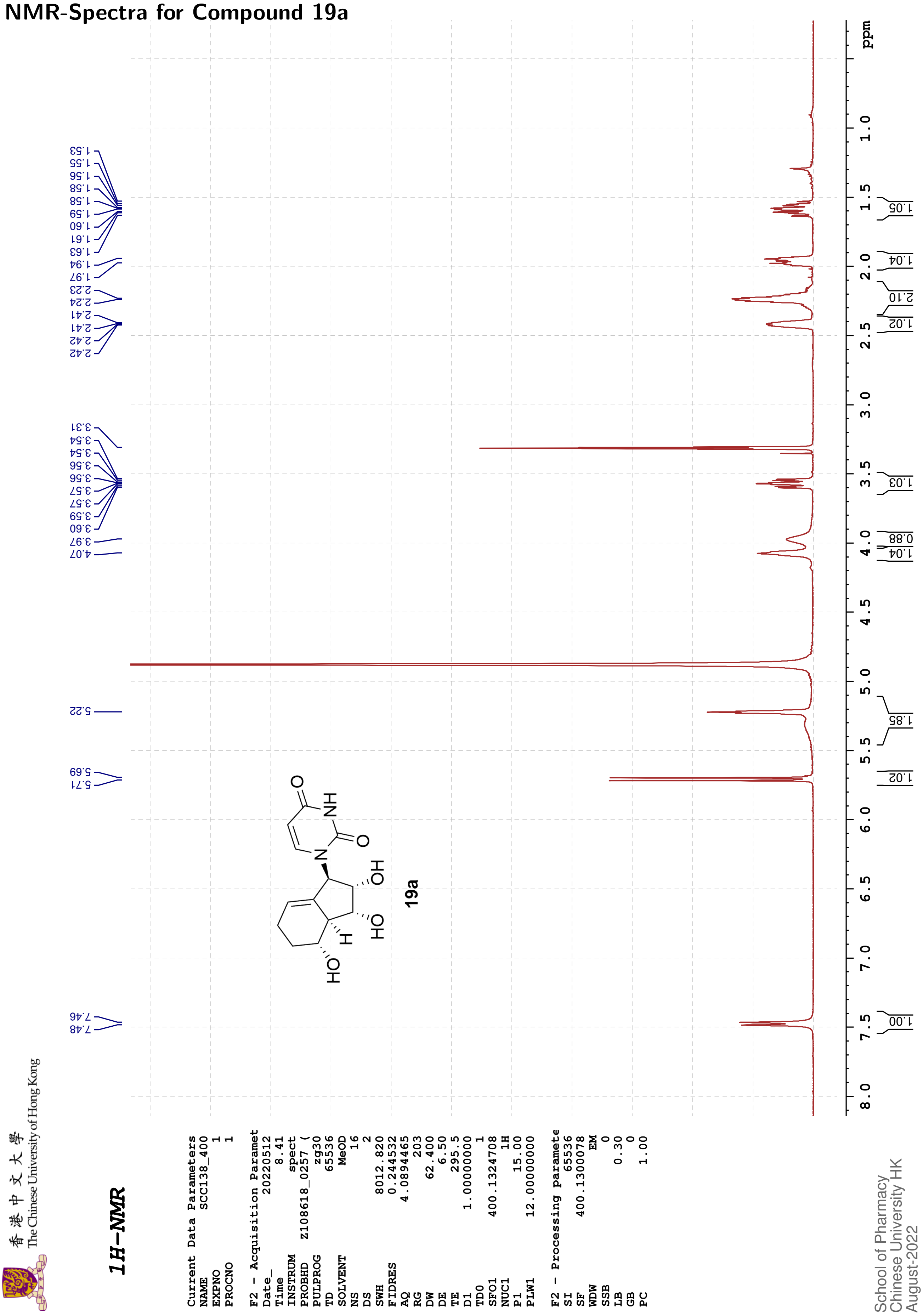

The Chinese University of Hong Kong

#### 13C{1H}-NMR

| Current Data Parameters | |
|---|---|
| NAME | SCC138_400 |
| EXPNO | 2 |
| PROCNO | 1 |

| F2 - Acquisition Paramet | |
|---|---|
| Date_ | 20220512 |
| Time | 8.58 |
| INSTRUM | spect |
| PROBHD | Z108618_0257 ( |
| PULPROG | zgpg30 |
| TD | 48074 |
| SOLVENT | MeOD |
| NS | 305 |
| DS | 4 |
| SWH | 24038.461 |
| FIDRES | 1.000061 |
| AQ | 0.9999392 |
| RG | 203 |
| DW | 20.800 |
| DE | 6.50 |
| TE | 296.2 |
| D1 | 2.00000000 |
| D11 | 0.03000000 |
| TD0 | 1 |
| SFO1 | 100.6228298 |
| NUC1 | 13C |
| P1 | 10.00 |
| PLW1 | 47.00000000 |
| SFO2 | 400.1316005 |
| NUC2 | 1H |
| CPDPRG[2 | waltz16 |
| PCPD2 | 90.00 |
| PLW2 | 12.00000000 |
| PLW12 | 0.33333001 |
| PLW13 | 0.16766000 |

| F2 - Processing paramete | |
|---|---|
| SI | 32768 |
| SF | 100.6126287 |
| WDW | EM |
| SSB | 0 |
| LB | 3.50 |
| GB | 0 |
| PC | 1.40 |

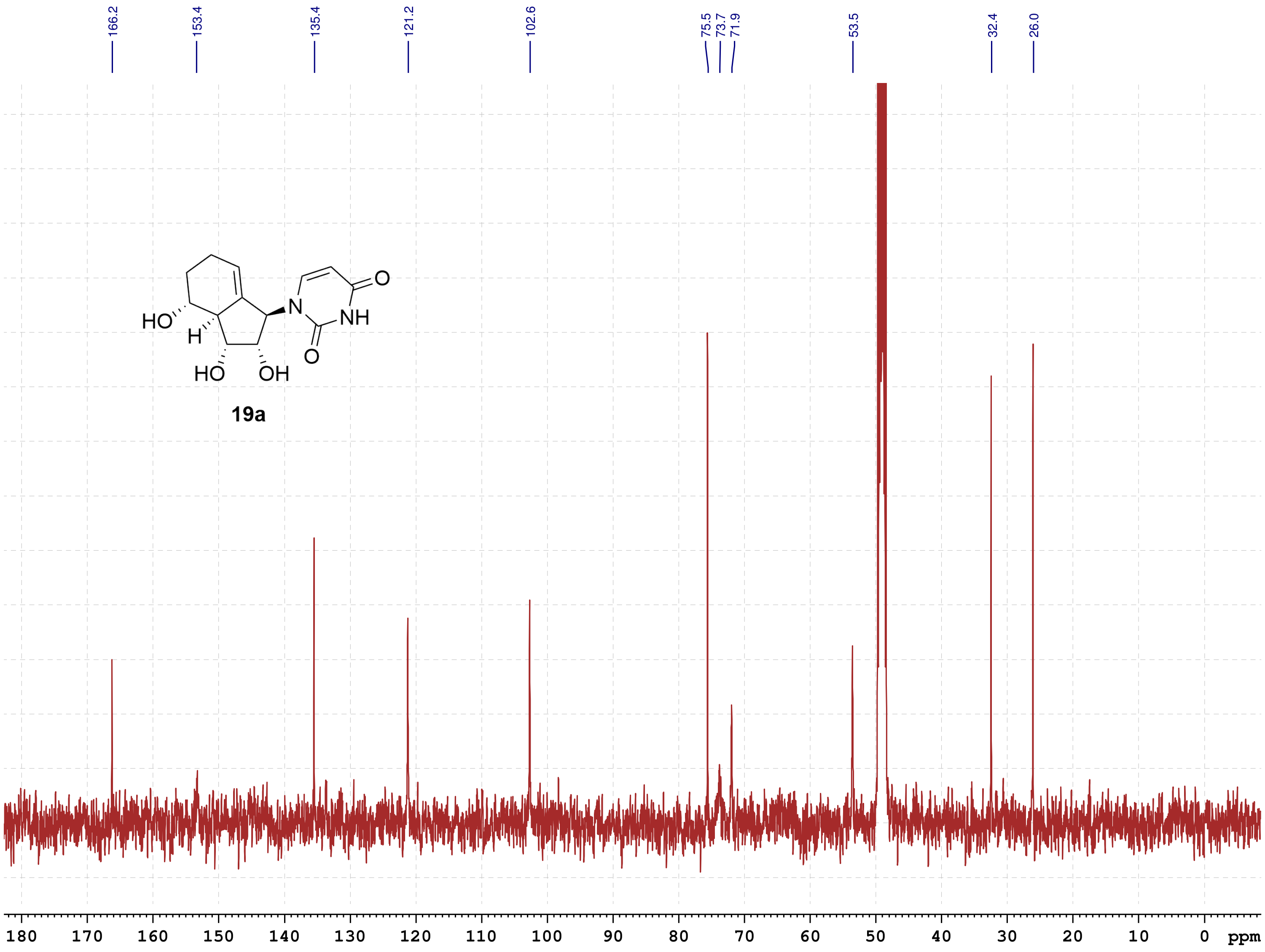

School of Pharmacy
Chinese University HK
August-2022

#### NMR-Spectra for Compound 19b

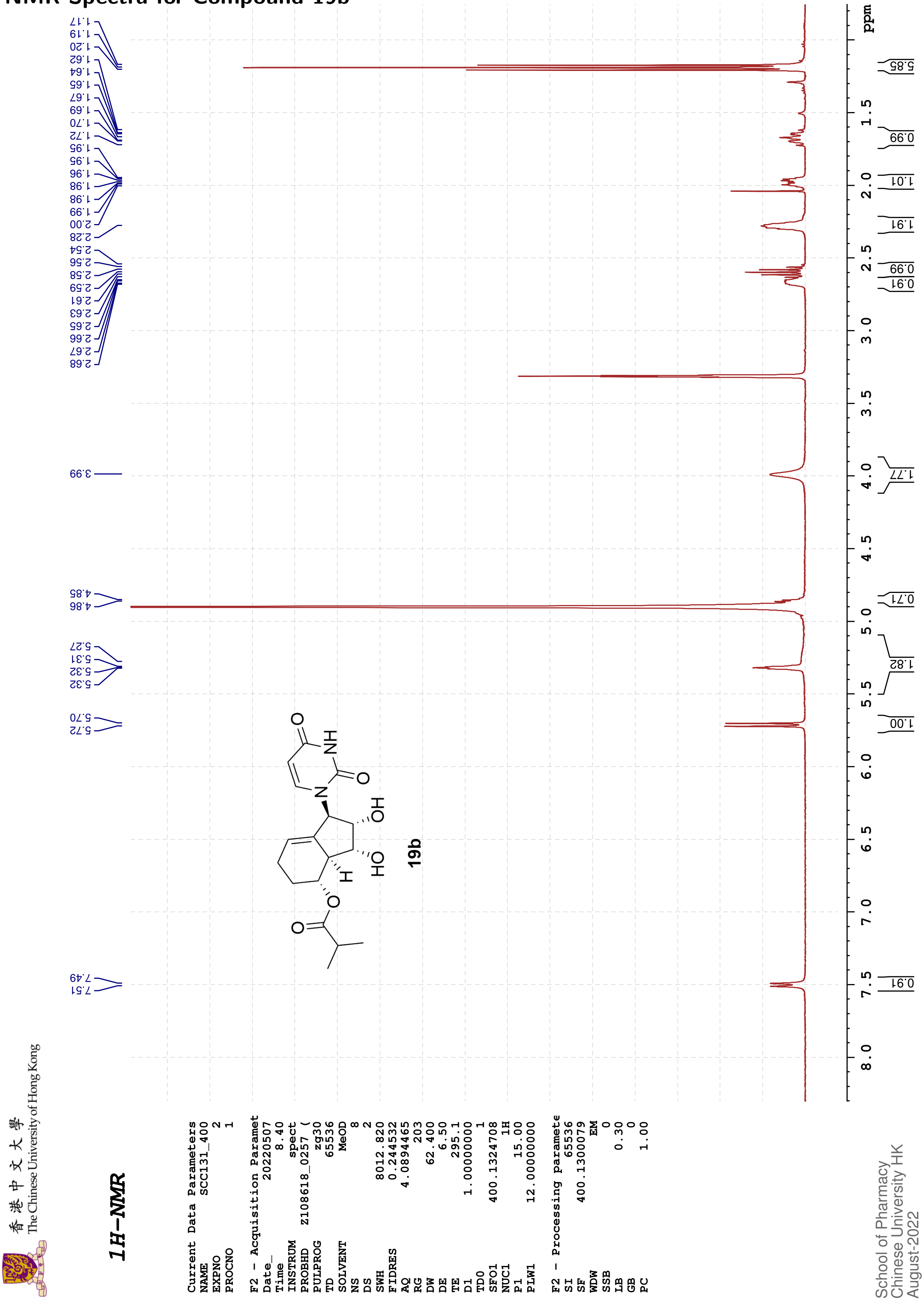

香港中文大學
The Chinese University of Hong Kong

#### 13C{1H}-NMR

```
Current Data Parameters
NAME       SCC056-F1_400
EXPNO                  3
PROCNO                 1

F2 - Acquisition Paramet
Date_           20220120
Time                8.14
INSTRUM            spect
PROBHD    Z108618_0257 (
PULPROG          zgpg30
TD                65536
SOLVENT            MeOD
NS                  197
DS                    4
SWH           24038.461
FIDRES         0.733596
AQ            1.3631488
RG                  203
DW               20.800
DE                 6.50
TE                294.9
D1           2.00000000
D11          0.03000000
TD0                   1
SFO1        100.6228298
NUC1                13C
P1                10.00
PLW1        47.00000000
SFO2        400.1316005
NUC2                 1H
CPDPRG[2        waltz16
PCPD2             90.00
PLW2        12.00000000
PLW12        0.33333001
PLW13        0.16766000

F2 - Processing paramete
SI                32768
SF          100.6126274
WDW                  EM
SSB                   0
LB                 1.00
GB                    0
PC                 1.40
```

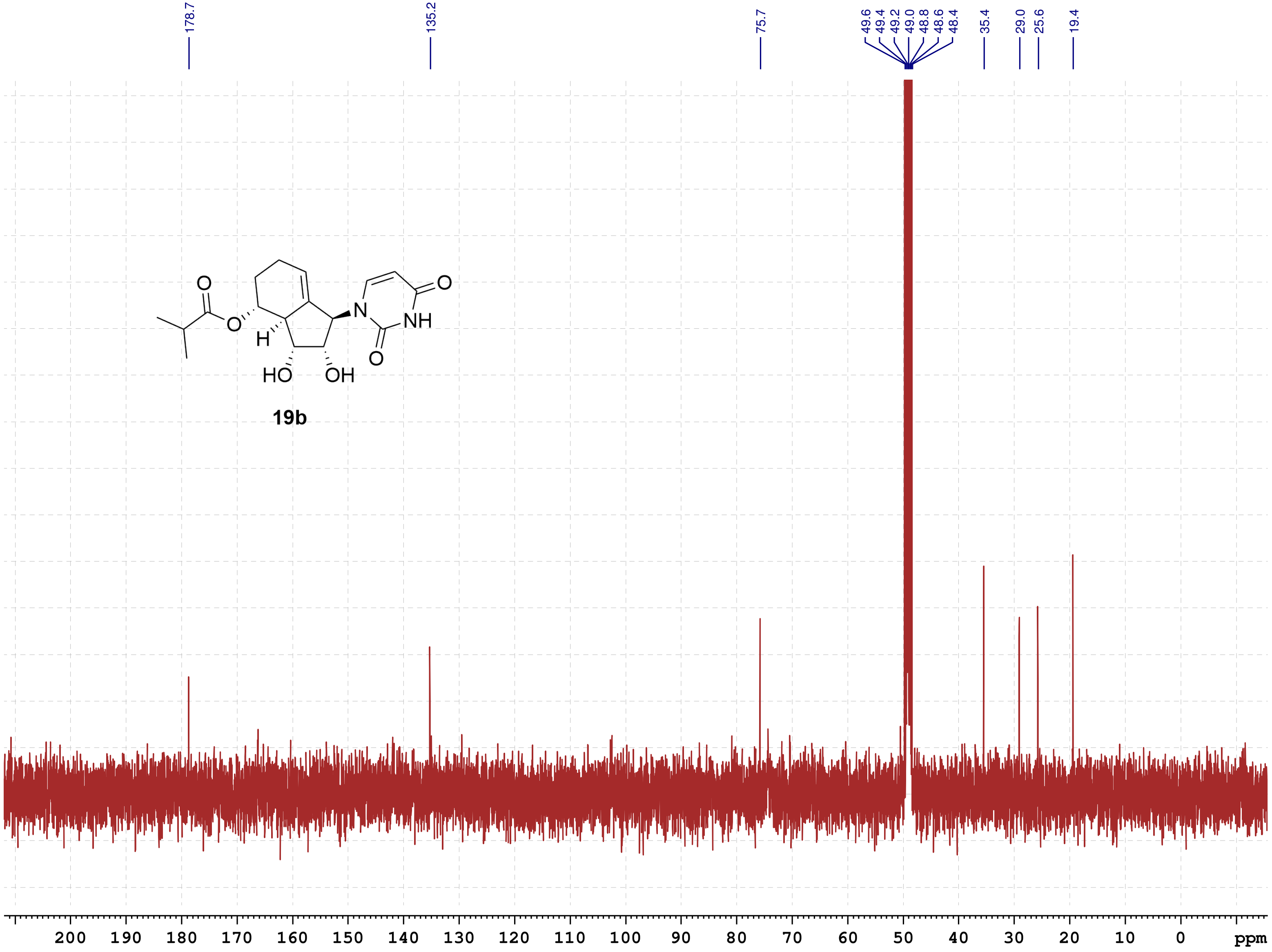

**19b**

School of Pharmacy
Chinese University HK
August-2022

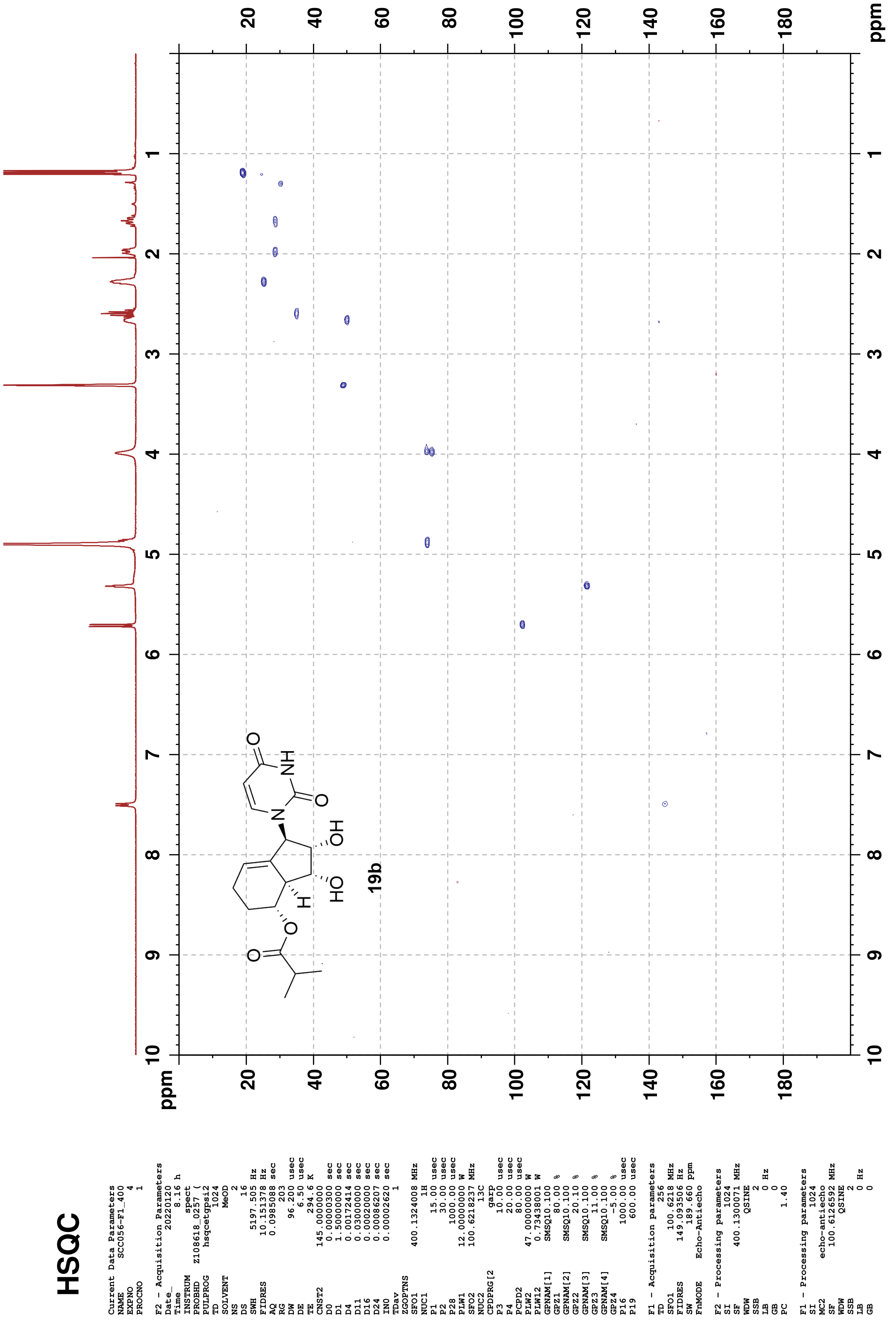
HSQC
Current Data Parameters
NAME SCC056-F1_400
EXPNO 4
PROCNO 1
F2 - Acquisition Parameters
Date_ 20220120
Time 8.16 h
INSTRUM spect
PROBHD Z108618_0257 (
PULPROG hsqcetgpsi2
TD 1024
SOLVENT MeOD
NS 2
DS 16
SWH 5197.505 Hz
FIDRES 10.151378 Hz
AQ 0.0985088 sec
RG 203
DW 96.200 usec
DE 6.50 usec
TE 294.6 K
CNST2 145.0000000
D0 0.00000300 sec
D1 1.50000000 sec
D4 0.00172414 sec
D11 0.03000000 sec
D16 0.00020000 sec
D24 0.00086207 sec
IN0 0.00002620 sec
TDav 1
ZGOPTNS
SFO1 400.1324008 MHz
NUC1 1H
P1 15.00 usec
P2 30.00 usec
P28 1000.00 usec
PLW1 12.00000000 W
SFO2 100.6218237 MHz
NUC2 13C
CPDPRG[2 garp
P3 10.00 usec
P4 20.00 usec
PCPD2 80.00 usec
PLW2 47.00000000 W
PLW12 0.73438001 W
GPNAM[1] SMSQ10.100
GPZ1 80.00 %
GPNAM[2] SMSQ10.100
GPZ2 20.10 %
GPNAM[3] SMSQ10.100
GPZ3 11.00 %
GPNAM[4] SMSQ10.100
GPZ4 -5.00 %
P16 1000.00 usec
P19 600.00 usec
F1 - Acquisition parameters
TD 256
SFO1 100.6218 MHz
FIDRES 149.093506 Hz
SW 189.660 ppm
FnMODE Echo-Antiecho
F2 - Processing parameters
SI 1024
SF 400.1300071 MHz
WDW QSINE
SSB 2
LB 0 Hz
GB 0
PC 1.40
F1 - Processing parameters
SI 1024
MC2 echo-antiecho
SF 100.6126592 MHz
WDW QSINE
SSB 2
LB 0 Hz
GB 0
19b
ppm

### HMBC

```
Current Data Parameters
NAME        SCC056-F1_400
EXPNO                   5
PROCNO                  1

F2 - Acquisition Parameters
Date_            20220120
Time                 8.31 h
INSTRUM             spect
PROBHD   Z108618_0257 (
PULPROG      hmbcgplpndqf
TD                   2048
SOLVENT              MeOD
NS                      2
DS                     16
SWH              5197.505 Hz
FIDRES           5.075689 Hz
AQ              0.1970176 sec
RG                    203
DW                 96.200 usec
DE                   6.50 usec
TE                  294.3 K
CNST2         145.0000000
CNST13         10.0000000
D0             0.00000300 sec
D1             1.50000000 sec
D2             0.00344828 sec
D6             0.05000000 sec
D16            0.00020000 sec
IN0            0.00002260 sec
TDav                    1
SFO1          400.1325208 MHz
NUC1                   1H
P1                  15.00 usec
P2                  30.00 usec
PLW1          12.00000000 W
SFO2          100.6228298 MHz
NUC2                  13C
P3                  10.00 usec
PLW2          47.00000000 W
GPNAM[1]        SMSQ10.100
GPZ1                50.00 %
GPNAM[2]        SMSQ10.100
GPZ2                30.00 %
GPNAM[3]        SMSQ10.100
GPZ3                40.10 %
P16               1000.00 usec

F1 - Acquisition parameters
TD                    128
SFO1             100.6228 MHz
FIDRES         345.685852 Hz
SW                219.870 ppm
FnMODE                 QF

F2 - Processing parameters
SI                   2048
SF            400.1300000 MHz
WDW                  SINE
SSB                     0
LB                      0 Hz
GB                      0
PC                   1.40

F1 - Processing parameters
SI                   1024
MC2                    QF
SF            100.6127685 MHz
WDW                  SINE
SSB                     0
LB                      0 Hz
GB                      0
```

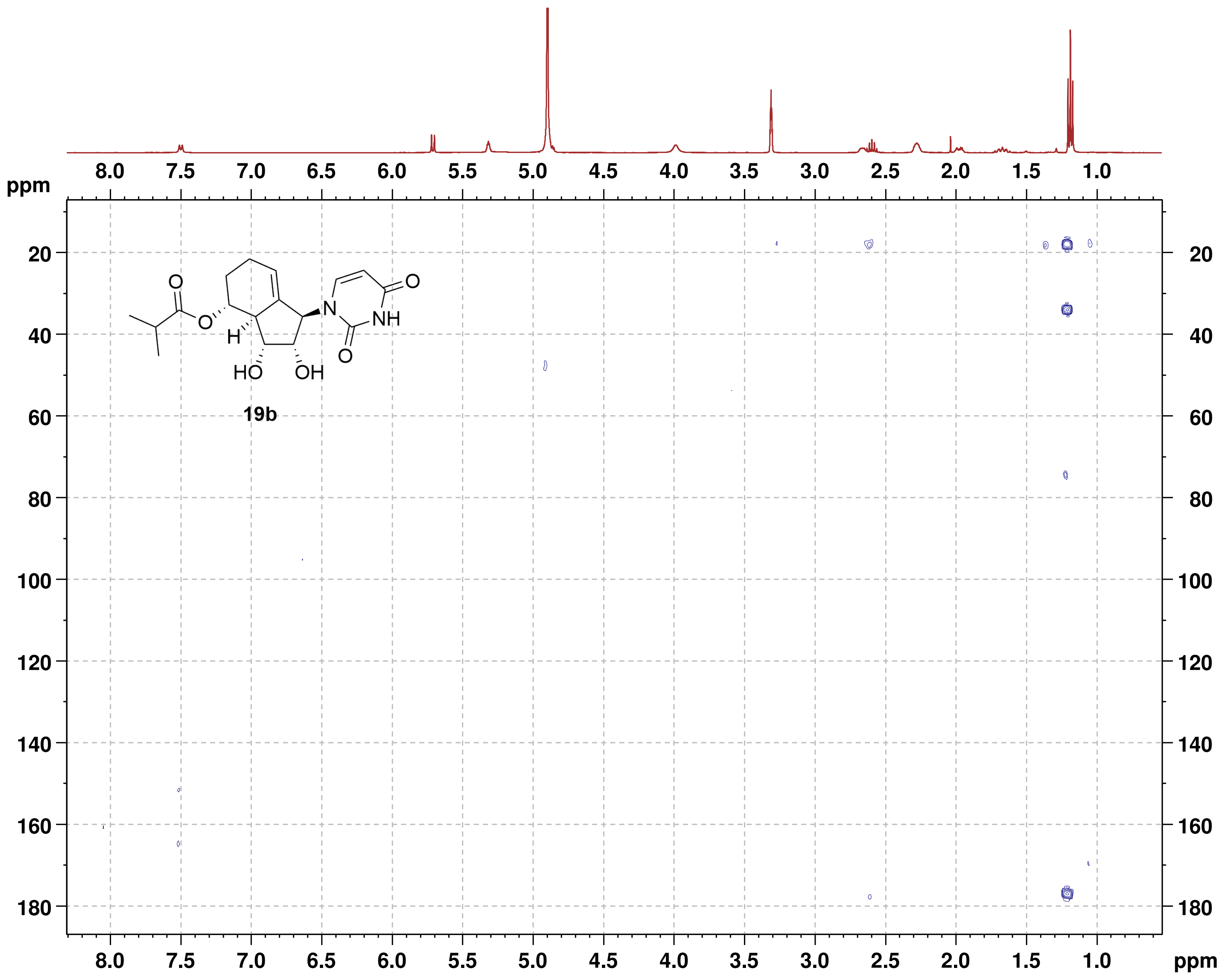

**NOESY**

```
Current Data Parameters
NAME          SCC131_400
EXPNO                  5
PROCNO                 1

F2 - Acquisition Parameters
Date_           20220507
Time                8.49 h
INSTRUM            spect
PROBHD   Z108618_0257 (
PULPROG       noesygpphpp
TD                  2048
SOLVENT             MeOD
NS                     2
DS                    16
SWH             6009.615 Hz
FIDRES          5.868765 Hz
AQ             0.1703936 sec
RG                   203
DW                83.200 usec
DE                  6.50 usec
TE                 295.1 K
D0         0.00006420 sec
D1         2.00000000 sec
D8         0.60000002 sec
D11        0.03000000 sec
D12        0.00002000 sec
D16        0.00020000 sec
IN0        0.00016660 sec
TDav                   1
SFO1         400.1320007 MHz
NUC1                  1H
P1                 15.00 usec
P2                 30.00 usec
P17              2500.00 usec
PLW1         12.00000000 W
PLW10         3.00000000 W
GPNAM[1]      SMSQ10.100
GPZ1               40.00 %
P16              1000.00 usec

F1 - Acquisition parameters
TD                   256
SFO1             400.132 MHz
FIDRES         46.893757 Hz
SW                15.001 ppm
FnMODE        States-TPPI

F2 - Processing parameters
SI                  1024
SF           400.1300072 MHz
WDW                QSINE
SSB                    2
LB                     0 Hz
GB                     0
PC                  1.00

F1 - Processing parameters
SI                  1024
MC2          States-TPPI
SF           400.1300068 MHz
WDW                QSINE
SSB                    2
LB                     0 Hz
GB                     0
```

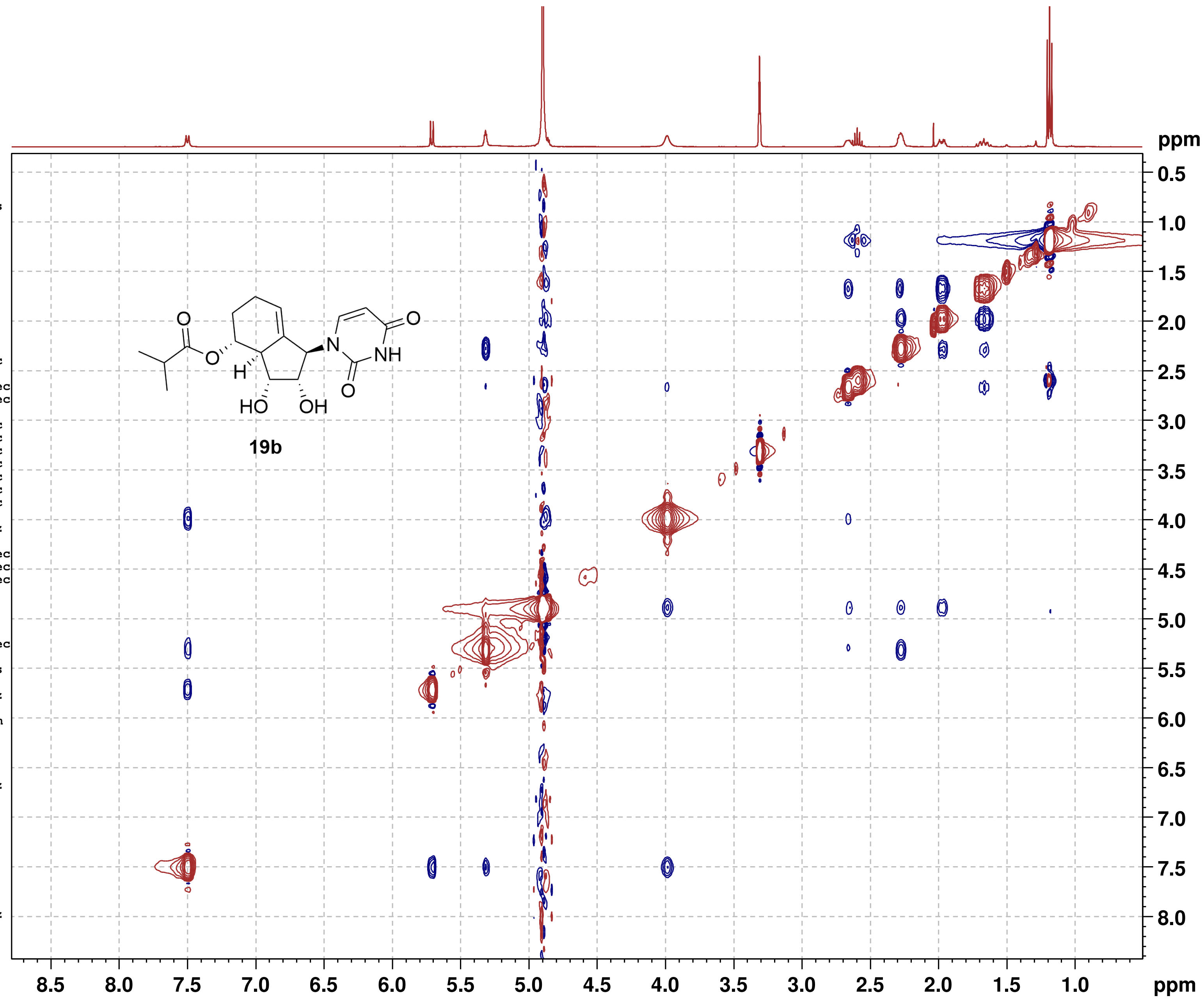

#### NMR-Spectra for Compound 20a

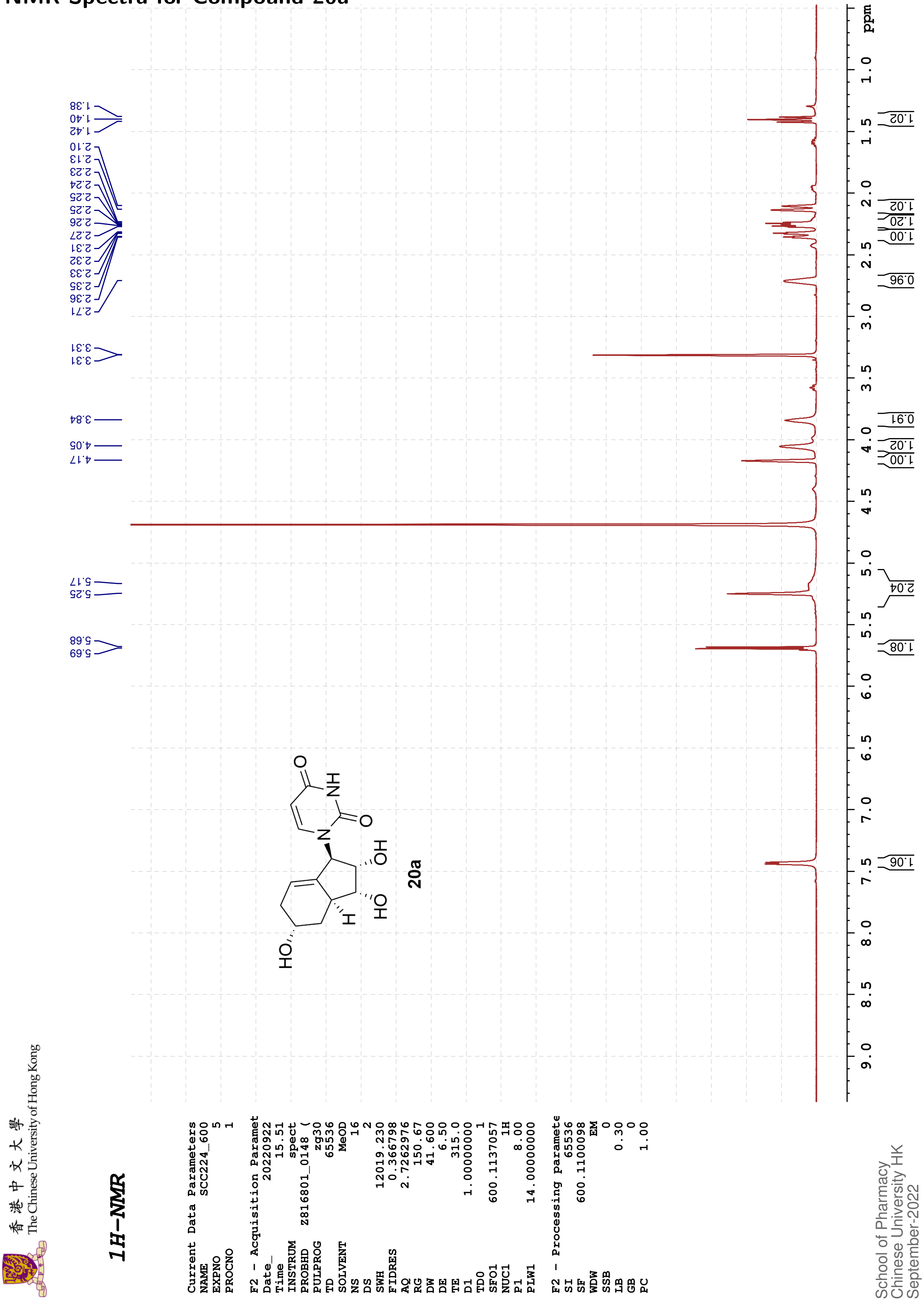

The Chinese University of Hong Kong

#### 13C{1H}-NMR

| | |
|---|---|
| Current Data Parameters | |
| NAME | SCC224_600 |
| EXPNO | 2 |
| PROCNO | 1 |
| F2 - Acquisition Paramet | |
| Date_ | 20220906 |
| Time | 15.04 |
| INSTRUM | spect |
| PROBHD | Z816801_0148 ( |
| PULPROG | zgpg30 |
| TD | 65536 |
| SOLVENT | MeOD |
| NS | 100 |
| DS | 4 |
| SWH | 36231.883 |
| FIDRES | 1.105709 |
| AQ | 0.9043968 |
| RG | 98.37 |
| DW | 13.800 |
| DE | 6.50 |
| TE | 315.0 |
| D1 | 2.00000000 |
| D11 | 0.03000000 |
| TD0 | 1 |
| SFO1 | 150.9128693 |
| NUC1 | 13C |
| P1 | 12.00 |
| PLW1 | 183.00000000 |
| SFO2 | 600.1124004 |
| NUC2 | 1H |
| CPDPRG[2 | waltz16 |
| PCPD2 | 70.00 |
| PLW2 | 14.00000000 |
| PLW12 | 0.18286000 |
| PLW13 | 0.09197600 |
| F2 - Processing paramete | |
| SI | 32768 |
| SF | 150.8975706 |
| WDW | EM |
| SSB | 0 |
| LB | 2.00 |
| GB | 0 |
| PC | 1.40 |

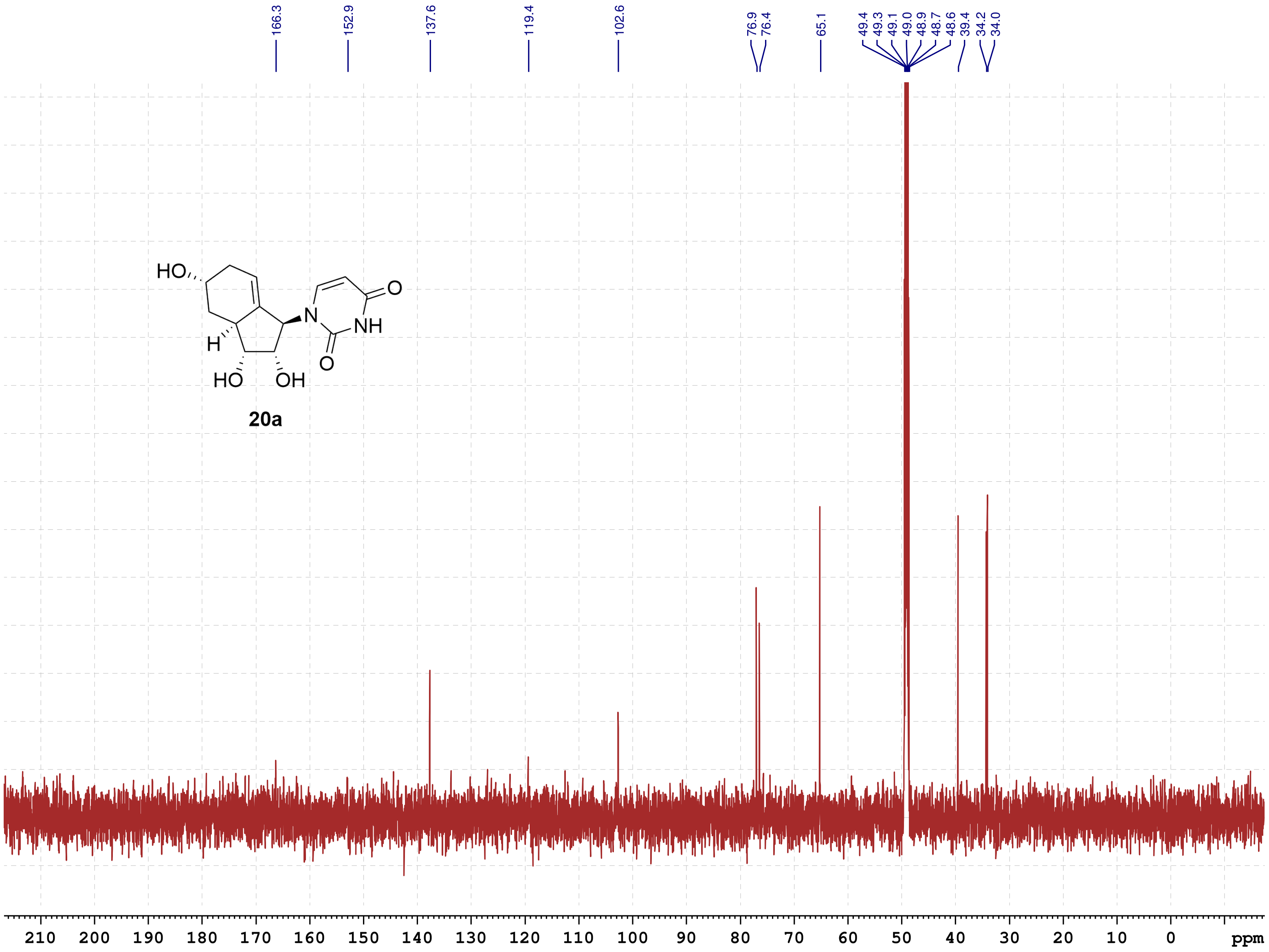

School of Pharmacy
Chinese University HK
September-2022

香港中文大學 The Chinese University of Hong Kong

**COSY**

Current Data Parameters
NAME SCC224_600
EXPNO 6
PROCNO 1

F2 - Acquisition Parameters
Date_ 20220922
Time 15.51 h
INSTRUM spect
PROBHD Z816801_0148 (
PULPROG cosygpppqf
TD 2048
SOLVENT MeOD
NS 1
DS 8
SWH 7812.500 Hz
FIDRES 7.629395 Hz
AQ 0.1310720 sec
RG 189.92
DW 64.000 usec
DE 6.50 usec
TE 315.0 K
D0 0.00000300 sec
D1 2.00000000 sec
D11 0.03000000 sec
D12 0.00002000 sec
D13 0.00000400 sec
D16 0.00020000 sec
IN0 0.00012800 sec
TDav 1
SFO1 600.1136007 MHz
NUC1 1H
P0 8.00 usec
P1 8.00 usec
P17 2500.00 usec
PLW1 14.00000000 W
PLW10 1.43359995 W
GPNAM[1] SMSQ10.100
GPZ1 10.00 %
P16 1000.00 usec

F1 - Acquisition parameters
TD 128
SFO1 600.1136 MHz
FIDRES 122.070312 Hz
SW 13.018 ppm
FnMODE QF

F2 - Processing parameters
SI 1024
SF 600.1100076 MHz
WDW QSINE
SSB 0
LB 0 Hz
GB 0
PC 1.40

F1 - Processing parameters
SI 1024
MC2 QF
SF 600.1100079 MHz
WDW QSINE
SSB 0
LB 0 Hz
GB 0

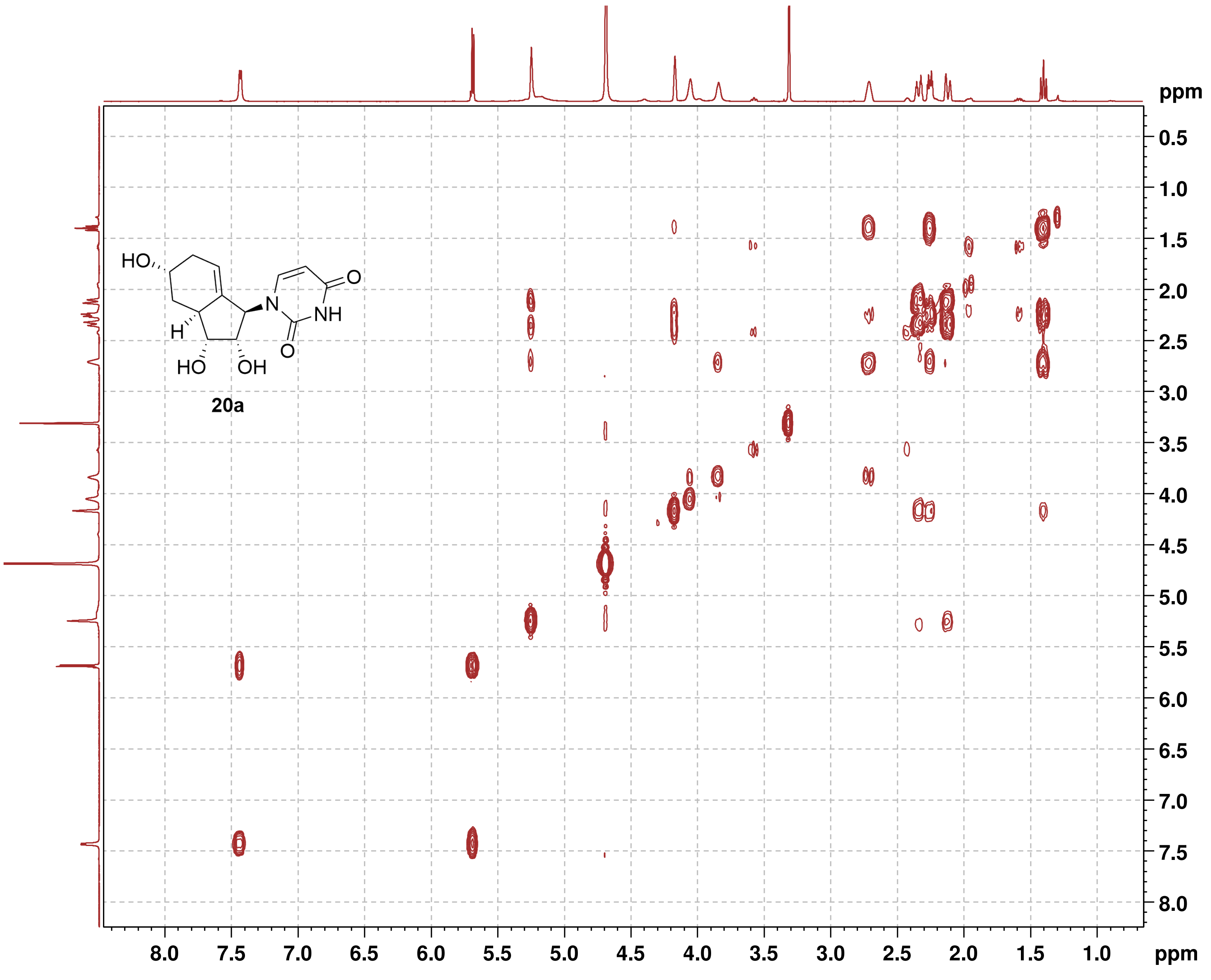

香港中文大學
The Chinese University of Hong Kong

**HSQC**

```
Current Data Parameters
NAME          SCC224_600
EXPNO                  3
PROCNO                 1

F2 - Acquisition Parameters
Date_           20220906
Time               15.06 h
INSTRUM            spect
PROBHD   Z816801_0148 (
PULPROG      hsqcetgpsi2
TD                  1024
SOLVENT             MeOD
NS                     3
DS                    16
SWH             7812.500 Hz
FIDRES         15.258789 Hz
AQ             0.0655360 sec
RG                189.92
DW                64.000 usec
DE                  6.50 usec
TE                 315.2 K
CNST2        145.0000000
D0            0.00000300 sec
D1            1.50000000 sec
D4            0.00172414 sec
D11           0.03000000 sec
D16           0.00020000 sec
D24           0.00086207 sec
IN0           0.00001660 sec
TDav                   1
ZGOPTNS
SFO1         600.1137057 MHz
NUC1                  1H
P1                  8.00 usec
P2                 16.00 usec
P28              1000.00 usec
PLW1         14.00000000 W
SFO2         150.9128693 MHz
NUC2                 13C
CPDPRG[2            garp
P3                 12.00 usec
P4                 24.00 usec
PCPD2              60.00 usec
PLW2        183.00000000 W
PLW12         7.32000017 W
GPNAM[1]      SMSQ10.100
GPZ1               80.00 %
GPNAM[2]      SMSQ10.100
GPZ2               20.10 %
GPNAM[3]      SMSQ10.100
GPZ3               11.00 %
GPNAM[4]      SMSQ10.100
GPZ4               -5.00 %
P16              1000.00 usec
P19               600.00 usec

F1 - Acquisition parameters
TD                   128
SFO1            150.9129 MHz
FIDRES        470.632538 Hz
SW               199.589 ppm
FnMODE      Echo-Antiecho

F2 - Processing parameters
SI                  1024
SF           600.1100094 MHz
WDW                QSINE
SSB                    2
LB                     0 Hz
GB                     0
PC                  1.40

F1 - Processing parameters
SI                  1024
MC2         echo-antiecho
SF           150.8975923 MHz
WDW                QSINE
SSB                    2
LB                     0 Hz
GB                     0
```

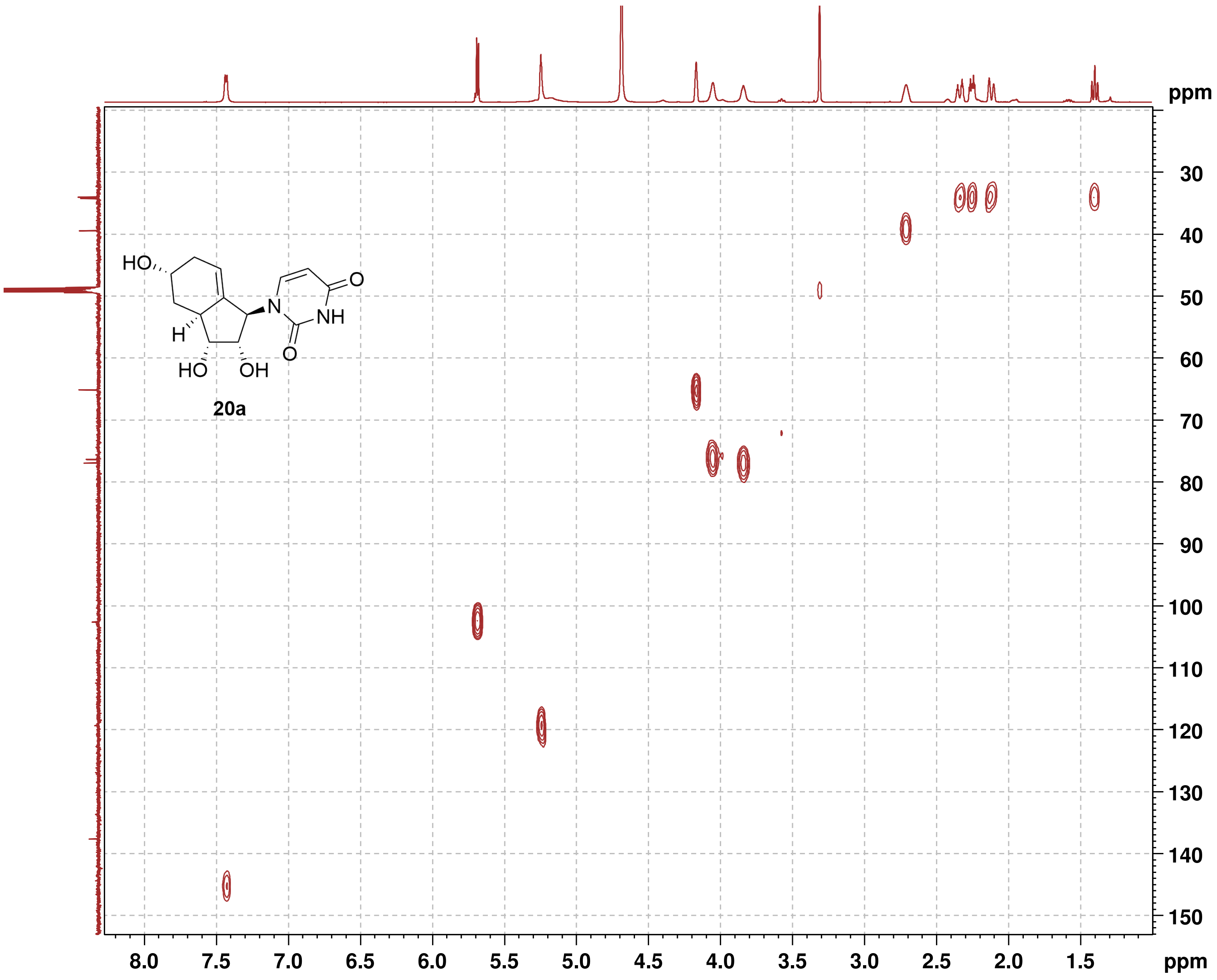

香港中文大學
The Chinese University of Hong Kong

**HMBC**

```
Current Data Parameters
NAME          SCC224_600
EXPNO                  4
PROCNO                 1

F2 - Acquisition Parameters
Date_           20220906
Time               15.17 h
INSTRUM            spect
PROBHD   Z816801_0148 (
PULPROG    hmbcgplpndqf
TD                  2048
SOLVENT             MeOD
NS                     4
DS                    16
SWH             6009.615 Hz
FIDRES          5.868765 Hz
AQ             0.1703936 sec
RG                189.92
DW                83.200 usec
DE                  6.50 usec
TE                 315.0 K
CNST2        145.0000000
CNST13        10.0000000
D0            0.00000300 sec
D1            1.50000000 sec
D2            0.00344828 sec
D6            0.05000000 sec
D16           0.00020000 sec
IN0           0.00001510 sec
TDav                   1
SFO1         600.1127005 MHz
NUC1                  1H
P1                  8.00 usec
P2                 16.00 usec
PLW1         14.00000000 W
SFO2         150.9128693 MHz
NUC2                 13C
P3                 12.00 usec
PLW2        183.00000000 W
GPNAM[1]      SMSQ10.100
GPZ1               50.00 %
GPNAM[2]      SMSQ10.100
GPZ2               30.00 %
GPNAM[3]      SMSQ10.100
GPZ3               40.10 %
P16              1000.00 usec

F1 - Acquisition parameters
TD                   128
SFO1            150.9129 MHz
FIDRES        517.384094 Hz
SW               219.415 ppm
FnMODE                QF

F2 - Processing parameters
SI                  2048
SF           600.1100083 MHz
WDW                 SINE
SSB                    0
LB                     0 Hz
GB                     0
PC                  1.40

F1 - Processing parameters
SI                  1024
MC2                   QF
SF           150.8975412 MHz
WDW                 SINE
SSB                    0
LB                     0 Hz
GB                     0
```

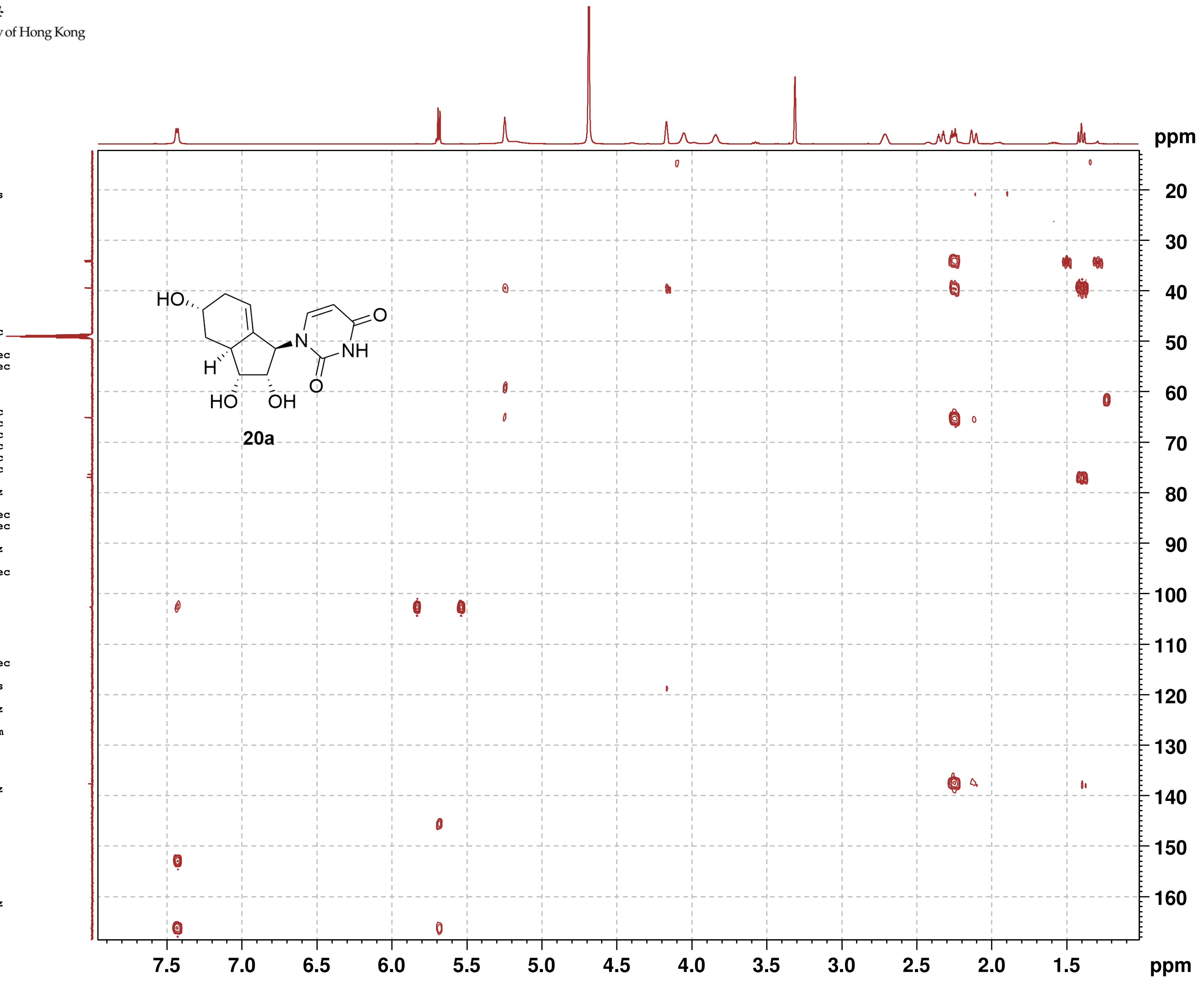

香 港 中 文 大 學
The Chinese University of Hong Kong

**NOESY**

```
Current Data Parameters
NAME          SCC224_600
EXPNO                  7
PROCNO                 1

F2 - Acquisition Parameters
Date_           20220922
Time               15.58 h
INSTRUM            spect
PROBHD   Z816801_0148 (
PULPROG        noesygpph
TD                  2048
SOLVENT             MeOD
NS                     2
DS                    16
SWH             6009.615 Hz
FIDRES          5.868765 Hz
AQ             0.1703936 sec
RG                189.92
DW                83.200 usec
DE                  6.50 usec
TE                 315.0 K
D0            0.00007301 sec
D1            2.00000000 sec
D8            0.75000000 sec
D16           0.00020000 sec
IN0           0.00016640 sec
TDav                   1
SFO1         600.1128205 MHz
NUC1                  1H
P1                  8.00 usec
P2                 16.00 usec
PLW1         14.00000000 W
GPNAM[1]      SMSQ10.100
GPZ1               40.00 %
P16              1000.00 usec

F1 - Acquisition parameters
TD                   128
SFO1            600.1128 MHz
FIDRES         93.900238 Hz
SW                10.014 ppm
FnMODE              TPPI

F2 - Processing parameters
SI                  1024
SF           600.1100100 MHz
WDW                QSINE
SSB                    2
LB                     0 Hz
GB                     0
PC                  1.00

F1 - Processing parameters
SI                  1024
MC2                 TPPI
SF           600.1100094 MHz
WDW                QSINE
SSB                    2
LB                     0 Hz
GB                     0
```

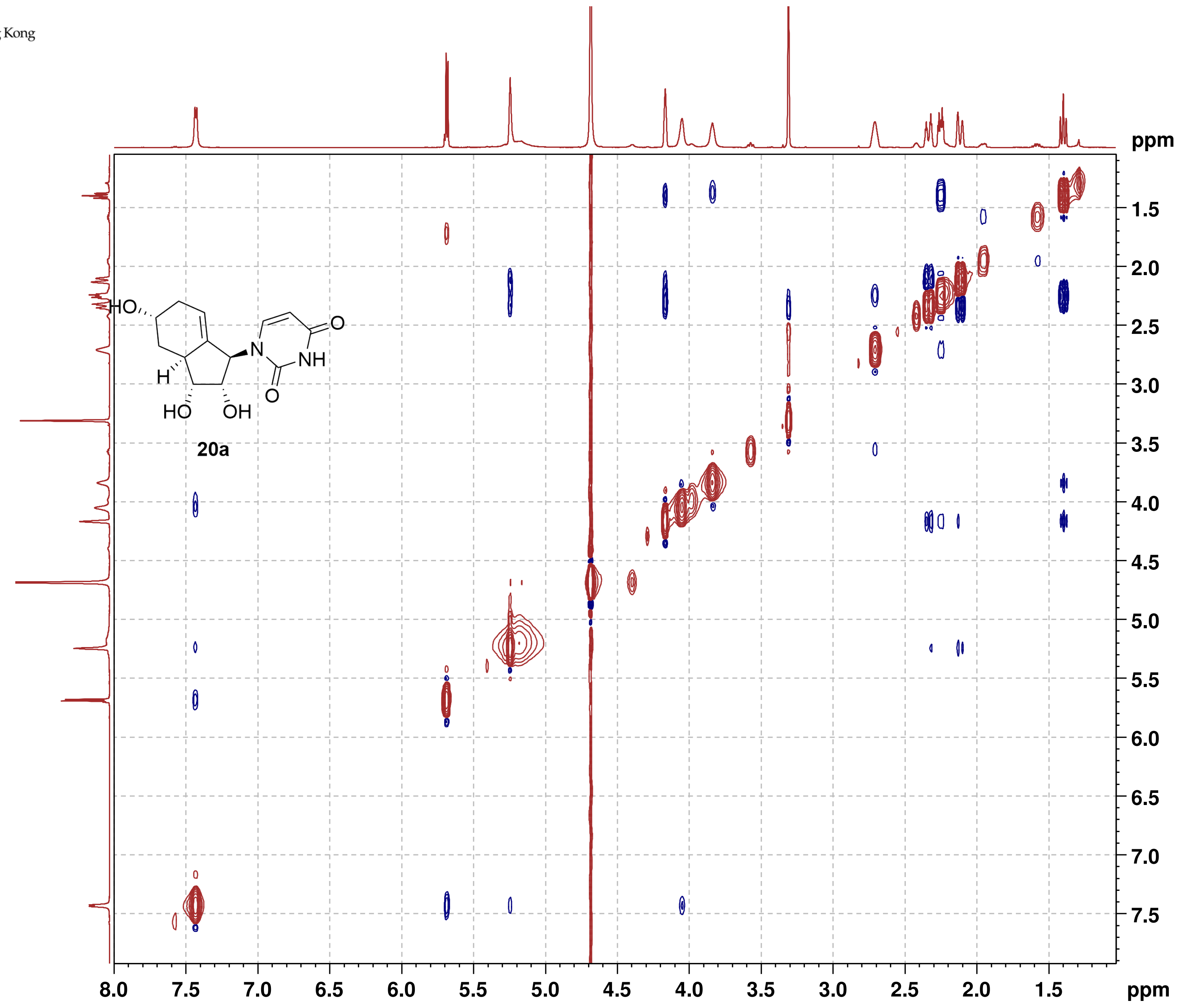

20a

#### NMR-Spectra for Compound 20b

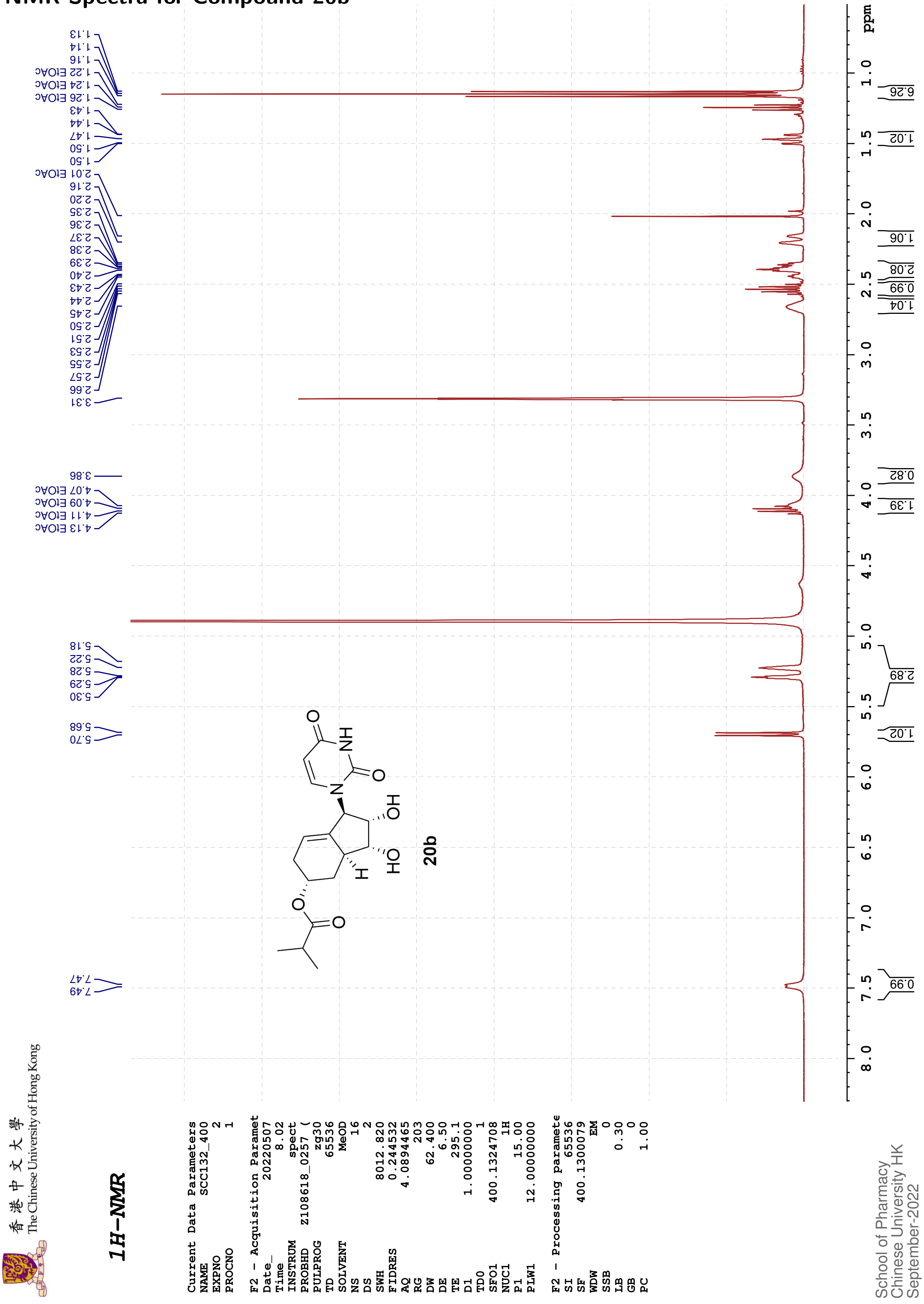

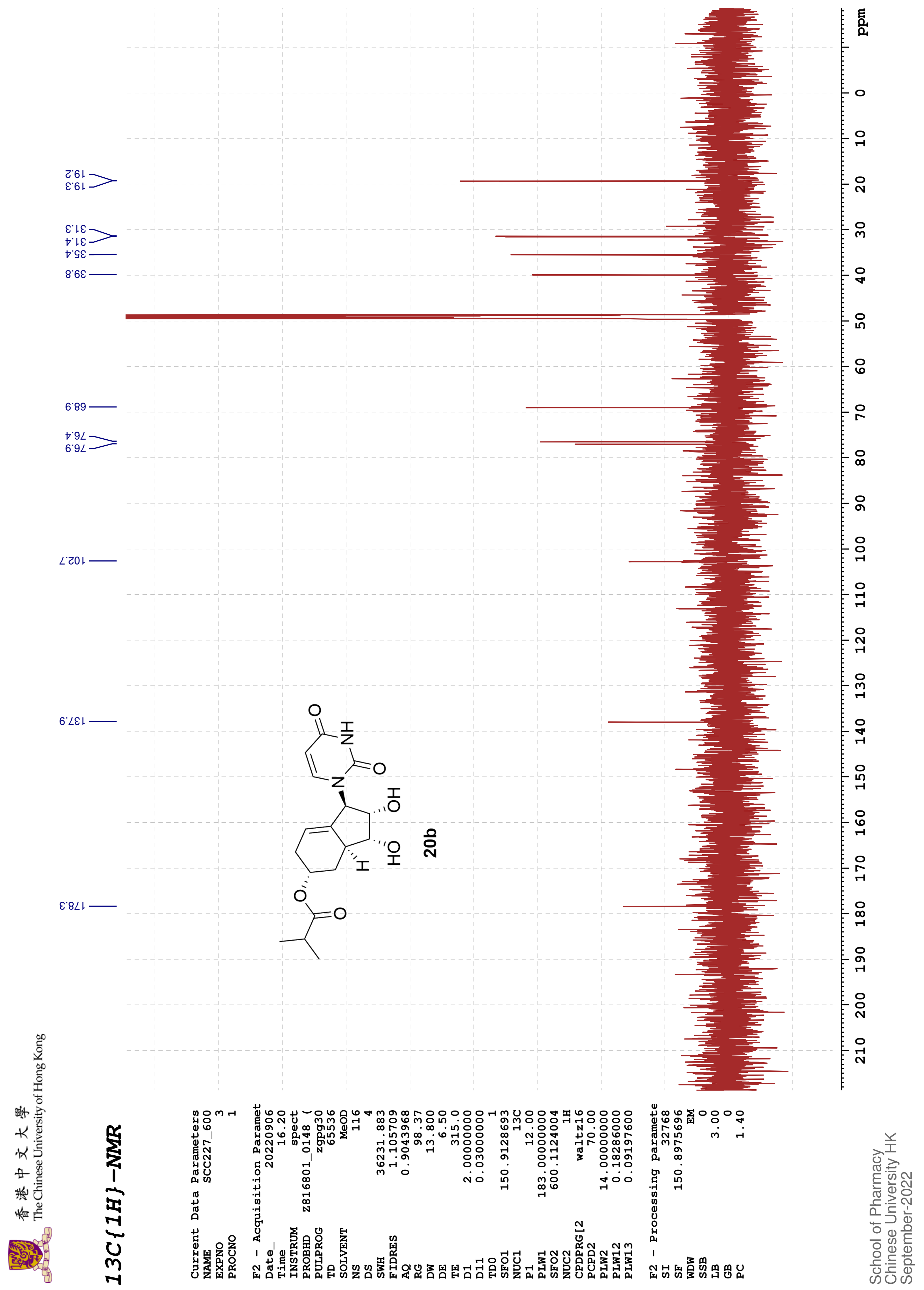
香港中文大學 The Chinese University of Hong Kong
13C{1H}-NMR
Current Data Parameters
NAME SCC227_600
EXPNO 3
PROCNO 1
F2 - Acquisition Paramet
Date_ 20220906
Time 16.20
INSTRUM spect
PROBHD Z816801_0148 (
PULPROG zgpg30
TD 65536
SOLVENT MeOD
NS 116
DS 4
SWH 36231.883
FIDRES 1.105709
AQ 0.9043968
RG 98.37
DW 13.800
DE 6.50
TE 315.0
D1 2.00000000
D11 0.03000000
TD0 1
SFO1 150.9128693
NUC1 13C
P1 12.00
PLW1 183.00000000
SFO2 600.1124004
NUC2 1H
CPDPRG[2 waltz16
PCPD2 70.00
PLW2 14.00000000
PLW12 0.18286000
PLW13 0.09197600
F2 - Processing paramete
SI 32768
SF 150.8975696
WDW EM
SSB 0
LB 3.00
GB 0
PC 1.40
School of Pharmacy
Chinese University HK
September-2022
178.3
137.9
102.7
76.9
76.4
68.9
39.8
35.4
31.4
31.3
19.3
19.2
20b
ppm

香港中文大學
The Chinese University of Hong Kong

COSY

```
Current Data Parameters
NAME          SCC132_400
EXPNO                  3
PROCNO                 1

F2 - Acquisition Parameters
Date_           20220507
Time                8.04 h
INSTRUM            spect
PROBHD   Z108618_0257 (
PULPROG         cosygpqf
TD                  2048
SOLVENT             MeOD
NS                     2
DS                     8
SWH             6009.615 Hz
FIDRES          5.868765 Hz
AQ             0.1703936 sec
RG                   203
DW                83.200 usec
DE                  6.50 usec
TE                 295.1 K
D0            0.00000300 sec
D1            1.48689198 sec
D13           0.00000400 sec
D16           0.00020000 sec
IN0           0.00016660 sec
TDav                   1
SFO1        400.1322007 MHz
NUC1                  1H
P0                 15.00 usec
P1                 15.00 usec
PLW1         12.00000000 W
GPNAM[1]       SINE.100
GPZ1               10.00 %
P16              1000.00 usec

F1 - Acquisition parameters
TD                   128
SFO1            400.1322 MHz
FIDRES         93.787514 Hz
SW                15.001 ppm
FnMODE                QF

F2 - Processing parameters
SI                  1024
SF           400.1300094 MHz
WDW                 SINE
SSB                    0
LB                     0 Hz
GB                     0
PC                  1.40

F1 - Processing parameters
SI                  1024
MC2                   QF
SF           400.1300095 MHz
WDW                 SINE
SSB                    0
LB                     0 Hz
GB                     0
```

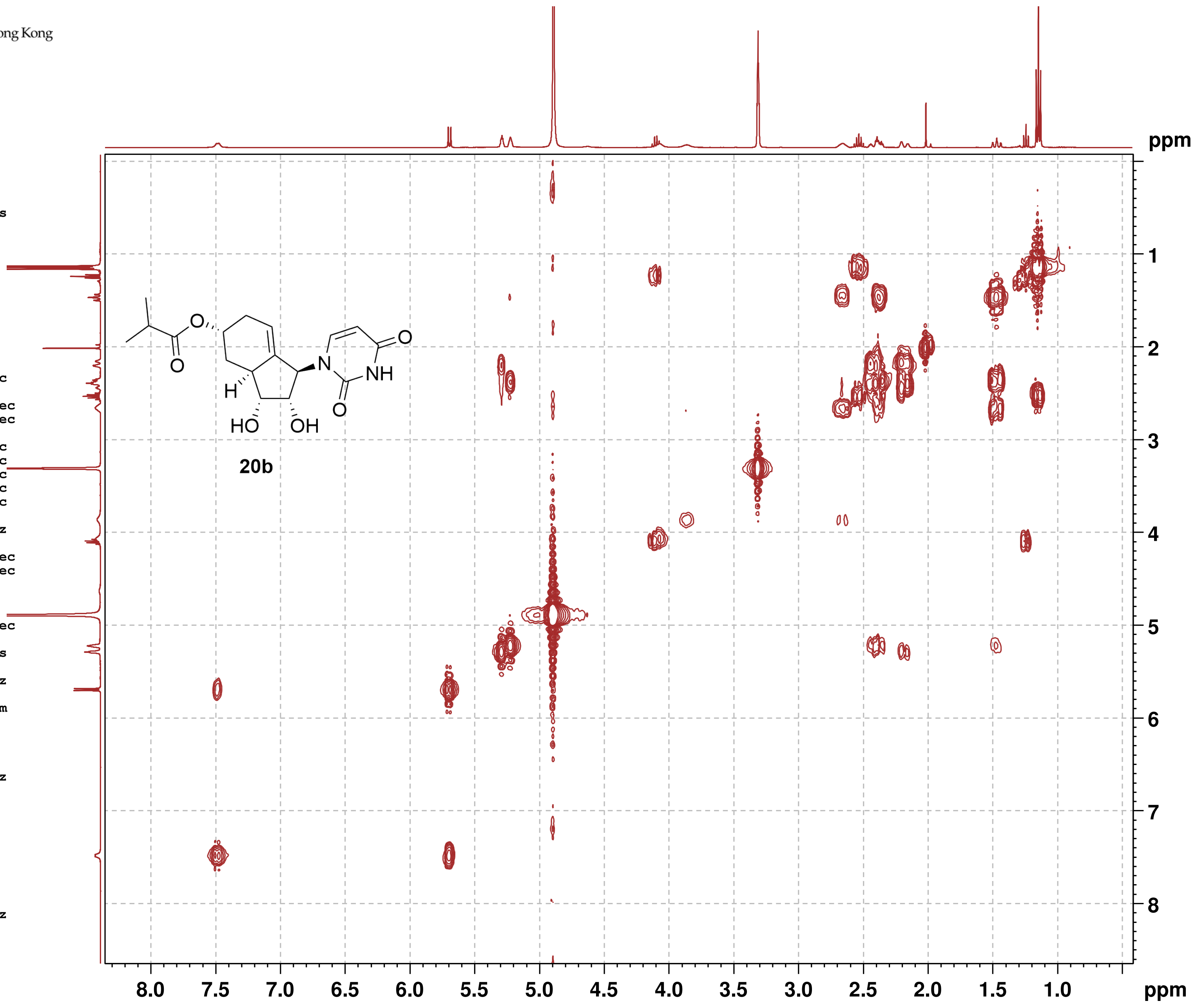

香港中文大學
The Chinese University of Hong Kong

#### HSQC

```
Current Data Parameters
NAME          SCC227_600
EXPNO                  4
PROCNO                 1

F2 - Acquisition Parameters
Date_           20220906
Time               16.22 h
INSTRUM            spect
PROBHD   Z816801_0148 (
PULPROG      hsqcetgpsi2
TD                  1024
SOLVENT             MeOD
NS                     3
DS                    16
SWH             7812.500 Hz
FIDRES         15.258789 Hz
AQ             0.0655360 sec
RG                189.92
DW                64.000 usec
DE                  6.50 usec
TE                 315.1 K
CNST2        145.0000000
D0            0.00000300 sec
D1            1.50000000 sec
D4            0.00172414 sec
D11           0.03000000 sec
D16           0.00020000 sec
D24           0.00086207 sec
IN0           0.00001660 sec
TDav                   1
ZGOPTNS
SFO1         600.1137057 MHz
NUC1                  1H
P1                  8.00 usec
P2                 16.00 usec
P28              1000.00 usec
PLW1         14.00000000 W
SFO2         150.9128693 MHz
NUC2                 13C
CPDPRG[2            garp
P3                 12.00 usec
P4                 24.00 usec
PCPD2              60.00 usec
PLW2        183.00000000 W
PLW12         7.32000017 W
GPNAM[1]      SMSQ10.100
GPZ1               80.00 %
GPNAM[2]      SMSQ10.100
GPZ2               20.10 %
GPNAM[3]      SMSQ10.100
GPZ3               11.00 %
GPNAM[4]      SMSQ10.100
GPZ4               -5.00 %
P16              1000.00 usec
P19               600.00 usec

F1 - Acquisition parameters
TD                   128
SFO1            150.9129 MHz
FIDRES        470.632538 Hz
SW               199.589 ppm
FnMODE      Echo-Antiecho

F2 - Processing parameters
SI                  1024
SF           600.1100095 MHz
WDW                QSINE
SSB                    2
LB                     0 Hz
GB                     0
PC                  1.40

F1 - Processing parameters
SI                  1024
MC2         echo-antiecho
SF           150.8976023 MHz
WDW                QSINE
SSB                    2
LB                     0 Hz
GB                     0
```

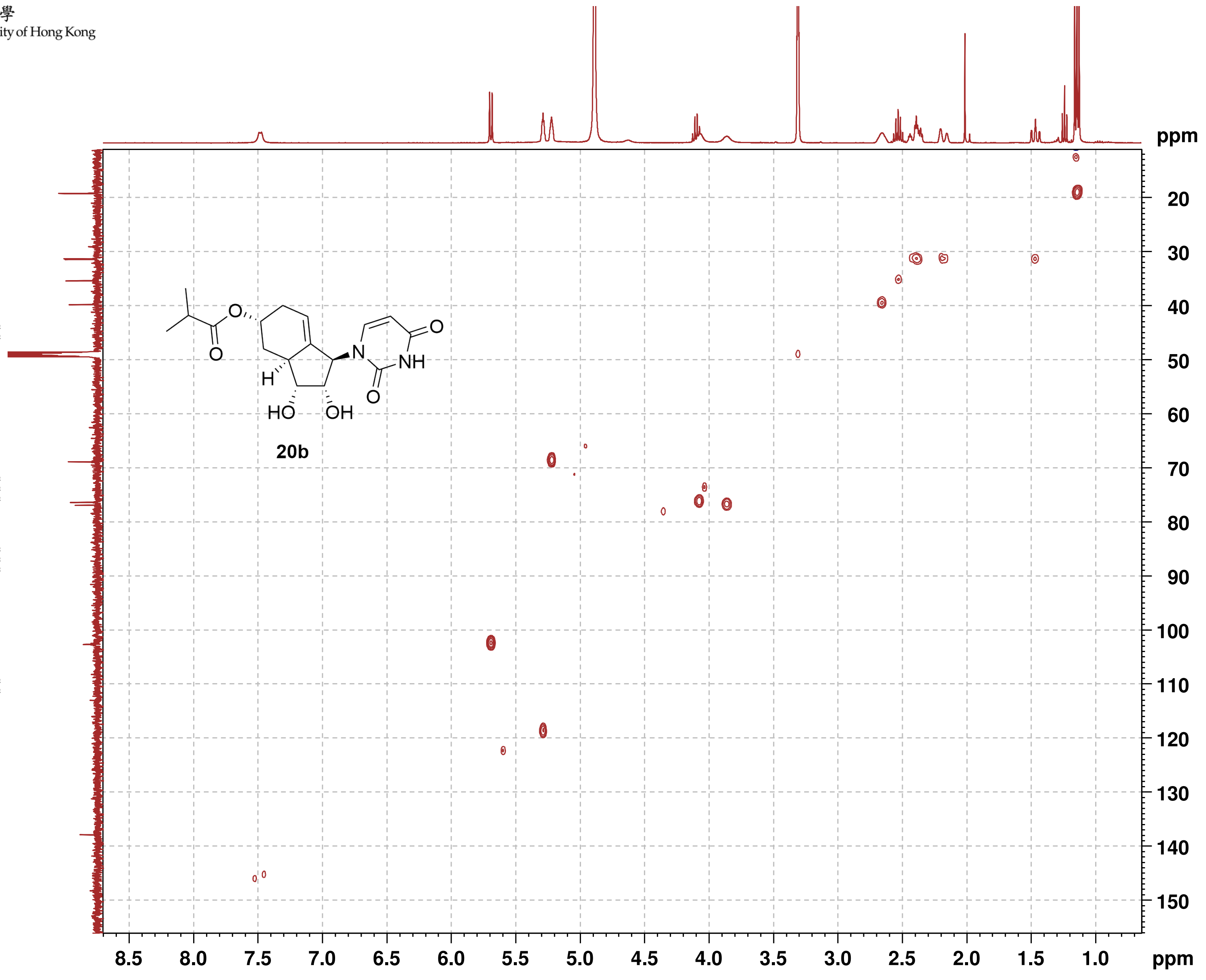

香港中文大學
The Chinese University of Hong Kong

**HMBC**

```
Current Data Parameters
NAME          SCC227_600
EXPNO                  5
PROCNO                 1

F2 - Acquisition Parameters
Date_           20220906
Time               16.34 h
INSTRUM            spect
PROBHD   Z816801_0148 (
PULPROG     hmbcgplpndqf
TD                  2048
SOLVENT             MeOD
NS                     4
DS                    16
SWH             6009.615 Hz
FIDRES          5.868765 Hz
AQ             0.1703936 sec
RG                189.92
DW                83.200 usec
DE                  6.50 usec
TE                 315.0 K
CNST2        145.0000000
CNST13        10.0000000
D0            0.00000300 sec
D1            1.50000000 sec
D2            0.00344828 sec
D6            0.05000000 sec
D16           0.00020000 sec
IN0           0.00001510 sec
TDav                   1
SFO1         600.1127005 MHz
NUC1                  1H
P1                  8.00 usec
P2                 16.00 usec
PLW1         14.00000000 W
SFO2         150.9128693 MHz
NUC2                 13C
P3                 12.00 usec
PLW2        183.00000000 W
GPNAM[1]      SMSQ10.100
GPZ1               50.00 %
GPNAM[2]      SMSQ10.100
GPZ2               30.00 %
GPNAM[3]      SMSQ10.100
GPZ3               40.10 %
P16              1000.00 usec

F1 - Acquisition parameters
TD                   128
SFO1            150.9129 MHz
FIDRES        517.384094 Hz
SW               219.415 ppm
FnMODE                QF

F2 - Processing parameters
SI                  2048
SF           600.1100070 MHz
WDW                 SINE
SSB                    0
LB                     0 Hz
GB                     0
PC                  1.40

F1 - Processing parameters
SI                  1024
MC2                   QF
SF           150.8975446 MHz
WDW                 SINE
SSB                    0
LB                     0 Hz
GB                     0
```

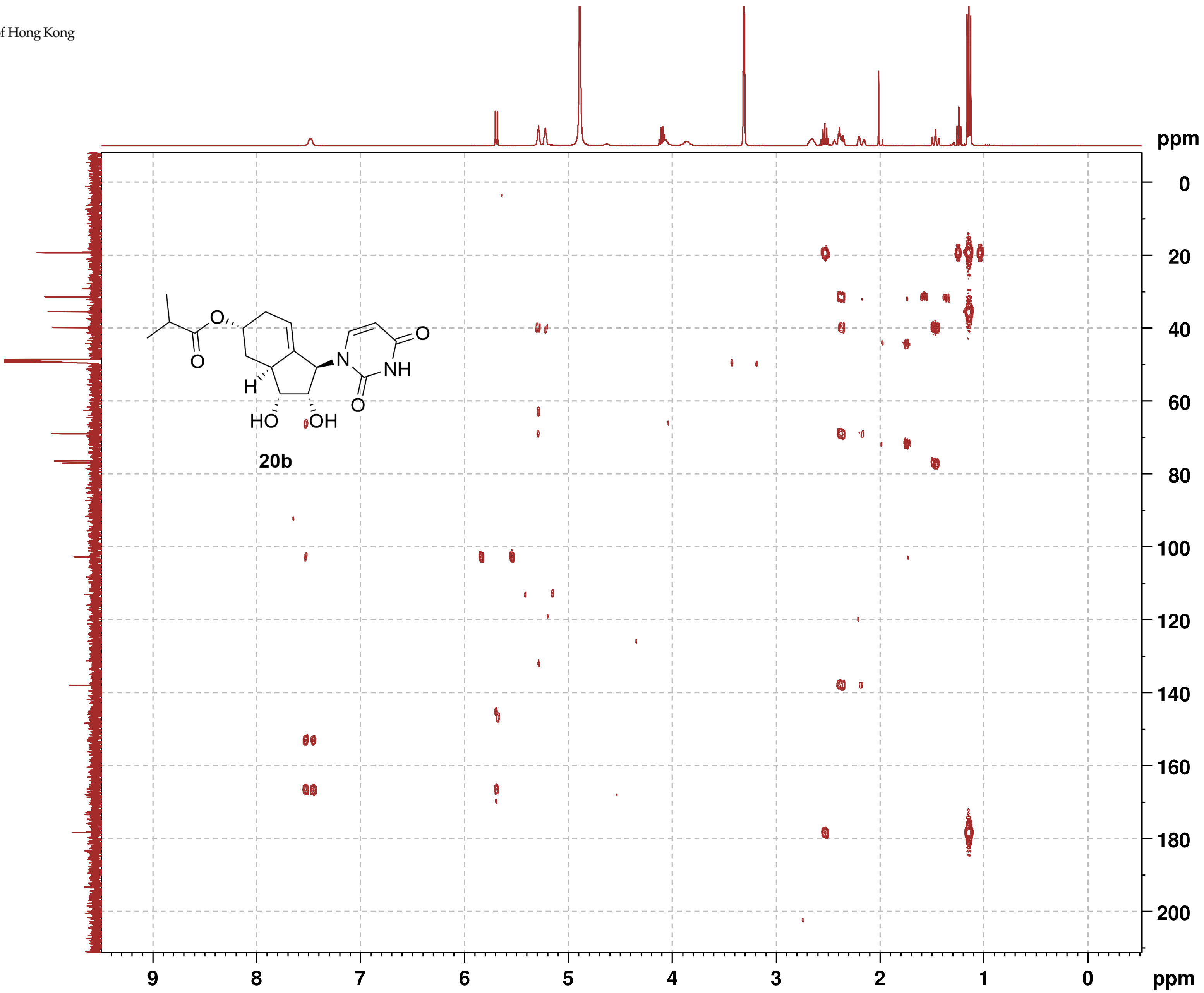

香港中文大學
The Chinese University of Hong Kong

#### NOESY

```
Current Data Parameters
NAME          SCC132_400
EXPNO                  4
PROCNO                 1

F2 - Acquisition Parameters
Date_           20220507
Time                8.13 h
INSTRUM            spect
PROBHD     Z108618_0257 (
PULPROG      noesygpphpp
TD                  2048
SOLVENT             MeOD
NS                     2
DS                    16
SWH             6009.615 Hz
FIDRES          5.868765 Hz
AQ             0.1703936 sec
RG                   203
DW                83.200 usec
DE                  6.50 usec
TE                 295.1 K
D0            0.00006420 sec
D1            2.00000000 sec
D8            0.60000002 sec
D11           0.03000000 sec
D12           0.00002000 sec
D16           0.00020000 sec
IN0           0.00016660 sec
TDav                   1
SFO1         400.1320007 MHz
NUC1                  1H
P1                 15.00 usec
P2                 30.00 usec
P17              2500.00 usec
PLW1         12.00000000 W
PLW10         3.00000000 W
GPNAM[1]       SMSQ10.100
GPZ1               40.00 %
P16              1000.00 usec

F1 - Acquisition parameters
TD                   256
SFO1             400.132 MHz
FIDRES         46.893757 Hz
SW                15.001 ppm
FnMODE        States-TPPI

F2 - Processing parameters
SI                  1024
SF           400.1300073 MHz
WDW                QSINE
SSB                    2
LB                     0 Hz
GB                     0
PC                  1.00

F1 - Processing parameters
SI                  1024
MC2          States-TPPI
SF           400.1300070 MHz
WDW                QSINE
SSB                    2
LB                     0 Hz
GB                     0
```

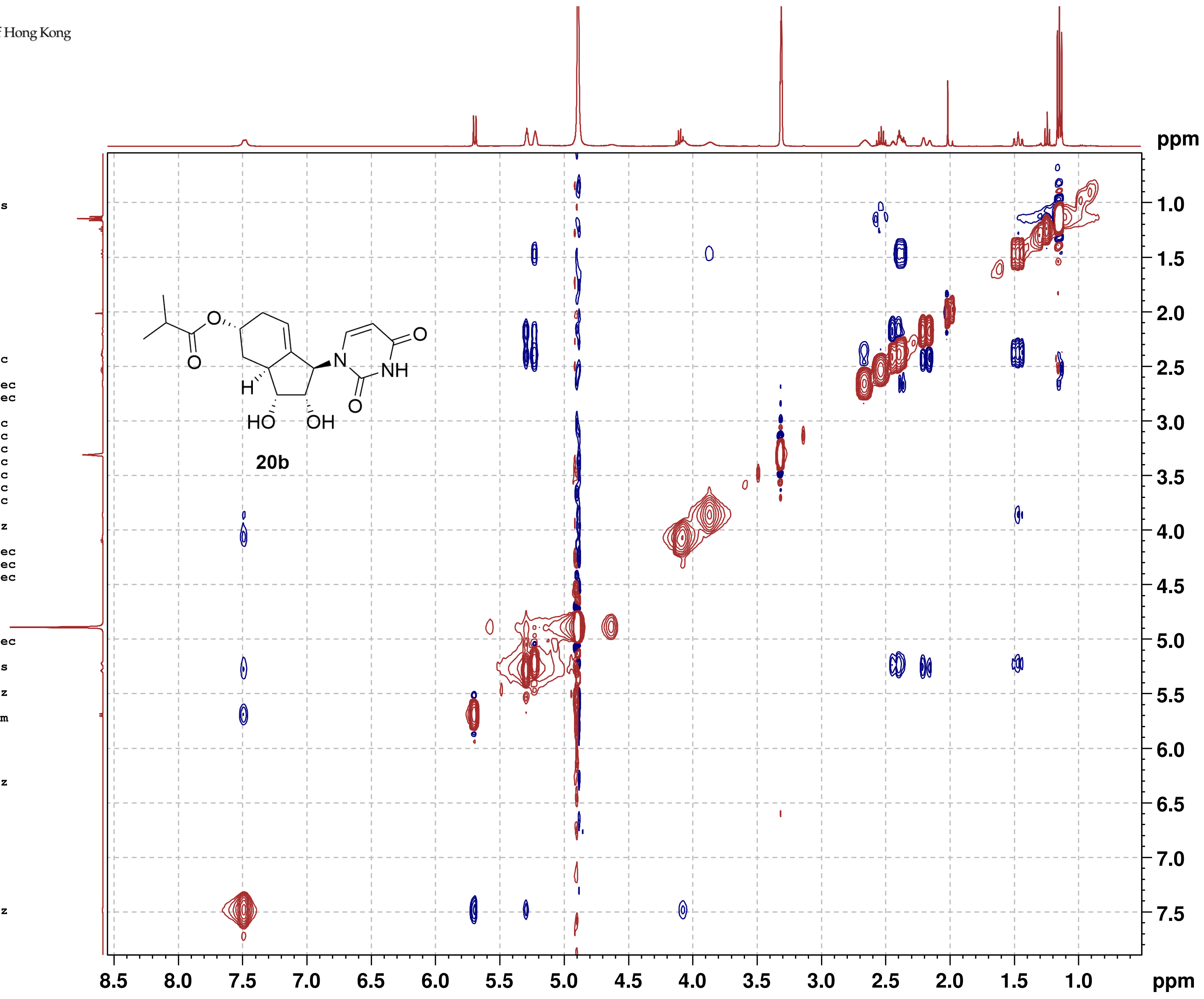

**20b**

#### NMR-Spectra for Compound 21b

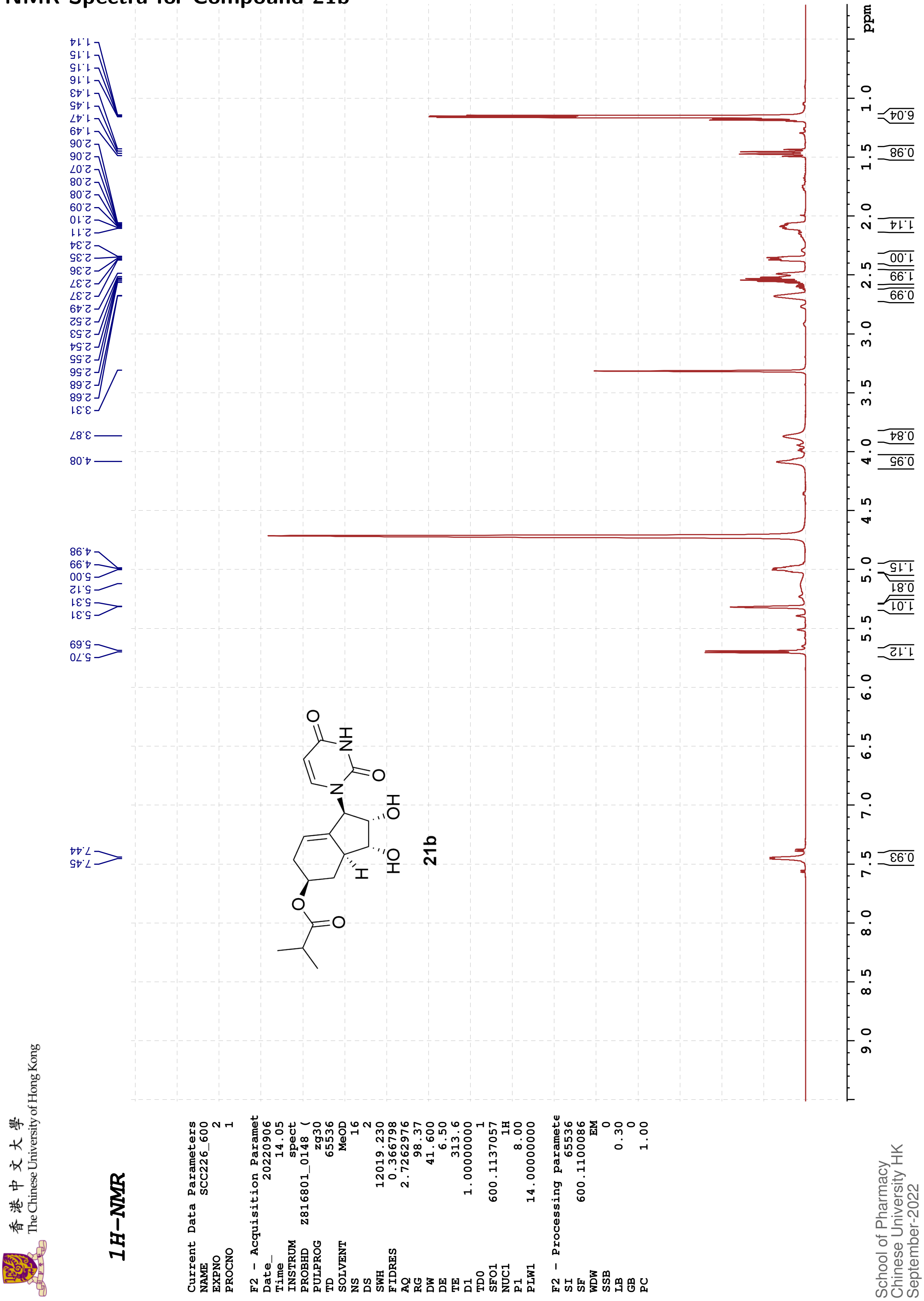

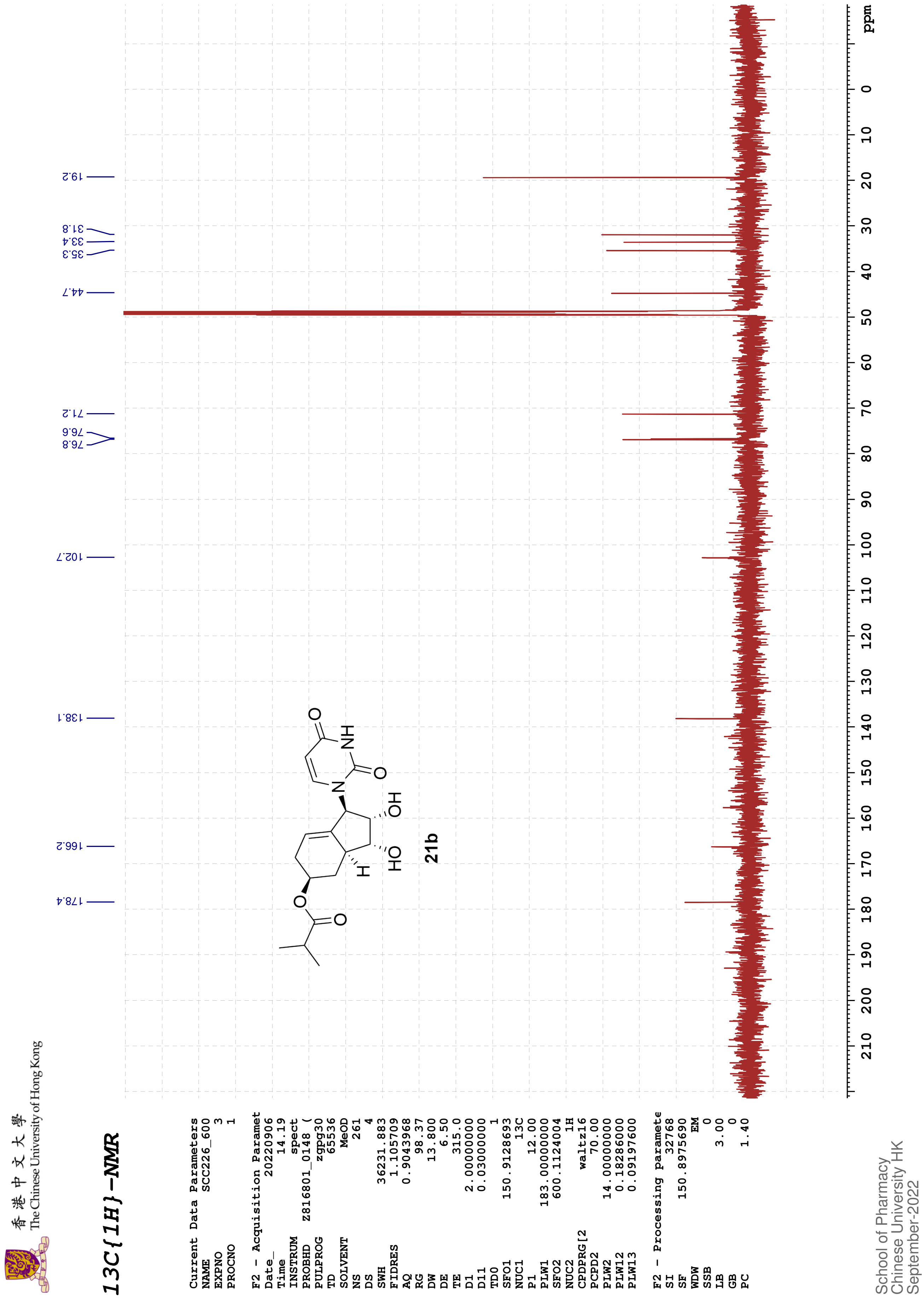
13C{1H}-NMR
The Chinese University of Hong Kong
178.4
166.2
138.1
102.7
76.8
76.6
71.2
44.7
35.3
33.4
31.8
19.2
21b
ppm
Current Data Parameters
NAME SCC226_600
EXPNO 3
PROCNO 1
F2 - Acquisition Paramet
Date_ 20220906
Time 14.19
INSTRUM spect
PROBHD Z816801_0148 (
PULPROG zgpg30
TD 65536
SOLVENT MeOD
NS 261
DS 4
SWH 36231.883
FIDRES 1.105709
AQ 0.9043968
RG 98.37
DW 13.800
DE 6.50
TE 315.0
D1 2.00000000
D11 0.03000000
TD0 1
SFO1 150.9128693
NUC1 13C
P1 12.00
PLW1 183.00000000
SFO2 600.1124004
NUC2 1H
CPDPRG[2 waltz16
PCPD2 70.00
PLW2 14.00000000
PLW12 0.18286000
PLW13 0.09197600
F2 - Processing paramete
SI 32768
SF 150.8975690
WDW EM
SSB 0
LB 3.00
GB 0
PC 1.40
School of Pharmacy
Chinese University HK
September-2022

香港中文大學
The Chinese University of Hong Kong

COSY

```
Current Data Parameters
NAME          SCC133_400
EXPNO                  4
PROCNO                 1

F2 - Acquisition Parameters
Date_           20220511
Time                7.55 h
INSTRUM            spect
PROBHD   Z108618_0257 (
PULPROG          cosygpqf
TD                  2048
SOLVENT             MeOD
NS                     2
DS                     8
SWH             6009.615 Hz
FIDRES          5.868765 Hz
AQ             0.1703936 sec
RG                   203
DW                83.200 usec
DE                  6.50 usec
TE                 295.3 K
D0            0.00000300 sec
D1            1.48689198 sec
D13           0.00000400 sec
D16           0.00020000 sec
IN0           0.00016660 sec
TDav                   1
SFO1         400.1322007 MHz
NUC1                  1H
P0                 15.00 usec
P1                 15.00 usec
PLW1         12.00000000 W
GPNAM[1]       SINE.100
GPZ1               10.00 %
P16              1000.00 usec

F1 - Acquisition parameters
TD                   128
SFO1            400.1322 MHz
FIDRES         93.787514 Hz
SW                15.001 ppm
FnMODE                QF

F2 - Processing parameters
SI                  1024
SF           400.1300074 MHz
WDW                 SINE
SSB                    0
LB                     0 Hz
GB                     0
PC                  1.40

F1 - Processing parameters
SI                  1024
MC2                   QF
SF           400.1300081 MHz
WDW                 SINE
SSB                    0
LB                     0 Hz
GB                     0
```

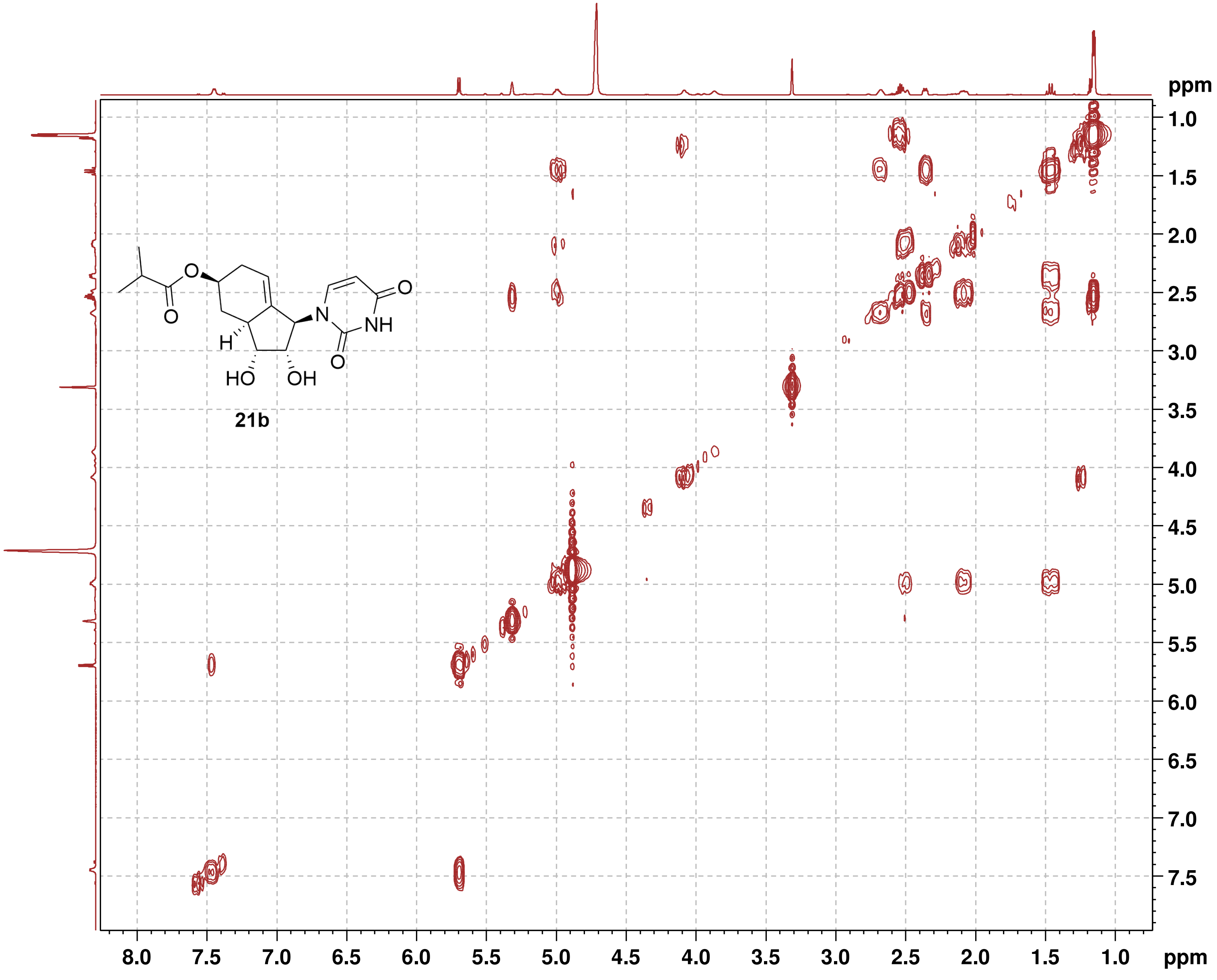

21b

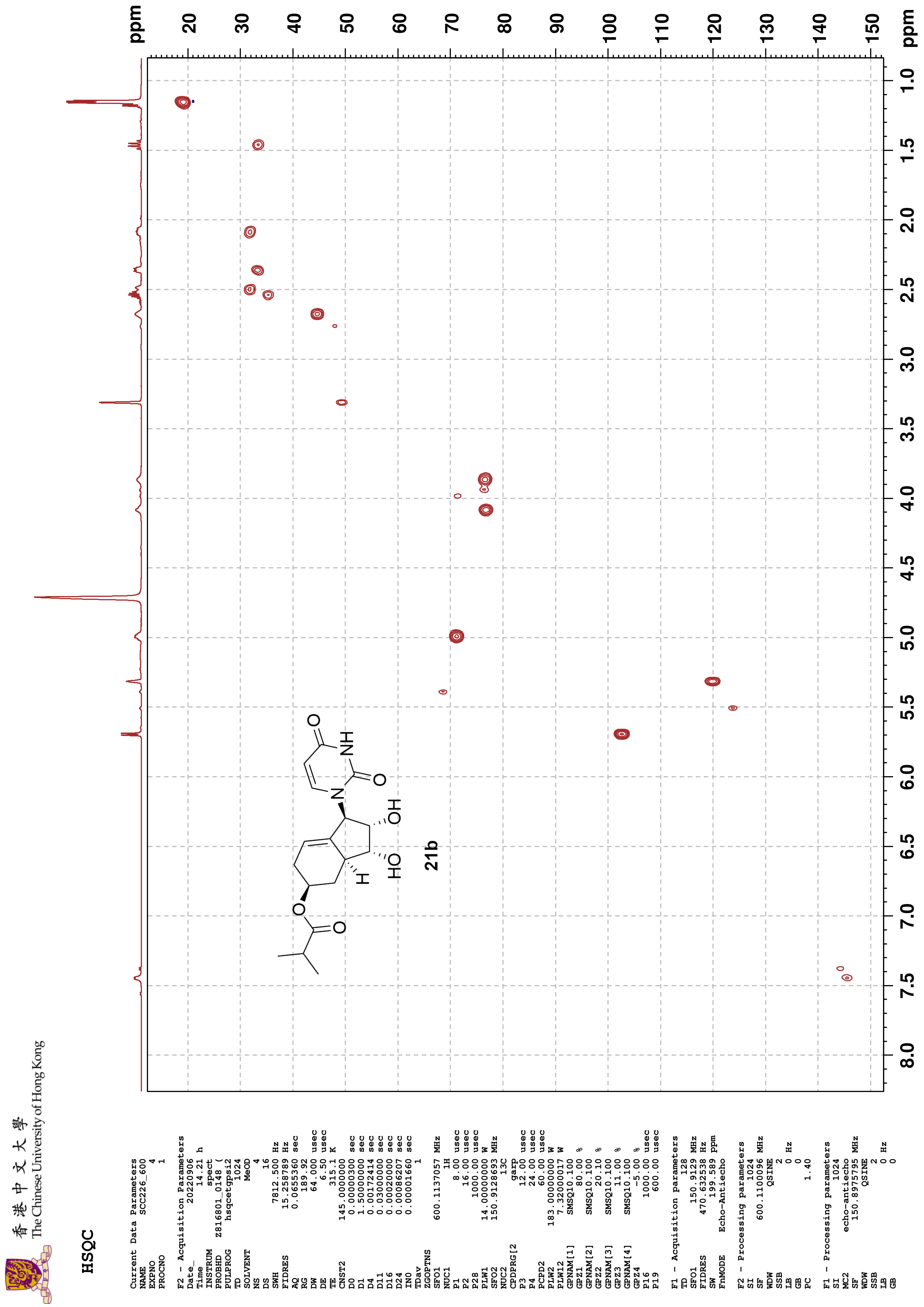

香港中文大學
The Chinese University of Hong Kong
HSQC
ppm
20
30
40
50
60
70
80
90
100
110
120
130
140
150
ppm
1.0
1.5
2.0
2.5
3.0
3.5
4.0
4.5
5.0
5.5
6.0
6.5
7.0
7.5
8.0
O
NH
N
O
OH
HO
H
O
O
21b
Current Data Parameters
NAME SCC226_600
EXPNO 4
PROCNO 1
F2 - Acquisition Parameters
Date_ 20220906
Time 14.21 h
INSTRUM spect
PROBHD Z816801_0148 (
PULPROG hsqcetgpsi2
TD 1024
SOLVENT MeOD
NS 4
DS 16
SWH 7812.500 Hz
FIDRES 15.258789 Hz
AQ 0.0655360 sec
RG 189.92
DW 64.000 usec
DE 6.50 usec
TE 315.1 K
CNST2 145.0000000
D0 0.00000300 sec
D1 1.50000000 sec
D4 0.00172414 sec
D11 0.03000000 sec
D16 0.00020000 sec
D24 0.00086207 sec
IN0 0.00001660 sec
TDav 1
ZGOPTNS
SFO1 600.1137057 MHz
NUC1 1H
P1 8.00 usec
P2 16.00 usec
P28 1000.00 usec
PLW1 14.00000000 W
SFO2 150.9128693 MHz
NUC2 13C
CPDPRG[2 garp
P3 12.00 usec
P4 24.00 usec
PCPD2 60.00 usec
PLW2 183.00000000 W
PLW12 7.32000017 W
GPNAM[1] SMSQ10.100
GPZ1 80.00 %
GPNAM[2] SMSQ10.100
GPZ2 20.10 %
GPNAM[3] SMSQ10.100
GPZ3 11.00 %
GPNAM[4] SMSQ10.100
GPZ4 -5.00 %
P16 1000.00 usec
P19 600.00 usec
F1 - Acquisition parameters
TD 128
SFO1 150.9129 MHz
FIDRES 470.632538 Hz
SW 199.589 ppm
FnMODE Echo-Antiecho
F2 - Processing parameters
SI 1024
SF 600.1100096 MHz
WDW QSINE
SSB 2
LB 0 Hz
GB 0
PC 1.40
F1 - Processing parameters
SI 1024
MC2 echo-antiecho
SF 150.8975795 MHz
WDW QSINE
SSB 2
LB 0 Hz
GB 0

香港中文大學
The Chinese University of Hong Kong

**HMBC**

```
Current Data Parameters
NAME          SCC226_600
EXPNO                  5
PROCNO                 1

F2 - Acquisition Parameters
Date_           20220906
Time               14.35 h
INSTRUM            spect
PROBHD   Z816801_0148 (
PULPROG    hmbcgplpndqf
TD                  2048
SOLVENT             MeOD
NS                     5
DS                    16
SWH             6009.615 Hz
FIDRES          5.868765 Hz
AQ             0.1703936 sec
RG                189.92
DW                83.200 usec
DE                  6.50 usec
TE                 314.9 K
CNST2        145.0000000
CNST13        10.0000000
D0            0.00000300 sec
D1            1.50000000 sec
D2            0.00344828 sec
D6            0.05000000 sec
D16           0.00020000 sec
IN0           0.00001510 sec
TDav                   1
SFO1         600.1127005 MHz
NUC1                  1H
P1                  8.00 usec
P2                 16.00 usec
PLW1         14.00000000 W
SFO2         150.9128693 MHz
NUC2                 13C
P3                 12.00 usec
PLW2        183.00000000 W
GPNAM[1]      SMSQ10.100
GPZ1               50.00 %
GPNAM[2]      SMSQ10.100
GPZ2               30.00 %
GPNAM[3]      SMSQ10.100
GPZ3               40.10 %
P16              1000.00 usec

F1 - Acquisition parameters
TD                   128
SFO1            150.9129 MHz
FIDRES        517.384094 Hz
SW               219.415 ppm
FnMODE                QF

F2 - Processing parameters
SI                  2048
SF           600.1100032 MHz
WDW                 SINE
SSB                    0
LB                     0 Hz
GB                     0
PC                  1.40

F1 - Processing parameters
SI                  1024
MC2                   QF
SF           150.8975846 MHz
WDW                 SINE
SSB                    0
LB                     0 Hz
GB                     0
```

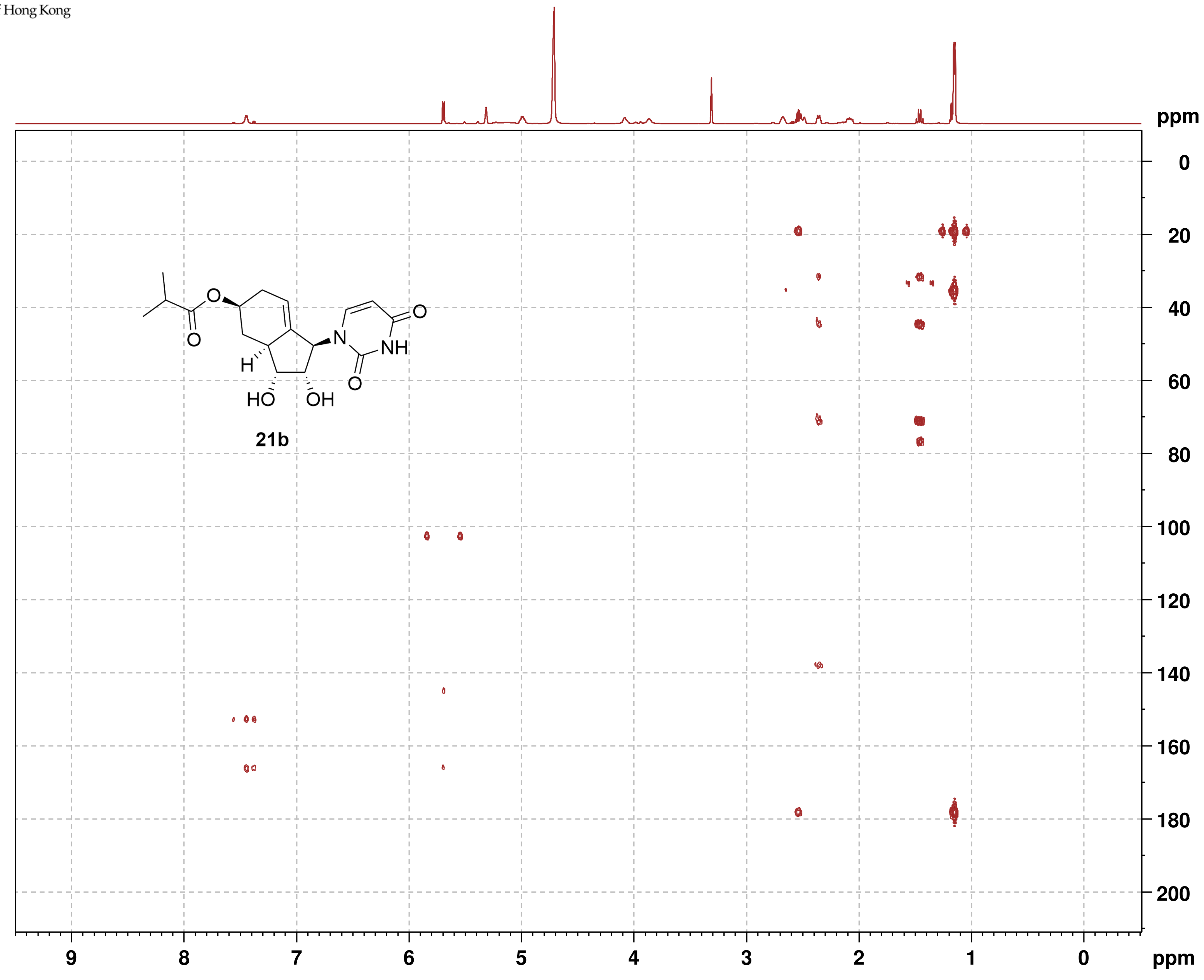

香港中文大學
The Chinese University of Hong Kong

**NOESY**

```
Current Data Parameters
NAME          SCC133_400
EXPNO                  5
PROCNO                 1

F2 - Acquisition Parameters
Date_           20220511
Time                8.04 h
INSTRUM            spect
PROBHD    Z108618_0257 (
PULPROG      noesygpphpp
TD                  2048
SOLVENT             MeOD
NS                     2
DS                    16
SWH             6009.615 Hz
FIDRES          5.868765 Hz
AQ             0.1703936 sec
RG                   203
DW                83.200 usec
DE                  6.50 usec
TE                 295.2 K
D0            0.00006420 sec
D1            2.00000000 sec
D8            0.60000002 sec
D11           0.03000000 sec
D12           0.00002000 sec
D16           0.00020000 sec
IN0           0.00016660 sec
TDav                   1
SFO1         400.1320007 MHz
NUC1                  1H
P1                 15.00 usec
P2                 30.00 usec
P17              2500.00 usec
PLW1         12.00000000 W
PLW10         3.00000000 W
GPNAM[1]        SMSQ10.100
GPZ1               40.00 %
P16              1000.00 usec

F1 - Acquisition parameters
TD                   256
SFO1             400.132 MHz
FIDRES         46.893757 Hz
SW                15.001 ppm
FnMODE       States-TPPI

F2 - Processing parameters
SI                  1024
SF           400.1300075 MHz
WDW                QSINE
SSB                    2
LB                     0 Hz
GB                     0
PC                  1.00

F1 - Processing parameters
SI                  1024
MC2          States-TPPI
SF           400.1300053 MHz
WDW                QSINE
SSB                    2
LB                     0 Hz
GB                     0
```

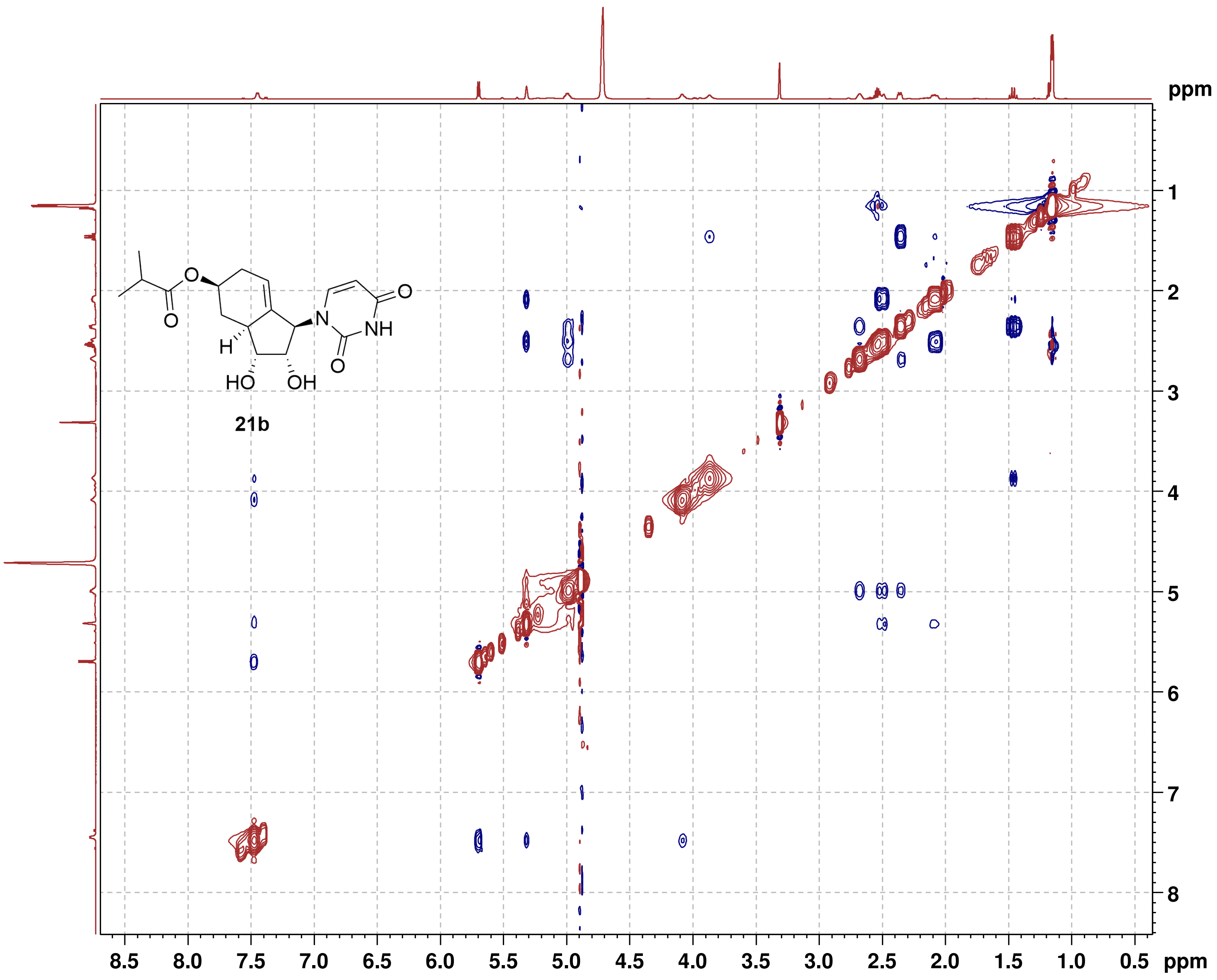

#### NMR-Spectra for Compound 21b

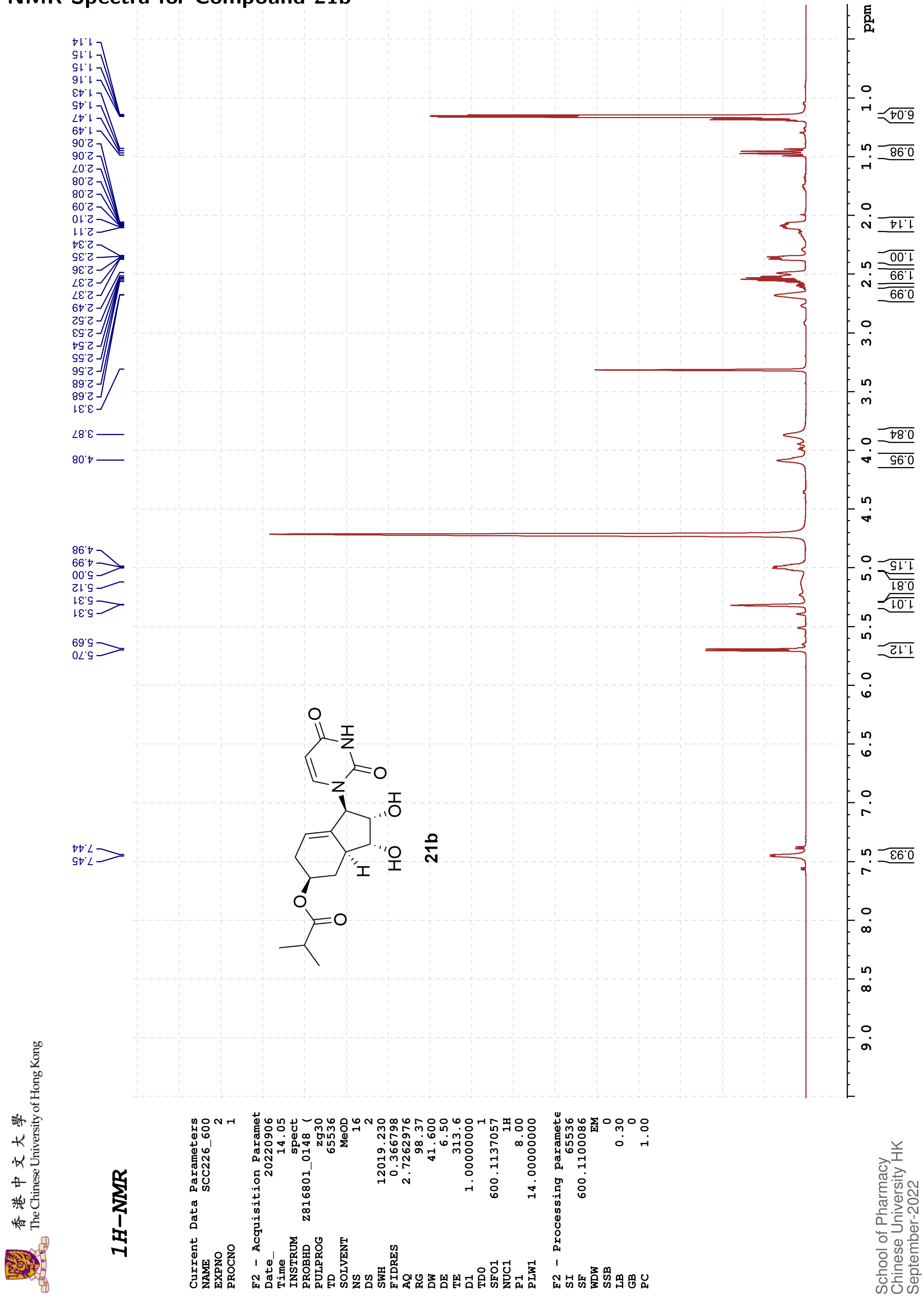

香港中文大學 The Chinese University of Hong Kong

#### 13C{1H}-NMR

```
Current Data Parameters
NAME            SCC226_600
EXPNO                    3
PROCNO                   1

F2 - Acquisition Paramet
Date_             20220906
Time                 14.19
INSTRUM              spect
PROBHD     Z816801_0148 (
PULPROG             zgpg30
TD                   65536
SOLVENT               MeOD
NS                     261
DS                       4
SWH              36231.883
FIDRES            1.105709
AQ               0.9043968
RG                   98.37
DW                  13.800
DE                    6.50
TE                   315.0
D1              2.00000000
D11             0.03000000
TD0                      1
SFO1           150.9128693
NUC1                   13C
P1                   12.00
PLW1          183.00000000
SFO2           600.1124004
NUC2                    1H
CPDPRG[2           waltz16
PCPD2                70.00
PLW2           14.00000000
PLW12           0.18286000
PLW13           0.09197600

F2 - Processing paramete
SI                   32768
SF             150.8975690
WDW                     EM
SSB                      0
LB                    3.00
GB                       0
PC                    1.40
```

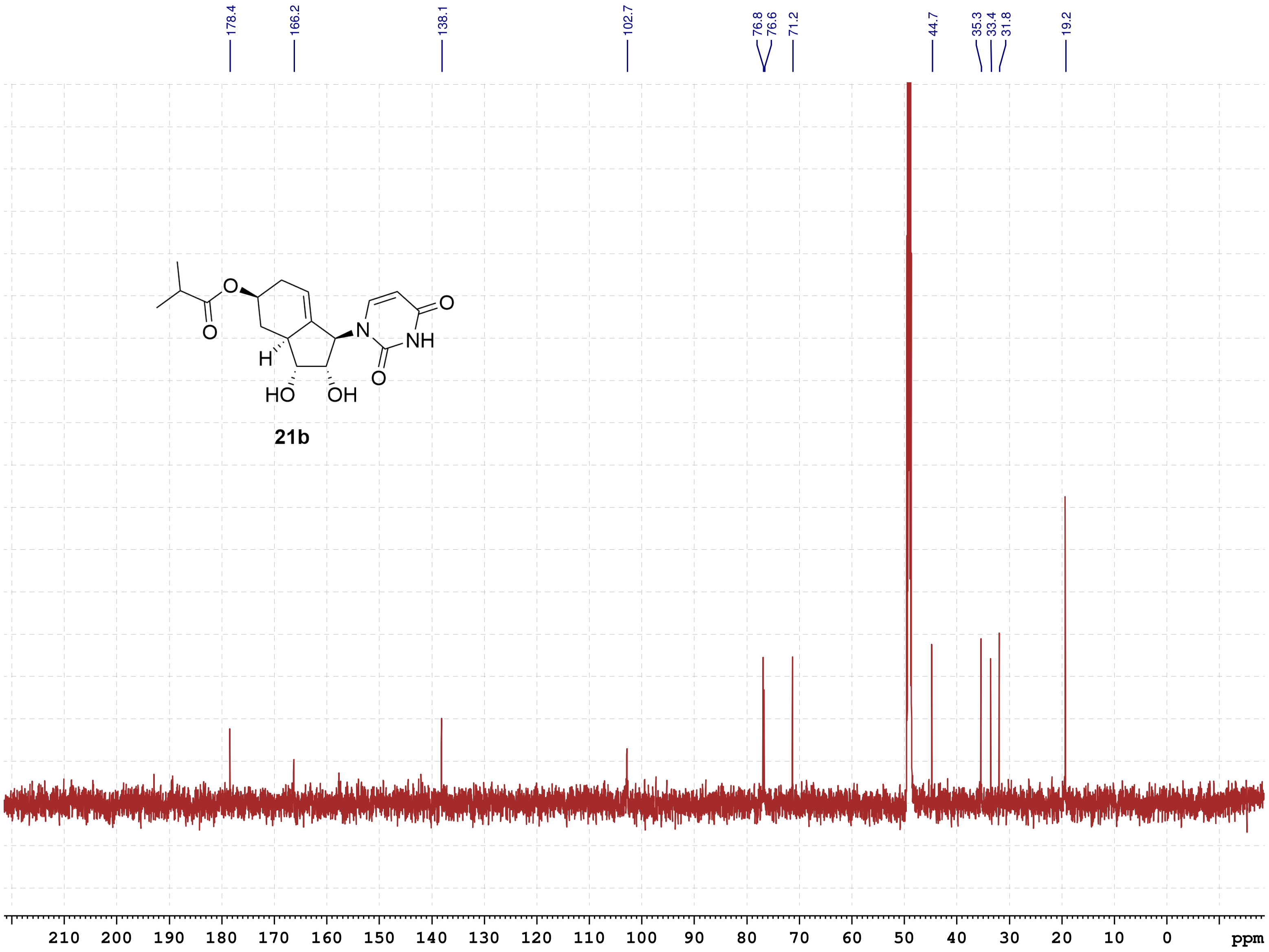

**21b**

School of Pharmacy
Chinese University HK
September-2022

香港中文大學
The Chinese University of Hong Kong

COSY

Current Data Parameters
NAME SCC133_400
EXPNO 4
PROCNO 1

F2 - Acquisition Parameters
Date_ 20220511
Time 7.55 h
INSTRUM spect
PROBHD Z108618_0257 (
PULPROG cosygpqf
TD 2048
SOLVENT MeOD
NS 2
DS 8
SWH 6009.615 Hz
FIDRES 5.868765 Hz
AQ 0.1703936 sec
RG 203
DW 83.200 usec
DE 6.50 usec
TE 295.3 K
D0 0.00000300 sec
D1 1.48689198 sec
D13 0.00000400 sec
D16 0.00020000 sec
IN0 0.00016660 sec
TDav 1
SFO1 400.1322007 MHz
NUC1 1H
P0 15.00 usec
P1 15.00 usec
PLW1 12.00000000 W
GPNAM[1] SINE.100
GPZ1 10.00 %
P16 1000.00 usec

F1 - Acquisition parameters
TD 128
SFO1 400.1322 MHz
FIDRES 93.787514 Hz
SW 15.001 ppm
FnMODE QF

F2 - Processing parameters
SI 1024
SF 400.1300074 MHz
WDW SINE
SSB 0
LB 0 Hz
GB 0
PC 1.40

F1 - Processing parameters
SI 1024
MC2 QF
SF 400.1300081 MHz
WDW SINE
SSB 0
LB 0 Hz
GB 0

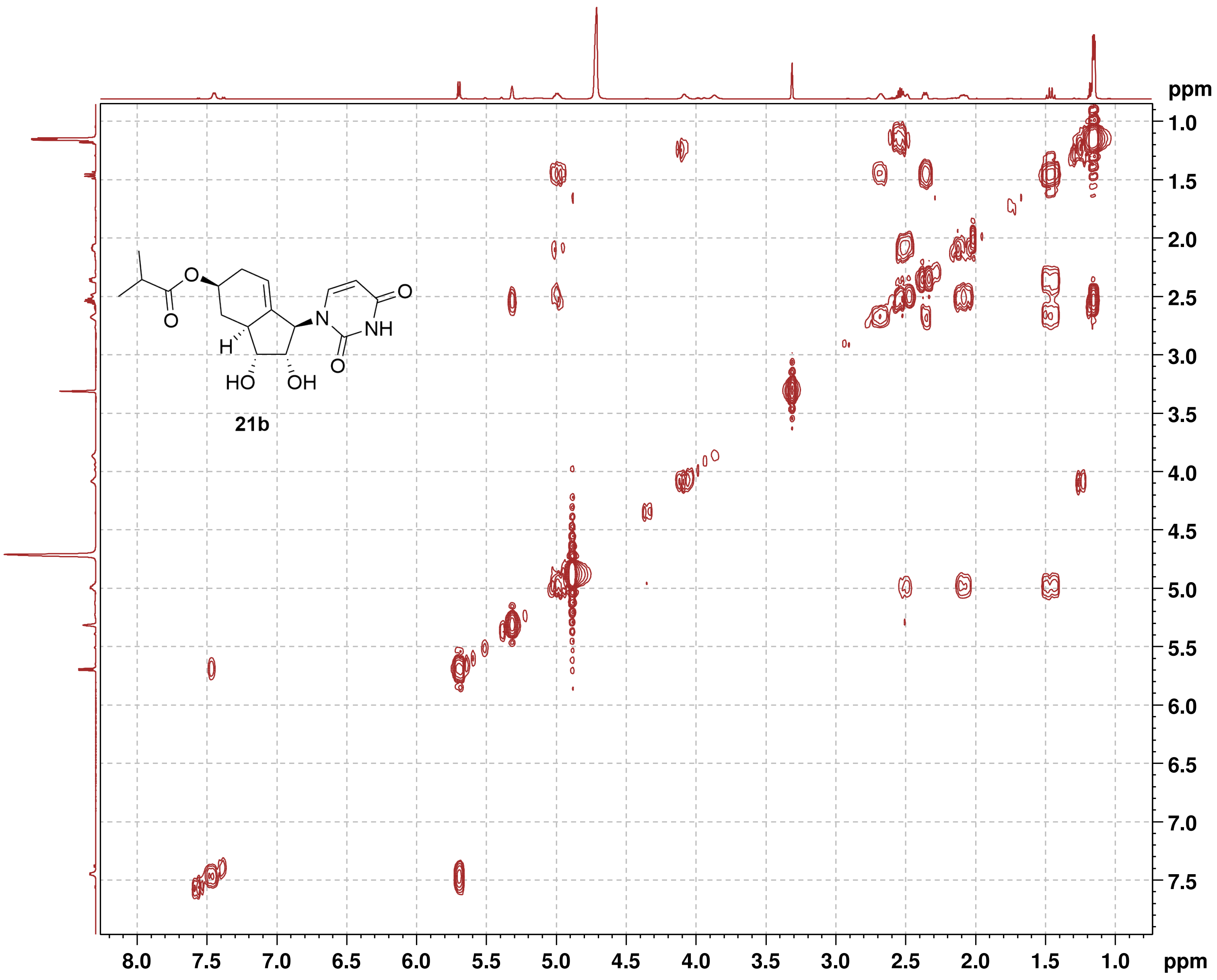

21b

香 港 中 文 大 學
The Chinese University of Hong Kong

**HSQC**

```
Current Data Parameters
NAME          SCC226_600
EXPNO                  4
PROCNO                 1

F2 - Acquisition Parameters
Date_           20220906
Time               14.21 h
INSTRUM            spect
PROBHD   Z816801_0148 (
PULPROG      hsqcetgpsi2
TD                  1024
SOLVENT             MeOD
NS                     4
DS                    16
SWH             7812.500 Hz
FIDRES         15.258789 Hz
AQ             0.0655360 sec
RG                189.92
DW                64.000 usec
DE                  6.50 usec
TE                 315.1 K
CNST2        145.0000000
D0            0.00000300 sec
D1            1.50000000 sec
D4            0.00172414 sec
D11           0.03000000 sec
D16           0.00020000 sec
D24           0.00086207 sec
IN0           0.00001660 sec
TDav                   1
ZGOPTNS
SFO1         600.1137057 MHz
NUC1                  1H
P1                  8.00 usec
P2                 16.00 usec
P28              1000.00 usec
PLW1         14.00000000 W
SFO2         150.9128693 MHz
NUC2                 13C
CPDPRG[2            garp
P3                 12.00 usec
P4                 24.00 usec
PCPD2              60.00 usec
PLW2        183.00000000 W
PLW12         7.32000017 W
GPNAM[1]      SMSQ10.100
GPZ1               80.00 %
GPNAM[2]      SMSQ10.100
GPZ2               20.10 %
GPNAM[3]      SMSQ10.100
GPZ3               11.00 %
GPNAM[4]      SMSQ10.100
GPZ4               -5.00 %
P16              1000.00 usec
P19               600.00 usec

F1 - Acquisition parameters
TD                   128
SFO1            150.9129 MHz
FIDRES        470.632538 Hz
SW               199.589 ppm
FnMODE      Echo-Antiecho

F2 - Processing parameters
SI                  1024
SF           600.1100096 MHz
WDW                QSINE
SSB                    2
LB                     0 Hz
GB                     0
PC                  1.40

F1 - Processing parameters
SI                  1024
MC2         echo-antiecho
SF           150.8975795 MHz
WDW                QSINE
SSB                    2
LB                     0 Hz
GB                     0
```

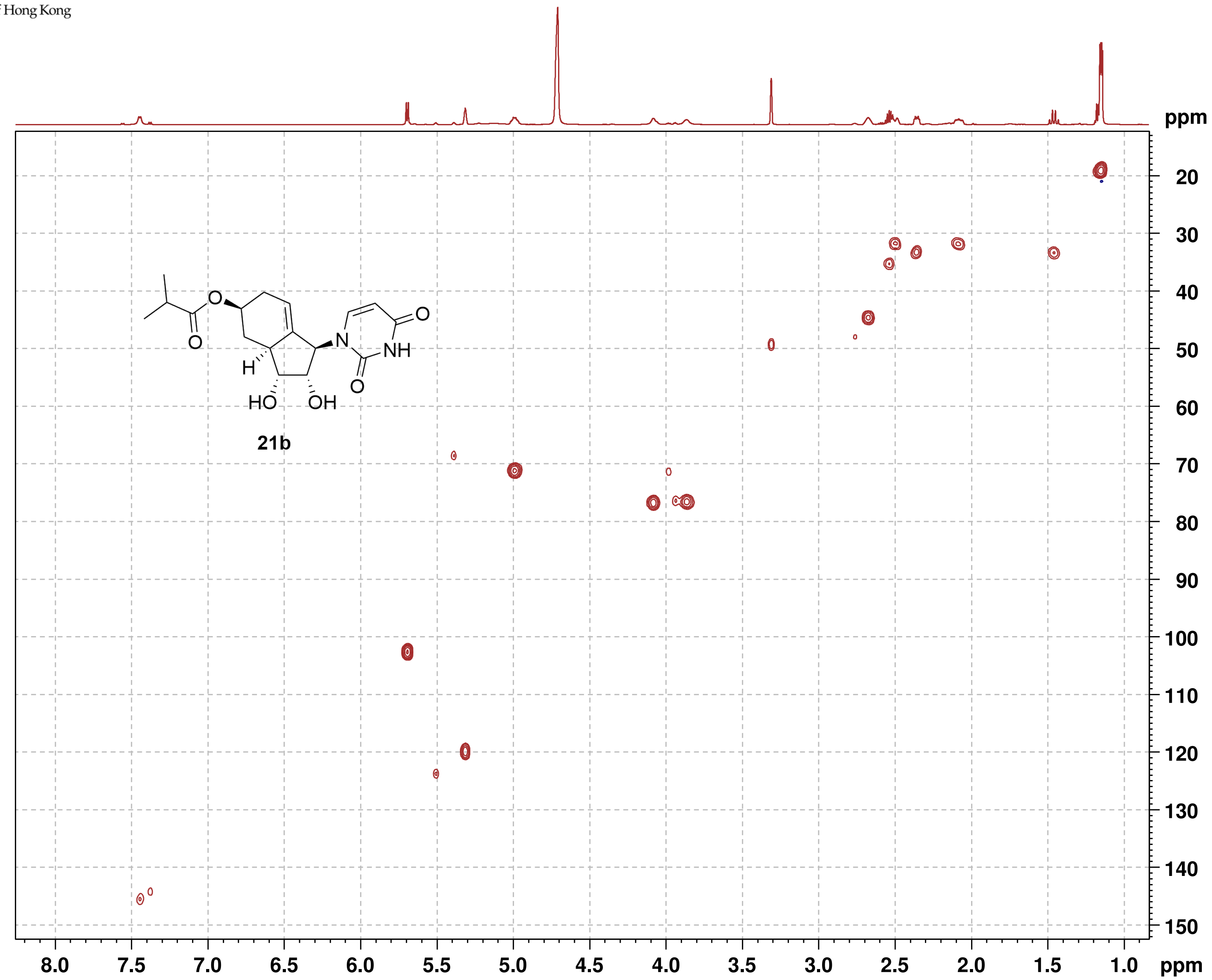

The Chinese University of Hong Kong 香港中文大學

**HMBC**

```
Current Data Parameters
NAME          SCC226_600
EXPNO                  5
PROCNO                 1

F2 - Acquisition Parameters
Date_          20220906
Time              14.35 h
INSTRUM           spect
PROBHD   Z816801_0148 (
PULPROG    hmbcgplpndqf
TD                 2048
SOLVENT            MeOD
NS                    5
DS                   16
SWH            6009.615 Hz
FIDRES         5.868765 Hz
AQ            0.1703936 sec
RG               189.92
DW               83.200 usec
DE                 6.50 usec
TE                314.9 K
CNST2       145.0000000
CNST13       10.0000000
D0           0.00000300 sec
D1           1.50000000 sec
D2           0.00344828 sec
D6           0.05000000 sec
D16          0.00020000 sec
IN0          0.00001510 sec
TDav                  1
SFO1        600.1127005 MHz
NUC1                 1H
P1                 8.00 usec
P2                16.00 usec
PLW1        14.00000000 W
SFO2        150.9128693 MHz
NUC2                13C
P3                12.00 usec
PLW2       183.00000000 W
GPNAM[1]     SMSQ10.100
GPZ1              50.00 %
GPNAM[2]     SMSQ10.100
GPZ2              30.00 %
GPNAM[3]     SMSQ10.100
GPZ3              40.10 %
P16             1000.00 usec

F1 - Acquisition parameters
TD                  128
SFO1           150.9129 MHz
FIDRES       517.384094 Hz
SW              219.415 ppm
FnMODE               QF

F2 - Processing parameters
SI                 2048
SF          600.1100032 MHz
WDW                SINE
SSB                   0
LB                    0 Hz
GB                    0
PC                 1.40

F1 - Processing parameters
SI                 1024
MC2                  QF
SF          150.8975846 MHz
WDW                SINE
SSB                   0
LB                    0 Hz
GB                    0
```

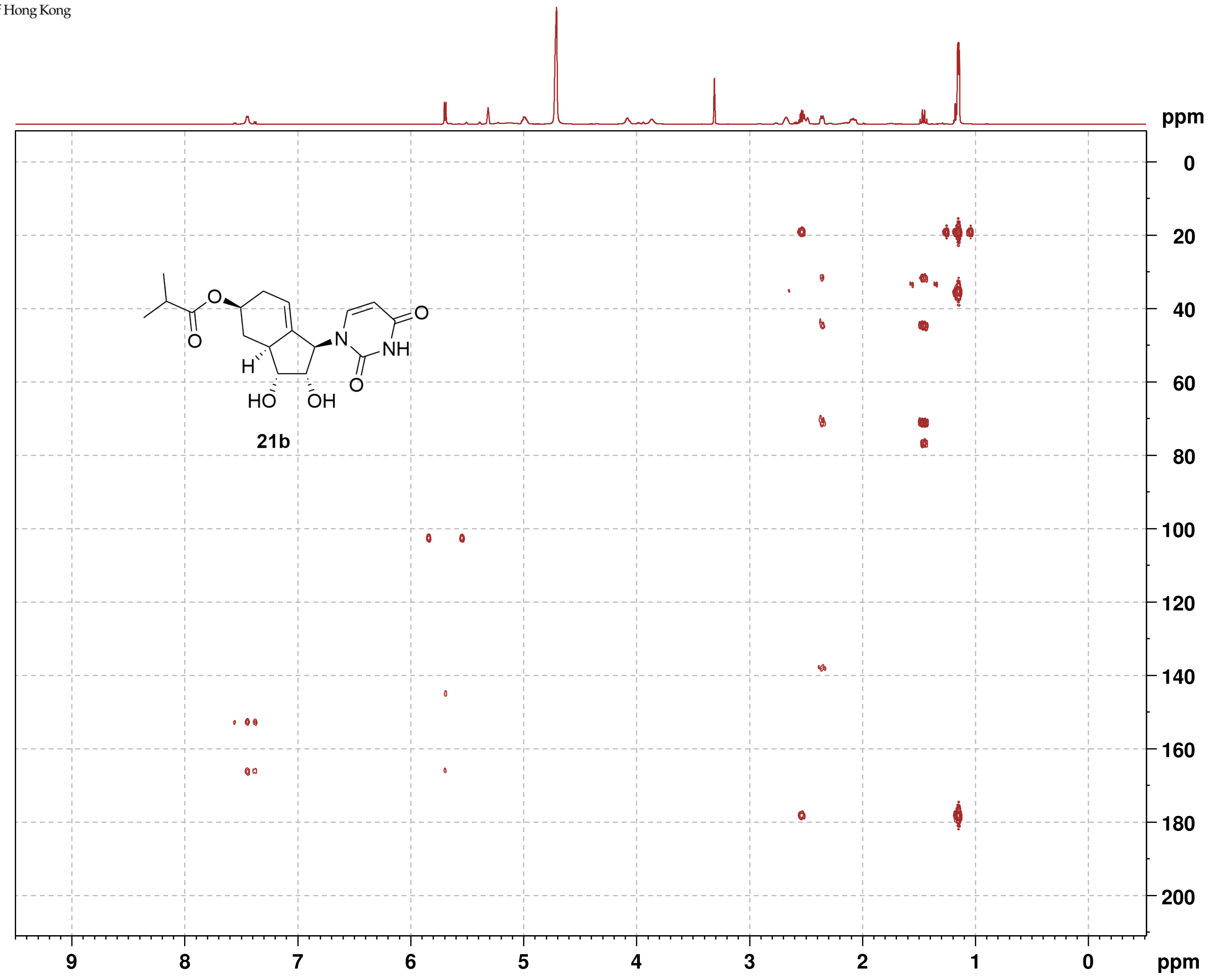

香港中文大學
The Chinese University of Hong Kong

**NOESY**

```
Current Data Parameters
NAME          SCC133_400
EXPNO                  5
PROCNO                 1

F2 - Acquisition Parameters
Date_           20220511
Time                8.04 h
INSTRUM            spect
PROBHD    Z108618_0257 (
PULPROG      noesygpphpp
TD                  2048
SOLVENT             MeOD
NS                     2
DS                    16
SWH             6009.615 Hz
FIDRES          5.868765 Hz
AQ             0.1703936 sec
RG                   203
DW                83.200 usec
DE                  6.50 usec
TE                 295.2 K
D0            0.00006420 sec
D1            2.00000000 sec
D8            0.60000002 sec
D11           0.03000000 sec
D12           0.00002000 sec
D16           0.00020000 sec
IN0           0.00016660 sec
TDav                   1
SFO1         400.1320007 MHz
NUC1                  1H
P1                 15.00 usec
P2                 30.00 usec
P17              2500.00 usec
PLW1         12.00000000 W
PLW10         3.00000000 W
GPNAM[1]       SMSQ10.100
GPZ1               40.00 %
P16              1000.00 usec

F1 - Acquisition parameters
TD                   256
SFO1             400.132 MHz
FIDRES         46.893757 Hz
SW                15.001 ppm
FnMODE       States-TPPI

F2 - Processing parameters
SI                  1024
SF           400.1300075 MHz
WDW                QSINE
SSB                    2
LB                     0 Hz
GB                     0
PC                  1.00

F1 - Processing parameters
SI                  1024
MC2          States-TPPI
SF           400.1300053 MHz
WDW                QSINE
SSB                    2
LB                     0 Hz
GB                     0
```

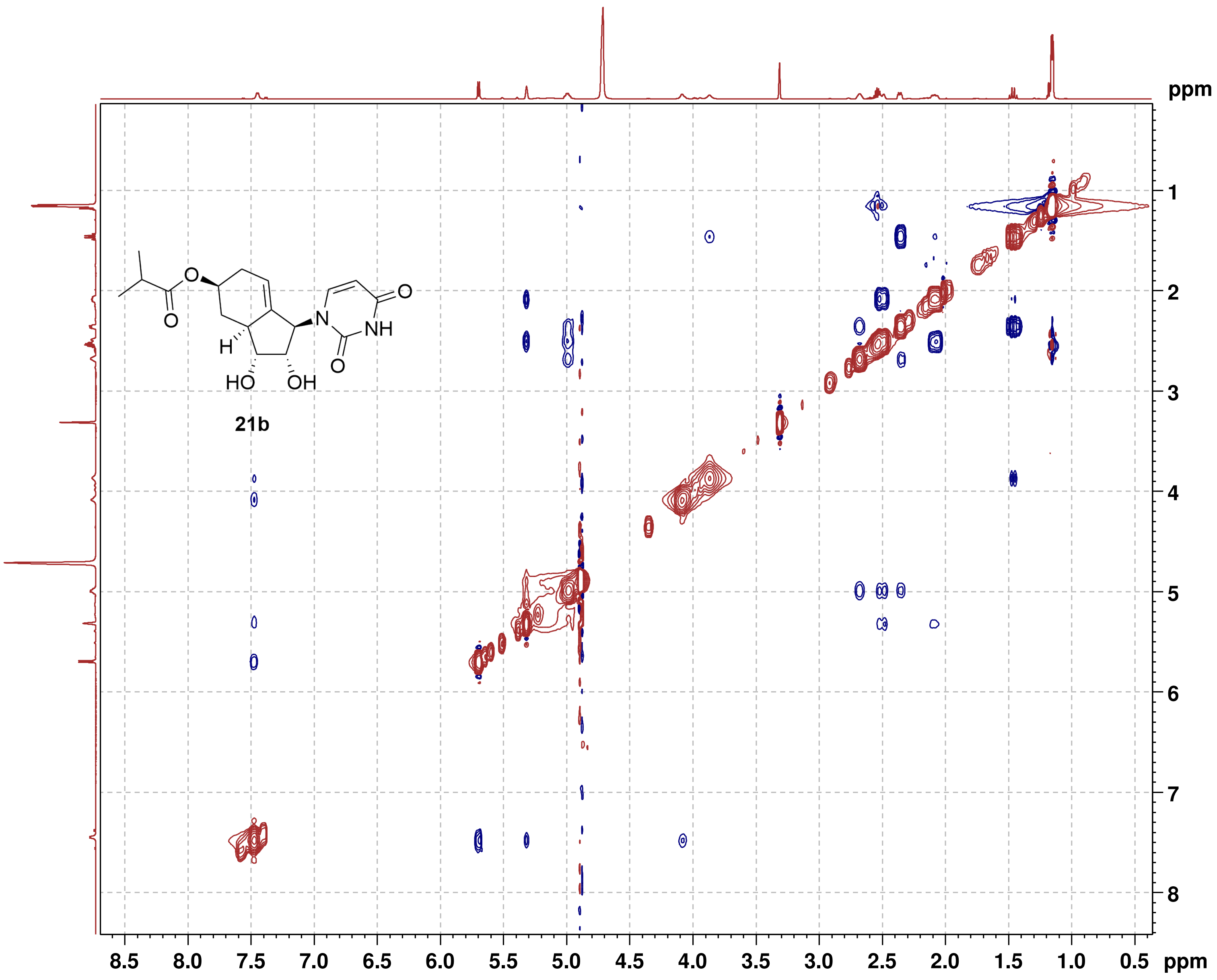
